# The structural landscape and diversity of *Pyricularia oryzae* MAX effectors revisited

**DOI:** 10.1101/2023.09.26.559520

**Authors:** Mounia Lahfa, Philippe Barthe, Karine De Guillen, Stella Cesari, Mouna Raji, Thomas Kroj, Marie Le Naour--Vernet, François Hoh, Pierre Gladieux, Christian Roumestand, Jérôme Gracy, Nathalie Declerck, André Padilla

## Abstract

*Magnaporthe* AVRs and ToxB-like (MAX) effectors constitute a family of secreted virulence proteins in the fungus *Pyricularia oryzae (syn. Magnaporthe oryzae)*, which causes blast disease on numerous cereals and grasses. In spite of high sequence divergence, MAX effectors share a common fold characterized by a ß-sandwich core stabilized by a conserved disulfide bond.

In this study, we investigated the structural landscape and diversity within the MAX effector repertoire of *P. oryzae.* Combining experimental protein structure determination and *in silico* structure modeling we validated the presence of the conserved MAX effector core domain in 77 out of 94 groups of orthologs (OG) identified in a previous population genomic study. Four novel MAX effector structures determined by NMR were in remarkably good agreement with AlphaFold2 (AF2) predictions. Based on the comparison of the AF2-generated 3D models we propose a classification of the MAX effectors superfamily in 20 structural groups that vary in the canonical MAX fold, disulfide bond patterns, and additional secondary structures in N- and C-terminal extensions. About one-third of the MAX family members remain singletons, without strong structural relationship to other MAX effectors. Analysis of the surface properties of the AF2 MAX models also highlights the high variability within the MAX family at the structural level, potentially reflecting the wide diversity of their virulence functions and host targets.

**Author summary:** MAX effectors are a family of virulence proteins from the plant pathogenic fungus *Pyricularia (syn. Magnaporthe) oryzae* that share a similar 3D structure despite very low amino-acid sequence identity. Characterizing the function and evolution of these proteins requires a detailed understanding of their structural diversity. With this in mind, we have determined the NMR structures of four new MAX effectors and shown a near-perfect match with the corresponding AlphaFold2 (AF2) models. We then applied a prediction pipeline based on similarity searches with structural modeling using the AF2 software to predict MAX effectors in a collection of 120 *P. oryzae* genomes. The resulting models and experimental structures revealed that the MAX core while preserved is highly permissive to secondary structure variations and may coexists with extensive structural diversity in terms of structured N- or C-terminal extensions permitting their classification. For a subset of AF2 models, we have also analyzed the physico-chemical properties of the core domain surfaces, adding another, more functional perspective, notably surface electrostatics and stickiness. This work constitutes a major step in understanding the relationships among MAX effectors by analyzing their structural landscape and cataloguing specific physico-chemical properties. It also provides valuable insights for guiding research into the putative targets of these effectors in infected plant hosts.

## Introduction

Fungal plant pathogens secrete small proteins, called effectors, which promote disease by targeting cellular processes in the host plant. There are hundreds of predicted effectors in the genomes of plant pathogenic fungi that are usually identified by their secretion signal and other characteristic features such as cysteine enrichment [1–3]. Some effectors are of particular interest since they constitute avirulence (AVR) factors that are detected by plant immune systems and render crops resistant to severe diseases. Fungal effectors usually show no amino-acid sequence homology to known proteins or protein domains, and consequently their biological function cannot be inferred from systematic *in silico* analysis (such as domain searches) but must be elucidated on a case-by-case basis. As a huge and growing number of putative effectors is being discovered, deciphering the functions and adaptive evolution of fungal effectors requires robust predictive tools for analyzing protein sequences and prioritizing candidates.

Recently, a combination of primary sequence pattern searches and structural modeling resulted in a major breakthrough in effector biology by revealing that fungal effector repertoires are actually dominated by a limited number of families sharing common structures despite extensive sequence variability [4–6]. One such family are the MAX (*Magnaporthe* AVRs and ToxB-like) effectors we identified in *Pyricularia oryzae* (synonym: *Magnaporthe oryzae*), the causal agent of blast disease in rice, wheat, and other cereals or grasses [7]. This pathogenic fungus is both a major threat to global food security [8] and a prime experimental model in plant pathology [9,10]. By solving the solution structure of two *P. oryzae* effectors, AVR1-CO39 and AVR-Pia, we discovered strong structural similarities between these sequence-unrelated effectors as well as with the ToxB effector from the wheat infecting fungus *Pyrenophora tritici-repentis* [7].

MAX effectors are specific to plant pathogenic ascomycete fungi, and they have undergone a major expansion in *P. oryzae*. Analysis of 120 isolates of *P. oryzae* identified ∼7800 putative MAX effectors that were grouped in 94 groups of orthologs (OGs) [11]. Individual isolates have 58 to 78 MAX effectors, corresponding to 5 to 10% of their effector repertoire. This high number suggests that MAX effectors have a critical role in the virulence of the blast fungus. This idea is further supported by the fact that MAX effectors are massively and specifically expressed during the early stages of plant infection and targeted by the plant immune system [7,10,11]. Indeed, nearly half of the cloned AVR genes of *P. oryzae* correspond to MAX effectors [7,12–15]. Analysis of the recognition of the MAX effectors AVR-Pia, AVR1-CO39, and AVR-Pik by the rice immune receptors RGA5 and Pik-1 suggests that they target small heavy metal-associated domain proteins (sHMAs), which show similarity to copper chaperones [16]. Another MAX effector, AvrPiz-t, targets four different host proteins involved in different cellular processes [17,18]. In comparison with other secreted proteins, MAX effectors show high presence/absence polymorphism and important sequence variability that is maintained by balancing selection [11]. Analysis of the MAX effectors AVR1-CO39, AVR-Pia and AVR-Pik indicates that non-synonymous polymorphisms frequently co-localize with residues interacting with immune receptors and, presumably, also with their host target proteins [11].

To better understand the function and evolution of *P. oryzae* MAX effectors, a systematic and robust analysis of their three-dimensional structure, especially outside the MAX core domain, is still needed. Indeed, in addition to the core, many MAX effectors possess N- and C-terminal extensions that could have critical roles, for instance, by establishing specific protein-protein interactions. Examining in more details these extensions, as well as other non-conserved structural features, may thus provide insights into the mechanism by which MAX effectors acquire new virulence capabilities, and allow a more comprehensive classification within the MAX family.

In the present study, we combined experimental and computational approaches to characterize finely the structural diversity of MAX effectors. We undertook structural studies of several MAX candidates and solved four new structures by Nuclear Magnetic Resonance (NMR). Comparison of these new experimental MAX structures with corresponding 3D models generated by template-based or *ab initio* approaches revealed the reliability of the MAX predictions. The highest accuracy was achieved with AlphaFold2 (AF2), which predicted the structure of MAX effectors, including non-conserved side-chains in terminal extensions that were not previously observed. We therefore revisited with the use of AF2 the structural landscape of *P. oryzae* MAX effectors, and validated the presence of a MAX core in 77 of the 94 previously defined MAX OGs [11]. Structural alignment of the AF2 models allowed us to refine the structural consensus and to explore the variability within the MAX family, including deviations from the canonical fold, disulfide bond pattern variations, additional secondary structures within N- and C-terminal extensions as well as variations in surface properties, such as stickiness and electrostatics of the core domain.

This work represents the most extensive structural analysis of a fungal effector family of a plant pathogen to date. It also provides valuable knowledge for analyses aimed at elucidating the function of MAX effectors, notably through the prediction of interaction sites within the MAX fold that could contribute to targeting host proteins during infection.

## Results

### The structure of MoToxB presents the canonical MAX fold

The MAX effector orthogroup OG33 of *P. oryzae* has high protein sequence similarity with the effector ToxB from *P. tritici-repentis* suggesting that the corresponding genes are orthologs [7,11]. We determined the structure of its representative in the Br58 isolate (S2 Table) by X-ray crystallography using molecular replacement at 1.38Å resolution (Supplementary Materials and Methods). The structure confirmed high structural similarity with ToxB from *P. tritici-repentis* and therefore this MAX effector was renamed MoToxB. The structure was also similar to the experimentally determined structures of five other *P. oryzae* MAX effectors that share less than 13% sequence identity with MoToxB (S1 Fig). Like other MAX effectors, MoToxB is structured as a 6-stranded ß-sandwich (ß1 to ß6) of two triple-stranded antiparallel ß-sheets with a ß6ß1ß2-ß3ß4ß5 topology. A disulfide bond that is conserved in almost all other MAX effectors forms a bridge between ß1 and the loop connecting ß4 and ß5. The two cysteines forming this bond are the only residues that are highly conserved in MAX effectors. In MoToxB a second disulfide bond connects ß2 and ß6 (Fig 1).

**Fig 1.**
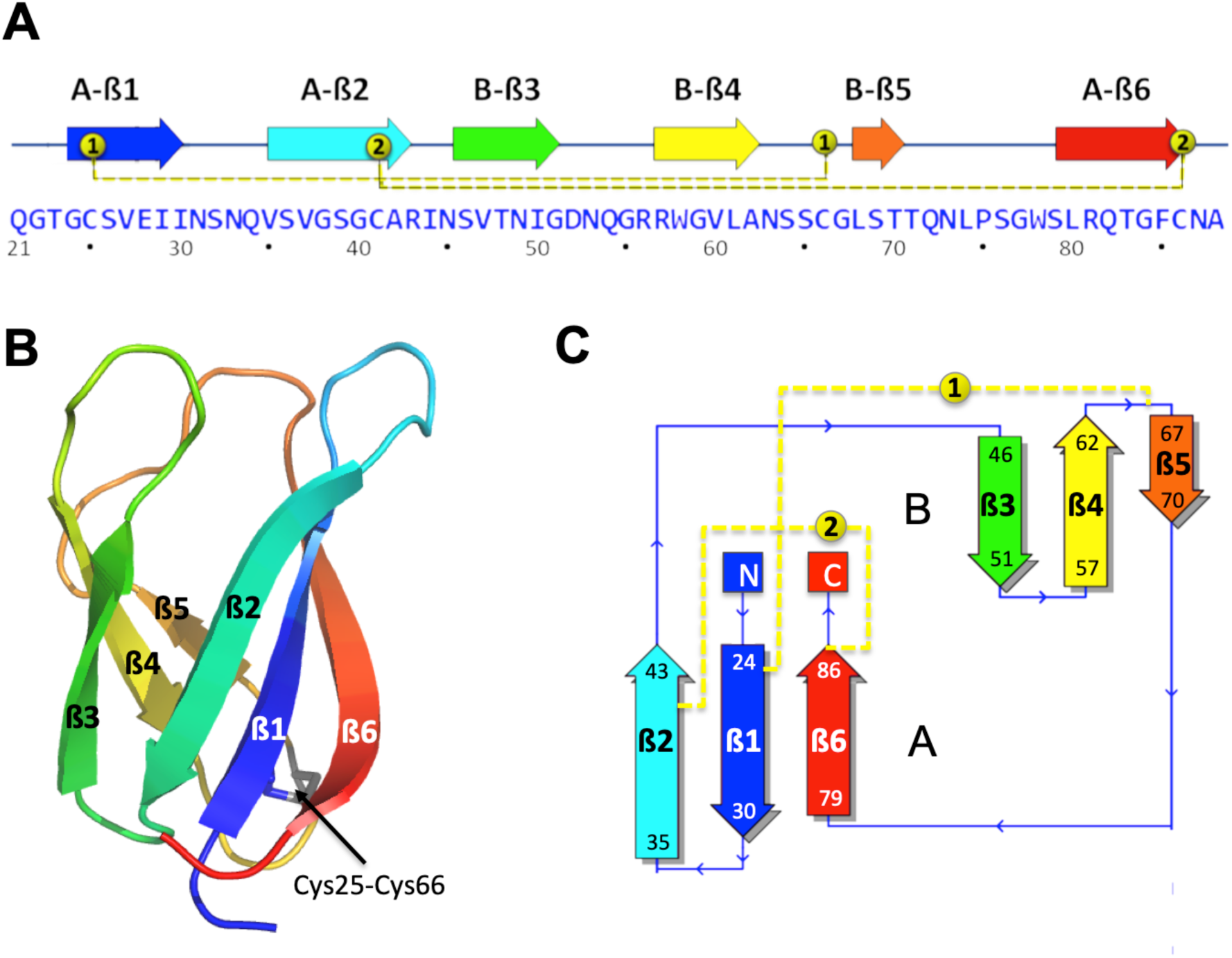
Structure of the M. oryzae ToxB (MoToxB) MAX effector. Primary and secondary structure of MoToxB showing the triple-stranded beta-sandwich forming the conserved MAX core with the two beta-sheets labeled by A and B, strands indicated by arrows and two disulfide bonds in yellow dotted lines. Disulfide bond SS “1” is almost strictly conserved in MAX effectors. (B) Cartoon representation of MoToxB crystal structure (PDB 6R5J) in rainbow color and the conserved disulfide bond “1” shown by sticks. (C) MoToxB topology diagram drawn by PDBsum and colored using the same color scheme as in A and B.

### NMR structures validate template-based modeling of MAX effectors

In our previous analysis of the MAX effector repertoire in *P. oryzae*, we used a combination of Hidden Markov Model (HMM) pattern searches and hybrid multiple Template Modeling (TM) for predicting the 3D structure of the conserved MAX core of each representative sequences of the 94 MAX effector OGs (S2 Table) defined in that study (OGs are provided in S1 Table) [11]. The reliability of the 3D models (referred as TM-pred models) was evaluated by the TM-pred score, which is an estimate of the TM-score that would be observed in a structural alignment of the TM-pred model with the corresponding experimentally resolved structure using TM-align. For these analyses eight experimental structures of MAX effectors were used as templates for homology modeling, and as a training data set for the TM-pred scoring function [11]. Almost 90% of the OG proteins were modeled at high confidence as MAX structures (TM-pred score > 0.6). Only three TM-pred models, those of OG22, OG77 and OG85, had a TM-pred score below 0.5 and were suspected to be no MAX effectors (S1 Fig). All 94 TM-pred models and associated TM-pred scores can be downloaded from the online table https://pat.cbs.cnrs.fr/magmax/model/ (see also Materials and Methods).

To deepen insight into MAX effectors and to assess the validity of the predictions, we attempted to resolve the experimental structures of 10 new MAX effector candidates with high expression during the biotrophic stage of infection (S3 Table) [11]. Using NMR spectroscopy, we successfully determined the structure of four of them (OG28, OG47, OG60 and OG67), and confirmed that they all have a MAX core fold (referred to as “MAX” instead of “OG clusters” from hereon). The 20 best-refined conformers obtained for each of these effectors were superimposed (Fig 2A), and the high quality of the NMR structures was supported by the low root mean square deviations (r.m.s.d). The complete structural statistics are given in S4 Table.

**Fig 2.**
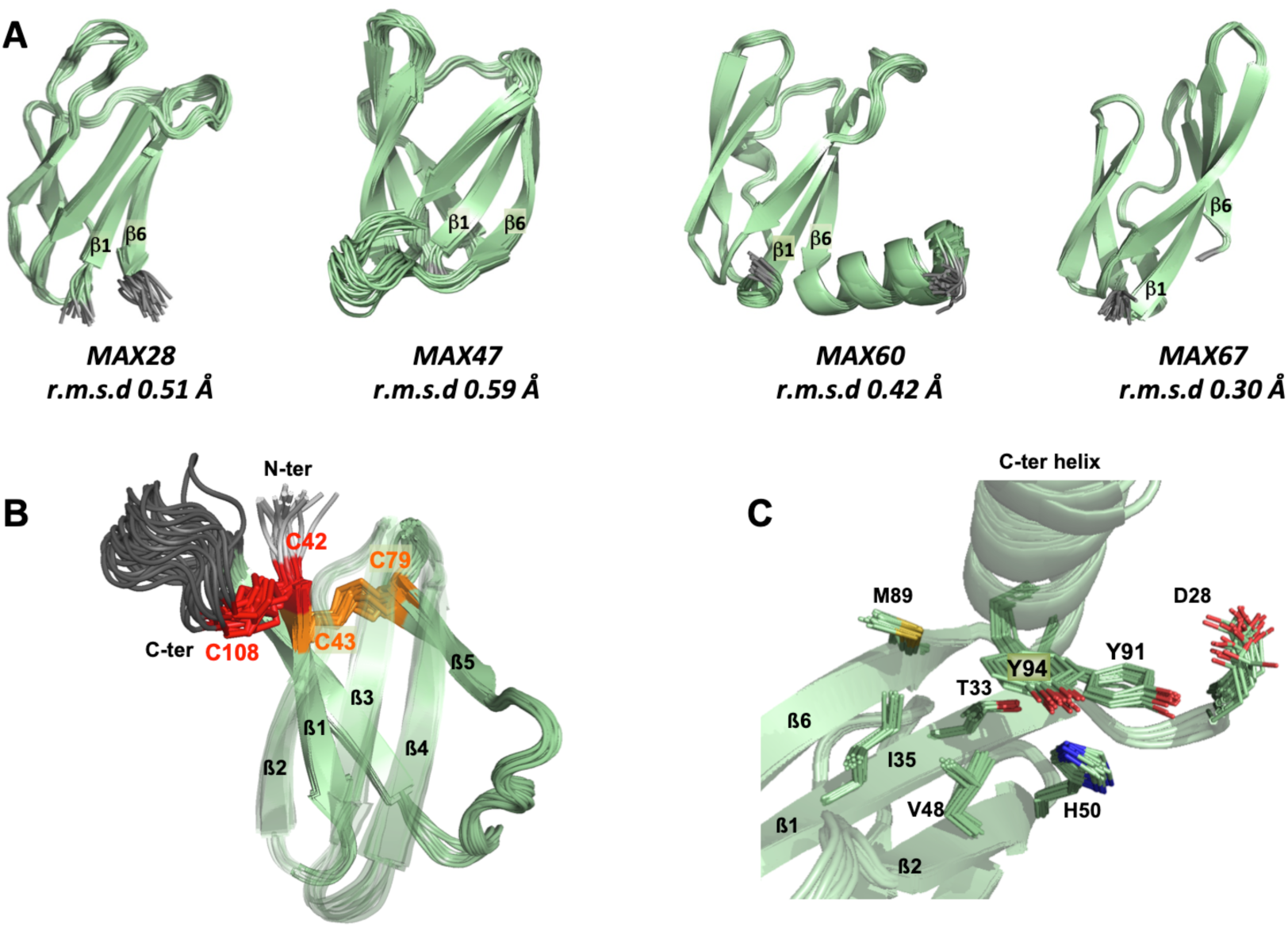
NMR structures of four MAX effectors. (A) NMR structures of MAX28, MAX47, MAX60 and MAX67 showing the superimposition and the r.m.s.d. of their 20 best conformers. The N- and C-terminal unstructured extensions before ß1 and after ß6, respectively, are not shown, except for the C-ter helix of MAX60. (B) View of the two disulfides bonds, C42-C108 (red) and C43-C79 (orange) for NMR structure of MAX47. The loop between the end of ß6 and the C-terminus is colored in dark grey. The ß2, ß3 and ß4 strands are transparent. (C) Local environment of the two residues Y91 and Y94 in the C-terminal helix of MAX60.

MAX28, MAX47 and MAX67 displayed the characteristic MAX ß-sandwich fold stapled through the conserved disulfide bridge linking ß1 and the ß4-ß5 loop (Fig. 1). The two cysteines forming this bond are the only residues that are highly conserved in MAX effectors. A particular feature of MAX67 was the exceptional length of the ß1 and ß2 strands (10 a.a.), which were longer than those in all other determined MAX effector structures. MAX60 diverged from the canonical MAX fold by the replacement of the ß5 strand by a helical turn, preventing the corresponding residues from forming a regular ß-sheet with ß4 and ß3.

In addition to the central MAX core, MAX28, MAX47 and MAX60 possess remarkable N- and/or C-terminal extensions. For MAX47, the 23 residue-long sequence extending before the ß1 strand was enriched in serine residues and was not resolved in the NMR structure. The ß1 strand started with two consecutive cysteine residues, which formed disulfide bonds that were well defined in the NMR structure (Fig 2B). The first cysteine made a disulfide bond with the last C-terminal cysteine residue (C42-C108). The second cysteine formed the disulfide bond with the cysteine in the ß4-ß5 loop (C43-C79), which is present, as already mentioned, in nearly all canonical MAX effectors, and named SS “1” disulfide bond in the following.

MAX60 has a C-terminal extension, which forms a well-defined α-helix that is attached to the structural core by hydrophobic contacts established by the aromatic rings of two tyrosine residues (Y91 and Y94). Nuclear Overhauser Effects (NOEs) in the NMR experiments revealed close contacts between tyrosine Y91 and residues D28 to T33 and H50, and between tyrosine Y94 and residues T33, I35, V48 and M89 (Fig 2C).

The resonances of the 42 residue-long C-terminal extension of MAX28 that contains lysine-repeated motifs (KxxxK) were not assigned in the NMR spectra. This is consistent with the prediction of this part of the protein as being unstructured.

The four new NMR structures of MAX effectors were superimposed using TM-align with the corresponding TM-pred models that we previously generated by template homology modeling (Fig 3A). The quality of the models was evaluated by the root-mean-square deviation (r.m.s.d) calculated between the observed and predicted structures and the TM-scores given by TM-align (a value of 1 meaning a perfect match). Comparison of the superimposed backbones showed that the overall MAX fold as well as the relative orientation of the two ß-sheets forming the central ß-sandwich were all well predicted. The prediction was particularly good for MAX28 whose MAX domain of the TM-pred model precisely matched the experimental structure (r.m.s.d.=2.11 Å), even for the loops joining the ß-strands. MAX28 was also the effector with the highest estimated TM-pred score (0.75), in remarkably good agreement with the true TM-score (0.74) of the TM-pred model aligned to the NMR structure. Structural predictions of the MAX core were also very good for MAX67, except for the long strands ß1 and ß2 in the NMR structure that could not be accurately modeled. This deviation explains the rather low TM-pred value for TM-pred_MAX67. The models of MAX47 and MAX60 showed also poor definition of certain ß-strands and exhibited strong divergence in connecting loops when compared to the NMR structures. For MAX47, the limited reliability of the TM-pred model was reflected by the low TM-pred score (0.63).

**Fig 3.**
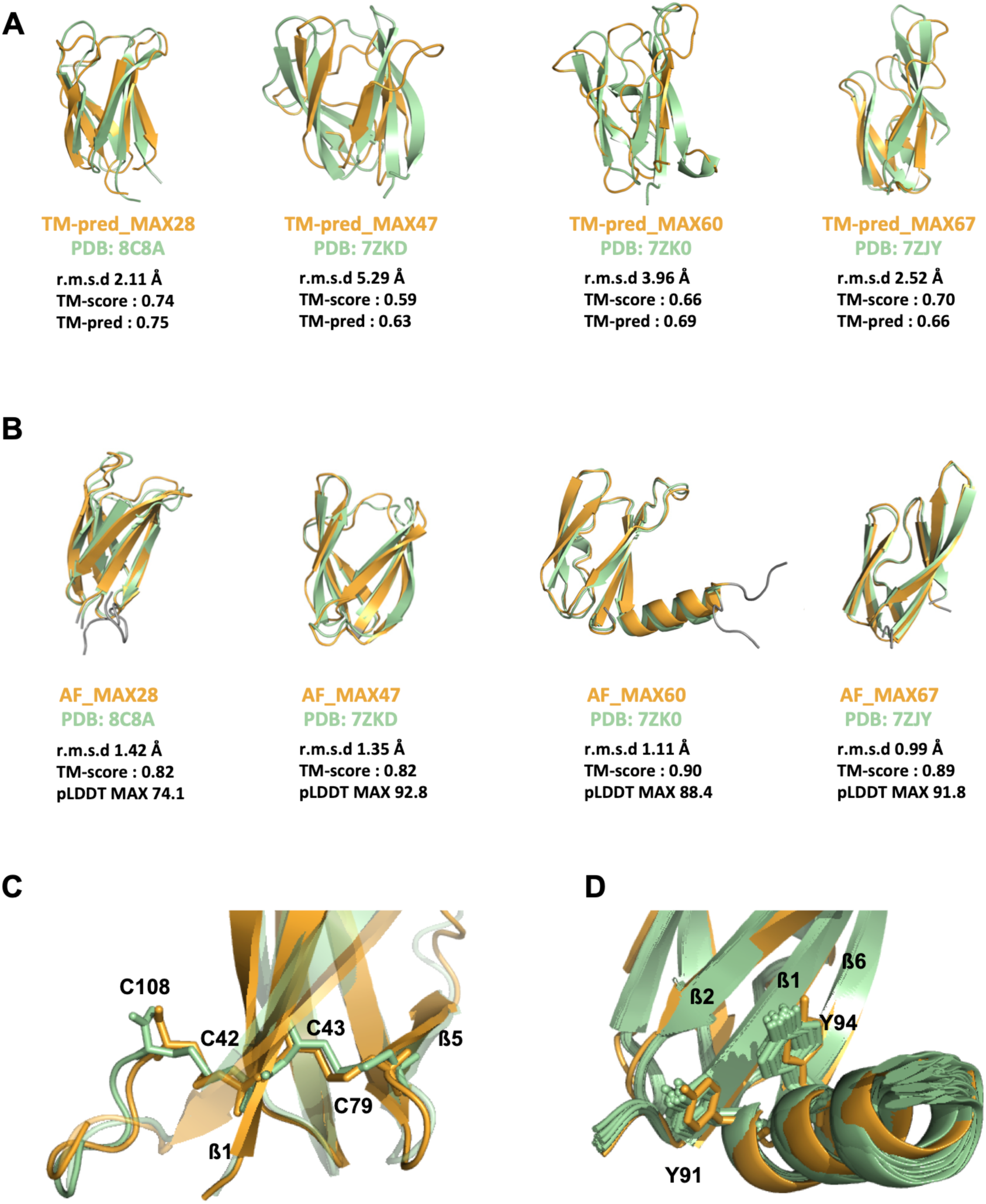
Comparison of newly determined NMR structures of MAX effectors with their TM-pred or AF predicted models. Superimposition of the four MAX effectors determined in this study by NMR shown by the best model (in green) and of their corresponding 3D models (in orange) predicted by hybrid multiple template modeling (TM-pred models shown in A) or AlphaFold2 (AF2 models shown in B). Metrics used for the quantitative assessment of the similarities between the predicted 3D models and their respective experimental structures are indicated: the root mean square deviation (r.m.s.d; the lowest, the best), the template modeling score (TM-score from TM-align, a value of 1 corresponding to a perfect match), TM-pred score (a predictive estimate of the TM-score). The N- and C-terminal boundaries were set according to the TM-pred models and did not include extensions determined in the NMR structures. (B) The r.m.s.d. between backbone heavy atoms of the superimposed NMR structure and AF2 models is given for the MAX domain only (S5 Table), as well as the predicted local distance difference test (pLDDT) that was used to estimate the reliability of the AF2 predictions in the MAX core (MAX pLDDT score, S5 Table). (C) View of the two disulfides bonds of MAX47, C42-C108 and C43-C79, as observed in the best NMR conformer and in the predicted AF_MAX47 model. (D) Position of the two tyrosyl side-chains of Y91 and Y94 in the C-terminal helix of MAX60 in the 20 NMR conformers and in the AF2 predicted model.

### AlphaFold2 reliably predicts MAX effectors core and extensions

The recent release of Artificial Intelligence (AI)-based protein structure modeling methods have demonstrated the high accuracy with which deep-learning softwares such as AlphaFold2 (AF2) can predict the 3D structure of most structured proteins [19]. To determine the accuracy of AF2 for the prediction of MAX effector structures, we used it to model MAX28, MAX47, MAX60 and MAX67 before the release of their NMR structures in the PDB. We used three different implementations of AF2 to generate the multiple sequence alignment (MSA) from which protein 3D models are predicted: the MMseqs2 and Jackhmmer implementations, which use MSAs generated automatically online or locally and that can include PDB templates, as well as the *Custom* MSA implementation that uses MSAs provided by the user and that was parameterized to not use PDB templates (see Materials and Methods). For each sequence query, the three MSAs were independently submitted to AF2, and the predictive quality of the top-ranked models was assessed thanks to the predicted local distance difference test (pLDDT) [19,20]. The pLDDT score, scaling from 0 to 100, is a residue-level accuracy score computed by AF2 that provides an estimate of the confidence of each residue’s predicted position in the protein structure. We considered the average pLDDT score for the overall protein, or only for residues within the predicted MAX core domain, hereafter, called the MAX pLDDT score. The MAX pLDDT score allows better filtering of the AF models, since it provides a quantitative measure of prediction quality focusing on the MAX core domain. For each MAX effector, the AF2 model having the highest MAX pLDDT score was selected and referred as its AF_MAX model (Fig 3).

The MAX core domain was predicted with high confidence in AF_MAX47, AF_MAX60 and AF_MAX67 according to their high MAX pLDDT scores, close to or exceeding 90 (Fig 3B and S5 Table). The best models were obtained in all three cases from the Jackhmmer AF2 implementation. A lower confidence score (74.1) was retrieved for the best AF2 model of MAX28, generated with the *Custom* MSA implementation. Nevertheless, AF_MAX28 was very close to the experimental structure of MAX28 according to the average r.m.s.d. value (1.42 Å) calculated from superimposing the MAX core backbone atoms of the NMR conformers. Indeed, for all four MAX effectors, the MAX domain of the best AF2 model displayed side-chain rotamers almost identical to those in the experimental structures and that were within the uncertainty of the NMR approach, i.e. density of NMR-derived constraints (Fig 3B).

AlphaFold2 modeling also succeeded in predicting details within the core domains. The cysteine residue side-chains forming the conserved SS1 disulfide bond were well defined in all four AF_MAX effector models. The same was true for those forming the additional disulfide bond, bridging the ß1 and the C-terminal extension, in the structure of MAX47 (Fig 3C). Another example of consistency between experimental structures and AF2 models was the remarkably well defined position and orientation of the C-terminal helix of MAX60, including the two tyrosine residues whose aromatic side chains stacked over the ß1-ß2-ß6 ß-sheet of the MAX core (Fig 3D). For MAX28, both AF2 model and NMR structure were consistent in predicting unstructured N- and C-terminal extensions. Moderate deviations from the experimental structure were only observed for residues in the C-terminus of MAX67 (S2 Fig).

### AlphaFold2 validates 77 out of 94 MAX OGs

Given the high quality of the AF2-generated models of MAX effectors, we applied the same AF modeling strategy to all other *P. oryzae* MAX OG representatives. The models were visualized to check the presence of the characteristic MAX core by inspecting the ß-strand topology and the presence of the conserved SS “1” disulfide bond (Pymol. v.1.6; Delano 2002). OG proteins showing significant topological deviations from this canonical MAX fold were discarded. The presence of the short ß5 strand was not used as a filtering criterion. Among the 80 AF2 models that matched the canonical MAX structure (Fig 4, S5 Table), 57 had MAX pLDDT scores greater than 80 and 20 had MAX pLDDT scores ranging from 60 to 80. Only three OG proteins, OG26, OG73 and OG94, exhibiting a central core compatible with a MAX fold had a MAX pLDDT score below 60 and were not kept in our final selection of 77 validated MAX structures. About 1/3^rd^ of the selected models was generated with the AF2 implementation using a *Custom* MSA constraining the alignment of the predicted ß1-ß4 strands and of the conserved cysteine residues in the SS “1” disulfide bond (S6 Table). An overview of the general structural features characterizing the 77 validated MAX effector AF models is given in S7 Table.

**Fig 4.**
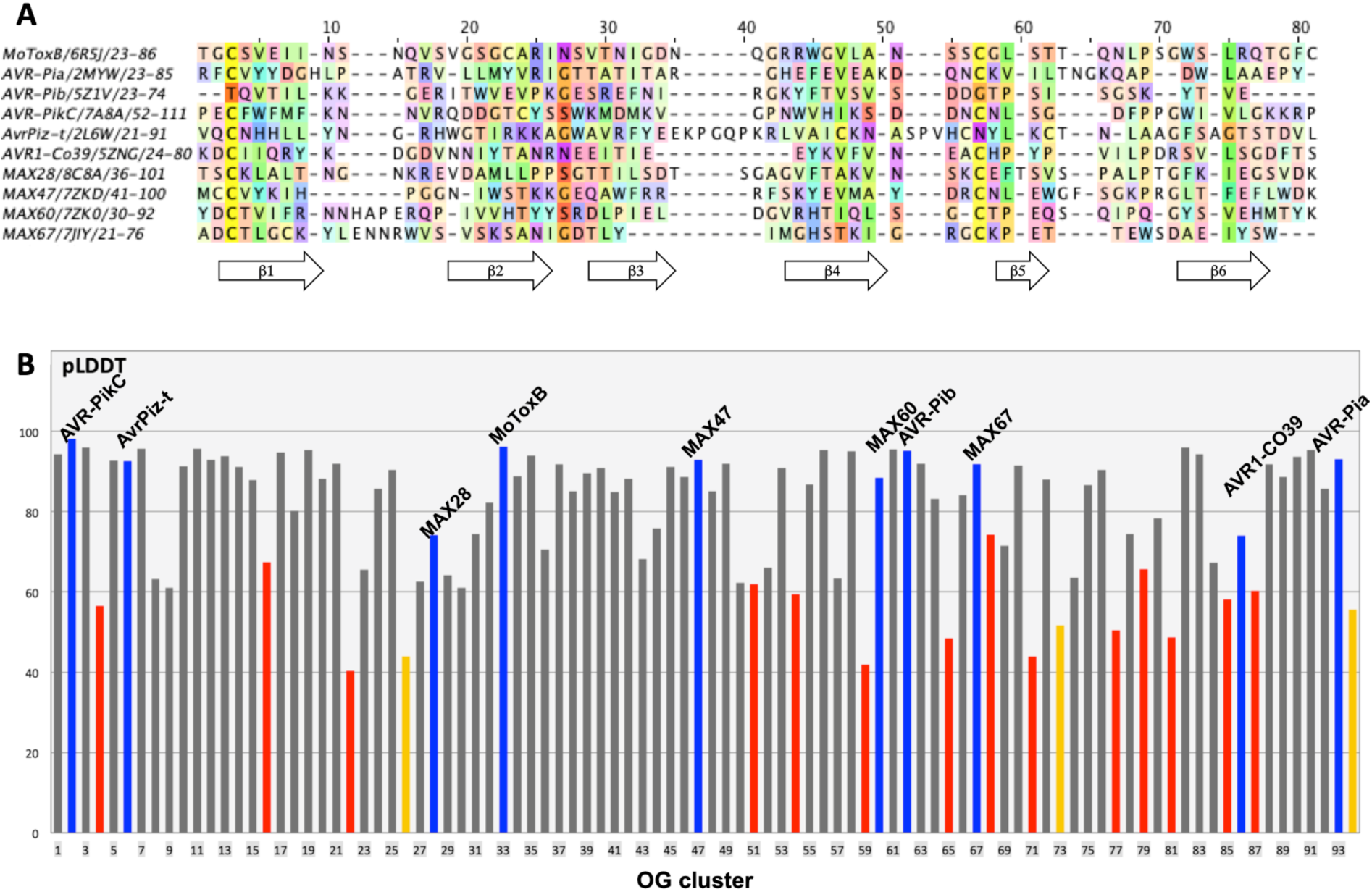
pLDDT scores of AF2 models for known and predicted MAX effectors. (A) Structural alignments of experimentally determined MAX effector structures using MoToxB structure for reference. Residues are colored according to the Taylor scheme [21](Taylor, 1997) and conservation (above 15% threshold) is used as a shading factor. (B) pLDDT or MAX pLDDT scores of the best AF2 models of the 94 OG representatives. OG representatives, whose models did not match the canonical MAX fold, have red bars showing the best overall pLDDT score. Bars of OG representatives with canonical MAX folds indicate the best MAX pLDDT score and are in blue for OGs with experimentally determined structures, in grey if the MAX pLDDT score was higher than 60 and in orange when it was below. Full data is available in S5 Table.

Among the 14 OG proteins that were not predicted to fold with the MAX topology, three (OG22, OG77 and OG85) were previously flagged as suspects based on their low TM-pred score, and five (OG04, OG59, OG65, OG68 and OG81) had a TM-pred score below 0.6 (S1 Fig). Other OG proteins such as OG51 and OG54 that exhibited a TM-pred score above 0.6 compatible with a MAX structure displayed significant distortions from the canonical MAX fold when modeled by AlphaFold2. For OG51, two models computed with MMseqs2 and the AF2 Jackhmmer implementation gave very similar models (backbone r.m.s.d. of 1.77Å) with pLDDT overall scores of 52.6 and 61.9, respectively. However, the C-terminal ß-strand of OG51 had a parallel orientation relative to the first ß1 strand that was not compatible with the MAX topology. The best OG54 model had a pLDDT score of 59.4 but deviated from the MAX topology by the absence of the C-terminal ß6 strand, which was not accurately modeled.

The 14 OG clusters that gave inconclusive AF2 models were submitted to two other protein structure prediction web-servers, RaptorX [22] and RosettaFold [23]. None of the computed models displayed the canonical MAX fold, with consistently low prediction scores, confirming the challenging nature of modeling these OG cluster sequences that may deviate from the MAX topology.

### Variations around the canonical MAX fold

From the analysis of the 77 AF models we could define more precisely the consensus structural elements that constitute the conserved MAX core. The average size of the ß strands and the connecting loops forming the canonical MAX fold given in S7 Table showed that ß1 and ß2 were usually of similar size and associated together with ß6 to form the longest anti-parallel ß sheet, while ß3 and ß4 strands were generally shorter (Fig 5A and S7 Table). Most variation occurred at the short, 2 to 8 residues long ß5 strand, which is preceded by a loop with variable length that can count up to 25 residues (MAX29). The ß5 strand can be associated with a short helix (MAX15, MAX60), totally absent (MAX78 and MAX83) or replaced by a helix (MAX20).

**Fig 5.**
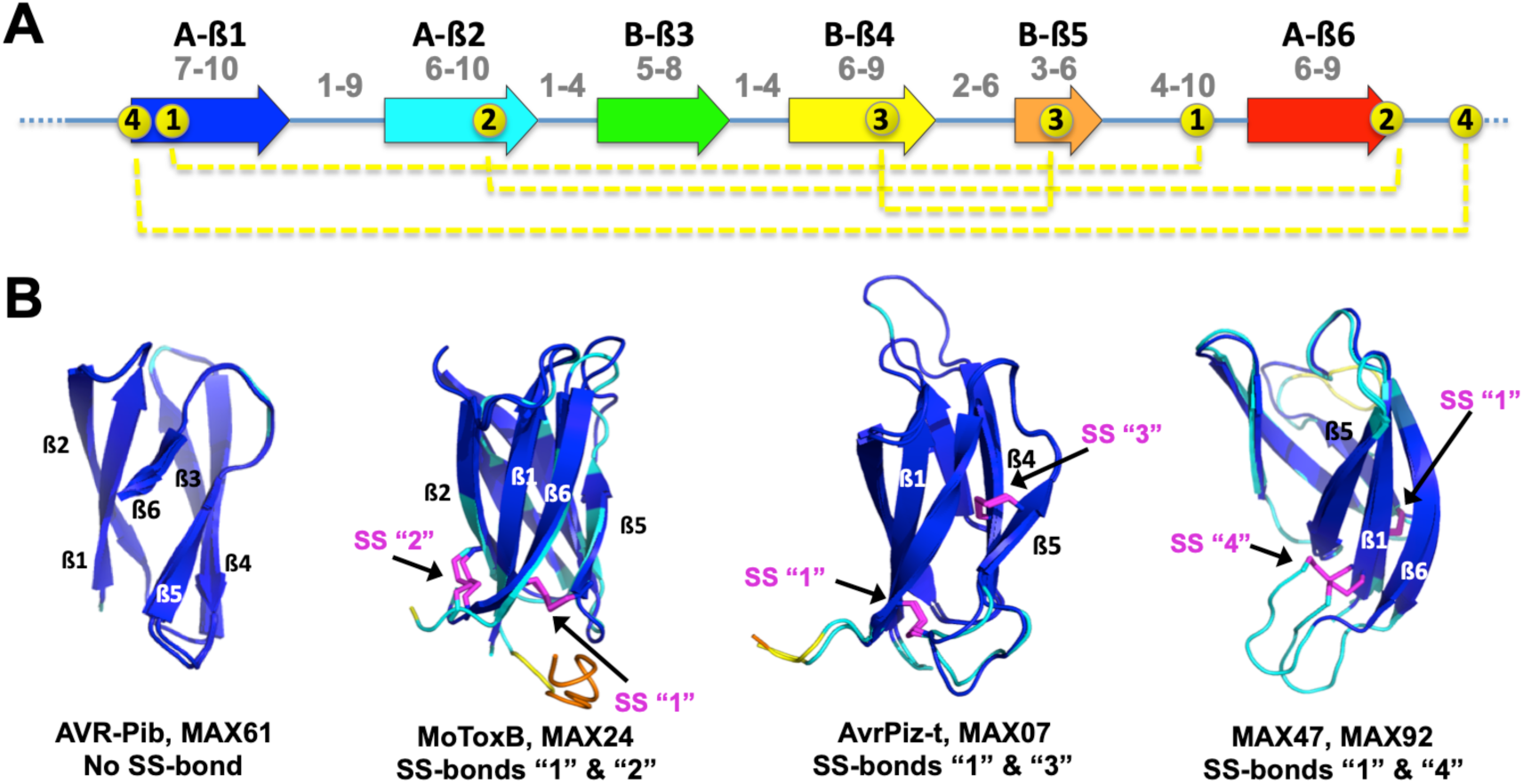
MAX domain structural features. (A) Average size range (indicated in grey) of ß strands (arrows) and connecting loops forming the central ß sandwich of the canonical MAX core, and of the N-ter and C-ter extensions. Average ranges are calculated in Table S7b. The 4 different types of disulfide bonds observed in MAX structures and models are indicated by dotted yellow lines. (B) Four different sets of structural models illustrating the variability of disulfide bond patterns. The AF2 models are colored according to their pLDDT score, by blue for high accuracy (>90), cyan for backbone at good accuracy (> 70), yellow for low confidence (> 50 and < 70) and orange for disordered (< 50). The disulfide bonds are shown in magenta, except for the AVR-Pib structure, which does not have a disulfide bond.

The number of disulfide bonds stabilizing the MAX protein can also greatly vary, from none in MAX61 and MAX62 (AVR-Pib) up to three in MAX46 or four in MAX52 (S7 Table). A unique member of the MAX family was MAX52, whose AF model consisted of two MAX core domains arranged in tandem and designated MAX52A and MAX52B in S7 Table, each having two disulfide bonds. Besides the conserved disulfide bond (SS “1”), which is a hallmark of the MAX domain, three types of additional disulfide bonds (SS “2”, “3” and “4”) were found in the experimental and AF2 MAX structures (Figs 5A and B). SS “4”, joining the N-terminus of ß1 to a C-terminal cysteine, is well defined in both MAX47 and MAX92. It was not present in any of the previously determined MAX 3D structures that could serve as template and was validated by our NMR structure of MAX47.

### N- and C-terminal extensions

Over two-third of the 77 AF_MAX models had peptide segments with 15 or more residues extending at one or both ends of the central MAX domain (S7 Table). The length of these extensions can vary among sequences belonging to the same OG cluster, especially for C-terminal, extensions (e.g. OG01, OG02 or OG15 clusters in S1a Table). C-terminal extensions were also more numerous and usually longer than N-terminal extensions. They were often modeled by AF2 with well-defined secondary structures, such as additional ß strands extending the ß2ß1ß6 sheet by one or two strands (e.g. MAX08, MAX12, MAX25), or a terminal helix as observed in the model and solution structure of MAX60. In many cases, terminal extensions appeared as unstructured regions that could not be modeled with high confidence by AF2. Long intrinsic disordered regions (IDRs) of more than 30 a.a. [24–26] may have diverse function in bacterial [27,28] and fungal effectors [29,30] and we therefore searched for IDR signatures in MAX effector sequences using ESpritz prediction software [31]. Long IDRs were predicted for ten MAX effectors and were unstructured in six AF_MAX models: in MAX15 (118 a.a.), MAX27 (36 a.a.) and MAX43 (43 a.a) as N-ter extensions, and in MAX28 (42 a.a.), MAX53 (38 a.a.) and MAX78 (43 a.a.) as C-ter extensions. It thus appears that long IDRs are a rare feature among *P. oryzae* MAX effectors, present in less than 8% of all modeled structures. The NMR solution structure of MAX28 validated the unstructured nature of its C-terminal extension.

### Two-thirds of the MAX effectors grouped in 20 structural classes

Here we performed hierarchical clustering of the selected 77 MAX effector models with two protein structure alignment software, Dali [32] and TM-align [33,34] which use different criteria for similarity scoring of superimposed structures. The Dali Z-score relies on secondary structure pairing and is a good estimate of topological conservation while the TM-score is computed for the whole alignment and weights paired residues with low r.m.s.d. more strongly than those that are more distant. When analyzed independently, the structural alignment trees retrieved from these two clustering approaches did not reveal clear sub-families of MAX structures, as shown by the lack of long internal branches in both trees (Fig 6). To facilitate the comparison of the trees, we differently colored the lines connecting each MAX model in both trees (see Materials and Methods).

**Fig 6.**
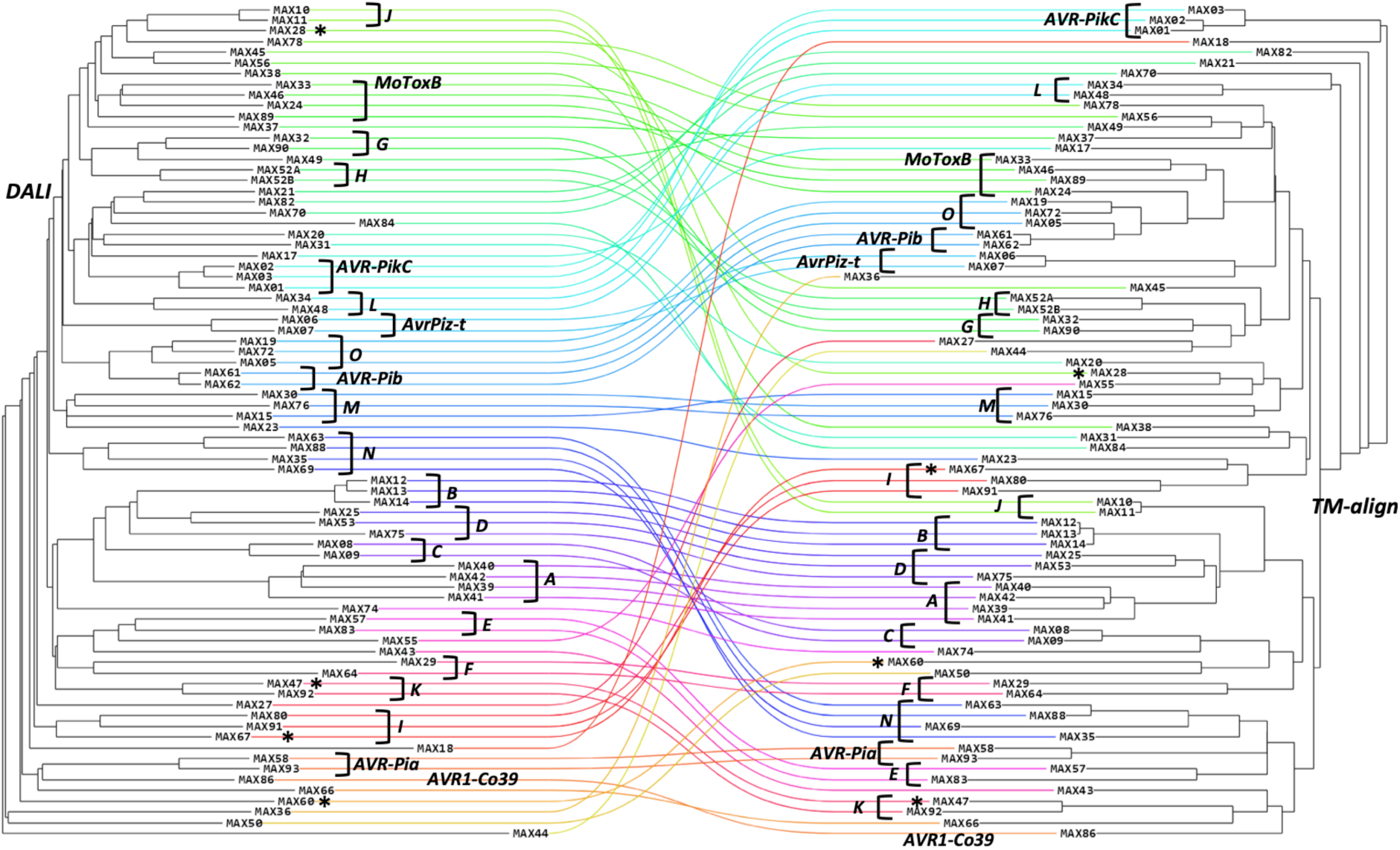
Comparison of the structural similarity trees of MAX effectors based on the Dali Z-score (left) and TM-align TM-score (right) of their superimposed AF models. Unstructured N- and C-terminal regions were removed from the AF_MAX models prior to the analysis. A line of a specific color connects each AF_MAX model in the Dali and TM-align trees. The MAX effectors with an experimental 3D structure are indicated by their name, or a star for the four novel MAX structures. The structural groups to which the MAX effectors were assigned are indicated by brackets and illustrated in Table 1 and S3 Fig.

**Table 1.**
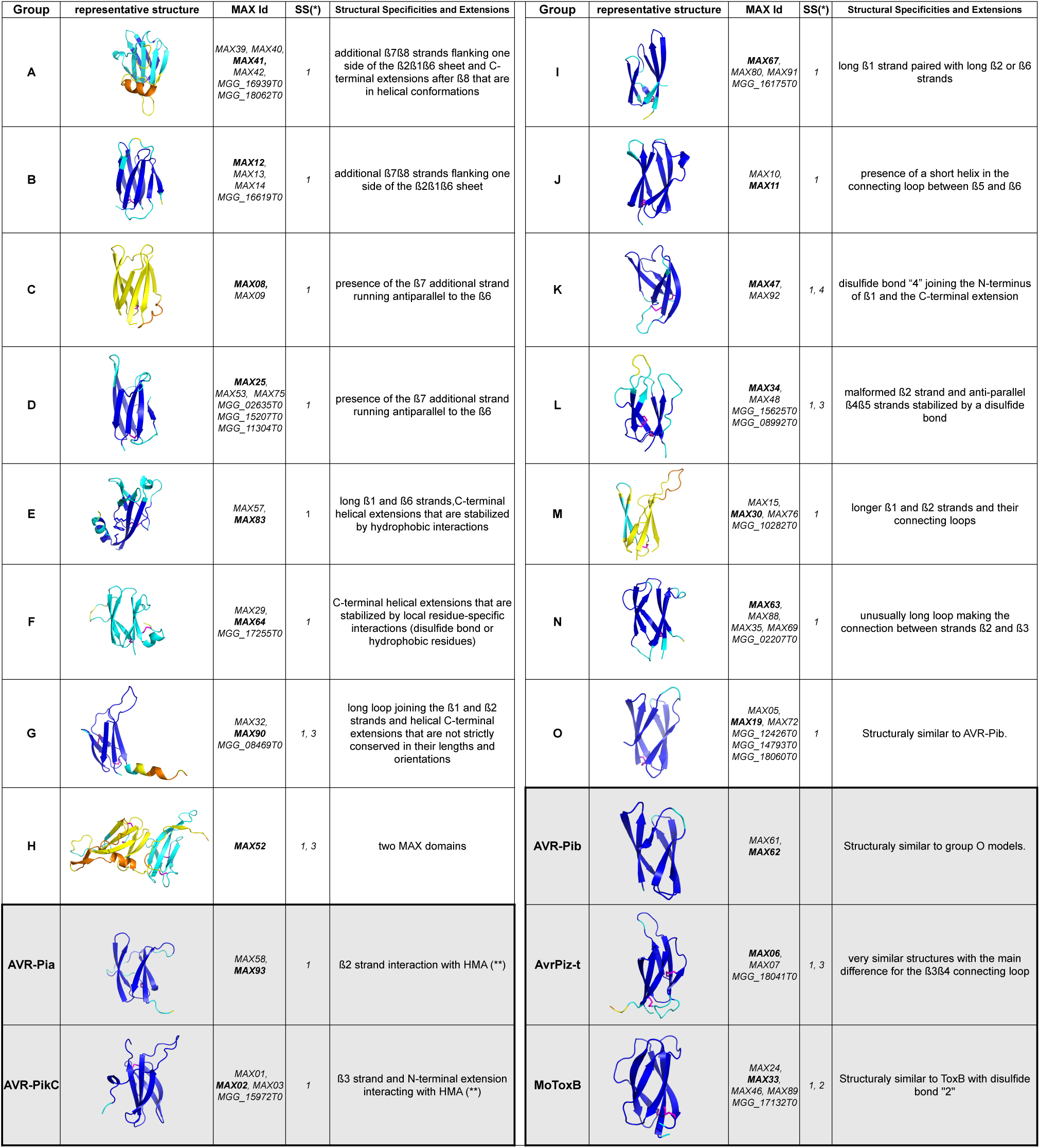
Structural groups identified in P. oryzae MAX effector family. The AF models are colored by their pLDDT scores (see legend of Fig 5). The groups with major structural variations (addition of secondary structures) are listed in the left-hand panels, including the MAX domain duplication of MAX52. The remaining groups, from I to O that do not have additional secondary structural elements but displayed specific structural variations of the MAX core domain itself are shown in the right-hand panels. To complete the overview, the five groups of well-established MAX effectors (AVR-Pia, AVR-Pib, AVR-PikC, AvrPiz-t and MoToxB) are shown at the bottom and highlighted in grey color. The MAX Id column gives the AF model identifiers of the MAX effector list reported in the present study (representative MAX model indicated in bold) and the corresponding MAX effectors reported by Seong & Krasileva, 2021 [4]; Seong and Krasileva, 2023 [5]; Yan and Talbot, 2023 [10] in the P. oryzae strain 70-15 are referred to by their MGG identifier (S8 Table). (*) Type of disulfide bond as defined in Fig 5 (**) from crystallographic structures of complexes

In this representation, any bundle of lines of similar colors highlights a possible structural similarity between the models that was common to both clustering methods and that could define a group of MAX models. Each group of at least two models was visually inspected for additional secondary structures that could add to the MAX core, the disulfide bond pattern as well as for specific structural features that fall outside the statistical average values reported in Fig 5A. Using this dual clustering method, we defined 15 groups of MAX models sharing common structural features, in addition to the 5 groups of well-established MAX effectors (AVR-Pia, AVR-Pib, AVR-PikC, AvrPiz-t and MoToxB) (Table 1). Together, these 20 groups comprised about two-thirds of the 77 MAX effectors. Groups A to G gather MAX models possessing major structured elements in addition to the canonical MAX core: groups A and B contain models with 2 extra strands, groups C and D contain models with 1 extra strand, and groups E to G contain models with C-terminal helical extensions after ß6. Group H consists of the MAX52 tandem domains connected by a structured linker. In contrast, groups I to O correspond to plain MAX structures, without other decoration but presenting variations of the MAX fold specific to each group. The characteristics of the 20 MAX structural groups are summarized in Table 1 and illustrated with more details in S3 Fig. Only 6 groups had a disulfide bond pattern with a specific SS bond adding to the conserved SS “1”, and only the AVR-Pib group had no SS bond.

### One-third of the MAX effectors are singletons

Singletons MAX effectors had no common structural features with other MAX effectors and constituted one-third of the AF_MAX effector models (26/77). The majority of them (17) consisted of a simple MAX core with unstructured N- and/or C-terminal extensions. Among them was AVR1-CO39 (MAX86). The remaining 9 singletons had diverse structured extensions (S4 Fig). In MAX49, N-terminal and C-terminal helices formed a helix bundle structure, not observed in other MAX effector models. MAX36, MAX43, MAX66 and MAX74 had an additional ß7 strand, which aligned anti-parallel to ß6, as observed for group D MAX effectors. Additional C-terminal helices were predicted for MAX44, MAX84, MAX55 and MAX60, in the last two cases with high confidence.

### MAX domains exhibit highly variegated surface properties

The comparison of the molecular surfaces of homologous proteins can highlight common or specific features related to their function. However, size differences or structural elements adding to their common fold can hamper such analysis. We therefore performed a detailed comparative analysis of the surface properties of the *bona fide* MAX effectors by focusing on the MAX core domains extracted from 49 AF_MAX models in which no structured regions interacted with the central core (S7a Table). For this subset of MAX domains, we computed SURFMAP [35] 2D projections of their molecular envelop, and compared the distribution of the following surface features: exposed secondary structures, electrostatic potential, stickiness and amino-acid polymorphism within the OG cluster to which belongs each representative MAX model (Fig 7).

**Fig 7.**
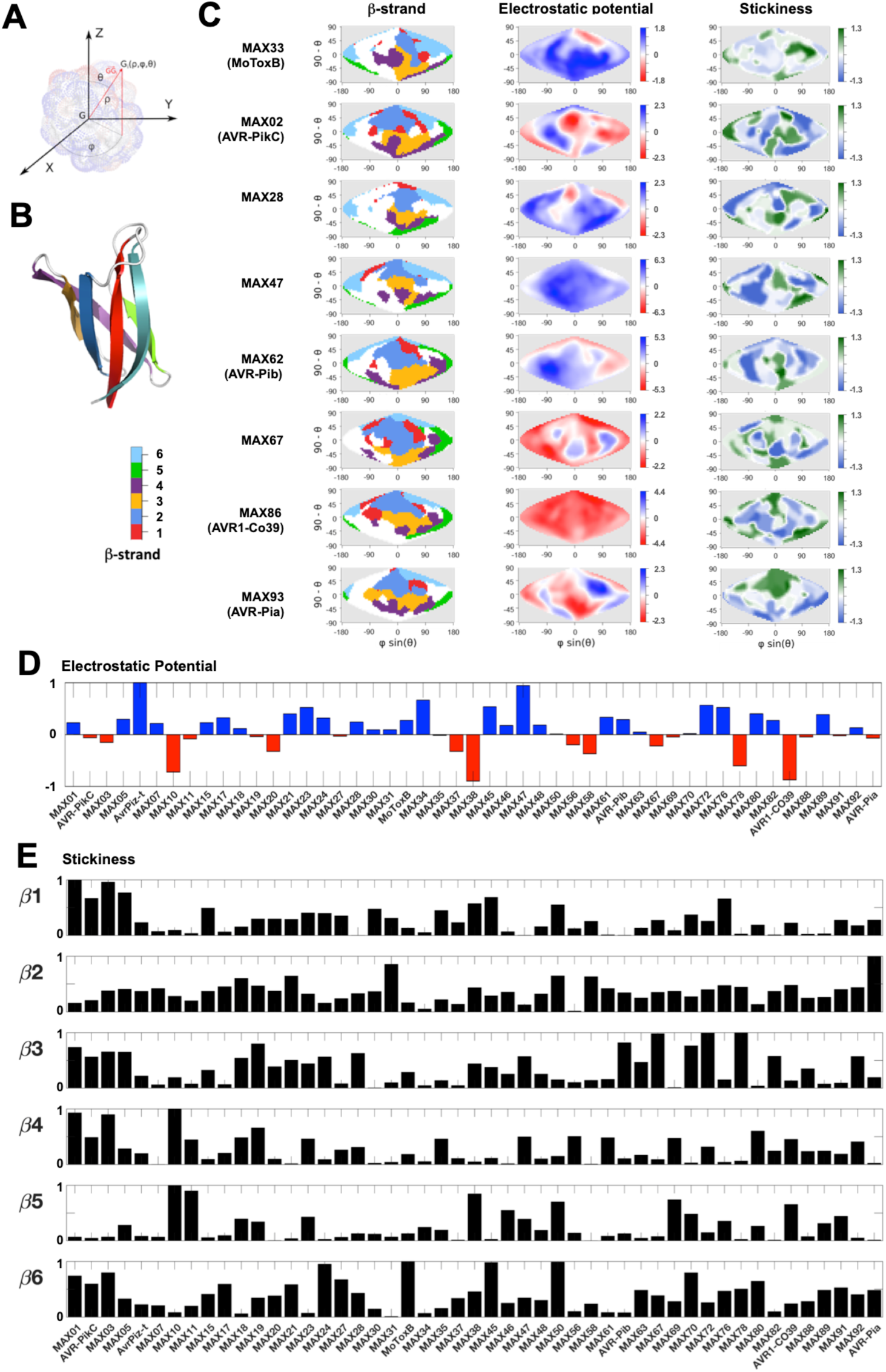
Surface properties of the MAX core domains. Surface properties of MAX core domains computed and represented using SURFMAP. A) Schematic of calculation of the spherical coordinates (from Schweke et al., 2022). The coordinates of each surface particle G_i_ is expressed in spherical coordinates (ρ, ϕ, θ), where ρ represents the distance of the particle G_i_ to the center of mass G of the protein, ϕ is the angle between the X axis and the projected vector *GG*_*l*_ in the plan (*GX*, *GY*), while θ is the angle between the vector *GG*_*l*_ and the Z axis. B) MoToxB structure showing the 6 ß-strands of the MAX core with the color code used for the surface representation in panel C. C) 2D maps of exposed ß-strands, electrostatic potential and stickiness of the molecular surfaces calculated by SURFMAP for MAX domains extracted from AF2 models of MAX effectors with known structure. N- and C-terminal extensions were discarded and the MAX core 3D models were all superimposed to AF_MAX33 (MoToxB) giving a reference frame for the Sanson-Flamsteed 2D projection computed surfaces. The ß-strand maps use the color code given in panel B. The ß1-exposed surface is mostly discontinuous around the ß2 surface that is located at the northern pole of the projection. The continuous ß3 surface lies below the ß2 surface and the discontinuous ß4 surface is found at the bottom of the projection. The ß6 surfaces are found to the west and east of the ß1-ß2 surface areas whereas ß5 surfaces are found on the most eastern areas. The electrostatic potential maps are scaled in the indicated kT/e units and the stickiness (related to hydrophobicity) scale is that defined by Levy E. D., 2010 [41]. D) Comparison of overall surface electrostatic potentials of MAX core domains, summed over the entire molecular surface and normalized by the highest absolute value calculated for the subset of 49 MAX effector models (S2 files). E) Relative surface stickiness of MAX core ß-strands. Stickiness values were summed for residues forming each of the 6 ß-strands of the MAX core domains and normalized by the highest value calculated for each strand in the subset of 49 MAX core models.

The electrostatic potential maps (Fig 7D and S2 files) revealed that MAX domain surfaces were more often positively charged than negatively charged or neutral, and that the molecular surfaces can appear entirely positive (e.g. MAX06 /AvrPiz-t, MAX23, MAX34, MAX47) or negative (e.g. MAX10, MAX38, MAX78, MAX86/AVR1-CO39), or present intense electrostatic patches (e.g. MAX02/AVR-Pik, MAX58, MAX62/AVR-Pib, MAX80). It is well established that positively charged regions in proteins are important for interaction with negatively charged macromolecules, such as nucleic acids and lipopolysaccharides [36], whereas negatively charged protein surfaces can be involved in membrane attachment or DNA mimicking functions [37–39]. In MAX47, we noticed that its unstructured N-terminal extension is rich in aspartic residues, suggesting that it could make transient interactions with the positively charged MAX core in the absence of its cellular target. In MAX62 (AVR-Pib), a surface loop region formed a strong positive patch (Fig 7C), which has been shown to be essential for the avirulence function of AVR-Pib and its nuclear localization in host cells [35]. Interestingly, a similar positive patch was visible on the surface of its structural homolog MAX61 belonging to the same structural group, as well as in MAX05 and MAX72 belonging to group O (Table1), suggesting that these effectors may also rely on this positive surface loop for their function. Similarly, AvrPiz-t displayed a positively charged surface mostly formed by lysine residues that are required for AvrPiz-t avirulence and virulence functions in rice [18]. Inversely, while the surface of MAX86 (AVR1-CO39) is strongly negative, that of MAX93 (AVR-Pia) is neutral, yet both AVR1-CO39 and AVR-Pia interact similarly through their ß2 strand with the HMA domain of the rice immune receptors RGA5 [40], respectively.

Wide divergence was also observed in the surface stickiness of the MAX domains. Surface stickiness is mostly related to surface hydrophobicity and reflects the propensity of amino acids to be involved in molecular interfaces [41]. For AVR-Pia, the HMA binding site correlated well with the presence of a large hydrophobic surface patch (Fig 7C and panel C in S5 Fig) and a very sticky ß2 strand, also present in MAX31 (Fig 7E). This was however not the case for AVR1-CO39 whose surface hydrophobicity was limited except in strand ß5. This strand is particularly variable in length and surface properties, exhibiting high stickiness in only a few MAX effectors (e.g. MAX10, MAX11 and MAX38) where it could contribute to the binding site of host target proteins. For the AVR-Pik group (MAX01, MAX02 and MAX03) surface stickiness was high in ß1 and sticky patches were also observed in strands ß3, ß4 and ß6. In all these effectors, the ß3 stickiness could serve in an interaction with strand ß4 of HMA domains, as observed in complexes of different MAX02 effectors with the HMA domain from Pikp-1 or from the rice protein OsHIPP19 targeted by AVR-PikF [12–14]. Other MAX effectors (e.g. MAX67, MAX72 and MAX78) possessed a highly sticky ß3 strand that could also associate with the ß-strand of an HMA domain or other type of protein domain. In the crystal structures of the AVR-Pik effectors, the anti-parallel ß1-ß6 strands of the MAX core make hydrophobic contacts with residues in their N-terminal extension which adopts a conserved extended conformation and considerably expands the binding interface with the HMA (S5 Fig). In MAX structural groups A to D (Table 1, not included in the present subset of Fig 7), a sticky ß6 strand was often associated with an anti-parallel ß7 extending the MAX core ß-sheet. Similarly, the highly hydrophobic ß6 strand present in MAX33 (MoToxB), MAX45 and MAX50 could interact with target proteins through an antiparallel ß-strand arrangement.

Altogether, this analysis highlights the very variegated surface properties exhibited by the *P. oryzae* MAX effectors. In spite of sharing a common fold, these sequence diverse proteins retain extensive diversity at the structural level. On the other hand, sequence conservation was high inside each OG cluster with average conservation scores in strands and loops close to the maximum conservation score of 9 (S7c Table). Only three clusters displayed low conservation scores in ß2, ß3 and ß6 strands for MAX47, in ß2 and the loop joining ß4 to ß5 for MAX63, and in strands ß4 and ß5 for MAX70, which resulted in all cases in increased polymorphism on their surfaces (S6 Fig and S2 Files).

## Discussion

### AF2 outcompetes other strategies for the prediction of MAX effector structures

In this study, we combined experimental structure determination and *in silico* modeling to elucidate the commonalities and variability of the three-dimensional structures in the MAX effector family of *Pyricularia oryzae*. Using X-ray crystallography we solved the structure of MoToxB and by NMR spectroscopy, we determined the structures of 4 new MAX effectors, MAX28 MAX47, MAX60 and MAX67, bringing to 10 the number of the experimental MAX effector structures from *P. oryzae*. This extended reference set, enabled us to evaluate the accuracy with which different modeling techniques can predict the structure of MAX effectors. In a previous work we had used a combination HMM pattern searches and hybrid multiple template modeling to predict the core of MAX effector structures and selecting best models according to their TM-pred score [11]. For the selected TM-pred models of MAX28, MAX47, MAX60 and MAX67 the overall structures of the MAX domain were properly modeled, and the observed deviations with their experimentally determined counterparts (TM score) were well predicted by the TM-pred scoring function (Fig 3A). However, the TM-pred models, although limited to the conserved MAX core, deviated substantially from the NMR structures (r.m.s.d ranging from 2.1 to 5.3 Å), highlighting the limits of template-based homology modeling [42], and the need of alternative strategies for the reliable modeling of MAX effectors.

Contrary to this, AF2 predicted with very high accuracy the new experimental structures. This was not only the case in the MAX effector core but also in structured regions outside the core. The average r.m.s.d values for superimposed backbone atoms of experimental and modeled structures were between 0,99 and 1,42 Å. Side chain rotamers and disulfide bond conformations were nearly identical, and interactions between secondary structure elements were predicted with high precision. Interestingly, this was true even in the case of MAX28 whose AF2 model had a MAX pLDDT score of only 74. These findings highlighted that AF2 predicts MAX effector structures in a highly reliable and precise manner even at relatively low pLDDT scores. Similar observations were made in other recent studies where experimentally resolved structures of fungal effectors were compared with AF2 predictions, as for example in the case of LARS [43], FOLD [44] and RALPH [45] effectors. Comparison with a published study testing *ab initio* approaches using Rosetta or the two web servers Robetta and QUARK to model MAX effectors with already known structures [46] confirmed that AF2 outcompetes other strategies for the prediction of MAX effector structures.

### Comparison with modeling of MAX effectors from P. oryzae strain 70-15

Three previous studies [4,5,10] aiming at identifying effector families in *P. oryzae* have used deep-learning methods such as TrRosetta or AF2 for systematically modeling the effector candidates from the reference isolate 70-15. In these three studies a total of either 11, 26 and 32 MAX effectors were detected, respectively. Ten effector candidates were identified as MAX effectors by all three studies, and another set of 22 MAX effectors were identified in only one or two (S8 Table). The high number of false negatives (18) in these studies that missed a large part of the MAX effectors of the 70-15 isolate highlights difficulties in model selection based on pTM-score and in detecting structural similarities using methods like CATH [47] and SCOPe [48], which were employed in these studies. All MAX effectors that were reported in these previous TrRosetta and AlphaFold studies were also identified in our analysis (Table 1 and S8 Table) with the exception of OG54 that we did not retain as a MAX effector. MAX members of the groups C, E and H were not found in the previous reports and members of groups J and K are absent from the genome of 70-15 for MAX. Taken together, these comparisons reveal that the combination of HMM-based pattern searches with systematic AF2 modeling is the most comprehensive approach for establishing the composition of fungal effector families, while structure modeling alone results in numerous false negatives, and underestimates the size of effector families.

### Classification of MAX effector AF2 models predicted in 120 P. oryzae genomes

The alignment of the 77 *bona fide* MAX effector models from AF2, NMR or X-ray crystallography permitted to determine with high precision the characteristics of the well-conserved structural core. It revealed for instance the mean length and variance of the conserved secondary structure elements and allowed the classification of possible cysteine bonding patterns. Only in exceptional cases, canonical structural features were missing or replaced. These cases mostly concerned cysteine bond SS “1” that was lacking in two MAX OGs or beta strand 5 that was replaced by an alpha helix or absent in five MAX OGs.

Comparison of the 77 MAX effector structures also showed that, beyond the well-conserved core, the MAX effector family harbors important structural diversity. Clustering of the structures distinguished 20 subfamilies comprising 51 orthogroups as well as 26 singletons that could not be classified. Major distinctive features are additional structured regions in the C-terminus characteristic of certain subfamilies. Groups A to D have one or two additional strands and groups E to G possess helical extensions. These regions presumably act in protein-protein interactions or contribute to overall functionality of the effectors. Notably, our study uncovered a domain duplication event within one of the MAX effector clusters (group H). Other dual-domain effectors have been described in recent studies, i.e. the Fol dual-domain effectors (FOLD) [44] and effectors predicted from *Puccinia graminis* [5]. The discovery of dual-domain effectors, including the domain duplication found in one of the MAX effector cluster, adds to our understanding of the diversity and complexity of fungal effector proteins. These dual-domain effectors likely have evolved to possess multiple functional domains that contribute to their virulence or interaction with host plants.

In addition to contributing to effector function, N- and C-terminal extensions of MAX effectors may have critical roles in protein folding. This hypothesis is supported by studies, where we used HP-NMR (high Hydrostatic Pressure NMR) [49,50] to analyze the folding/unfolding of AVR-Pia, AVR-Pib [51] and MAX60 [52]. While the MAX effector core of AVR-Pia and AVR-Pib folded similarly around a ß3ß4 intermediate, MAX60 had an early folding intermediate formed by ß1, ß6 and the C-terminal helix, a specific extension of this MAX effector. Mutants lacking this helix were not sufficiently stable to be purified. These findings show how additional sequences outside the core can have profound impacts on the folding of MAX effectors.

Beside structured extensions, long intrinsically disordered regions (IDRs) were only observed in six MAX effectors. IDR regions lack a stable 3D structure and exhibit conformational flexibility, allowing them to interact with multiple binding partners and fulfill various functions [53] but are difficult to precisely predict from the sequence [54]. Modest structural variations were detected within the MAX core of the seven subgroups I to O and distinguished these groups from each other. The classification provided in Table 1 can be further improved by incorporating surface properties. Thus, Avr-Pib and O groups could be combined, since they are structurally similar and members of the O group have electrostatic surfaces related to AVR-Pib.

### MAX effectors of Venturia inaequalis are distinct from P. oryzae MAX effectors

*V. inaequalis* is an ascomycete fungus, in the *Venturiaceae* family, responsible for apple scab disease. Although it is only very distantly related to *P. oryzae*, V. *inaequalis* has also an extended MAX effector family as revealed by systematic modeling of its effector repertoire [55]. However, none of the *V. inaequalis* MAX effectors fitted into any of the *P. oryzae* MAX subfamilies. Indeed, all *V. inaequalis* MAX effectors present three conserved disulfide bonds, of which one is the canonical SS “1” bond characteristic of MAX effectors of *P. oryzae* and other fungi, while the remaining two were not found in MAX effectors of any other species. Moreover, MAX-like effectors of *V. inaequalis* usually possess a C-terminal helical extension connected to the MAX core domain *via* a specific disulfide bond. This defines these newly discovered MAX-like effectors from *V. inaequalis* as a distinct subfamily with unique sequence and structure features [55]. *V. inaequalis* colonizes the leaf surface by growing below the cuticle, and releases effectors in this sub-cuticular host environment without penetrating the underlying epidermal cells. Due to this specific life style, the function and host targets of the *V. inaequalis* MAX effectors, are presumably fundamentally different from those of *P. oryzae*.

### Conclusion and perspectives

Structural information tremendously extends the insight, which can be obtained from primary sequence, and expands our understanding of biological processes or evolution to the atomic level. However, corresponding analyses are far from being trivial especially in rapidly evolving protein families with high sequence diversity, as it is the case of fungal effector proteins. Our study shows that HMM pattern searches associated with AF2 structure modeling provide a solid basement for establishing in a comprehensive manner effector families in fungi. In addition to providing detailed insight into effector family diversity, such studies generate solid bases for the functional investigation of fungal effectors and for the investigation of their evolutionary trajectories. The combination of structural modeling and population genomics provides exciting perspectives for accelerated and deepened investigation of the molecular evolution of fungal effector proteins [11]. Ongoing and rapid improvements in *in silico* protein-protein interaction analysis, such as improved prediction of the structures of protein complexes [56] or screening of interacting proteins [57], are opening a new era in the functional investigation of effectors by generating at high throughput hypothesis for effector function and evolution. However, for both of these research areas experimental structure determination remains critical, since large parts of fungal effectoromes can still not be modeled with good confidence, and models of protein complexes generally lack precision and therefore only provide limited insight into the identity and conformation of the directly interacting residues and the details of the binding interface.

## Materials and Methods

### Experimental Structures

#### MAX28 Protein expression and purification

Protein expression and purification experimental details for MAX47, MAX60 and MAX67 are available in [58] Mounia et al. 2022. For MAX28 we followed essentially the same protocol than for the other MAX effectors for producing the ^15^N-labelled NMR sample but the His_6_-tag was not cleaved to keep the protein soluble. Uniformly labeled ^15^N MAX28 was expressed in E. coli BL21 (DE3) cells (Invitrogen, Thermo Fisher Scientific, Waltham, USA) from a homemade plasmid pDB-his-CCDB-3C (courtesy of Frederic Allemand, CBS Montpellier, France). Protein expression was carried out in ^15^NH_4_Cl (1 g/l) enriched M9 medium. Cells were grown at 37 °C until reaching an OD600 = 0.8 and then, expression proceeded overnight at 30 °C after induction by addition of 0.3 mM IPTG. Cells were harvested by centrifugation, re-suspended in denaturing buffer (50 mM Tris, 300 mM NaCl, 1 mM DTT (dithiothreitol), 8 M urea, pH 8) and lysed by ultra-sonication. The supernatant containing the unfolded protein was applied to a HisTrap HP 5 ml affinity column (Cytiva, Freiburg im Breisgau, Germany). The His_6_-tagged protein was eluted in 50 mM Tris, 300 mM NaCl, 1 mM DTT, 8 M urea, pH 8 with an imidazole gradient up to 500 mM. At this step, MAX28 was directly dialyzed against 10 mM Na Phosphate, 2 mM DTT, 150 mM NaCl, pH 6.8 buffer in order to remove imidazole and urea, allowing the refolding of the protein. The MAX28 samples were then concentrated using Amicon Ultra Centrifugal Filter Devices (MW cutoff 3000 Da), (Merck Millipore, Burlington, USA) prior to size exclusion chromatography (SEC) using HiLoad 16/600 Superdex 75 pg column (Cytiva). Fractions containing protein were pooled, concentrated to 0.4 mM and stored at −20°C. All NMR experiments were carried out at 27°C on a Bruker Avance III 800 MHz or Bruker Avance III 700 MHz spectrometer, both equipped with 5 mm z-gradient TCI cryoprobe. NMR samples consisted on approximately 0.4 mM ^15^N-labeled protein dissolved in 10 mM Na-Phosphate buffer (pH 6.8) and 150 mM NaCl with 5% D_2_O for the lock.

#### NMR Structure determination of MAX28, MAX47, MAX60 and MAX67

^1^H chemical shifts were directly referenced to the methyl resonance of DSS, while ^15^N chemical shifts were referenced indirectly to the absolute ^15^N/^1^H frequency ratio. All NMR spectra were processed with Topspin 3.6 (Bruker) and analyzed with Cindy 2.1 (Padilla, www.cbs.cnrs.fr). Assignments for MAX28, MAX47, MAX60 and MAX67 have been deposited to and are available from the BMRB data bank under the accession entry 34782, 34731, 34730 and 34729, respectively.

The NMR structures were determined from the NMR constraints listed in S4 Table that were obtained as follow. NOE cross-peaks identified on 3D [^1^H, ^15^N] NOESY-HSQC (mixing time 150 ms) were assigned through automated NMR structure calculations with CYANA*3* [59,60]. Hydrogen bond restraints were derived using standard criteria on the basis of the amide ^1^H / ^2^H exchange experiments and NOE data. When identified, the hydrogen bond was enforced using the following restraints: ranges of 1.8–2.0 Å for d(N-H,O), and 2.7–3.0 Å for d(N,O). Dihedral restraints were obtained from TALOS-N [61] analysis of backbone atom chemical shifts for MAX47, MAX60 and MAX67. For the final list of restraints, distance values redundant with covalent geometry were eliminated and disulfide bonds that were consistent with short distances between cysteine residues were added.

A total of 200 three-dimensional structures were generated using the torsion angle dynamics protocol of CYANA*3* from NOEs, hydrogen bonds and disulfide bond restraints (S4 Table). The 20 best structures (based on the final target penalty function values) were minimized with CNS 1.2 according to the RECOORD procedure [62] and analyzed with PROCHECK [63]. The rmsds were calculated with MOLMOL [64]. All statistics are given in S4 Table.

The structure coordinates have been deposited at the Protein Data Bank under the following accession codes: MAX28 (PDB_8C8A), MAX47 (PDB_7ZKD), MAX60 (PDB_7ZK0), MAX67 (PDB_7ZJY),).

### TM-pred MAX models Web Table

Homology models of each OG representative sequence relative to each 3D template were built using MODELLER v9.1 [65] with several alternative query-template threading alignments as described in Le Naour—Vernet et al., 2023 [11]. The top-5 models were selected according to the TM-pred scoring function and are given in the Web page: https://pat.cbs.cnrs.fr/magmax/model/

The Web table has the following columns: Group (OG cluster), Score (composite evaluation score of the best model -best scores are in red, worst are in blue-), Dfire, Goap, Qmean, E1D, E2D, E3D (individual evaluation scores of the best model), Alignment (aligned identifier sequences), Identifier (protein identifier of the orthologs prioritizing *Oryzae* infecting strains). Structural models and alignments are available by clicking on each OG cluster in the first column. A multiple sequence alignment of non-redundant orthologous proteins is displayed at the top of the page, starting with the modeled representative sequence. For each OG cluster given in [11] we further filtered out redundant sequences using CD-HIT [66] (S1 Table). The representative sequence of each OG cluster was determined to be the sequence sharing the highest sequence identity with a consensus sequence derived from the OG cluster sequence alignment by MAFFT [67] (S2 Table). Signal peptide prediction before the cleaved residue (cs.SIGNALP41), mean hydrophobic index (ih.MEAN), sequence conservation (1D.CSRV), alternative alignments and model secondary structures (2d.STRIDE) are displayed below the sequence alignment. Sequence conservation scoring implemented in PAT [68] was calculated according to Sonnhammer and Hollich, 2005 [69] with a Gonnet matrix [70]. At the bottom of the page, the 5 best models can be displayed using different representations and color schemes. Models can also be downloaded in PDB format.

### Modeling by AlphaFold

For each OG representative sequence we computed three AF models differing by the way of building multiple sequence alignment (MSA). The MMseqs2 MSA was obtained from the MMseqs2 [71,72] web server as implemented in the ColabFold version of AlphaFold 2.0 [73]. We also used the version of AlphaFold 2.2.0 that builds MSAs by Jackhmmer on uniclust, mgnify and uniref90 databases. These two implementations use PDB templates. Finally, a *Custom* MSA was build from Muscle_v3.8.31 [74] by inserting (-profile option) the query sequence on top of a previously computed MSA, termed ß1ß4_MSA. The ß1ß4_MSA was build from a Muscle alignment of the OG sequences (S1 Table) by filtering out those having the two flanking cysteine residues in the ß1 and in the loop between ß4 and ß5 strands not correctly aligned to the 8 3D template sequences. The ß1ß4_MSA was further processed by truncating the sequences by eliminating residues (-2 included) before and (+2 included) after the first and last aligned cysteine residues, respectively, and filtering out for redundant sequences by CD-HIT, giving a total size of 247 aligned sequences (S6 Table). For each query, the consistency of the *Custom* MSA was determined by checking the correct alignment of the cysteine residues in the query and in the appended ß1ß4_MSA. When consistent the *Custom* MSA was converted to a3m format by the reformat.pl script [75] and directly used as input in the ColabFold implementation of AlphaFold 2.0 calculations that was setup without the use of PDB templates. *Custom* MSAs could not be built for OG61 and OG62 from absence of cysteine residues in their primary sequences and were not consistent for OG15, OG27, OG71, OG81, OG85 and OG92.

The quality of each model was assessed by the pLDDT overall score [76]. The correct MAX topology was verified by visual inspection (Pymol v.1.6 Delano 2002). For models having the MAX topology, a MAX pLDDT score that was an average score of residues in the MAX core domain (including residues from ß1 to ß6) was calculated. For each query, the best AF2 MAX model was selected when the MAX pLDDT score was above 60. The complete set of AF2_generated models for the 77 validated MAX effectors is available in S1 Files.

### Dali implementation and search

A standalone implementation of DaliLite.v5 [77] was used for this work. For the all-to-all clustering by Dali we first discarded *unstructured* stretches in each model. For this, the *structured* domain of each model was defined by taking the STRIDE [78] output, and filtering for the first residue in the first and last residue in the last secondary structure (helix or strand), respectively (S7 Table). The model of MAX52 was split in two chains A and B each containing a MAX domain. All these *structured* domains were used for clustering with Dali Z-scores excluding *de-facto* unresolved protein regions without loosing important structural information.

#### TM-align scoring and side-by-side plot with Dali Z-score tree

The distance between each pair of AF2 models that were used for Dali clustering was estimated by the TM-score obtained from TM-align after pairwise model superposition. A classification tree was then inferred from these pairwise distances using FastME [79]. Finally, Fig 6 was obtained by joining identical models in the FastME tree and in the Dali tree, respectively, by a line of the same color.

### Surface properties of MAX core domains

A subset of 49 MAX effector AF2 models, each consisting of a MAX core domain and optional N- and/or C-*unstructured* extensions (S7 Table) was defined by discarding AF2 models having N- and/or C-terminal *structured* extensions (listed in the groups A to H in Table 1 and in S4 Fig). All MAX core domains of the 49 AF2 models were superimposed to the reference MoToxB structure with their ß1 strand vertically aligned to the Z Cartesian axe giving a reference frame for the Sanson-Flamsteed 2D projection computed using SURFMAP [35]. Their surface properties including stickiness and electrostatics (APBS) [80] were computed by SURFMAP and are given in S2 File. The temperature factor column of the PDB files was used to encode the color of the exposed surface of the six ß-strands, from 1 to 6, respectively. The sum of the surface stickiness positive values of each individual ß-strand was computed by filtering SURFMAP surface stickiness output and are reported in Fig 7E. The amino-acid conservation scores given for each OG cluster to which belongs each representative MAX model were used to color encode the surface from white for high conservation score of 9, light blue colors for intermediate conservation scores (from 8 to 6), sky-blue for low conservation score of 5 and darker blue colors indicating highly polymorphic positions with conservation scores of 4 and below.

## Author Contributions

M.L., and K.d.G. prepared the ^15^N-^13^C-labeled protein samples (sub-cloning, protein expression and purification). P.B. made the NMR resonance assignment of the four proteins in the study conditions. J.G. wrote scripts for protein sequence analysis and homology modeling. C.R. and A.P. supervised the project, conceived experiments and modeling, participated in the interpretation, and M.L., A.P., N.D., C.R., S.C., P.G. wrote the article. A.P. also contributed to the funding acquisition with P.G. and T.K. All authors have read and agreed to the published version of the manuscript.

## Acknowledgements

We are grateful to Liisa Holm for feedback in implementing Dali. The authors are grateful to J. Maidment (CBS, Montpellier) for careful proof reading and language correction of our manuscript. We are grateful to Matthew Bowler and Didier Nurizzo at the European Synchrotron Radiation Facility (ESRF), Grenoble, France for providing assistance in using beamline ID30A-1. The authors gratefully acknowledge the ESRF for provision of synchrotron radiation facilities via Block Allocation Group beamtime.

## Supporting information

S1 Fig. Template-based modeling of MAX effector sequences

S2 Fig. NMR structures and AF models

S3 Fig. Groups of AlphaFold models

S4 Fig. Singletons MAX effectors with structured extensions

S5 Fig. Hydrophobic residues on the surface of AVR-Pik and AVR-Pia

S6 Fig. Amino-acid polymorphism mapped on the surface of AF MAX models

S1 Table. List of Pyricularia (syn. Magnaporthe) oryzae entries in the 94 MAX effector orthogroup (OG) clusters

S2 Table. Representative sequences of the 94 MAX effector orthogroup clusters

S3 Table. Report on structural studies of 10 putative MAX effectors (OG proteins)

S4 Table. Refinement statistics of MAX effector NMR structures and AF model superimposition

S5 Table. Summary of AlphaFold2 modeling statistics and list of validated MAX effectors

S6 Table. ß1ß4 MSA

S7 Table. Structural features of 77 AF-modeled structures of validated MAX effector, statistics and conservation scores.

S8 Table. Comparison of our selection of MAX effectors with previous modelisation studies

Supplementary Materials_and_Methods. Crystallographic structure determination of MoToxB

S1 File. Zipped folder of AF MAX models

S2 File. Zipped folder of AF MAX model surfaces

