## Supplementary material for "The structural landscape and diversity of *Pyricularia oryzae* MAX effectors revisited": Figure_S1

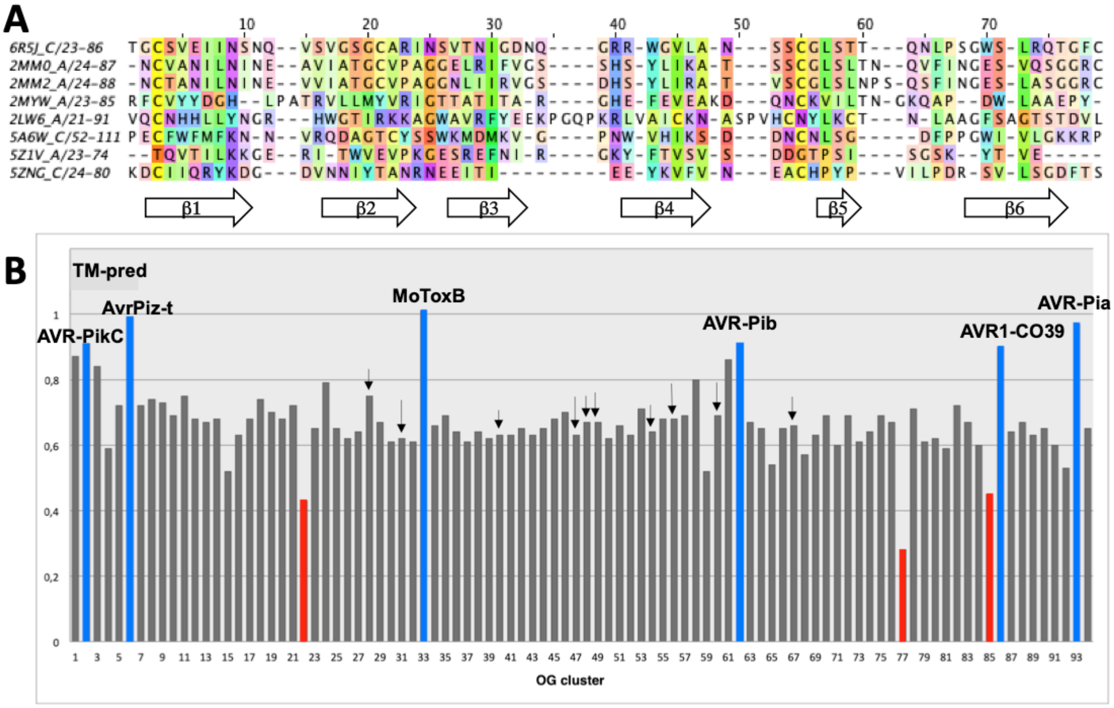


**S1 Fig.** ***Template-based modeling of MAX effector sequences.*** *(A) Structural alignments of the eight experimentally-determined MAX effector templates used in the TM-pred score training data set. The PDB codes are reported for MoToxB (6R5J, top sequence), ToxB and variant (2MM0 and 2MM2 respectively), AVR-Pia (2MYW), AvrPiz-t (2LW6), AVR-PikD (5A6W), AVR-Pib (5Z1V) and AVR1-CO39 (5ZNG) and underscoring indicates the chain Id. The percentage of conservation (above 15% threshold) was used as a shading factor over Taylor color-coded residues using JALVIEW version 2.11.2.4. MoToxB and ToxB have homologous structures with 32% of identical aligned residues, Dali Z-score and TM-align score of 10.3 and 0.82, respectively. The average residue identity between MoToxB and all other 5 MAX sequences (excluding ToxB and variant) is 13%. The white arrows indicate consensus strand positions. (B) Best TM-pred scores of the representative sequences of all OG clusters. The scores of experimentally determined structures (including AVR-PikC - PDB 7A8X) are shown in blue. OG clusters having the lowest TM-pred scores are shown in red. Black arrows identify the OG proteins that were submitted to expression and purification trials (S3 Table).*
