## Supplementary material for "The structural landscape and diversity of *Pyricularia oryzae* MAX effectors revisited": Figure_S2

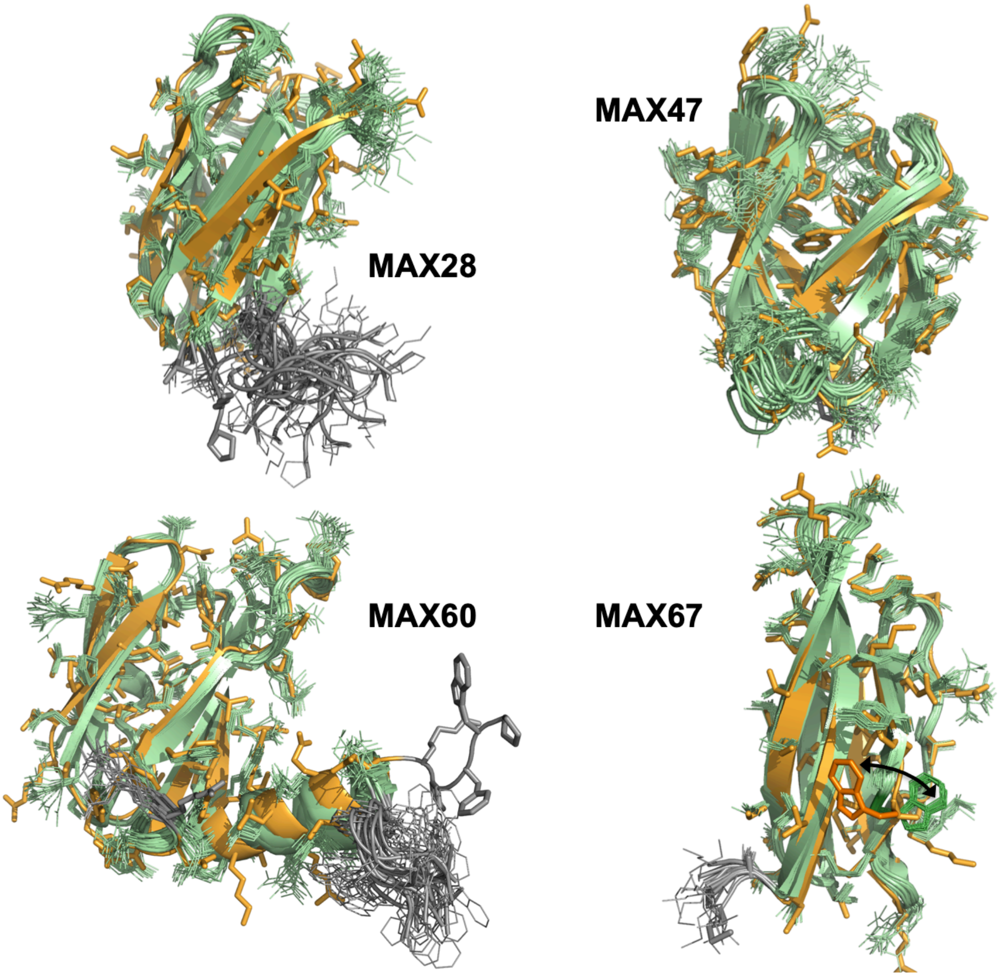


***S2 Fig. NMR structures and AF models***

*Superimposition of the 20 conformers of the NMR structure (green) and AF model (orange). Side-chains are shown by lines and sticks. The black arrow shows the different orientation of the C-terminal W77 in the AF model and in the NMR structure of MAX67.*
