## Supplementary material for "The structural landscape and diversity of *Pyricularia oryzae* MAX effectors revisited": Figure_S3

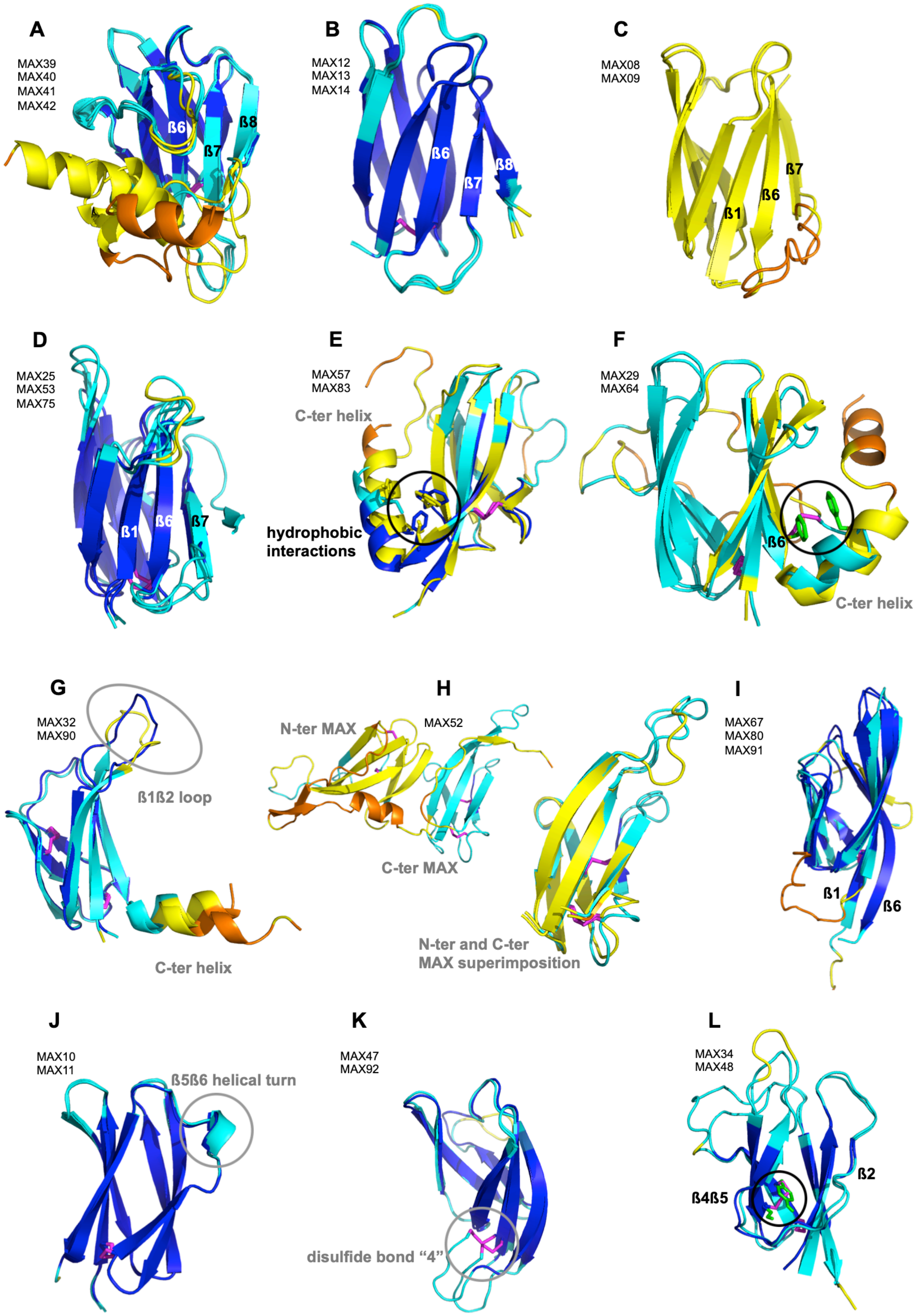
A
MAX39
MAX40
MAX41
MAX42
ß6
ß7
ß8
B
MAX12
MAX13
MAX14
ß6
ß7
ß8
C
MAX08
MAX09
ß7
ß6
ß1
D
MAX25
MAX53
MAX75
ß1
ß6
ß7
E
MAX57
MAX83
C-ter helix
hydrophobic
interactions
F
MAX29
MAX64
ß6
C-ter helix
G
MAX32
MAX90
ß1ß2 loop
C-ter helix
H
N-ter MAX
MAX52
C-ter MAX
N-ter and C-ter
MAX superimposition
I
MAX67
MAX80
MAX91
ß1
ß6
J
MAX10
MAX11
ß5ß6 helical turn
K
MAX47
MAX92
disulfide bond "4"
L
MAX34
MAX48
ß2
ß4ß5

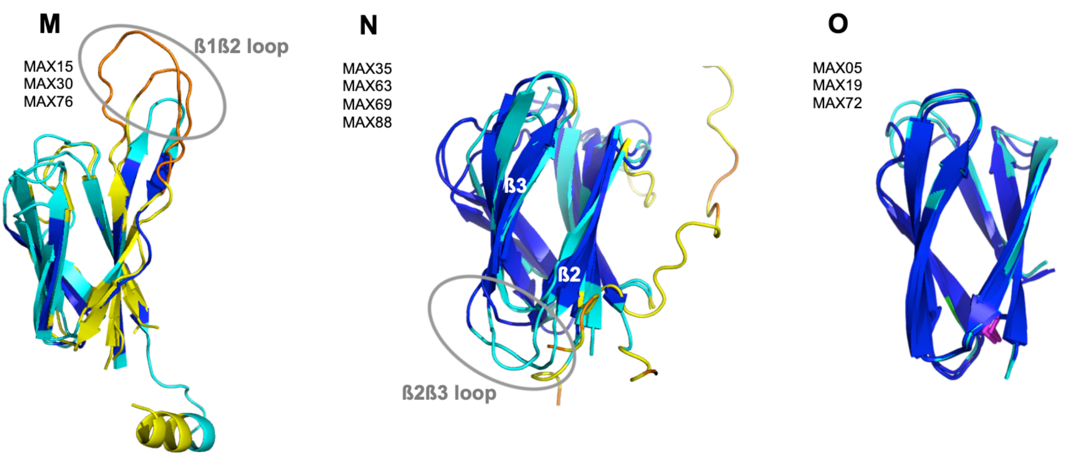


***S3 Fig. Groups of AlphaFold models***

*Superimposition of MAX models belonging to each of the groups that are labeled from A to O as indicated in Table 1.*

*The two groups **A** and **B** have additional strands flanking one side of the ß2ß1ß6 sheet. The group A specificity is the presence of C-terminal extensions after ß8 that are in helical conformations but differ in their lengths and relative orientations. The two groups **C** and **D** possess an additional ß7 strand that extends the first ß-sheet beyond ß6. The group C models have lower accuracy (pLDDT MAX score slightly above the 60 cutoff) than group D models.*

*Various inter-residue stabilizing interactions between the helical C-terminal extensions are observed for the groups **E** and **F**. Hydrophobic interactions between a conserved tryptophan residue in ß6 and hydrophobic residues in the helix are found in the group E models. Similarly, in the group F the position of the C-terminal helix is well-defined according to specific interactions taking place between residues in ß6 and the helix (phenylalanine stacking or a disulfide bond). For group **G** models the orientation of the C-terminal helices are more variable and display low pLDDT scores. A unique feature of the MAX family is the MAX52, whose ALPHAFOLD model consists of two MAX core modules (group **H**). The two MAX domains of MAX52 are structurally similar as illustrated by the snapshot of their superimposition (TM-score of 0.87 and DALI Z-score of 11.7 with 1.1 Å r.m.s.d and 38 % sequence identity).*

*Group **I** MAX structures have long ß1 flanked by similarly long ß2 or ß6 strands. Group **J** MAX structures have an helical turn in the loop joining ß5 to ß6. Group **K** MAX structures have a C-terminal extension with the disulfide bond "4". Group **L** MAX structures have a badly defined ß2 strand and stabilizing interaction between ß4 and ß5 strands by a disulfide bond (in magenta) or hydrogen bond (in green). The groups **M** and **N** MAX structures have long loops joining ß1 to ß2 and ß2 to ß3, respectively. Group **O** MAX structures are very similar to AVR-Pib structure but have the conserved disulfide bond "1".*
