## Supplementary material for "The structural landscape and diversity of *Pyricularia oryzae* MAX effectors revisited": Figure_S4

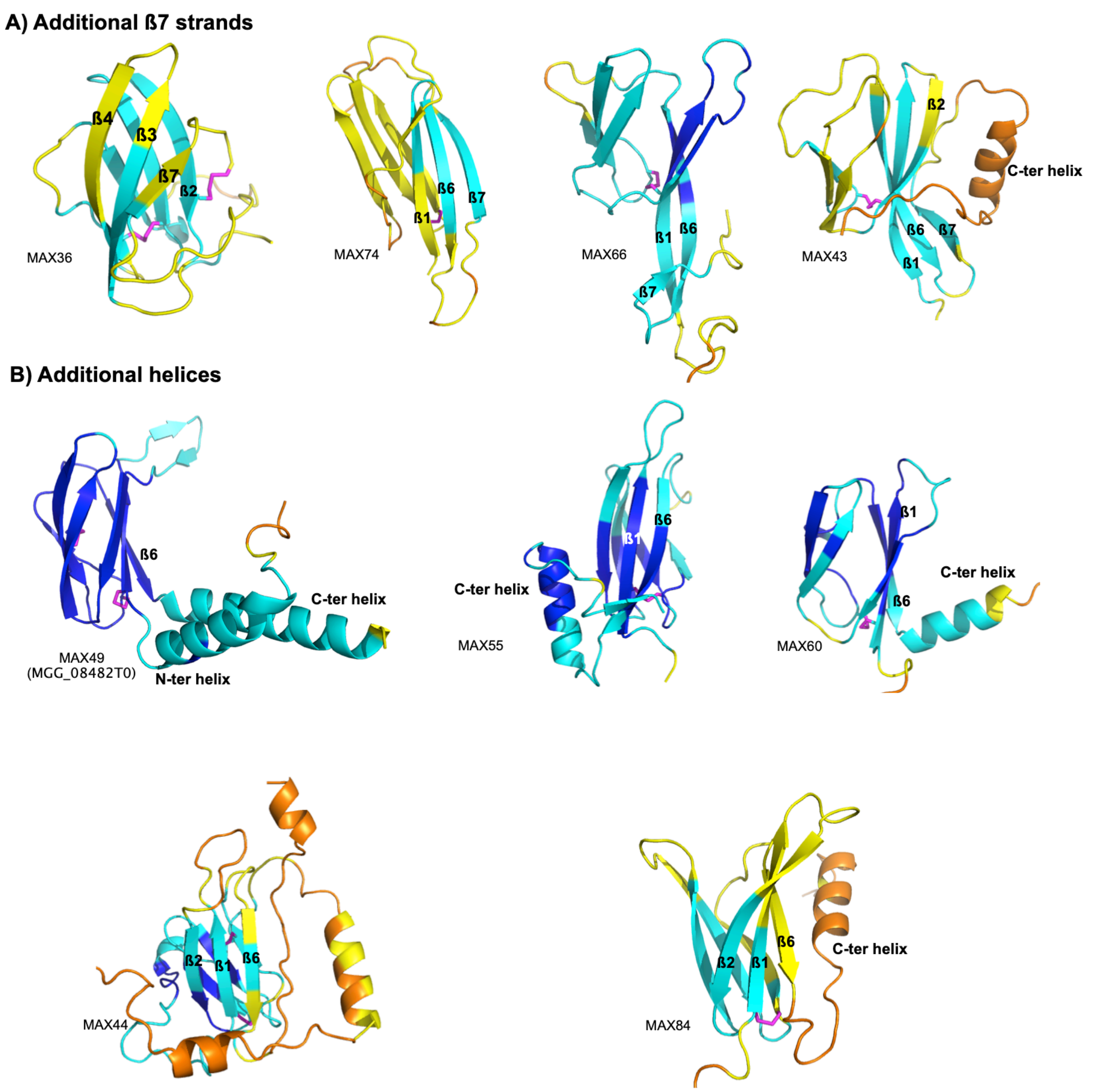


***S4 Fig. Singletons MAX effectors with structured extensions***

*A) MAX effectors with ß7 additional strand. For MAX36 the ß7 strand runs parallel to ß3 and the C-terminal cysteine residue makes a disulfide bond with a cysteine residue in ß2.*

*B) MAX effectors with additional helices. In the case of MAX49 the N- and C-terminal helices are packed against each other. C-terminal helices of MAX55 and MAX60 are well defined in the AlphaFold models while they have low pLDDT scores in MAX44 and MAX84 models.*
