## Supplementary material for "The structural landscape and diversity of *Pyricularia oryzae* MAX effectors revisited": Figure_S5

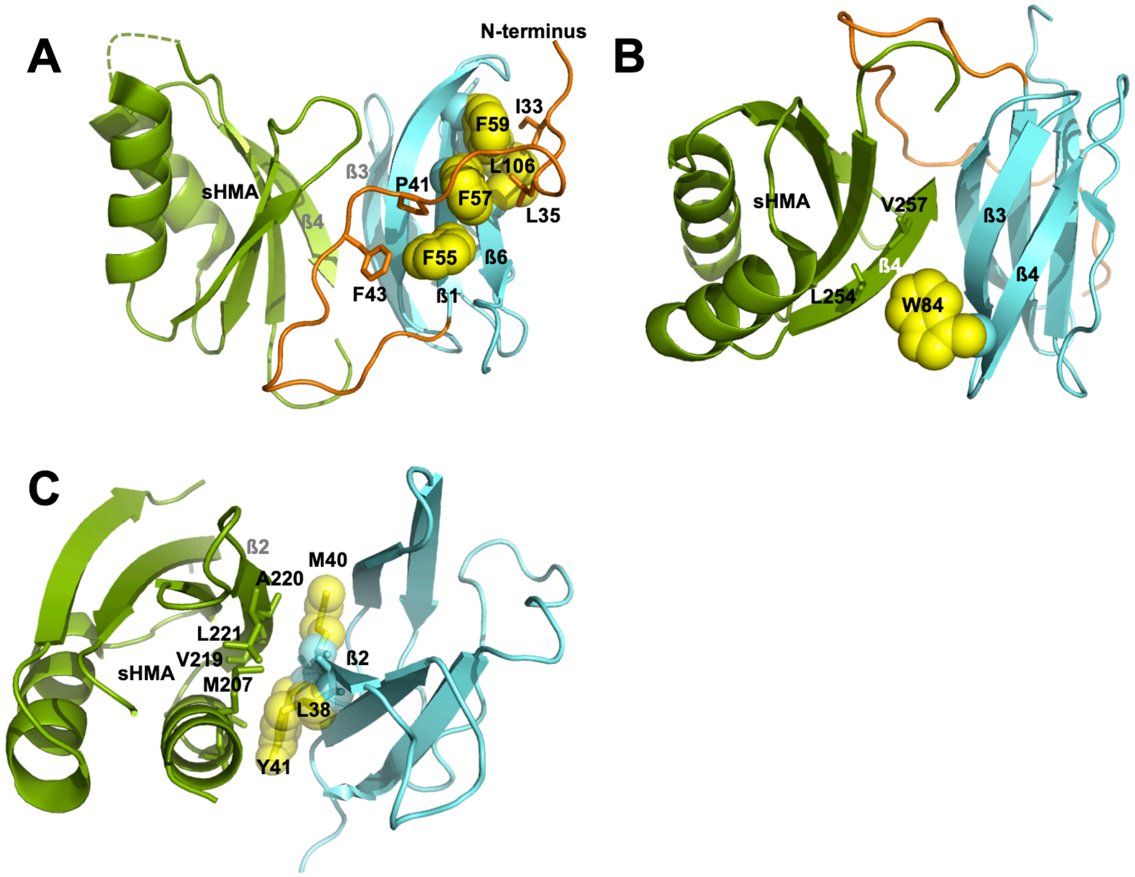


***S5 Fig. Hydrophobic residues on the surface of AVR-Pik and AVR-Pia.***

*(A) and (B) AVR-PikD is in cyan and sHMA-Pikp1 is in green, from the complex structure (5A6W). The main backbone – backbone interaction involves hydrogen bonds between ß3 strand of the AVR-PikC and the ß4 strand of the sHMA-Pikp [1].*

*(A) Hydrophobic residues in ß1 and ß6 of AVR-PikC are shown by yellow spheres for their side-chains with strictly conserved residues labeled, except F59 that is replaced by Y59 in 48% of the OG01, OG02 and OG03 sequences. The N-terminal stretch before the ß1 strand is shown in orange and residues that make hydrophobic interaction with the ß1-ß6 surface are labeled and strictly conserved excepted F43 that is replaced by V43 in 22% of the sequences.*

*(B) Hydrophobic W84 residue (replaced by F84 in 43% of the sequences) on the ß4 surface of AVR-PikC that makes interaction with L254 and V257 in the ß4 strand of sHMA-Pik.*

*(C) AVR-Pia is in cyan and sHMA-Pikp1 is in green, from the complex structure (6Q76). The main backbone – backbone interaction involves hydrogen bonds between ß2 strand of the AVR-Pia and the ß2 strand of the sHMA-Pikp [2]. Hydrophobic residues in ß2 of AVR-Pia are shown by yellow spheres for their side-*

*chains and by sticks for side-chains in ß2 of the sHMA-Pikp.*
