## Supplementary material for "The structural landscape and diversity of *Pyricularia oryzae* MAX effectors revisited": Figure_S6

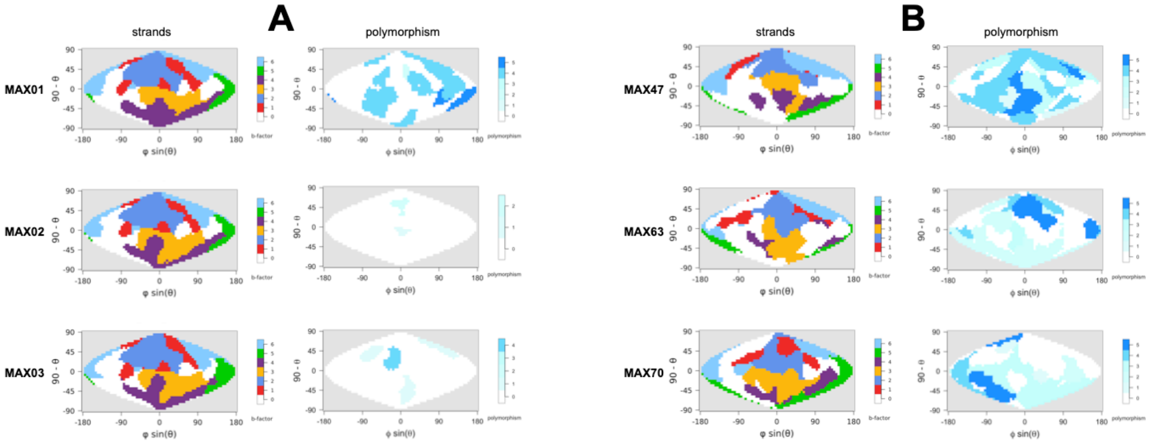


***S6 Fig. Amino-acid polymorphism mapped on the surface of AF MAX models***

*The color code for strands is the same as used in Fig 7. The polymorphism color scale was derived from the amino-acid conservation scores given for each OG cluster by coding from white (low polymorphism) for high conservation score of 9, and darker blue colors indicating highly polymorphic positions with conservation scores of 4 and below. A) The amino-acid polymorphism for AVR-Pik group including MAX01, MAX02 (AVR-PikC) and MAX03. B) The surface of the three MAX effectors displaying high polymorphism.*
