## Supplementary material for "The structural landscape and diversity of *Pyricularia oryzae* MAX effectors revisited": S4_Table

### S4a Table: Refinement statistics of MAX effector NMR structures

| | MAX47 | MAX60 | MAX67 | MAX28 |
|---|---|---|---|---|
| **NMR distance and dihedral constraints** | | | | |
| Distance constraints | | | | |
| Total NOE | 1379 | 2115 | 1389 | 1269 |
| Intra-residue | 337 | 475 | 246 | 221 |
| Inter-residue | | | | |
| Sequential ($\|i-j\|$ = 1) | 413 | 556 | 376 | 350 |
| Medium-range ($\|i-j\|$ < 4) | 124 | 393 | 164 | 159 |
| Long-range ($\|i-j\|$ > 5) | 505 | 691 | 603 | 539 |
| Hydrogen bonds | 56 | 64 | 64 | 48 |
| Disulfide bonds restraints | 6 | 3 | 3 | 3 |
| Total dihedral angle restraints | | | | |
| $\phi$ | 55 | 64 | 50 | 0 |
| $\psi$ | 55 | 64 | 50 | 0 |
| $\chi_1$ | 21 | 43 | 23 | 0 |
| **Structure statistics** | | | | |
| Violations (mean and s.d.) | | | | |
| Max. distance constraint violation (Å) | 0.16 ± 0.03 | 0.16 ± 0.02 | 0.12 ± 0.02 | 0.22 ± 0.04 |
| Max. dihedral angle violation (º) | 2.04 ± 0.90 | 2.38 ± 0.76 | 1.35 ± 0.42 | n. d. |
| Deviations from idealized geometry | | | | |
| Bond lengths (Å) | 0.0116 ± 0.0005 | 0.0117 ± 0.0003 | 0.0116 ± 0.0003 | 0.0118 ± 0.0003 |
| Bond angles (º) | 1.1135 ± 0.0316 | 1.2043 ± 0.0305 | 1.1140 ± 0.0287 | 1.1340 ± 0.0354 |
| Impropers (º) | 1.3765 ± 0.0865 | 1.3153 ± 0.0618 | 1.2634 ± 0.0786 | 1.4213 ± 0.0804 |
| **Ramachandran plot (%)** | | | | |
| Most favoured region | 82.8 | 86.1 | 89.5 | 79.0 |
| Additionally allowed region | 16.2 | 13.5 | 10.5 | 20.4 |
| Generously allowed region | 0.6 | 0.4 | 0.0 | 0.3 |
| Disallowed region | 0.4 | 0.0 | 0.0 | 0.3 |
| **Average pairwise** r.m.s.**d (Å)** (1) | | | | |
| Backbone atoms of 20 NMR conformers | 0.59 ± 0.12 | 0.42 ± 0.10 | 0.30 ± 0.08 | 0.51 ± 0.11 |
| Heavy atoms of 20 NMR conformers | 1.31 ± 0.19 | 0.95 ± 0.12 | 0.87 ± 0.13 | 0.94 ± 0.13 |

(1) Average pairwise root mean square deviation (r.m.s.d) between backbone atoms of the 20 best refined NMR conformers calculated for residues 42-100 (MAX47), 29-102 (MAX60), 22-76 (MAX67) and 38-99 (MAX28).

**S4b Table: AF model and NMR structure superimposition**

**MAX28:** AlphaFold model *versus* 20 NMR conformers

| **Average pairwise *r.m.s.* deviation** (Å)** | |
|---|---|
| Backbone | 1.42 ± 0.05 |
| Heavy | 2.84 ± 0.04 |

** " Pairwise r.m.s.d. calculated among 20 refined structures for residues 38-99 against the AlphaFold model."

**MAX47:** AlphaFold model *versus* 20 NMR conformers

| **Average pairwise *r.m.s.* deviation** (Å)** | |
|---|---|
| Backbone | 1.35 ± 0.11 |
| Heavy | 2.17 ± 0.10 |

** " Pairwise r.m.s.d. calculated among 20 refined structures for residues 42-100 against the AlphaFold model."

**MAX60:** AlphaFold model *versus* 20 NMR conformers

| **Average pairwise *r.m.s.* deviation** (Å)** | |
|---|---|
| Backbone | 1.11 ± 0.07 |
| Heavy | 2.06 ± 0.07 |

** " Pairwise r.m.s.d. calculated among 20 refined structures for residues 29-102 against the AlphaFold model."

**MAX67:** AlphaFold model *versus* 20 NMR conformers

| **Average pairwise *r.m.s.* deviation** (Å)** | |
|---|---|
| Backbone | 0.99 ± 0.04 |
| Heavy | 2.07 ± 0.04 |

** " Pairwise r.m.s.d. calculated among 20 refined structures for residues 22-76 against the AlphaFold model."
