## Supplementary_Materials_and_Methods for "The structural landscape and diversity of *Pyricularia oryzae* MAX effectors revisited"

### *Crystallographic structure determination of MoToxB*

The structure was determined at 1,38 Å resolution by molecular replacement.

***Protein expression and purification.*** The sequence for the mature protein (residues 21-90 for MoToxB, UniProt L7J9X7: identical to the *P. oryzae* gene strain Br58_090910T0 or the TH16_EuGene_00135161 gene in GEMO database [1]) was inserted into the pET-SP vector [1,2]. The plasmid pET-SP-MoToxB was used to transform E. coli BL21 (DE3) cells (Invitrogen). Bacteria were grown on auto inducing minimal media C-750501 [3] containing ampicillin 100 µg/mL. Cells were grown with shaking at 200 rpm at 30˚C for 24h. To generate isotopically labeled samples for NMR spectroscopy, we used $^{15}NH_4Cl$ as the primary nitrogen. Cells were harvested by centrifugation 5000 x g for 20 minutes using Thermo Scientific SORVALL LYNX 6000 and re-suspended in lysis buffer (200mM TrisHCl pH8, 200mM Sucrose, 0.05mM EDTA, 50µM lysozyme). After 30 minutes incubation, cell debris were removed by centrifugation at 12 000 g for 30 min at 4°C. The His6-tagged proteins were isolated from the filtrate using a 5 mL Ni-NTA HisTrap HP (Healthcare Life Sciences) with an imidazole gradient from 5mM to 500 mM over 10 column volumes (CV). Fractions containing the protein were identified by SDS-PAGE gel analysis and pooled. The protein was further purified by gel filtration using a Superdex S75 26/60 (GE Healthcare) column in buffer A (20 mM TrisHCl, pH 8.0, 150 mM NaCl, 1mM DTT) and pure fractions were pooled. The protein was cleaved using TEV protease at 35 µg/ml during an overnight dialysis bath in buffer A at 4°C. The cleaved protein recovered after further gel filtration and verification of fractions by SDS PAGE was pooled. All chromatography steps were performed using an AKTA purifier (GE Healthcare Life Sciences).

**Crystallogenesis**. The purified protein was concentrated to 10 mg/ml in 20 mM TrisHCl, pH 8.0, 150 mM NaCl, 1mM DTT. Crystallization trials were performed at 25°C using the hanging-drop vapor-diffusion method in 96 microplates and Mosquito HTS robot with 100 nl of protein mixed with 100 nl of reservoir. After 3 weeks, crystals were obtained from condition 0,1M Hepes pH 7.5, 1.6M Ammonium Sulfate,

1,6M PEG 1000. Crystals were cryo-protected with oil (Paratone, Hampton research), and freezed in liquid $N_2$.

***X-ray Data collection, processing and structure determination***.

X-ray diffraction data were collected on a Pixel detector (PILATUS3 2M) by the autonomous ESRF beamline MASSIF-1 [4] using automatic protocols for the location and optimal centering of crystals. The beam diameter was selected automatically to match the crystal volume of highest homogeneous quality. Strategy calculations accounted for flux and crystal volume in the parameter prediction for complete data sets. Data were auto-processed by XDS [5], AIMLESS [6], from the CCP4/CCP4i programs suite [7,8]. *MotoxB* crystals belong to the *C*121 space group and contain two molecules in the asymmetric unit. The structure was determined at 1,38 Å resolution by Molecular replacement using PHASER from PHENIX (Table 1) and a preliminary NMR model (see below). After model building using ARP/WARP [9], COOT [10] and refinement by REFINE from PHENIX [11] the final structure had a $R(\%)/R(\%)_{free}$ ratio of 0.143 / 0.171.

***Preliminary Solution Structure of MoToxB used for molecular replacement.***

The $^{15}$N-labeled protein sample was dialyzed against 20 mM Na-Phosphate buffer (pH 6.8), 150 mM NaCl and 1 mM DTT, overnight at 4°C. The sample was concentrated to 760 µM in 220 µL volume with 5% $D_2O$ for the lock. NMR experiments were carried out at 10°C on a Bruker Avance III 700 MHz spectrometer, equipped with 5 mm z-gradient TCI cryoprobe. Backbone and side chains $^1$H and $^{15}$N resonance assignments were obtained through 3D [$^1$H,$^{15}$N] NOESY-HSQC (mixing time 150 ms) and TOCSY-HSQC (isotropic mixing: 50 ms) experiments. Water suppression was achieved with the WATERGATE sequence [12]. $^1$H chemical shifts were directly referenced to the methyl resonance of DSS, while $^{15}$N chemical shifts were referenced indirectly to the absolute $^{15}$N/$^1$H frequency ratio. All NMR spectra were processed with Topspin 3.6 (Bruker) and analyzed with Cindy 2.1 (Padilla, www.cbs.cnrs.fr). Assignments of amide $^1$H, $^{15}$N chemical shifts are given in Figure 1. NOE cross-peaks identified on the 3D [$^1$H, $^{15}$N] NOESY-HSQC experiment were assigned through several runs of automated NMR structure calculations with CYANA 3 [13,14]. For the final list of restraints, distance values redundant with covalent geometry were eliminated and two disulfide bonds that were consistent with short distances between cysteine residues Cys25-Cys66 and Cys41-Cys86 were added. A total of 200 three-dimensional structures were generated using the torsion angle dynamics protocol of CYANA 3 from the NOEs and 2 disulfide bond restraints. The best structure (based on the final target penalty function value) was used for molecular replacement.

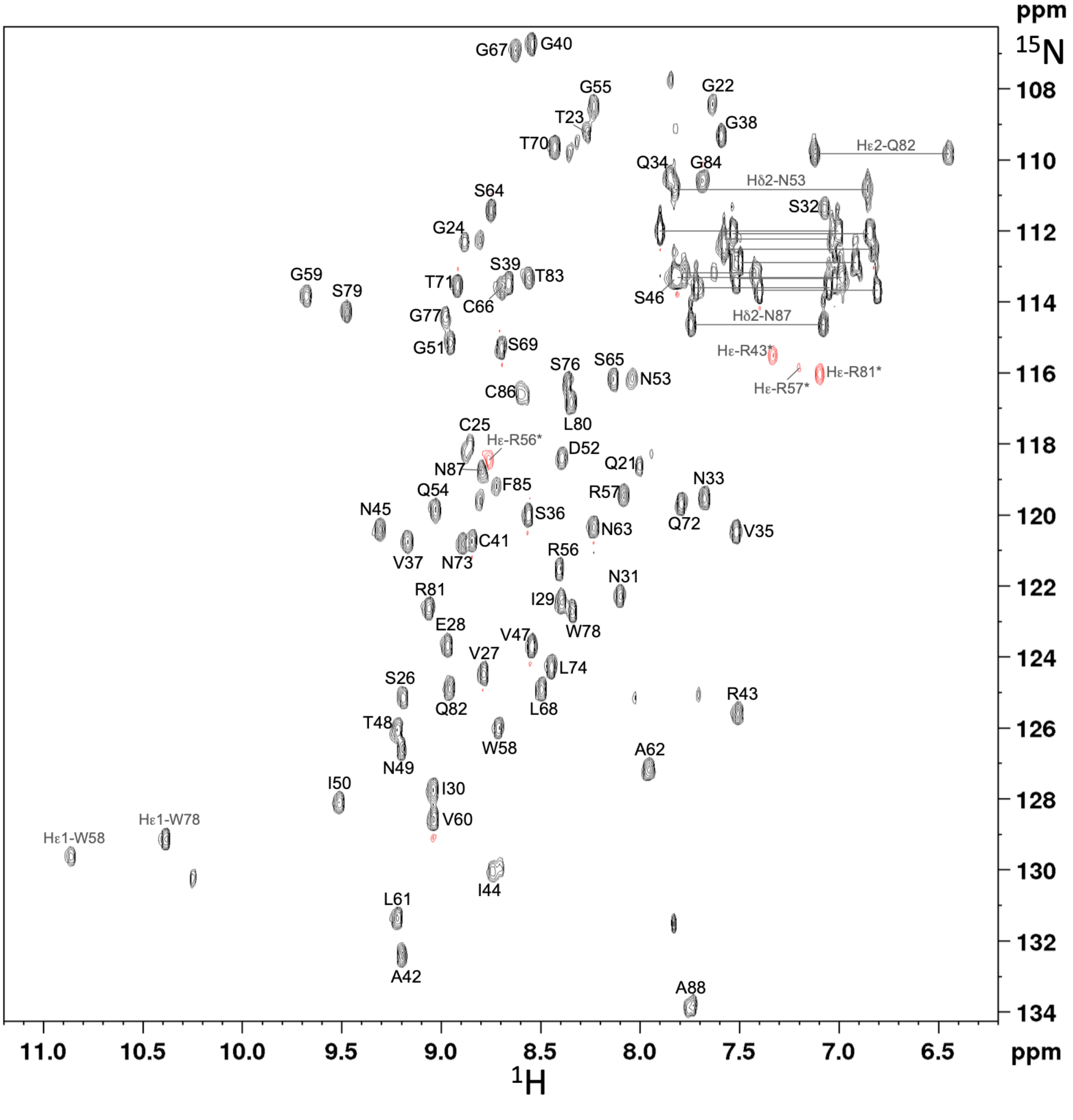


**Figure 1.** [$^1$H, $^{15}$N] HSQC of $^{15}$N-labeled MoToxB (0.76 mM) in 20 mM Na-phosphate pH 6.8, 150 mM NaCl, 1 mM DTT recorded at 283K and at 700 MHz.

**Table 1. X-ray data collection and refinement statistics.**

| | MoToxB |
|---|---|
| Wavelength | 0.996 |
| Resolution range (Å) | 34.59-1.38 (1.43-1.38) |
| Space group | *C*121 |
| Unit cell | |
| *a, b, c* (Å) | 73.341, 24.760, 68.613 |
| *α, β, γ* (°) | 90, 100.88, 90 |
| Total reflections | 59102 (2877) |
| Unique reflections | 23545 (1686) |
| Multiplicity | 2.5 (1.7) |
| Completeness (%) | 93.7 (70) |
| Mean I/sigma(I) | 14 (4,5) |
| Wilson B-factor (Å$^2$) | 12 |
| R-merge | 0.038 (0.11) |
| $CC_{1/2}$ | 0.997 (0.967) |
| Reflections used in refinement | 22356 |
| Reflections used for R-free | 1189 |
| R-work | 0.143 |
| R-free | 0.171 |
| Number of non-hydrogen atoms | 1118 |
| macromolecules | 970 |
| Heterogen atoms | 19 |
| solvent | 120 |
| Protein residues | 135 |
| RMS(bonds Å) | 0.0062 |
| RMS(angles °) | 0.98 |
| Ramachandran favored (%) | 100 |
| Ramachandran allowed(%) | 0 |
| Ramachandran outliers (%) | 0 |
| Rotamer outliers (%) | 0 |
| Clashscore | 3.06 |
| Average B-factor (Å$^2$) | 15.03 |
| macromolecules | 13.04 |
| solvent | 35.56 |

#$R_{merge} = S_{hkl}\, S_i\, |I_i(hkl) - (I(hkl))| / S_{hkl}\, S_i\, I_i(hkl)$, where $I_i(hkl)$ is the *i*th observation of reflection *hkl* and <*I(hkl)*> is the weighted average intensity for all observations of reflection *hkl*.
