## Supplementary figures and images for "The structural landscape and diversity of *Pyricularia oryzae* MAX effectors revisited"

### MAX01_polymorphism.png

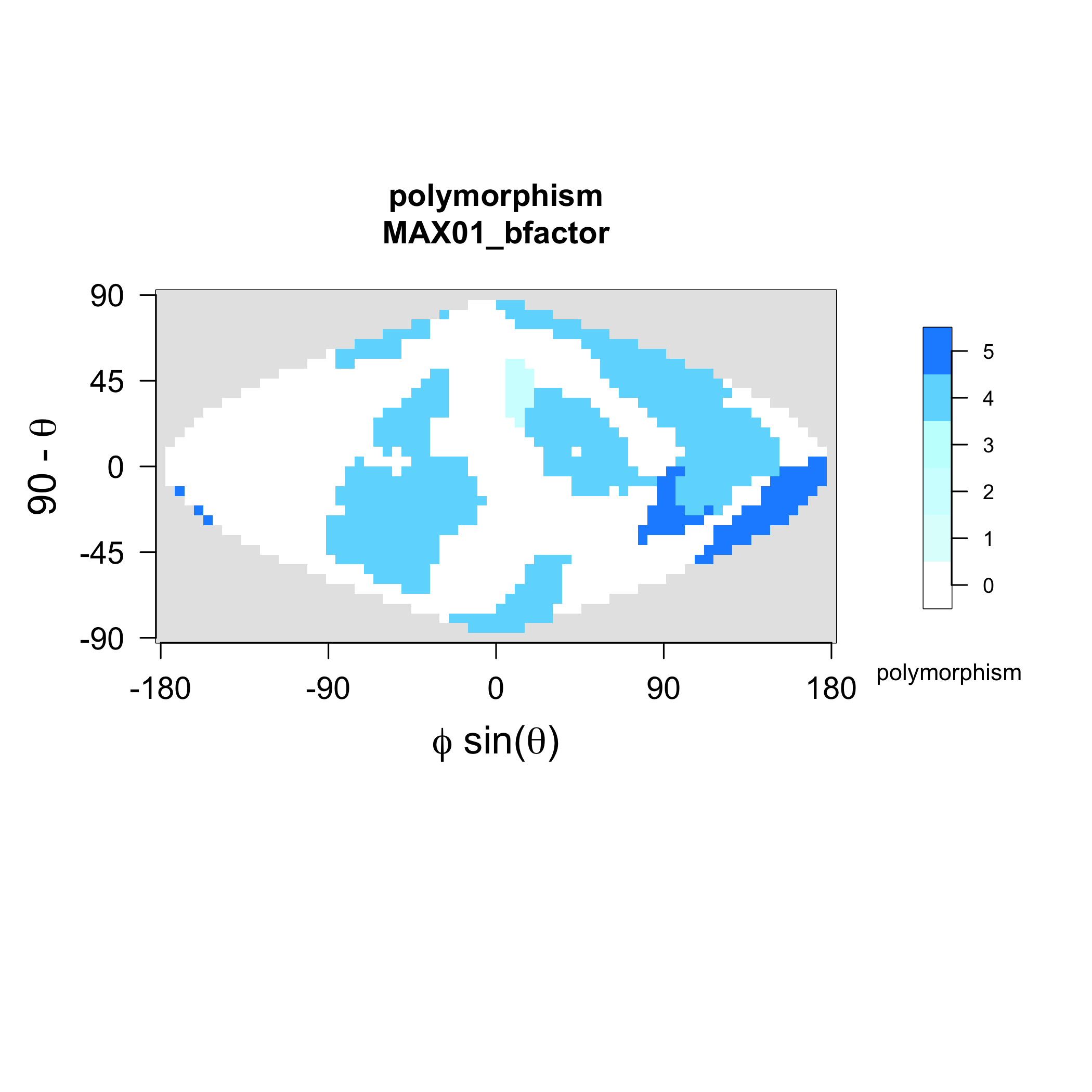

### MAX01_strands.png

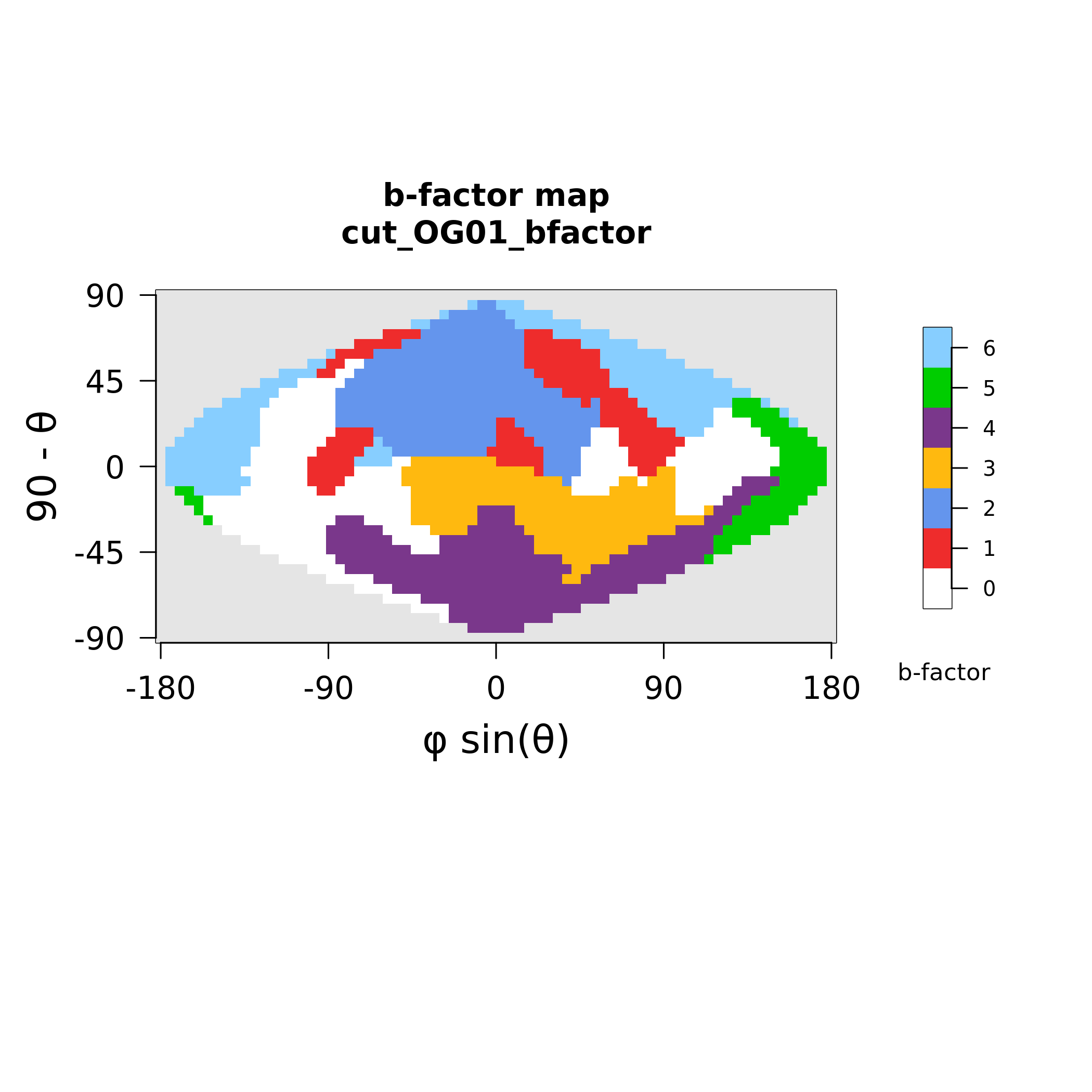

### MAX02_polymorphism.png

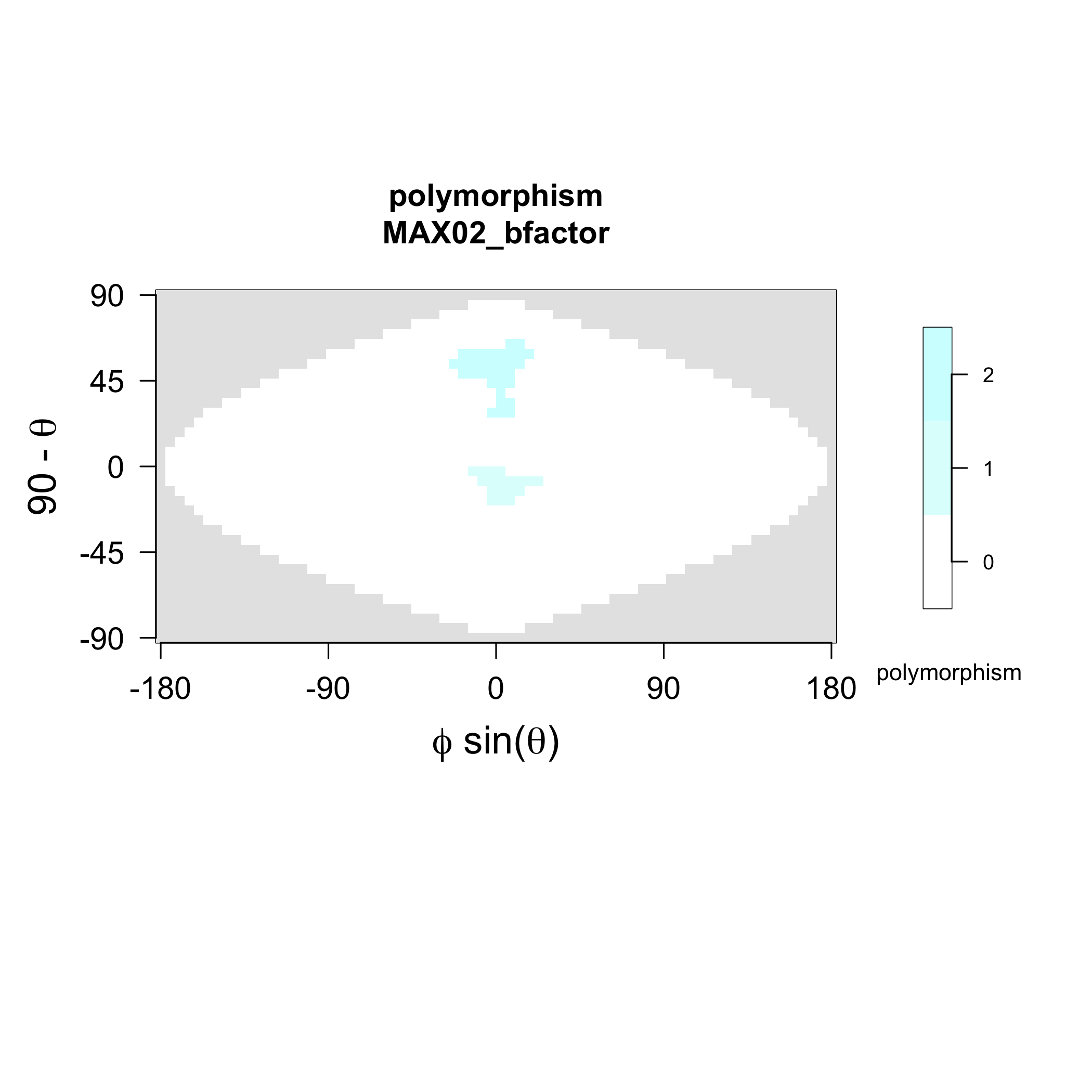

### MAX02_strands.png

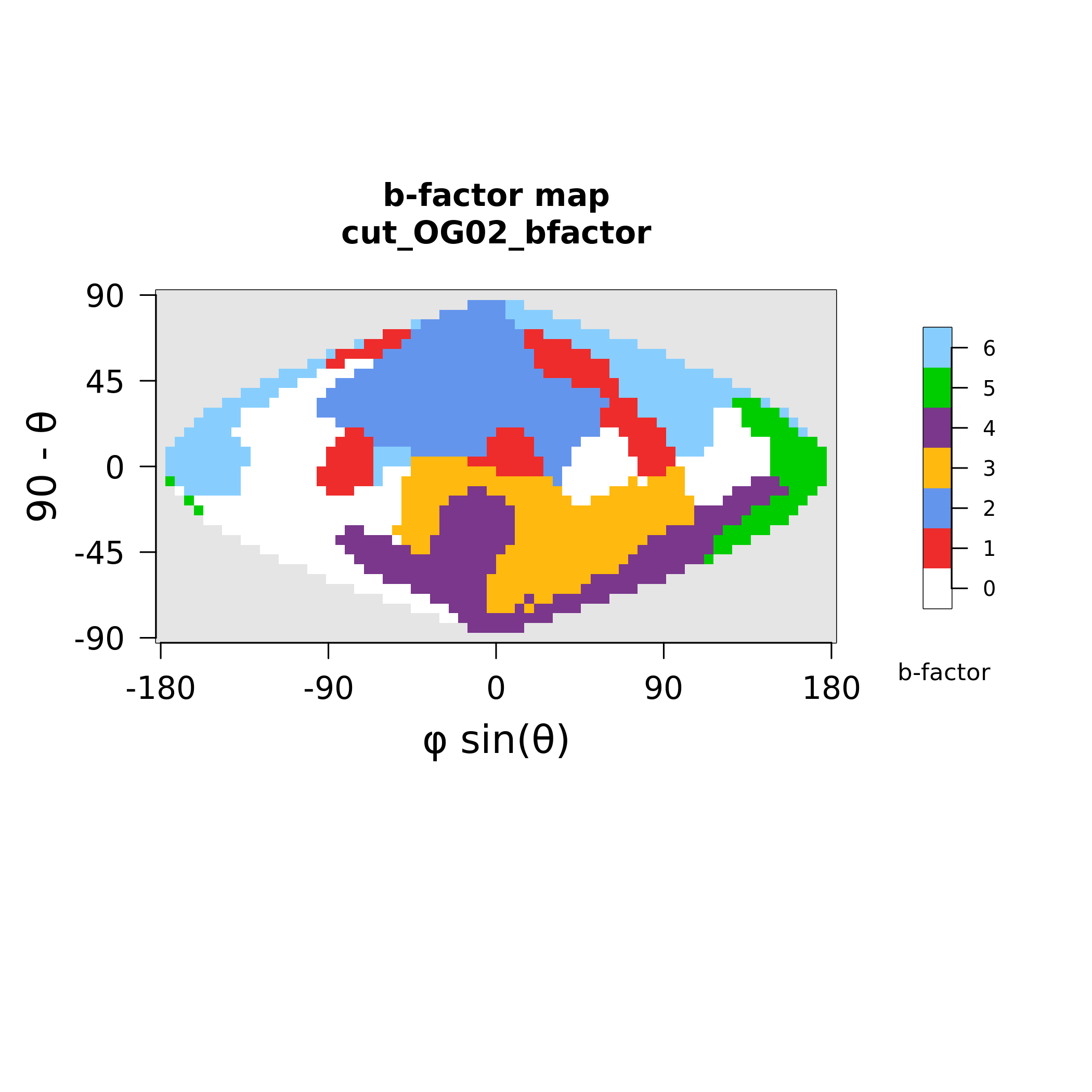

### MAX03_polymorphism.png

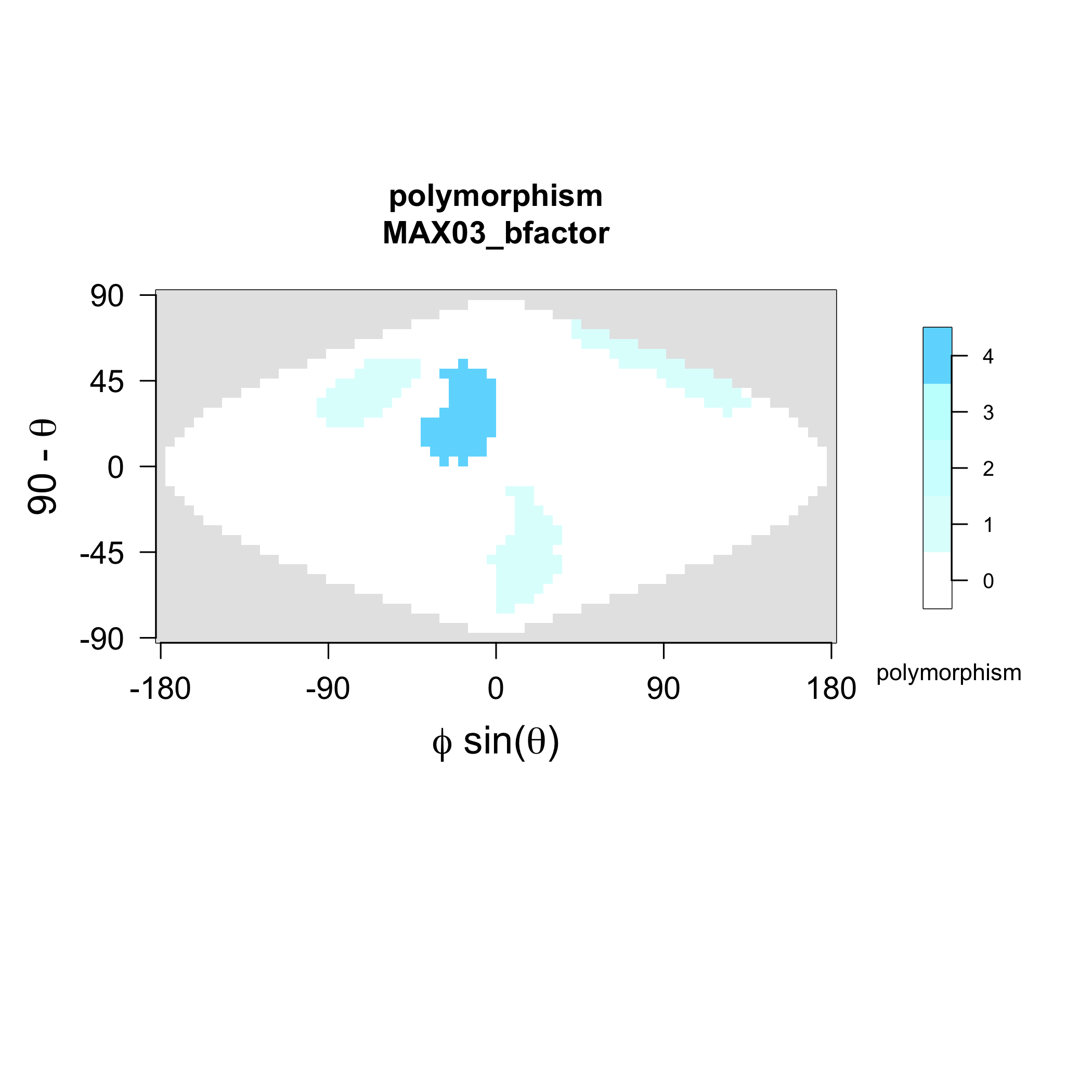

### MAX03_strands.png

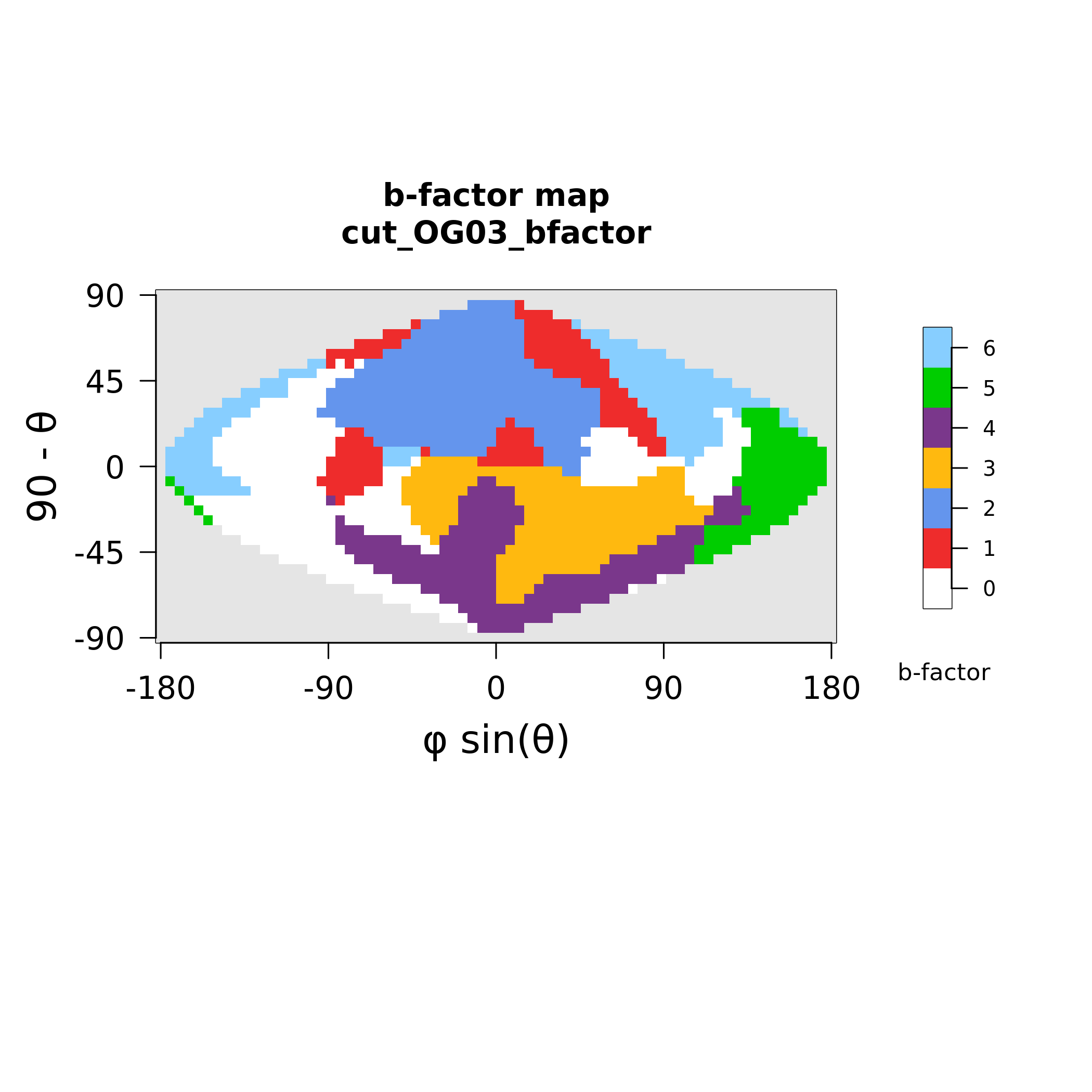

### MAX05_polymorphism.png

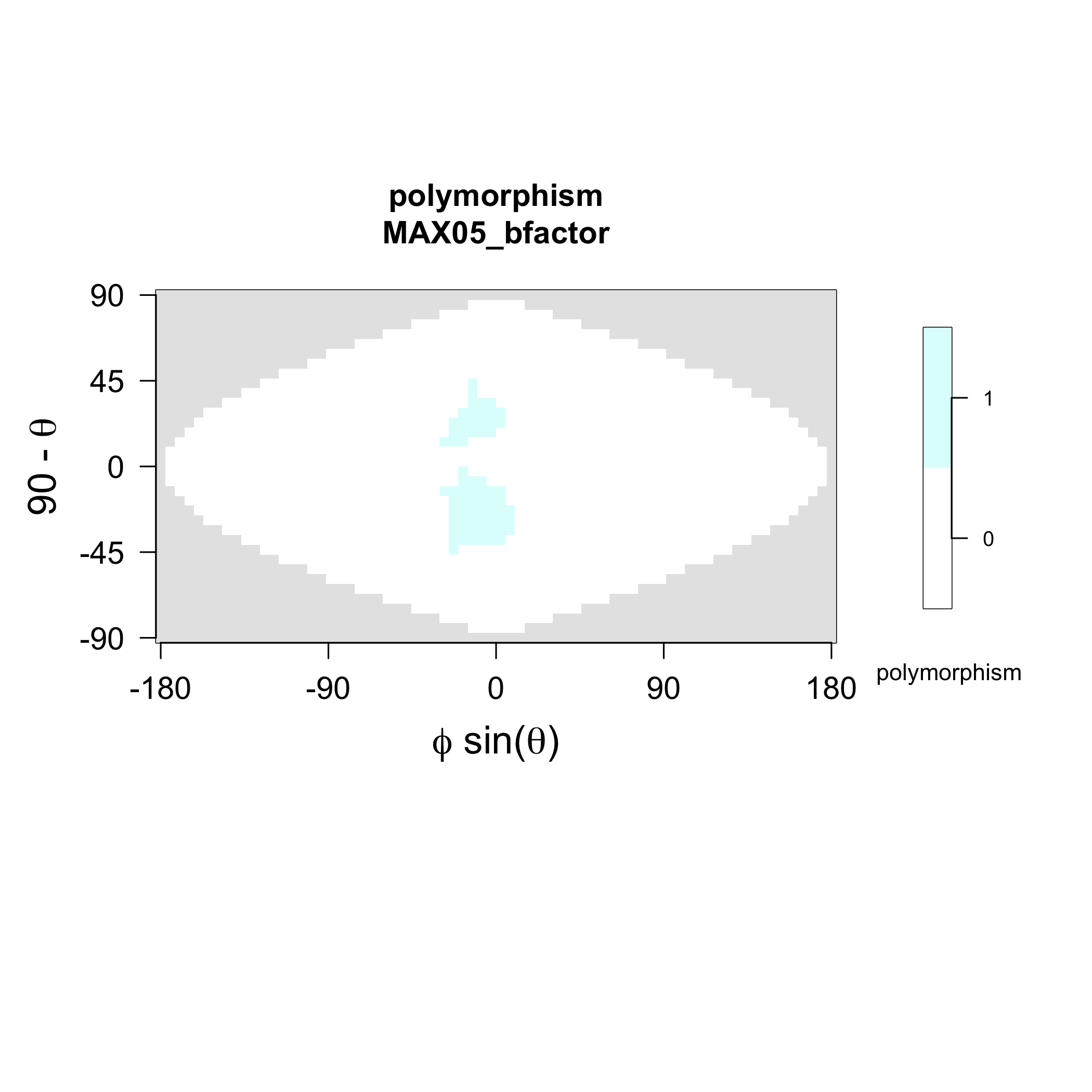

### MAX05_strands.png

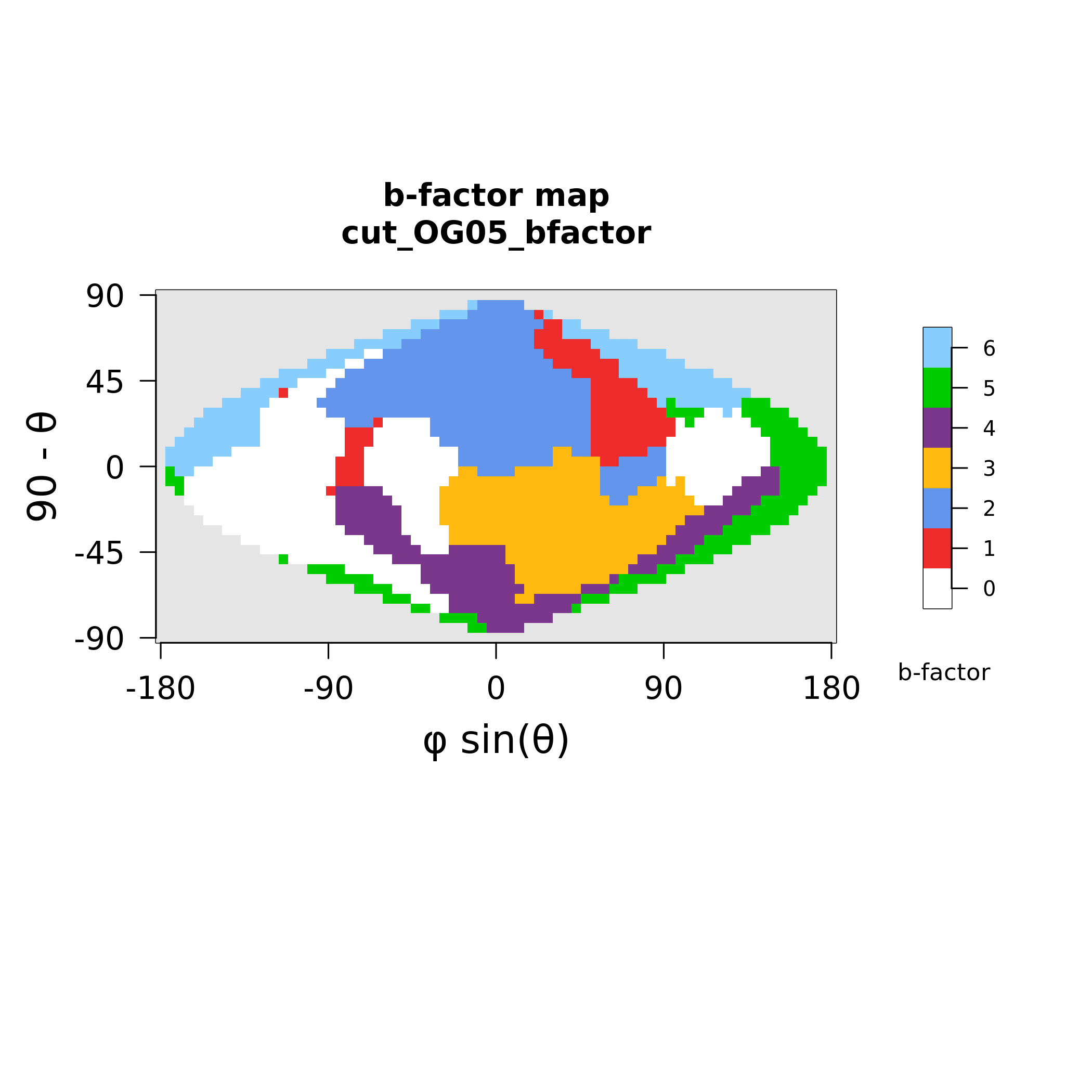

### MAX06_polymorphism.png

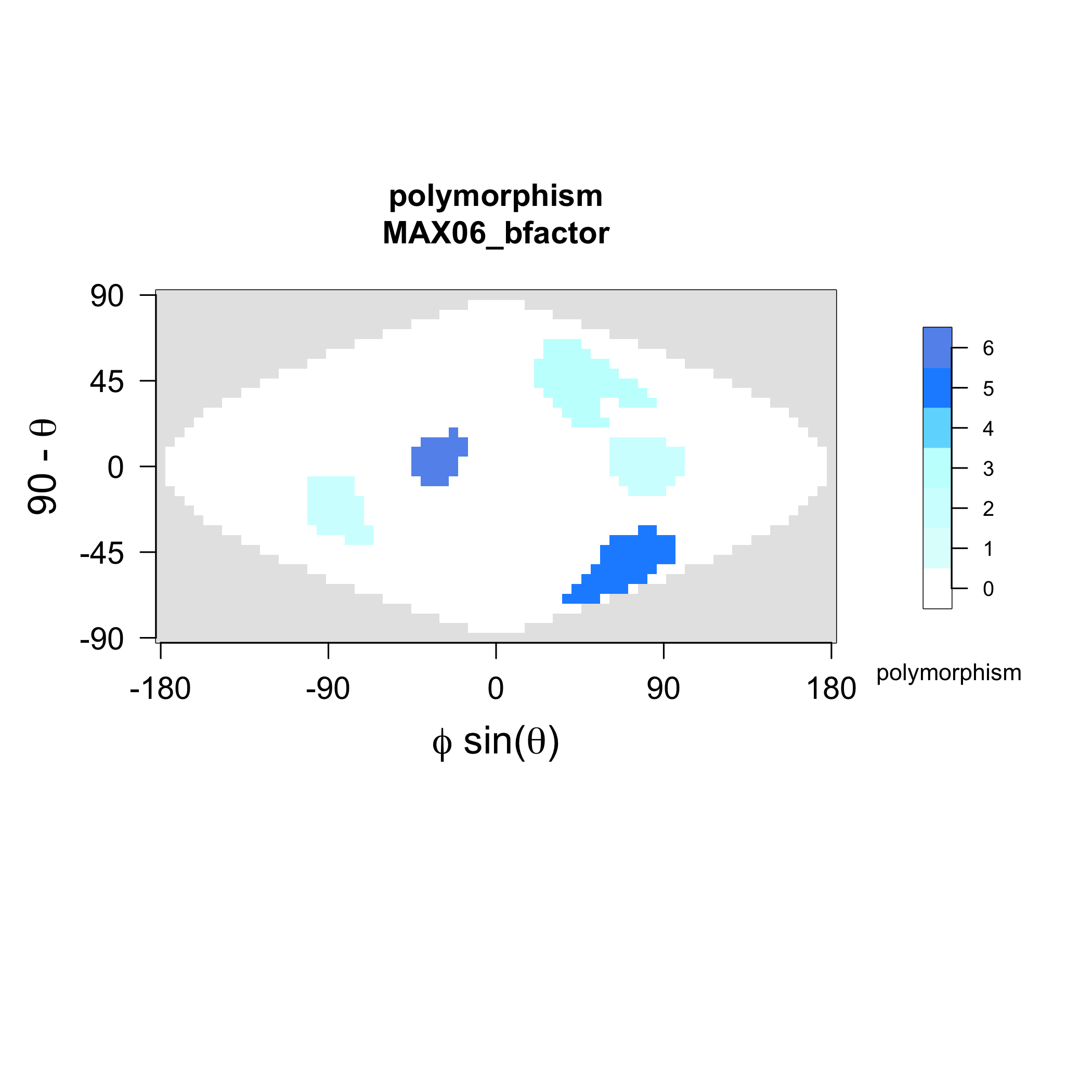

### MAX06_strands.png

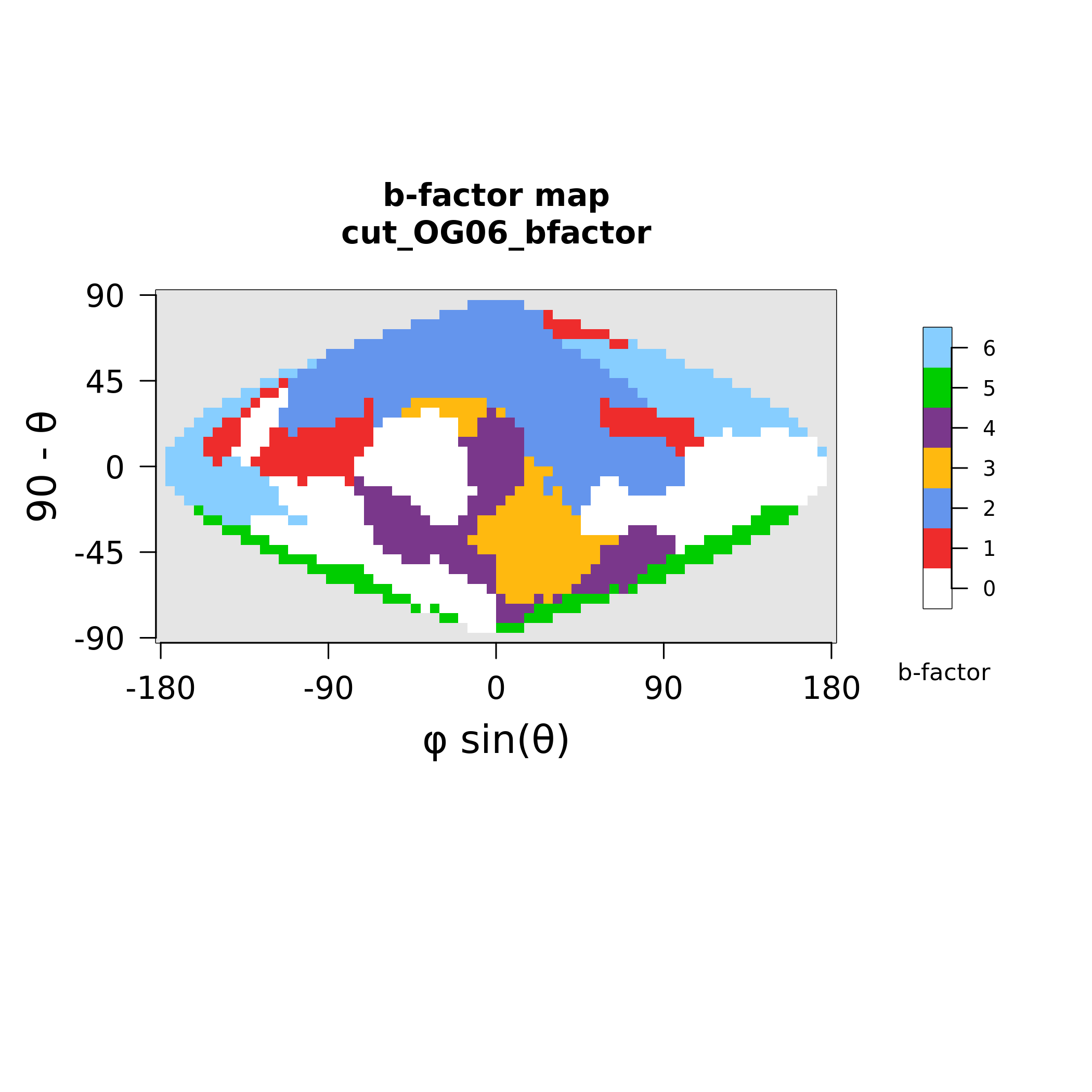

### MAX07_polymorphism.png

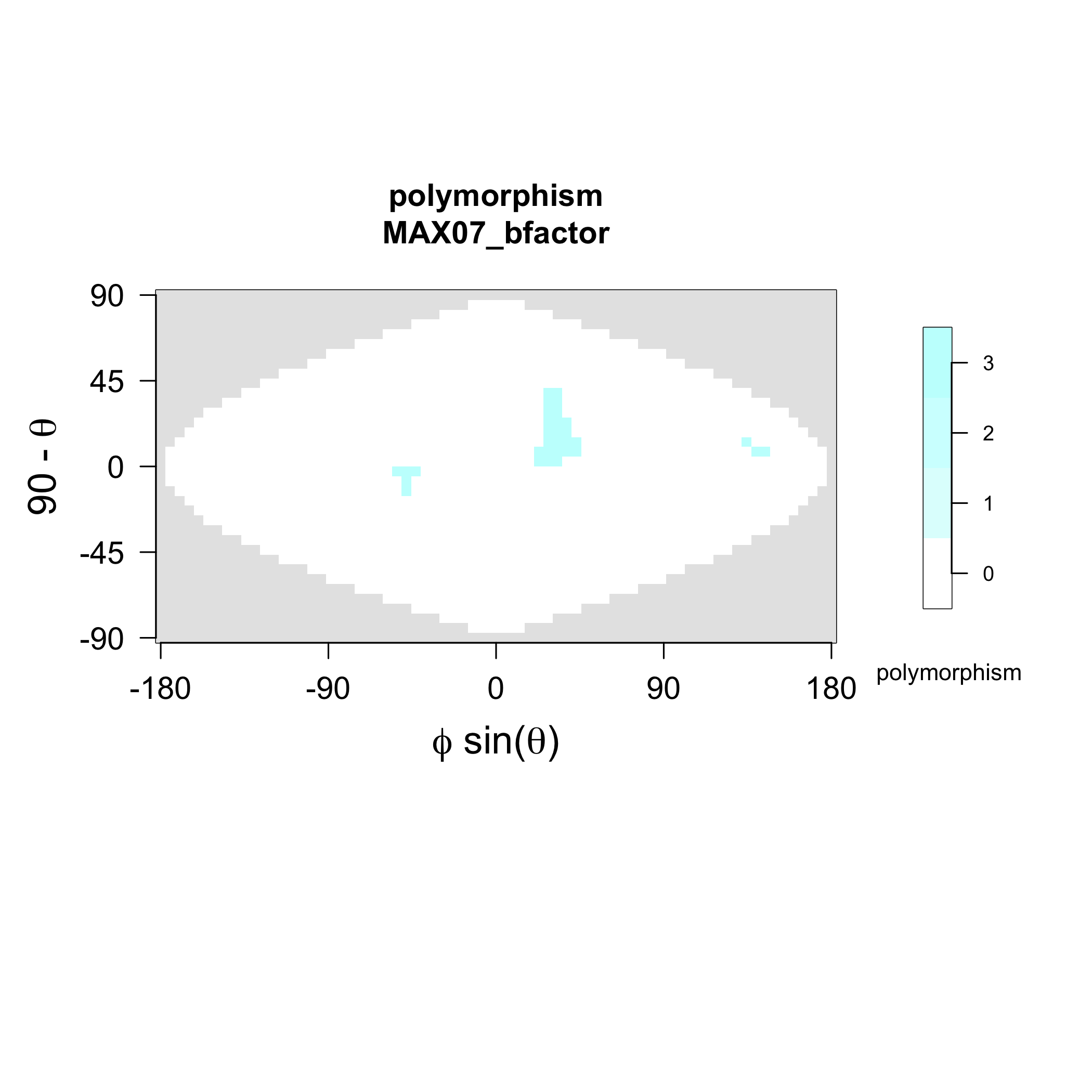

### MAX07_strands.png

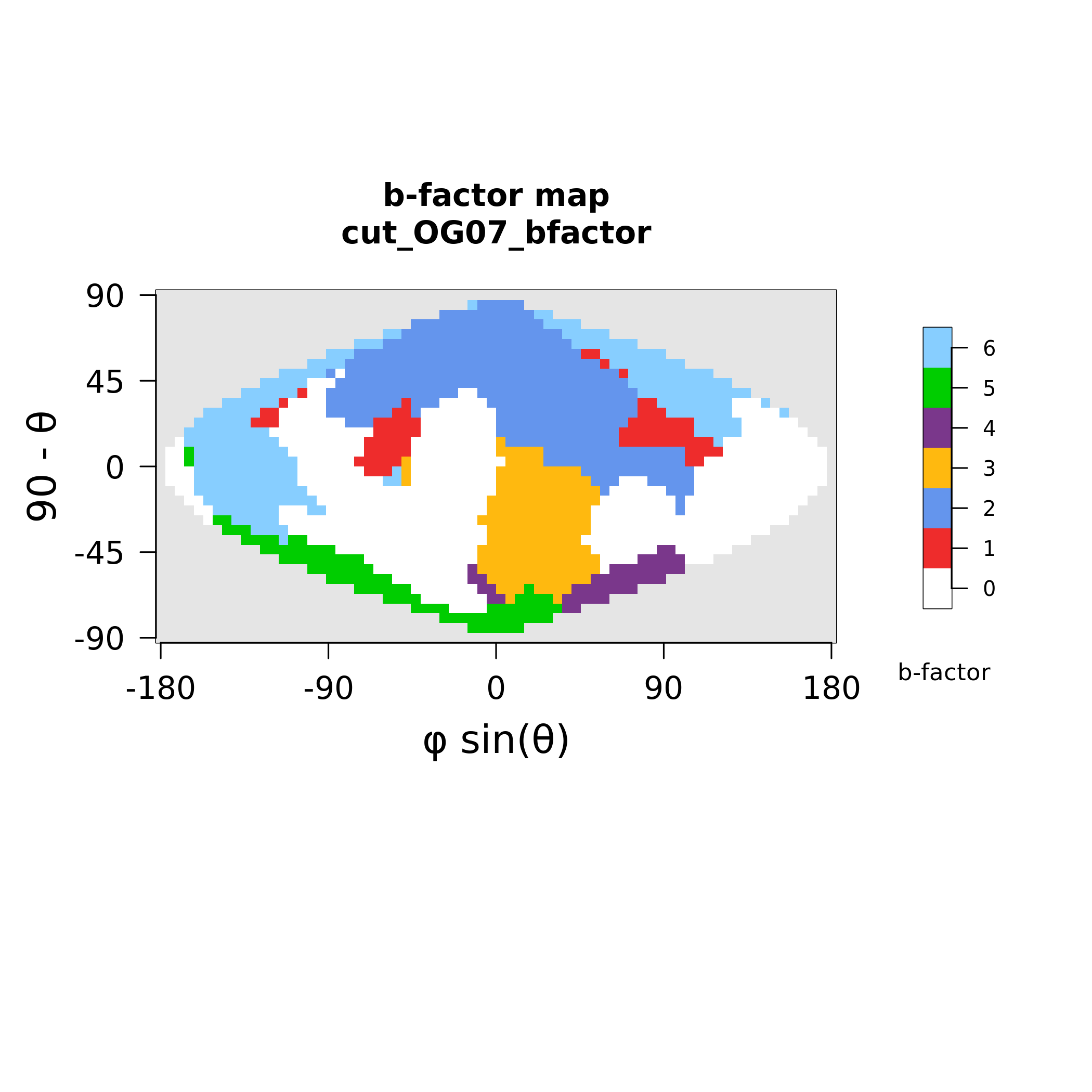

### MAX10_polymorphism.png

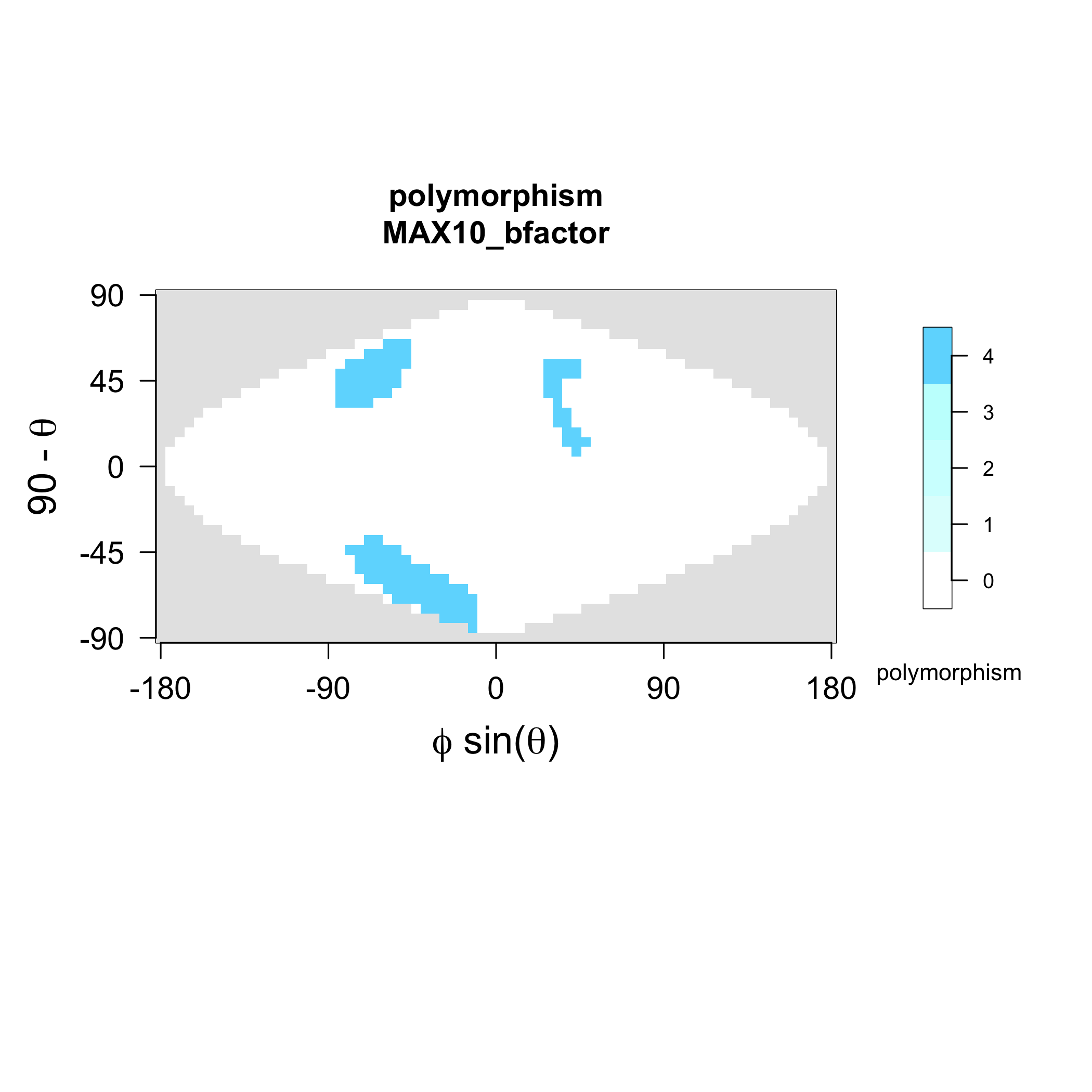

### MAX10_strands.png

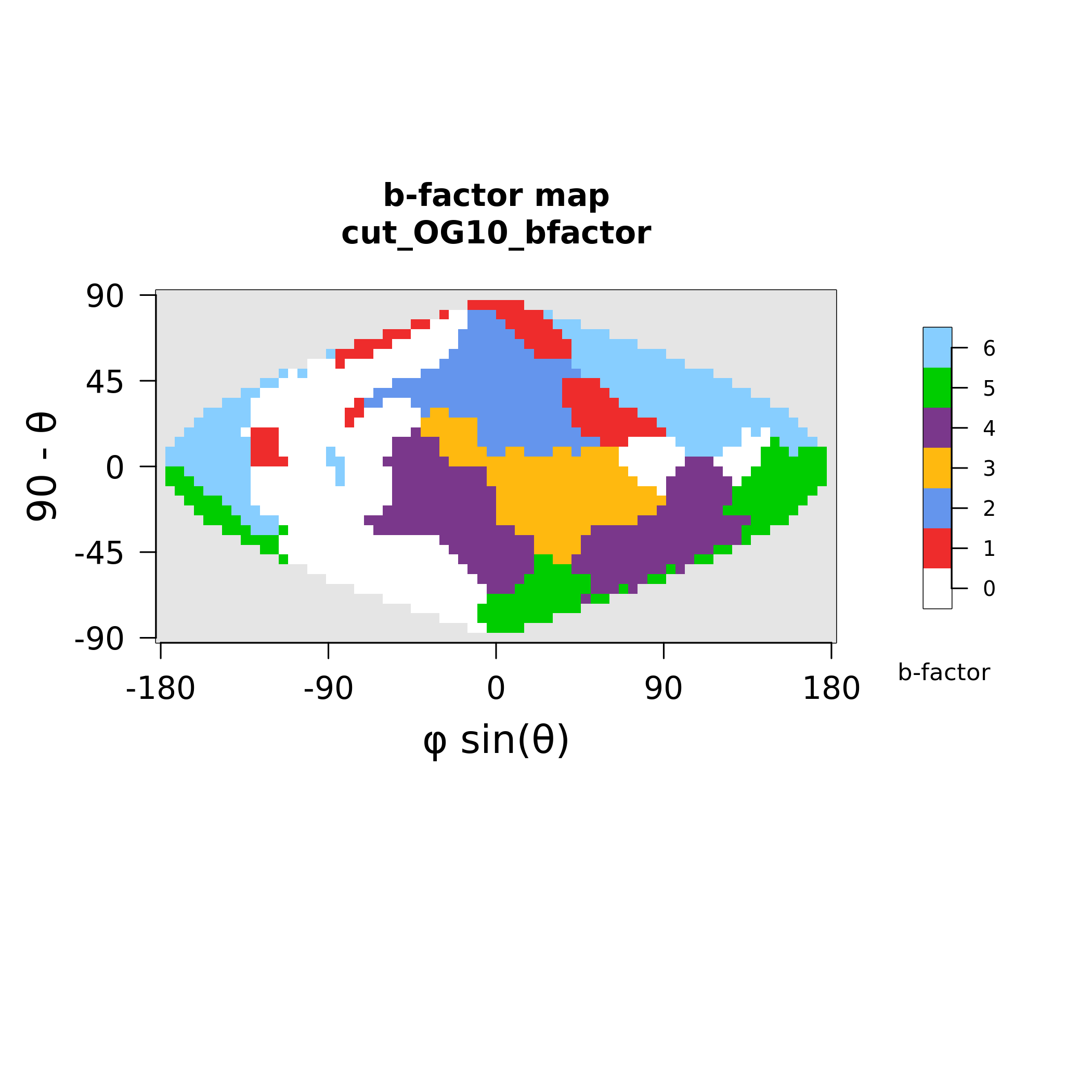

### MAX11_polymorphism.png

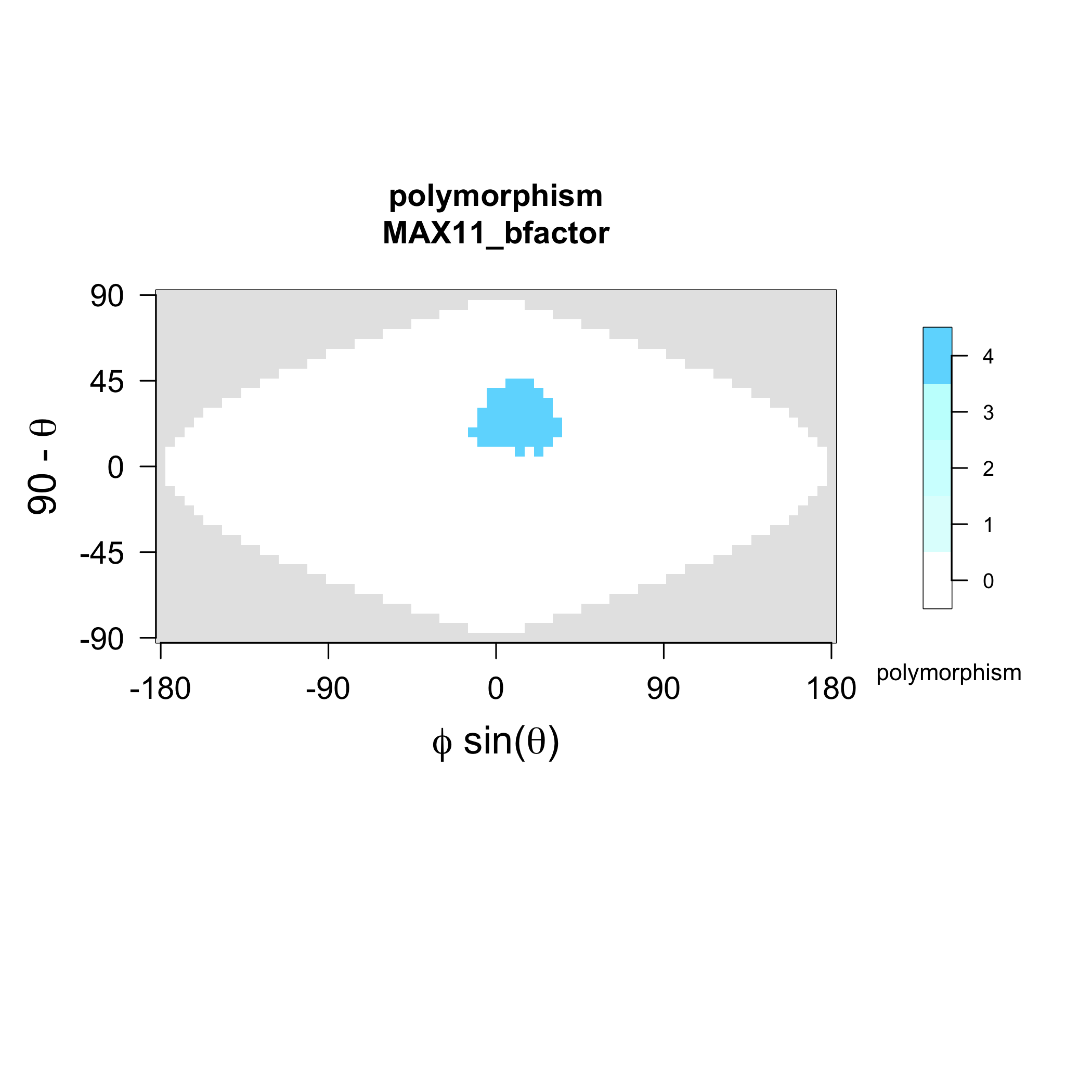

### MAX11_strands.png

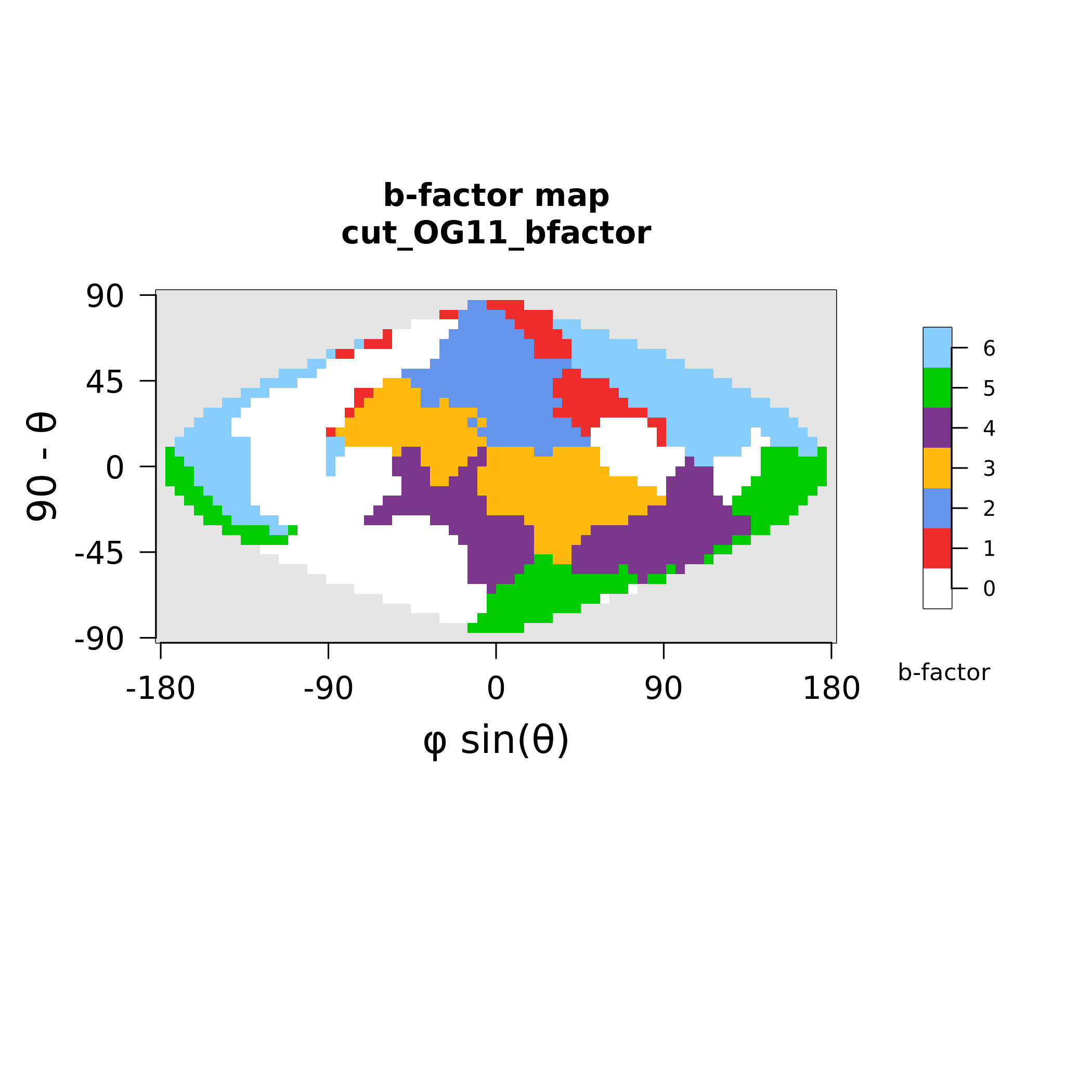

### MAX15_polymorphism.png

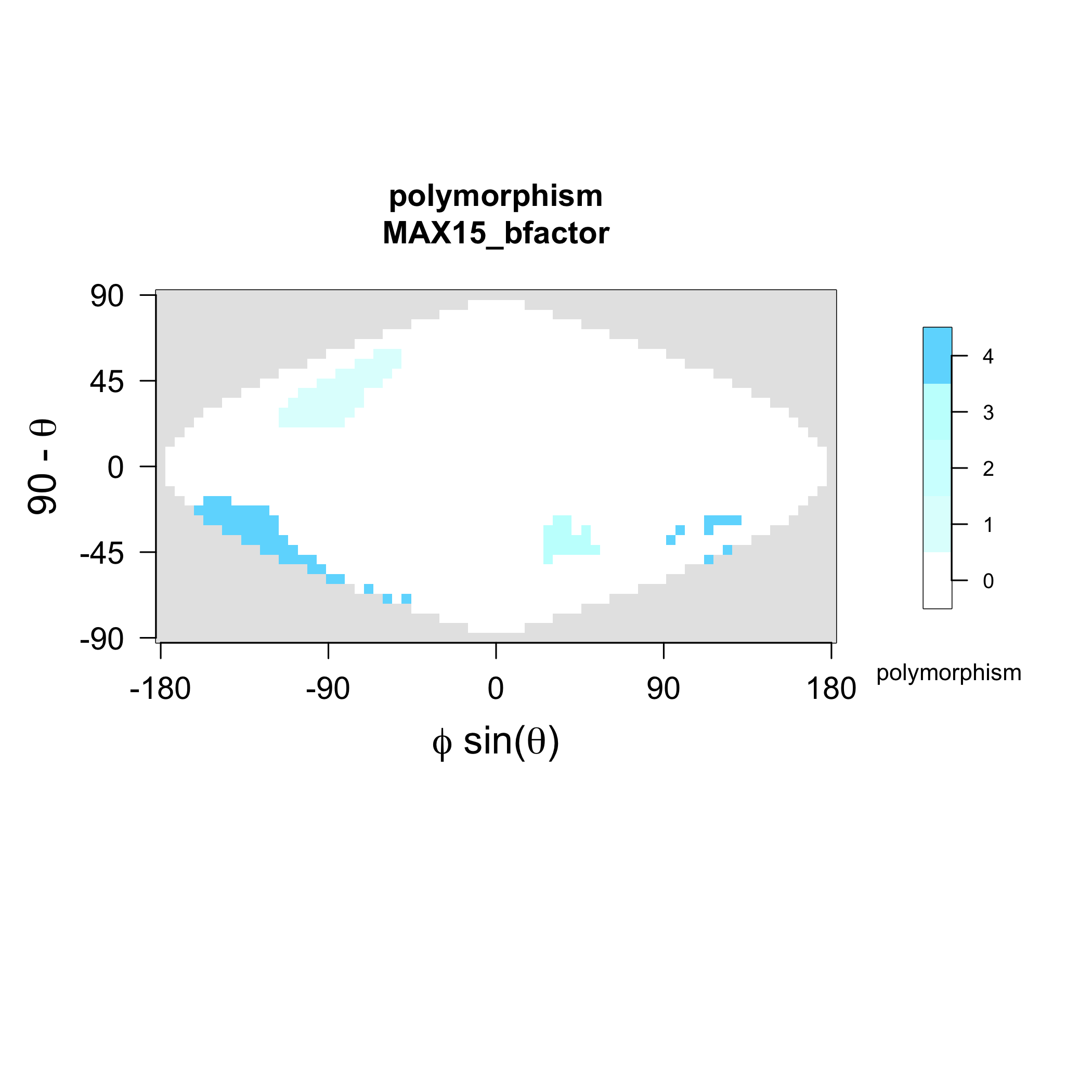

### MAX15_strands.png

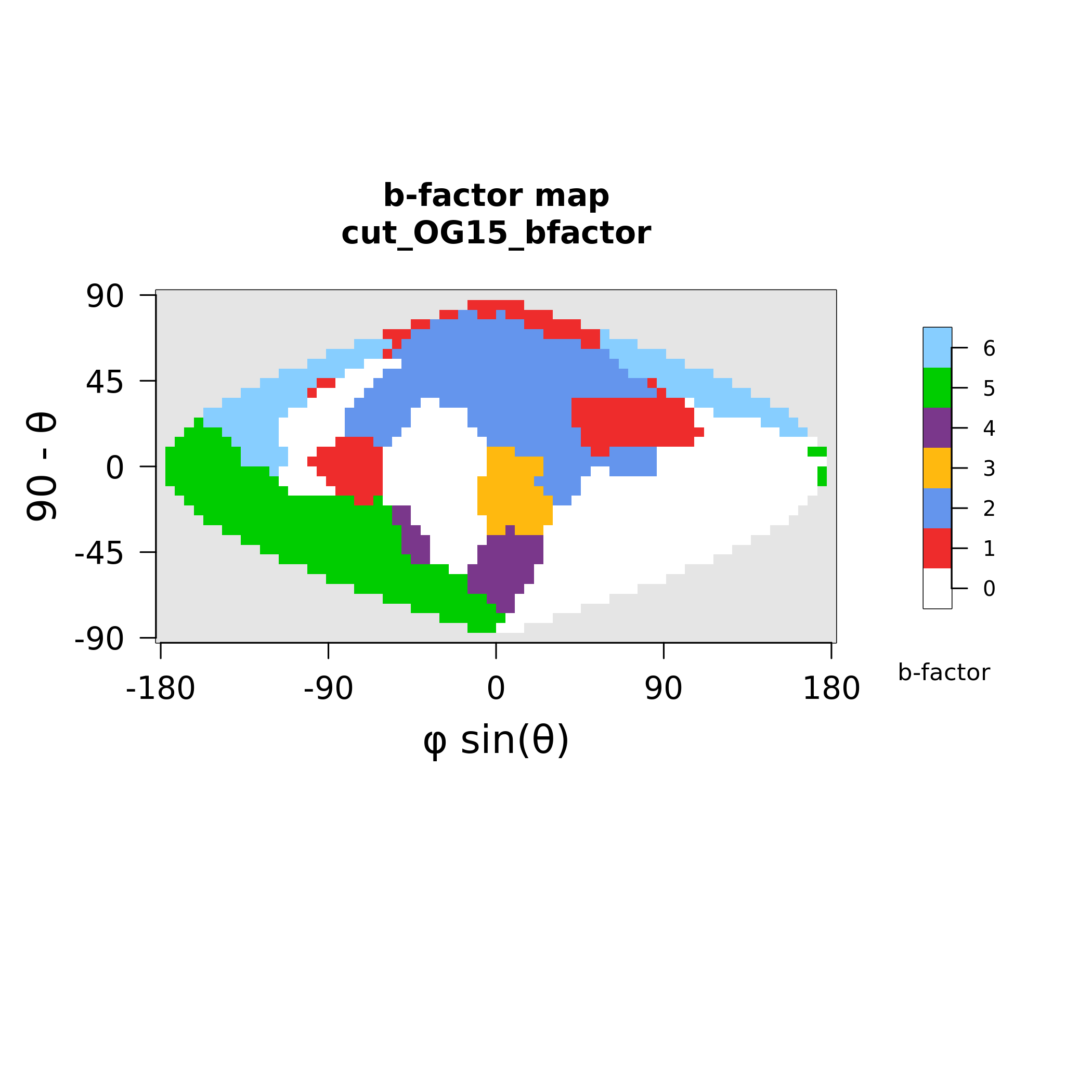

### MAX17_polymorphism.png

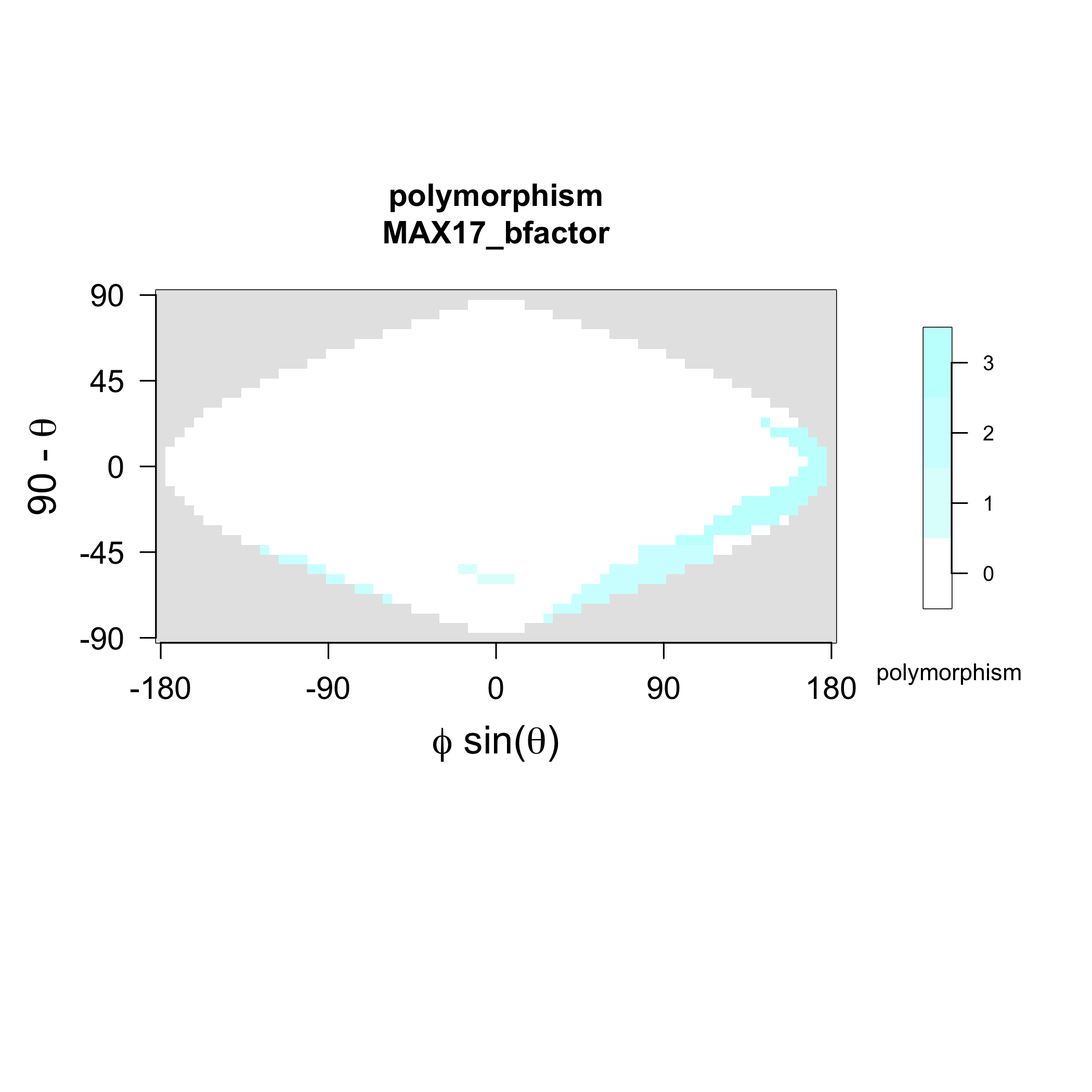

### MAX17_strands.png

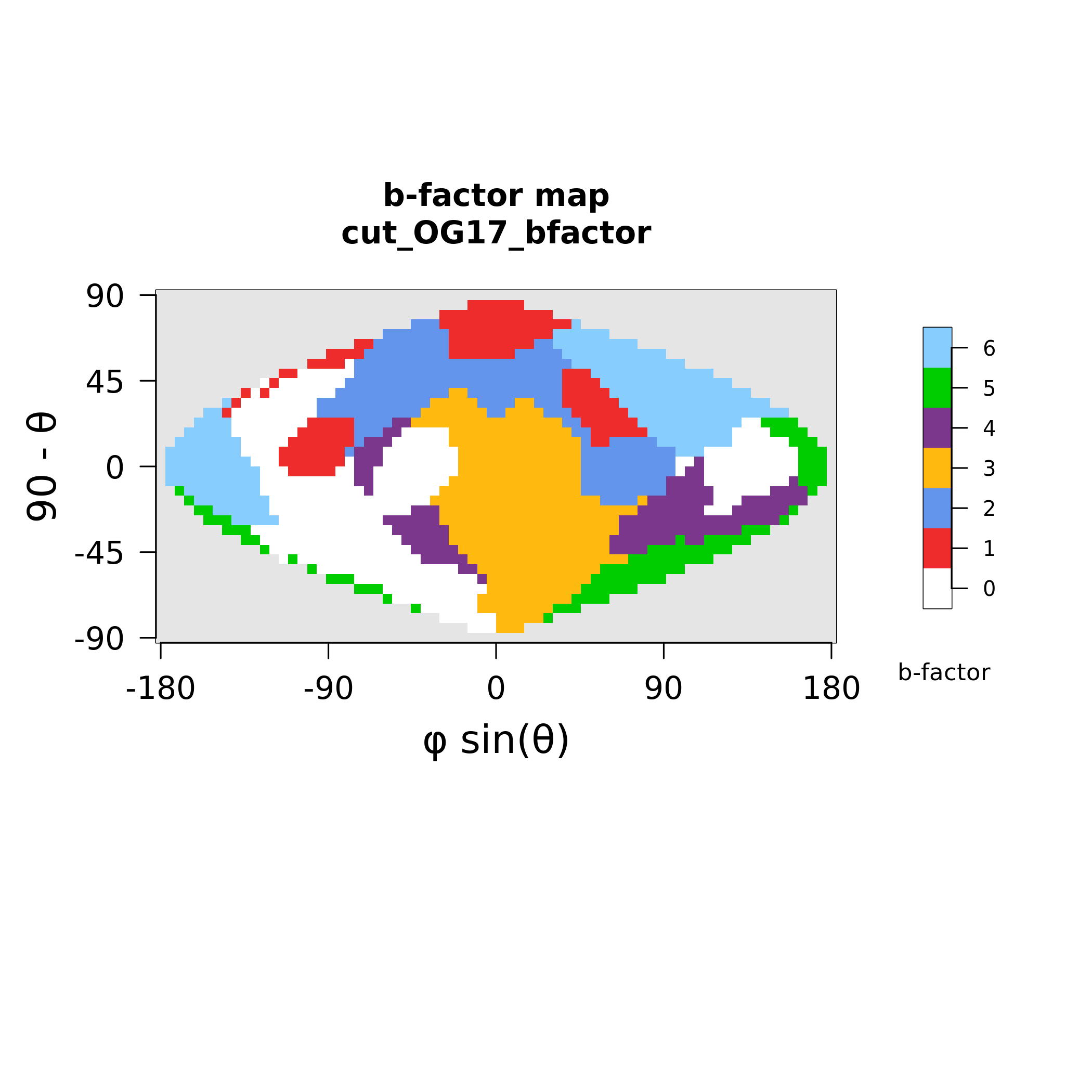

### MAX18_polymorphism.png

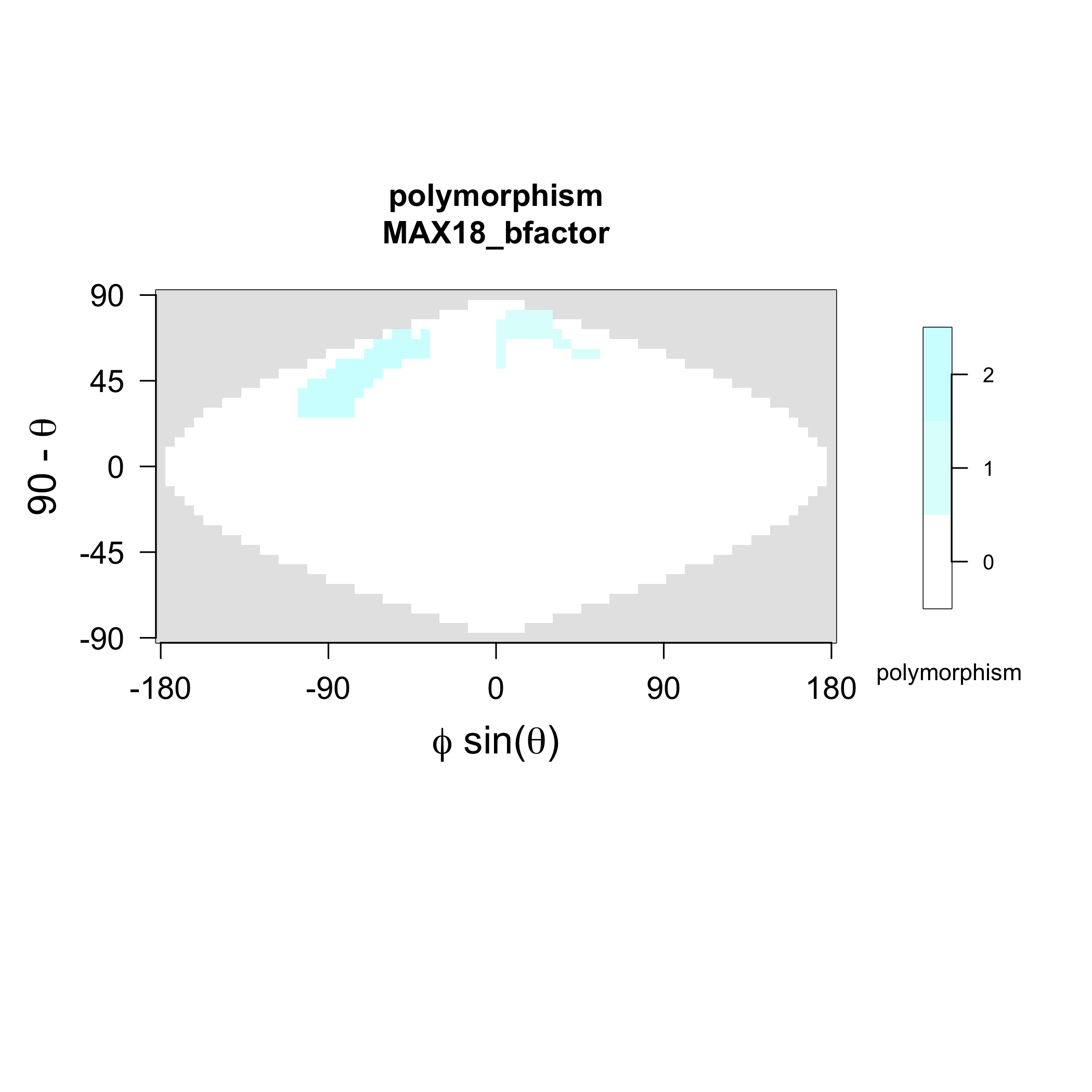

### MAX18_strands.png

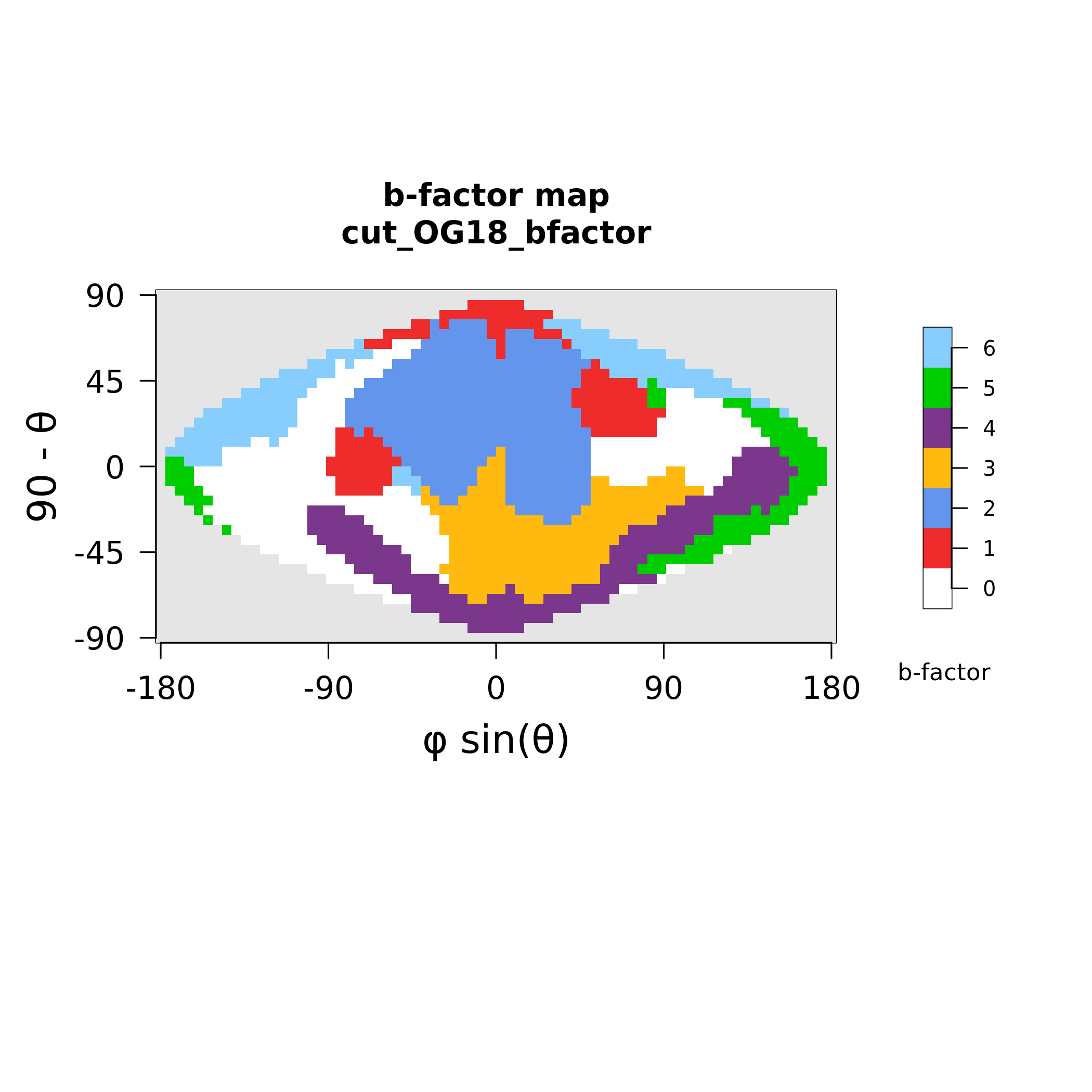

### MAX19_polymorphism.png

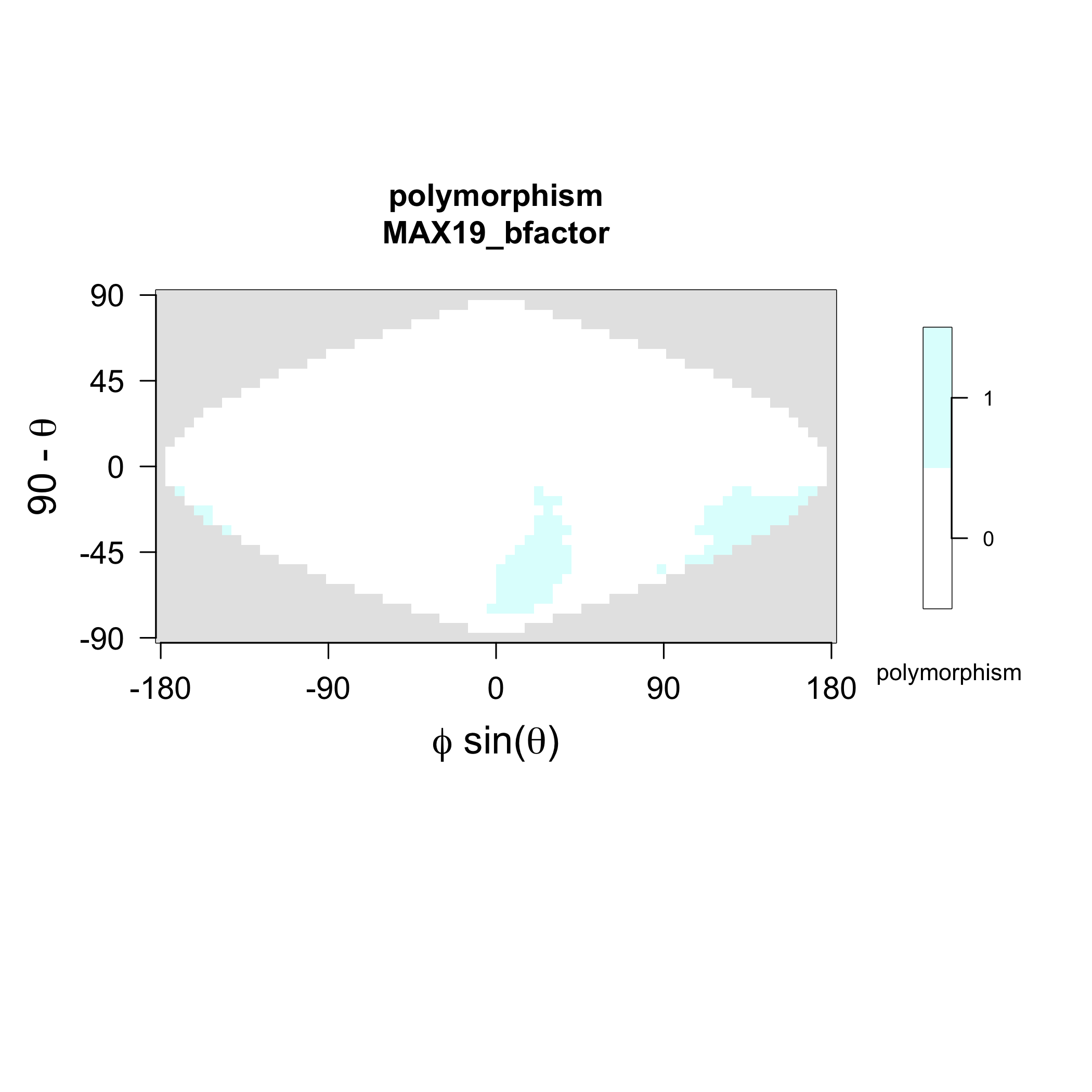

### MAX19_strands.png

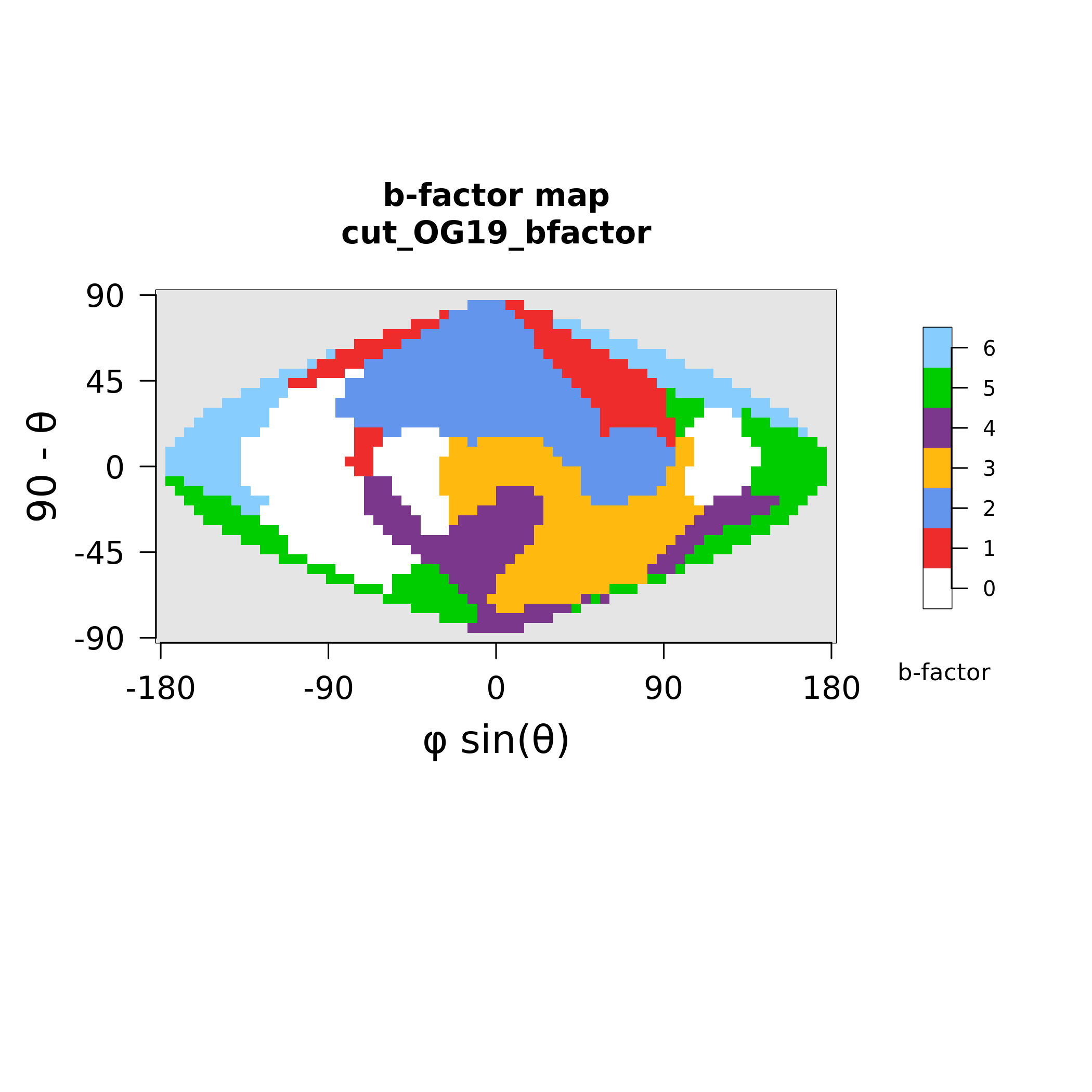

### MAX20_polymorphism.png

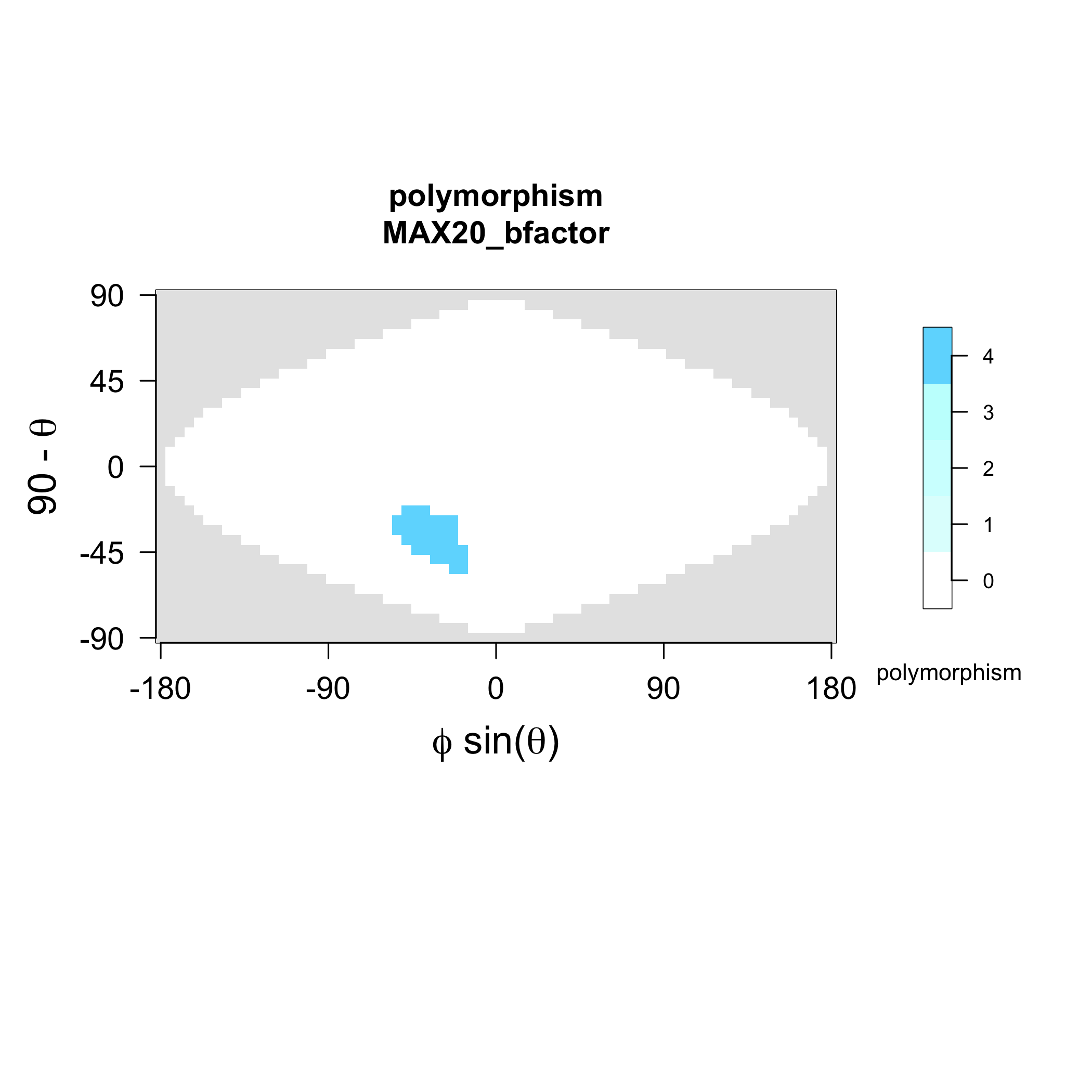

### MAX20_strands.png

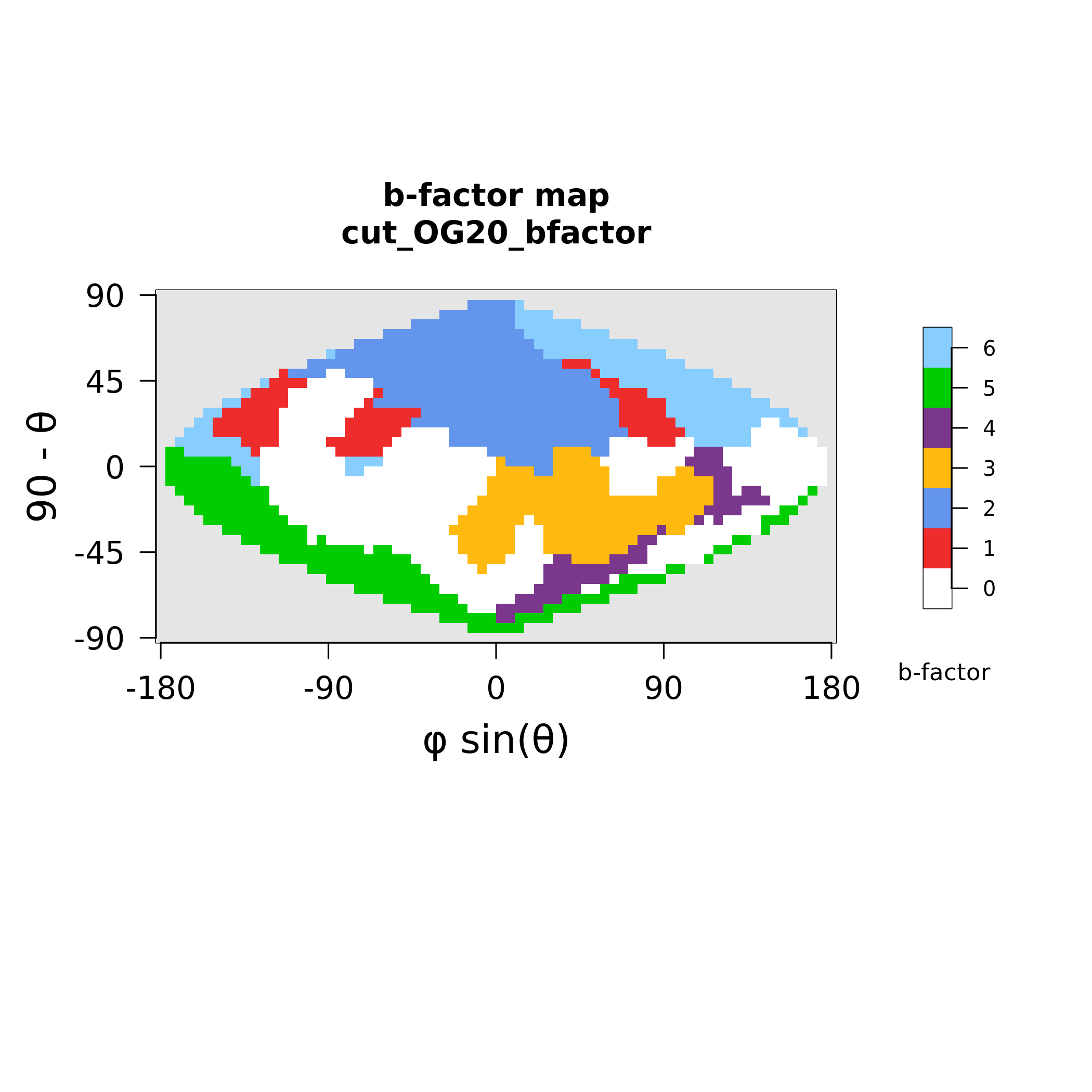

### MAX21_polymorphism.png

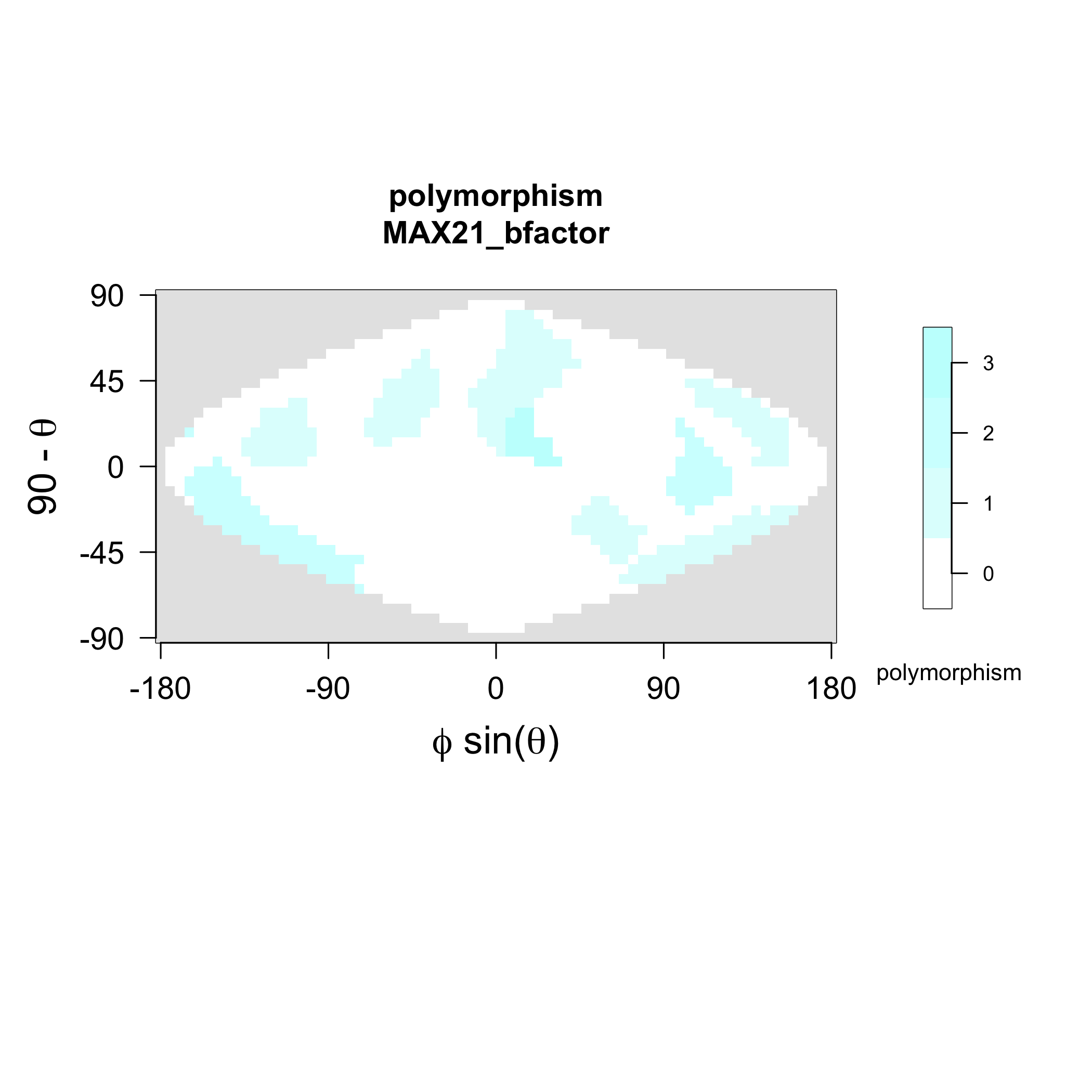

### MAX21_strands.png

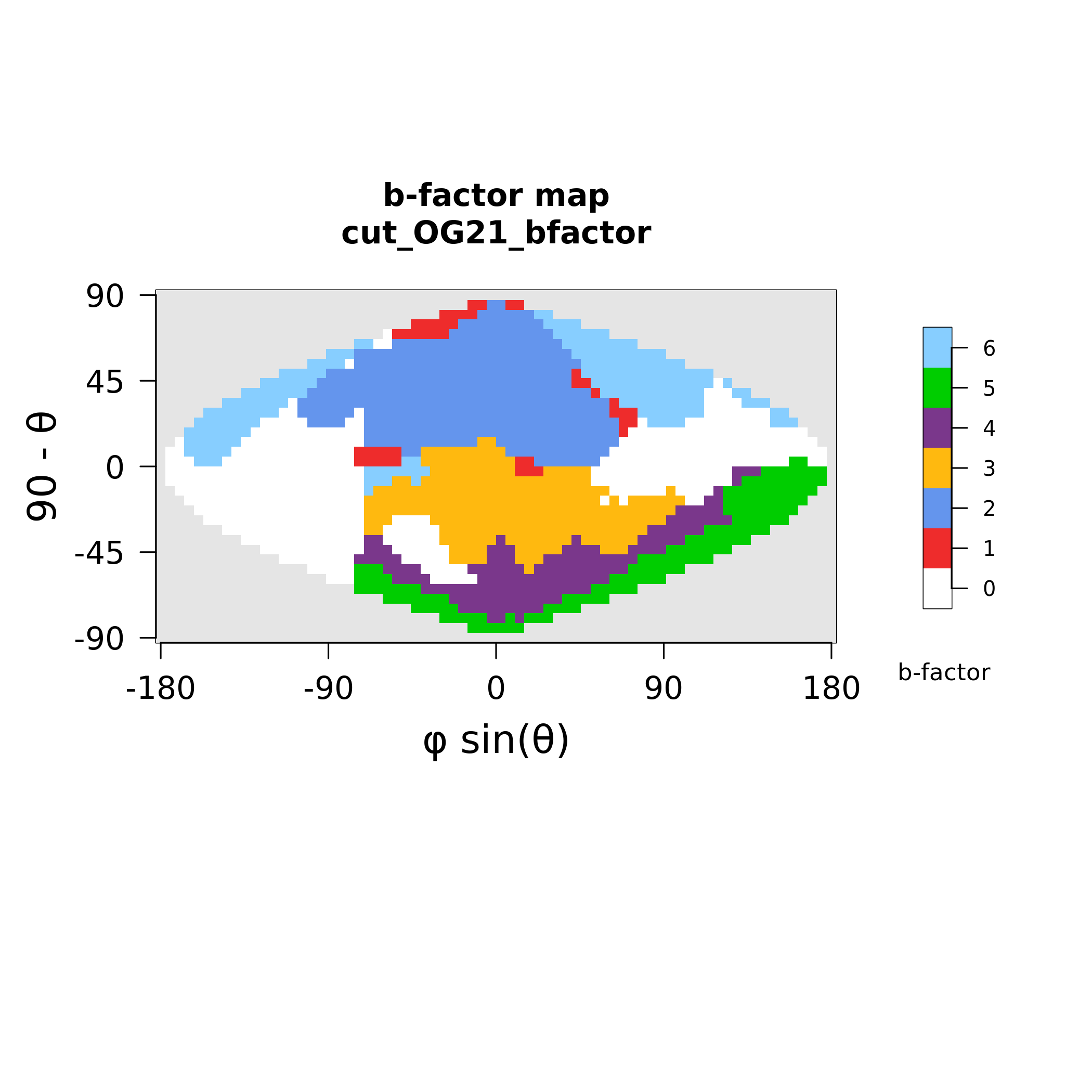

### MAX23_polymorphism.png

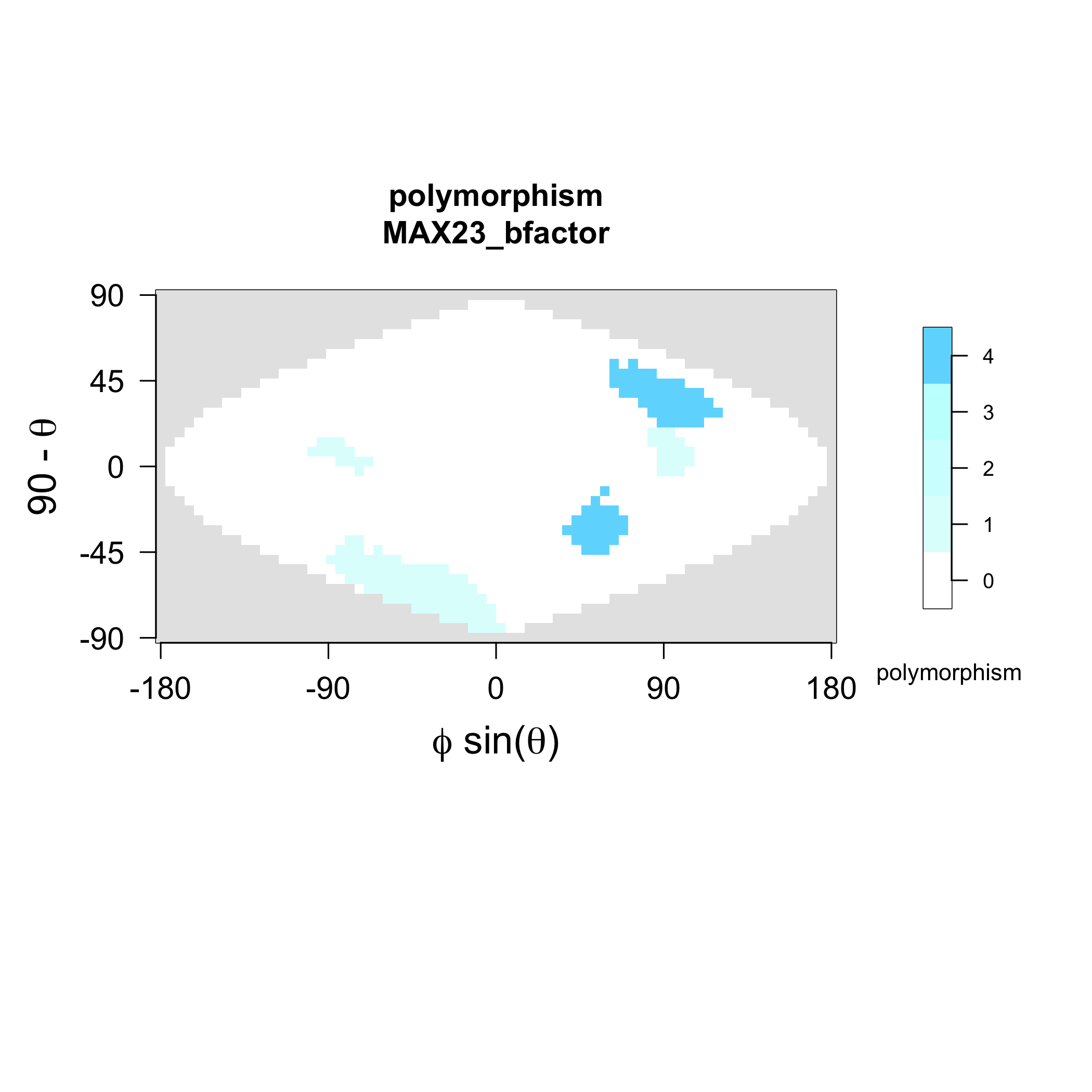

### MAX23_strands.png

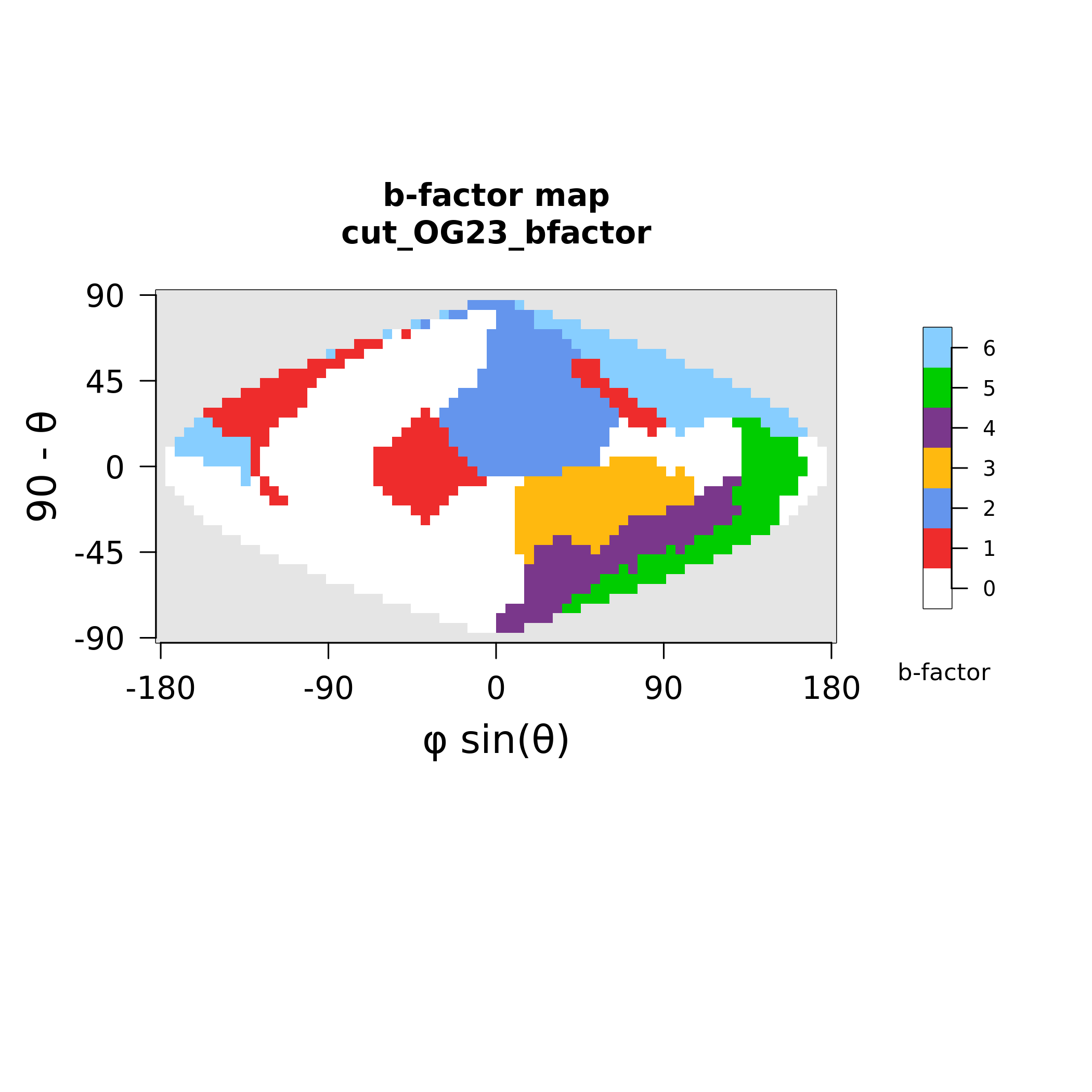

### MAX24_polymorphism.png

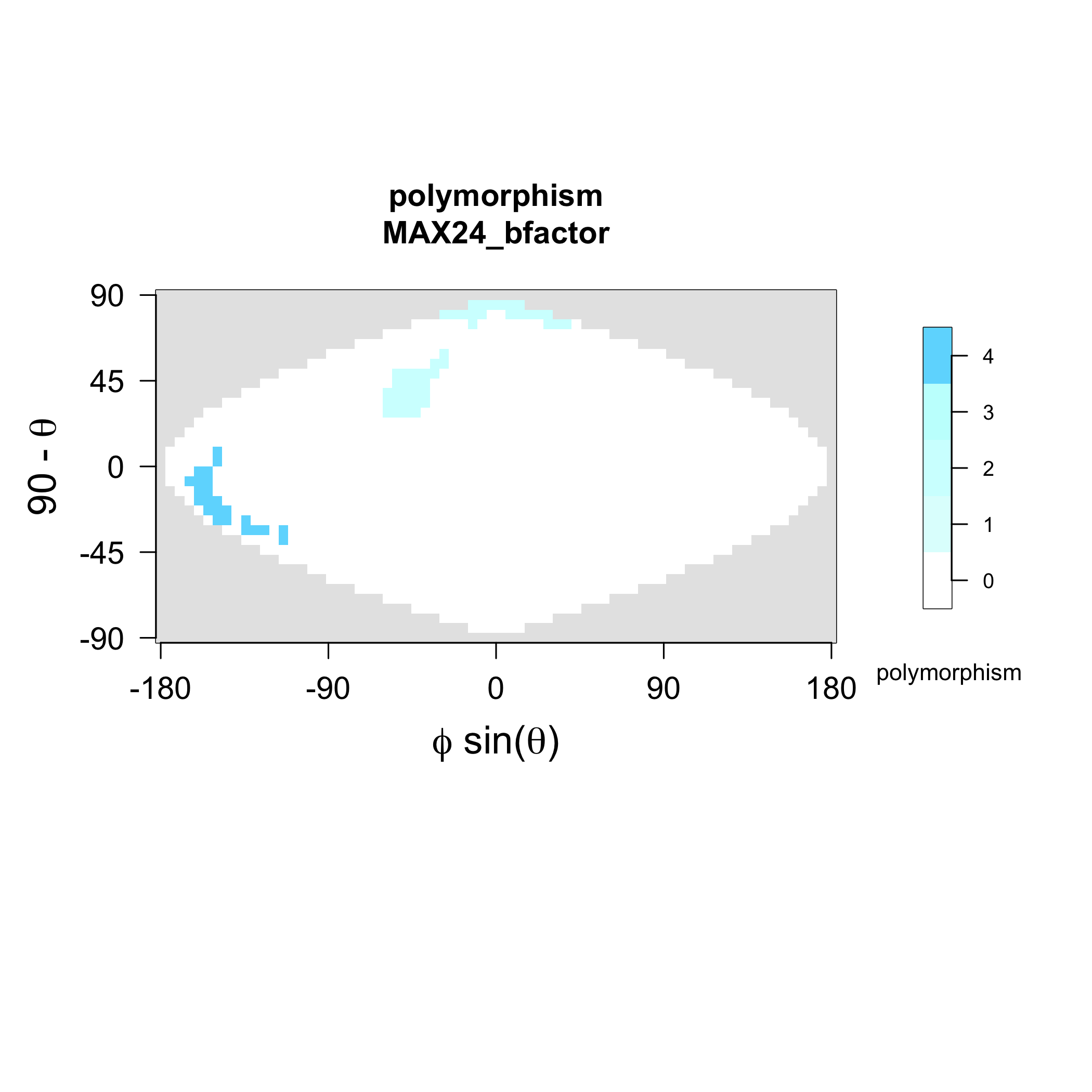

### MAX24_strands.png

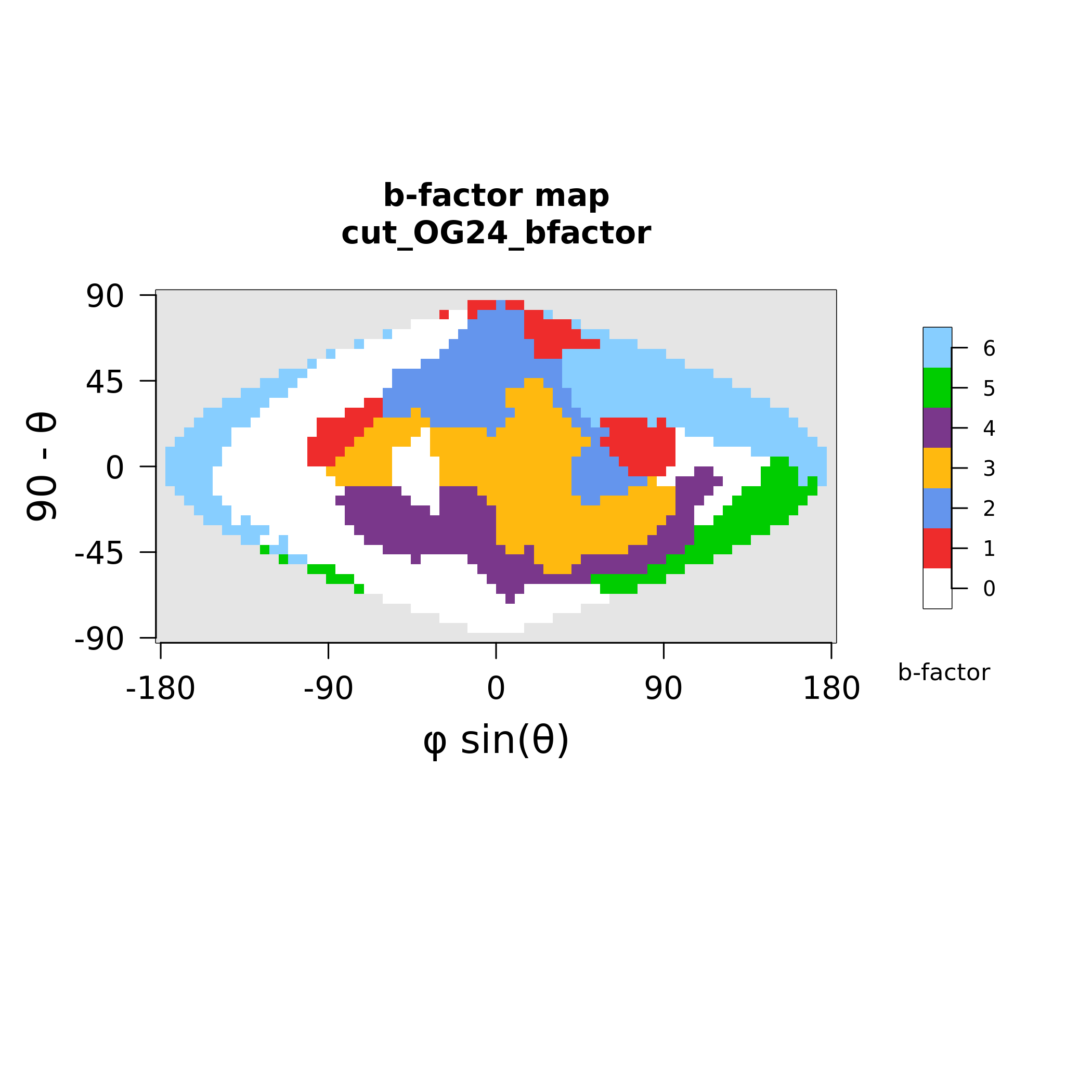

### MAX27_polymorphism.png

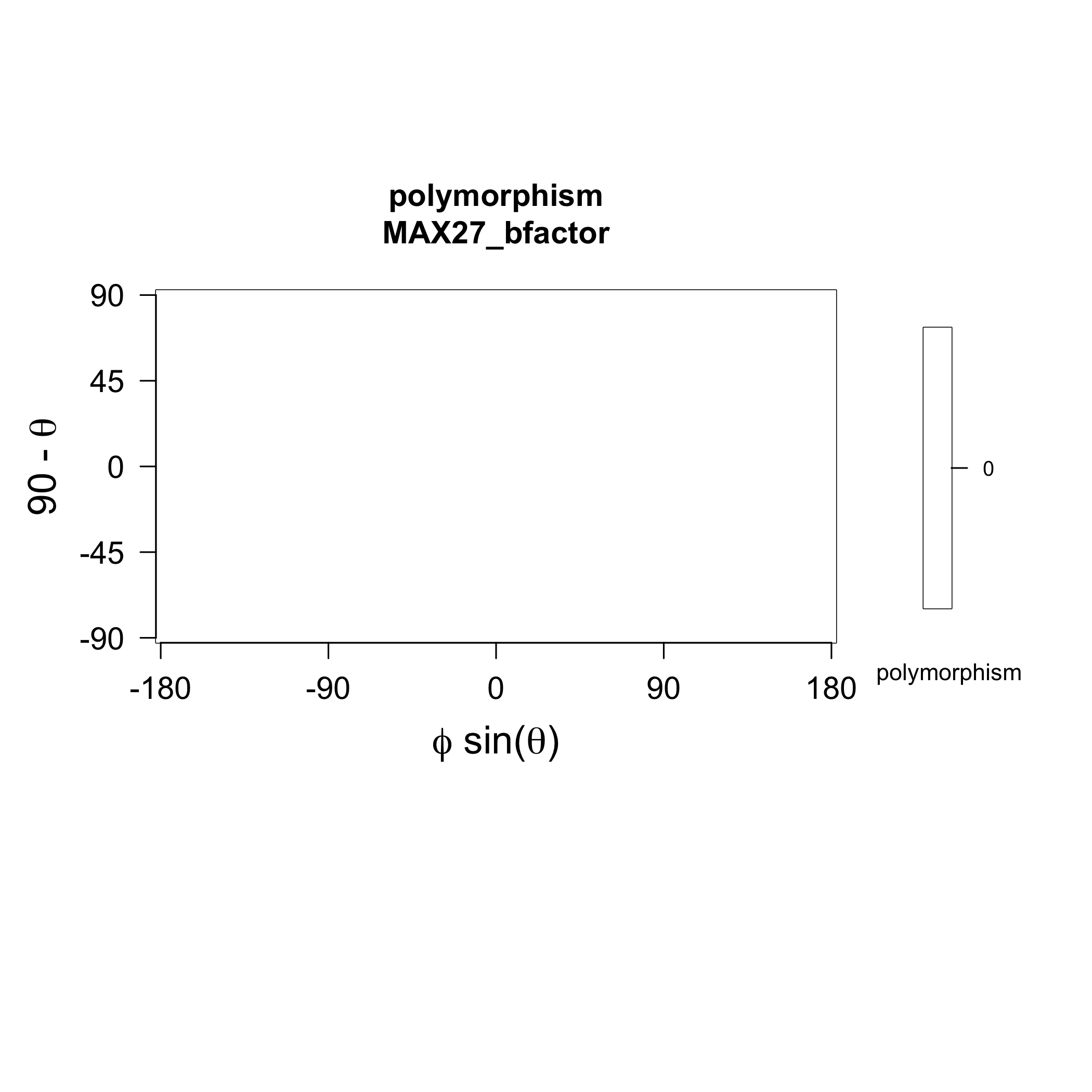

### MAX27_strands.png

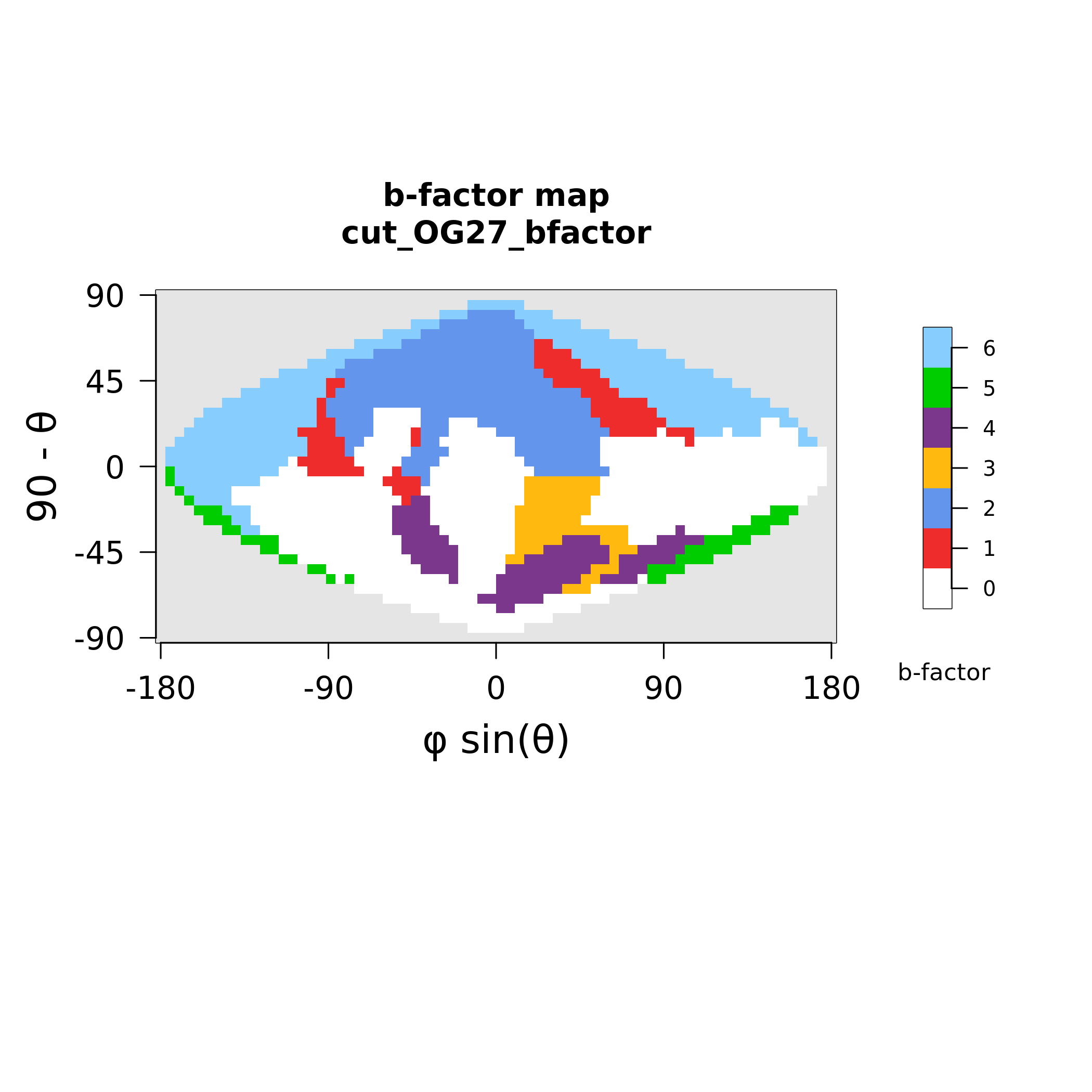

### MAX28_polymorphism.png

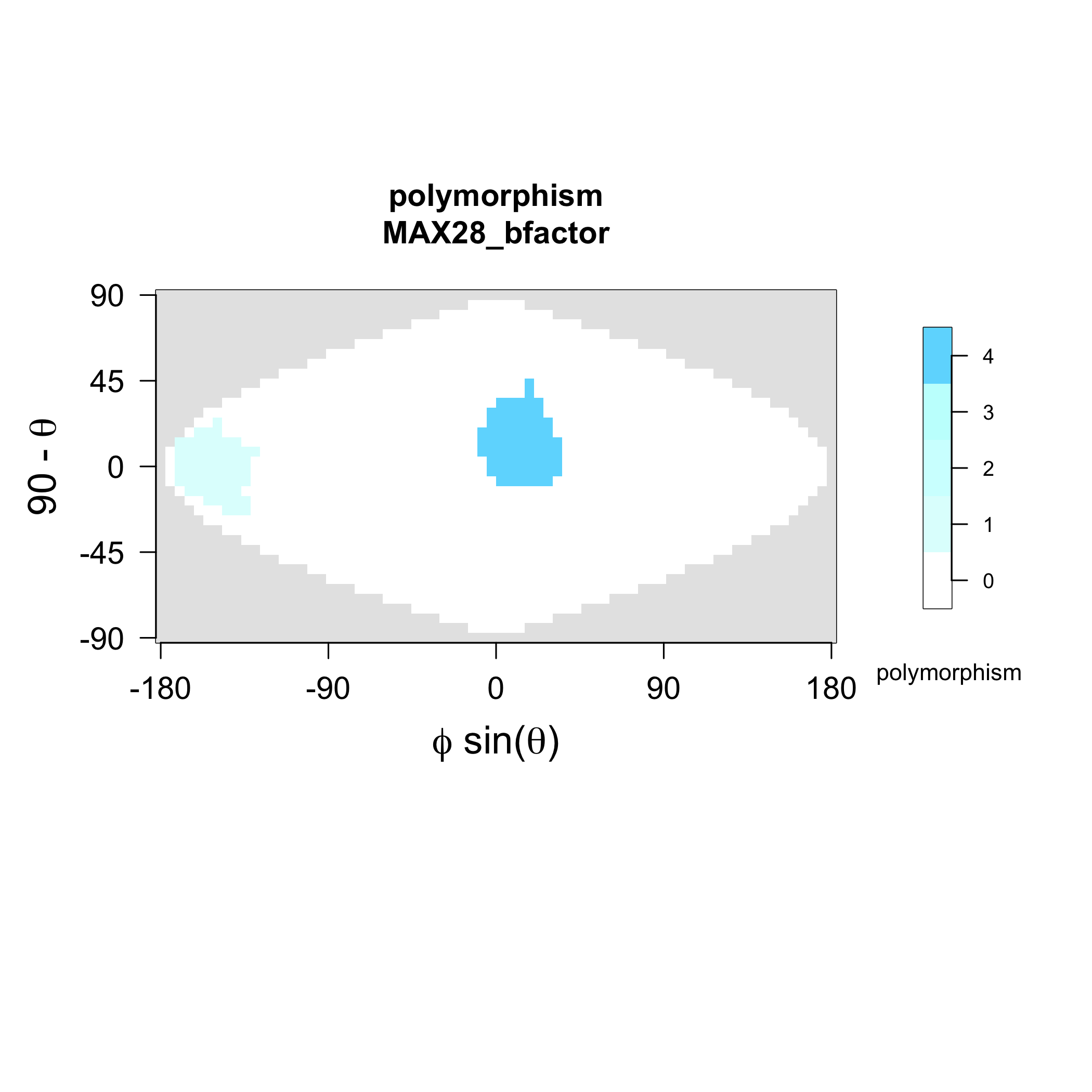

### MAX28_strands.png

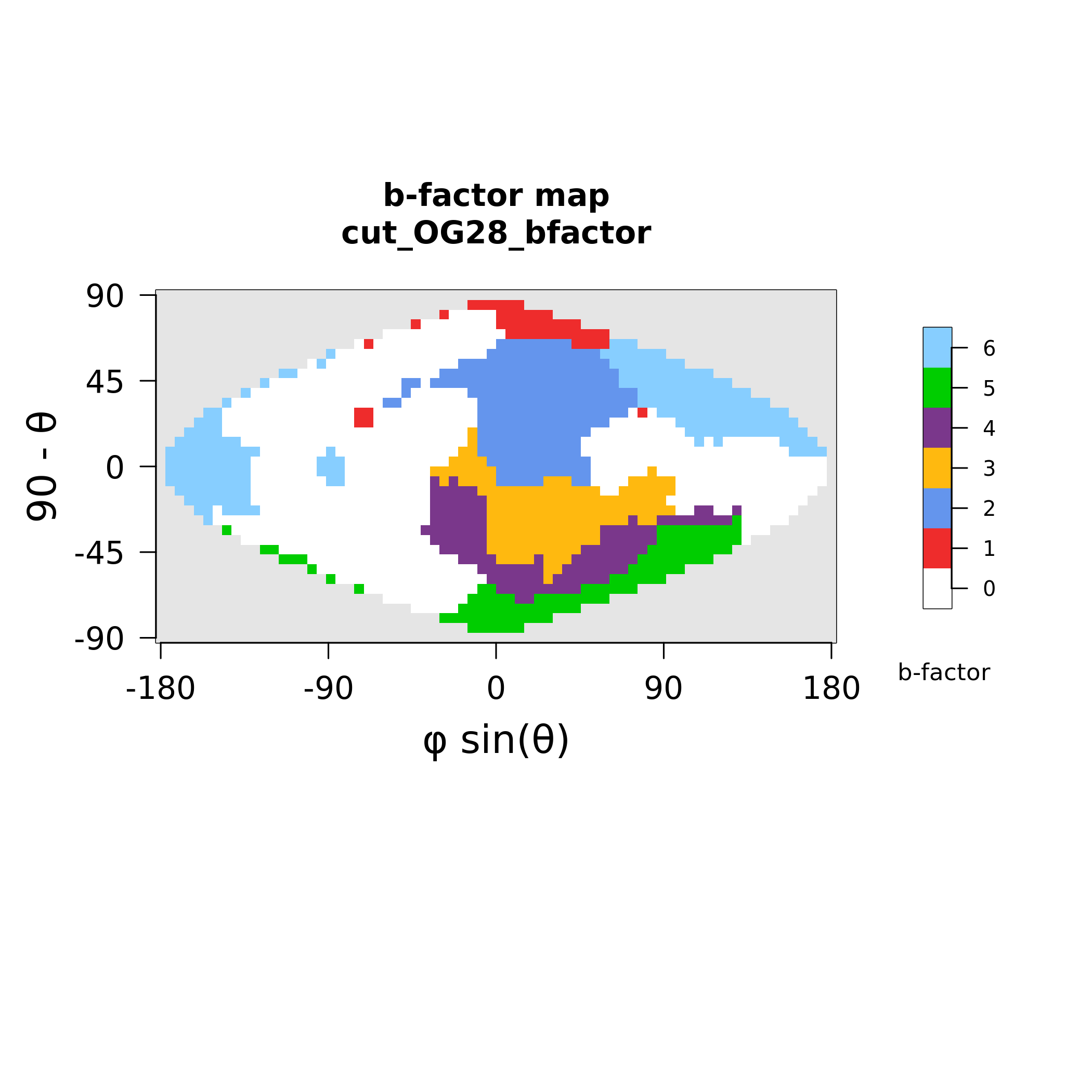

### MAX30_polymorphism.png

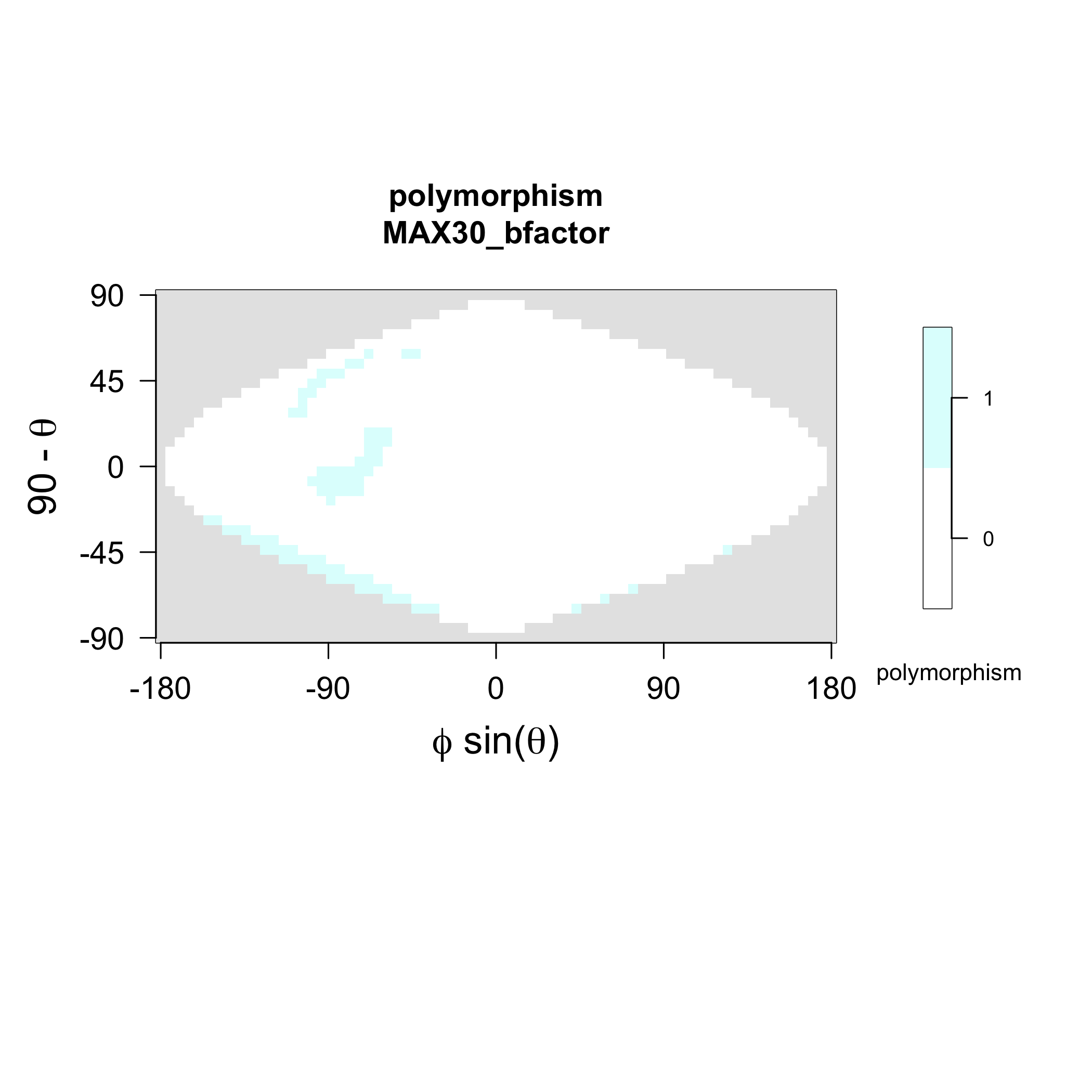

### MAX30_strands.png

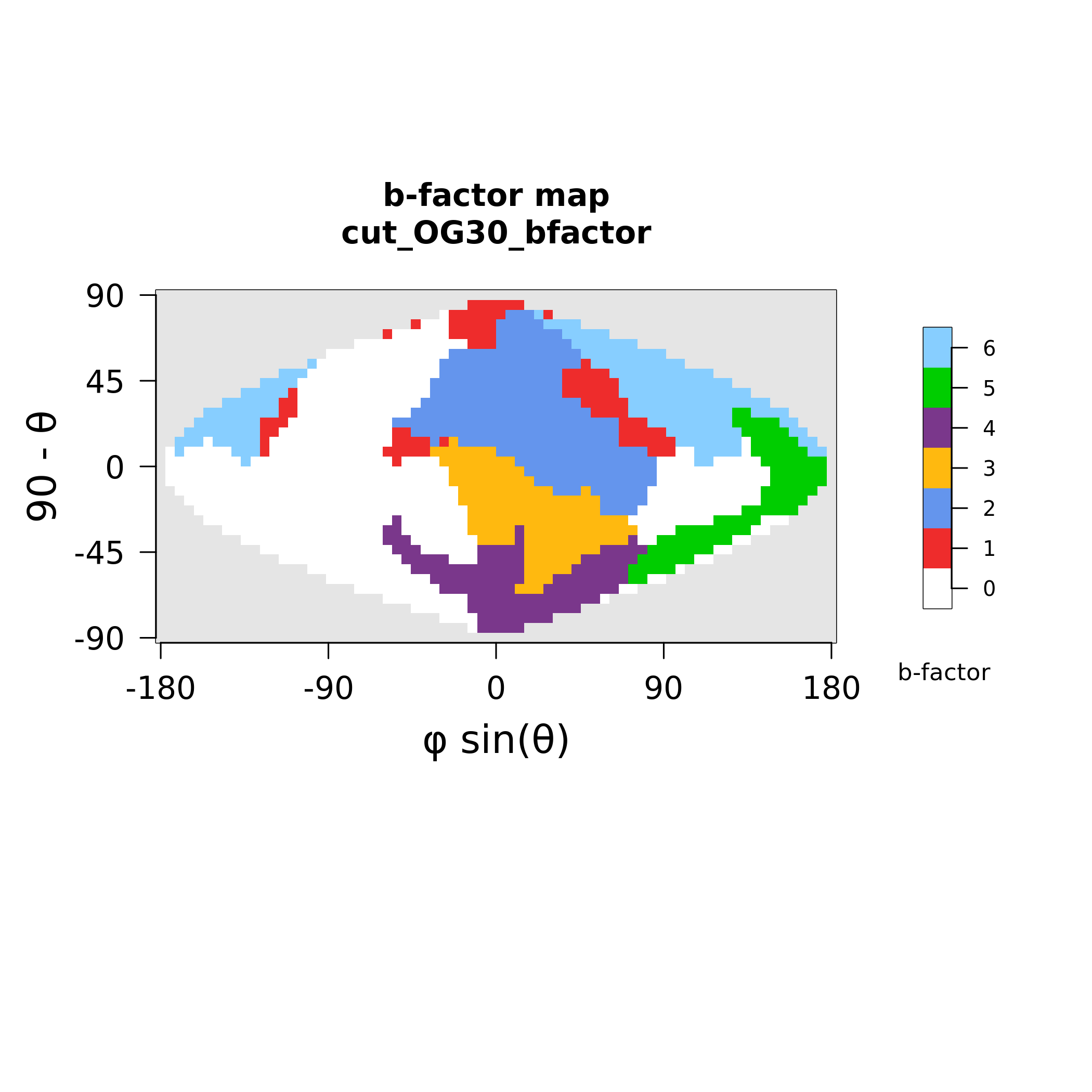

### MAX31_polymorphism.png

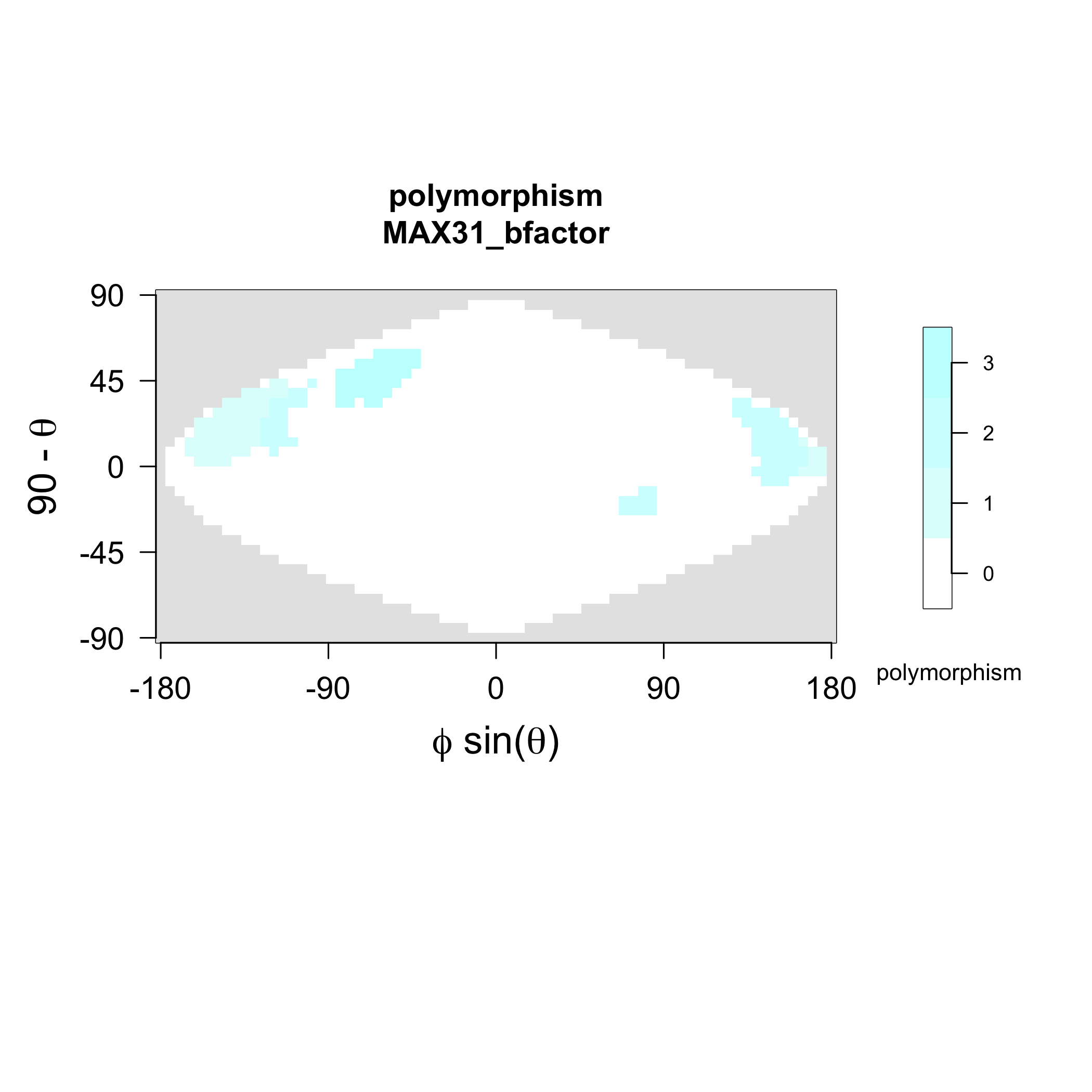

### MAX31_strands.png

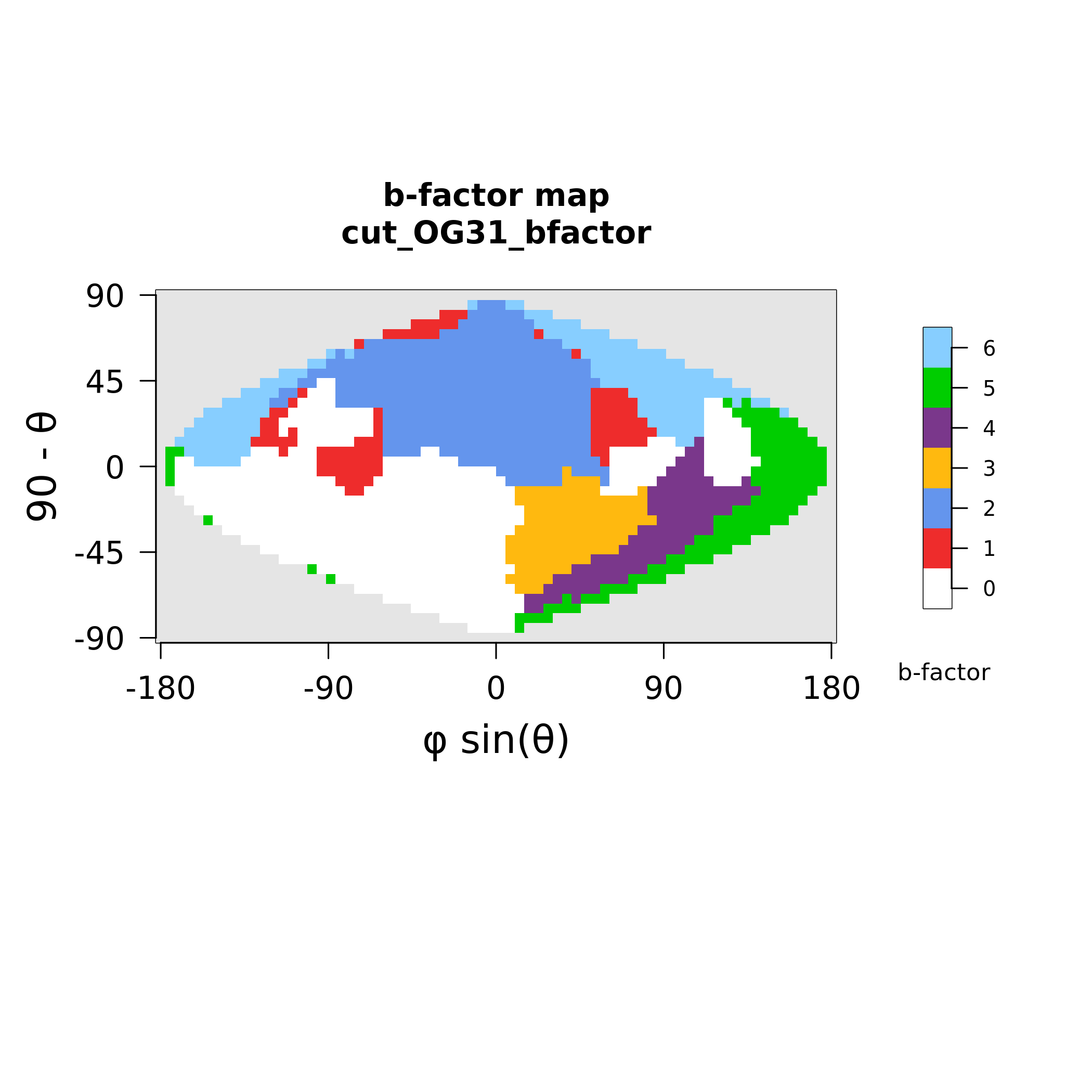

### MAX33_polymorphism.png

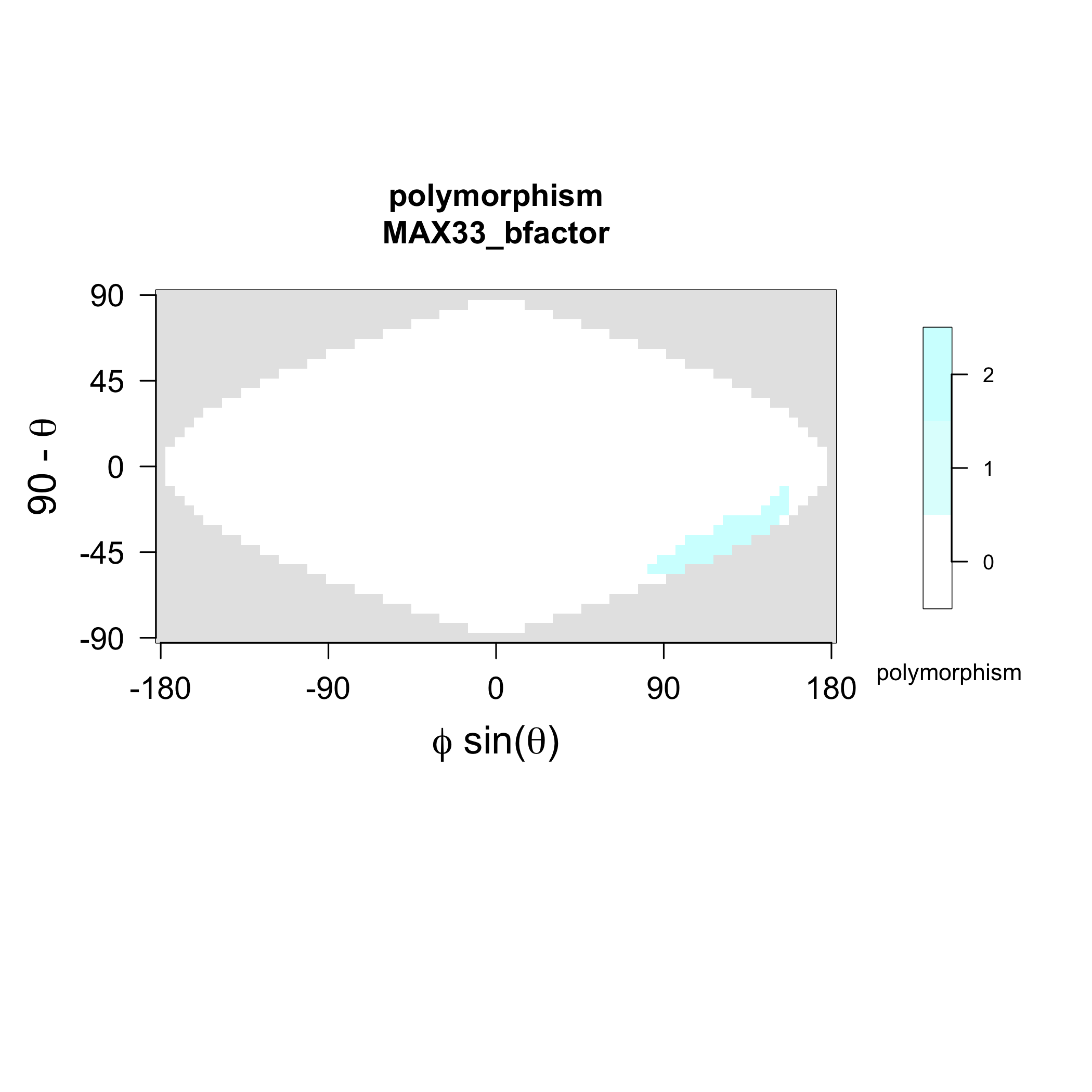

### MAX33_strands.png

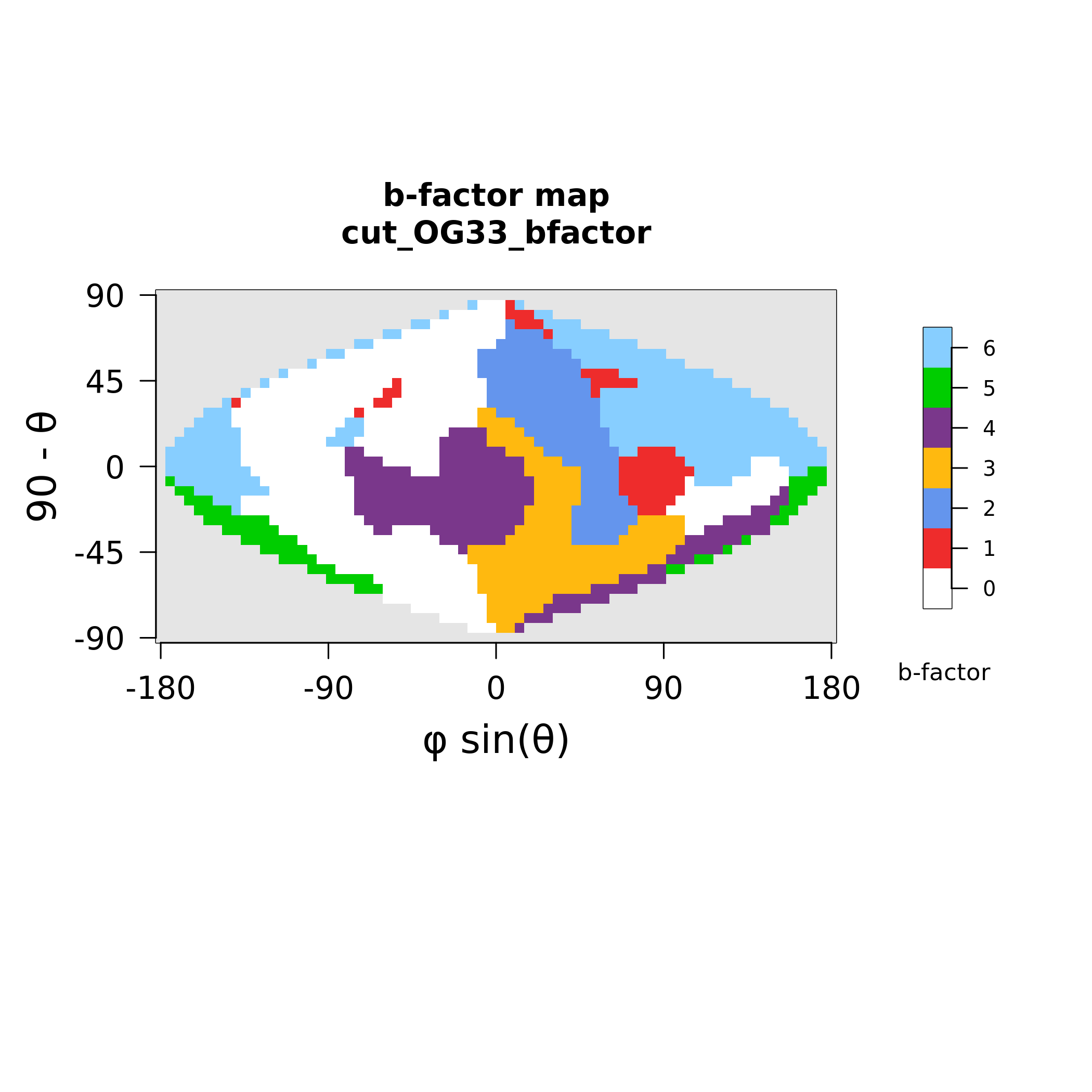

### MAX34_polymorphism.png

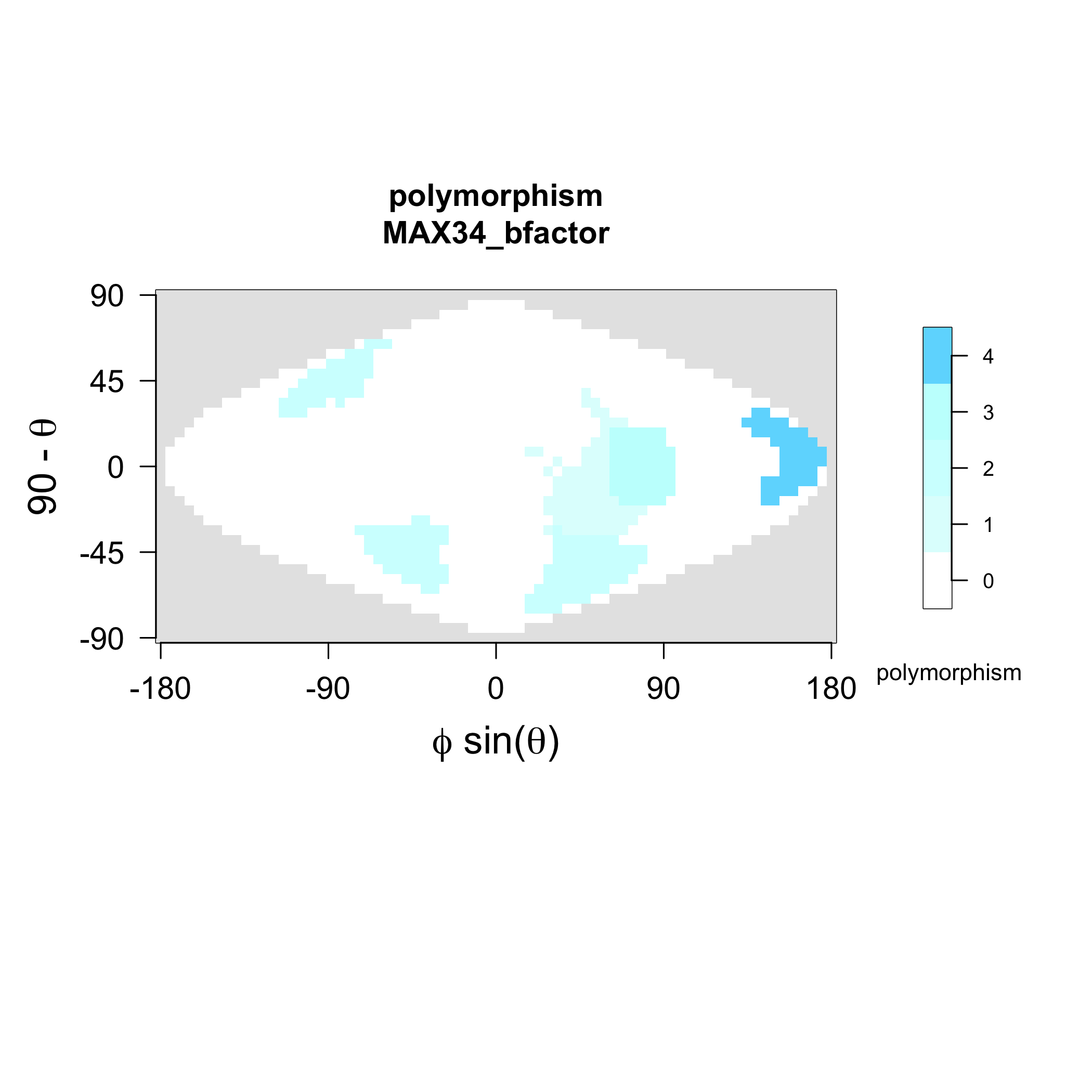

### MAX34_strands.png

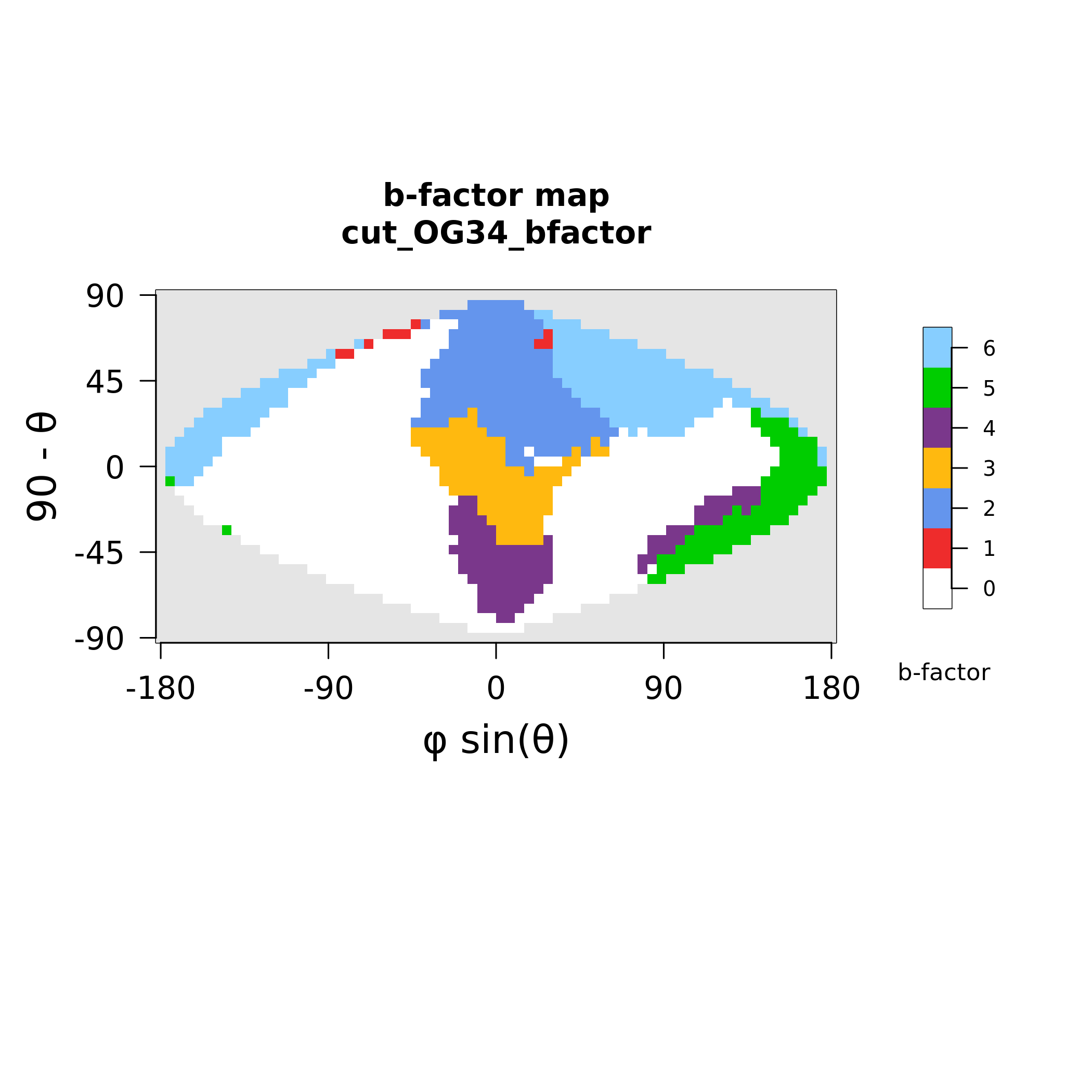

### MAX35_polymorphism.png

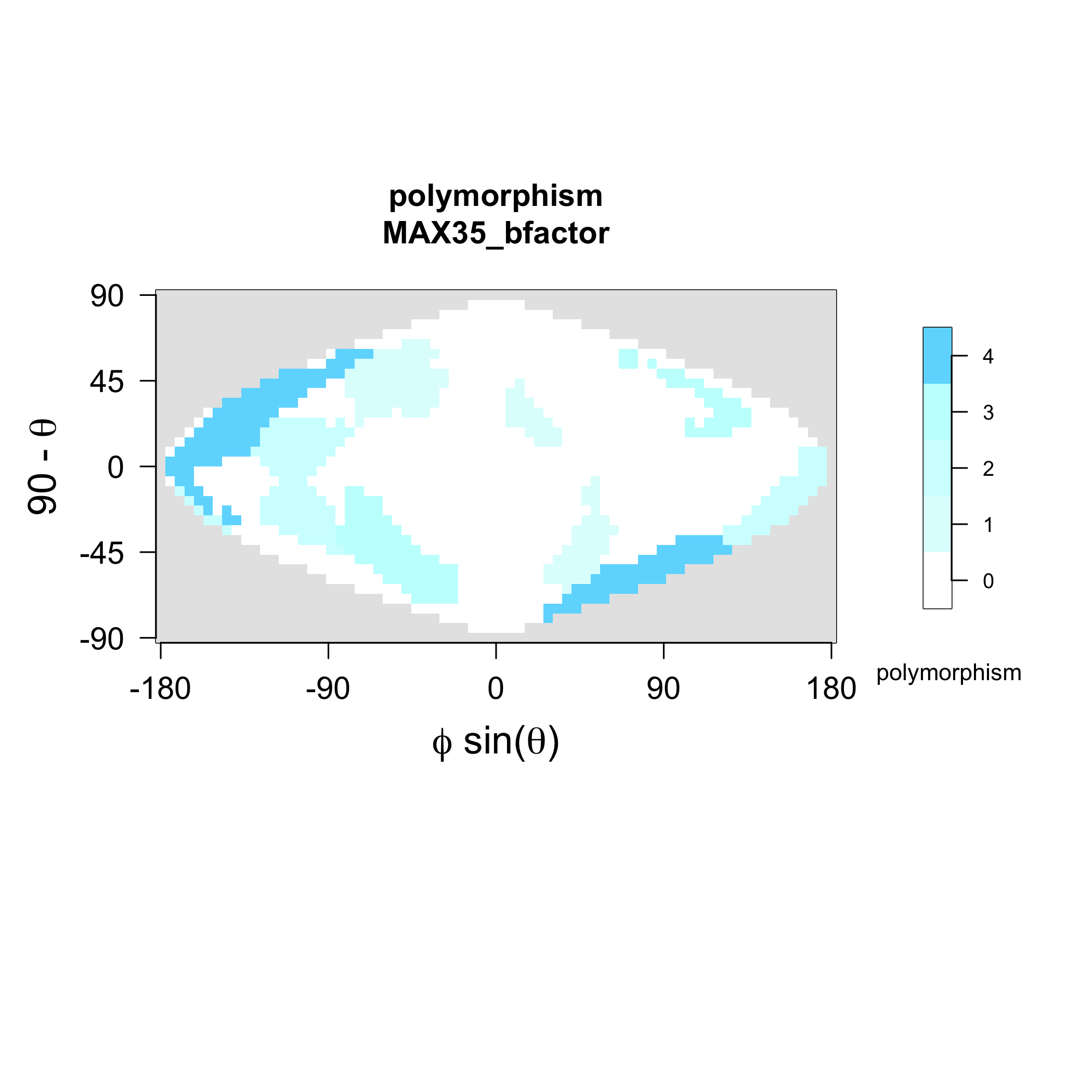

### MAX35_strands.png

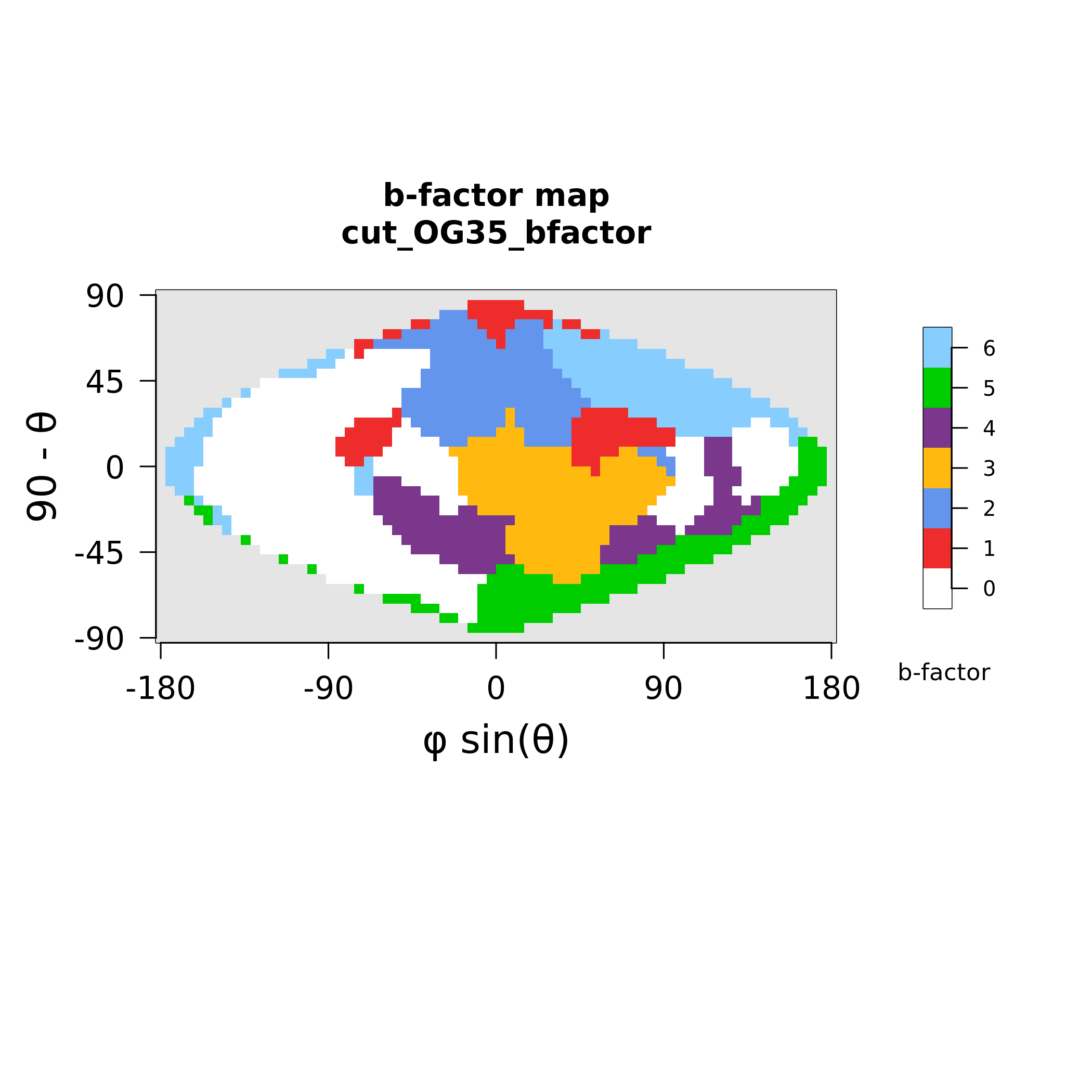

### MAX37_polymorphism.png

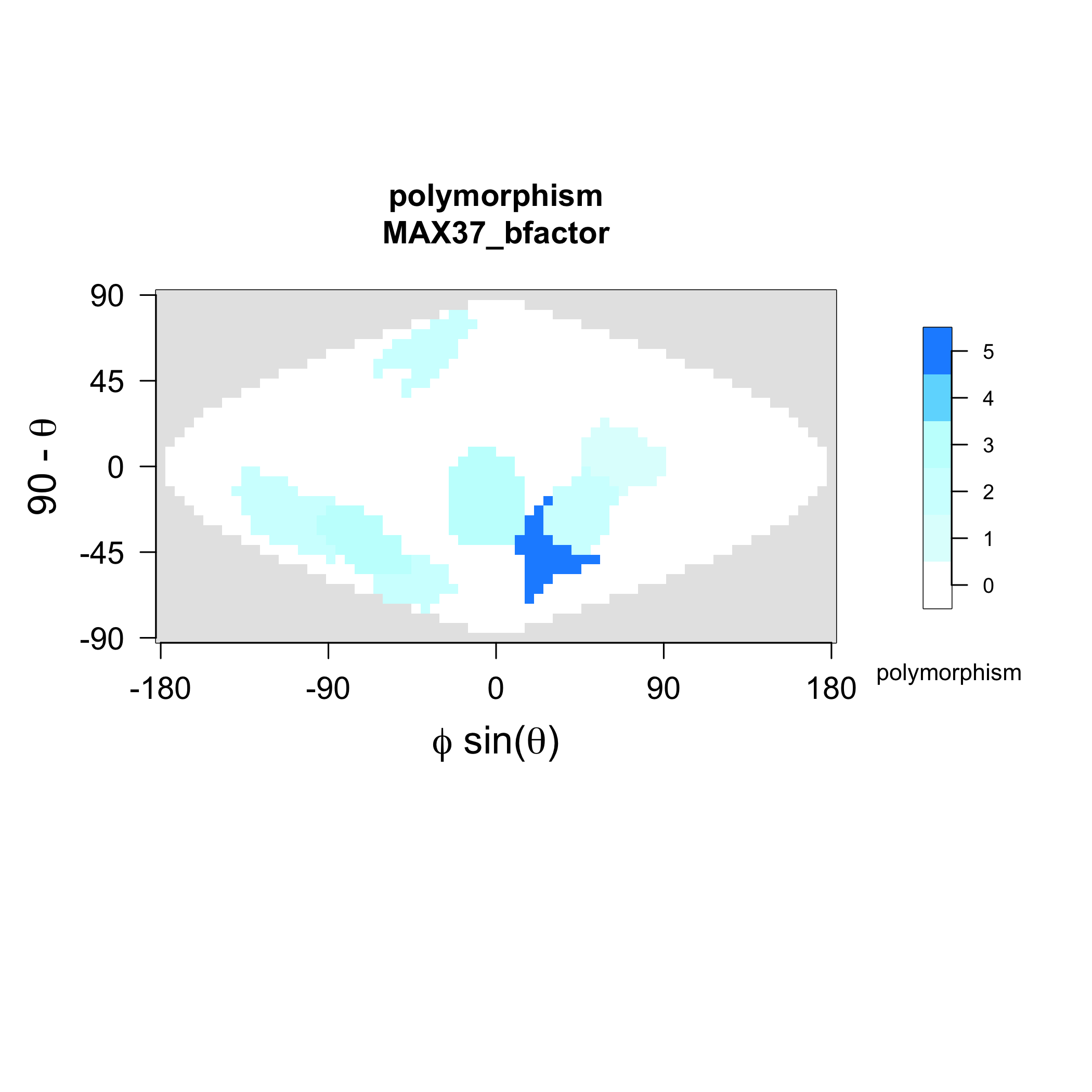

### MAX37_strands.png

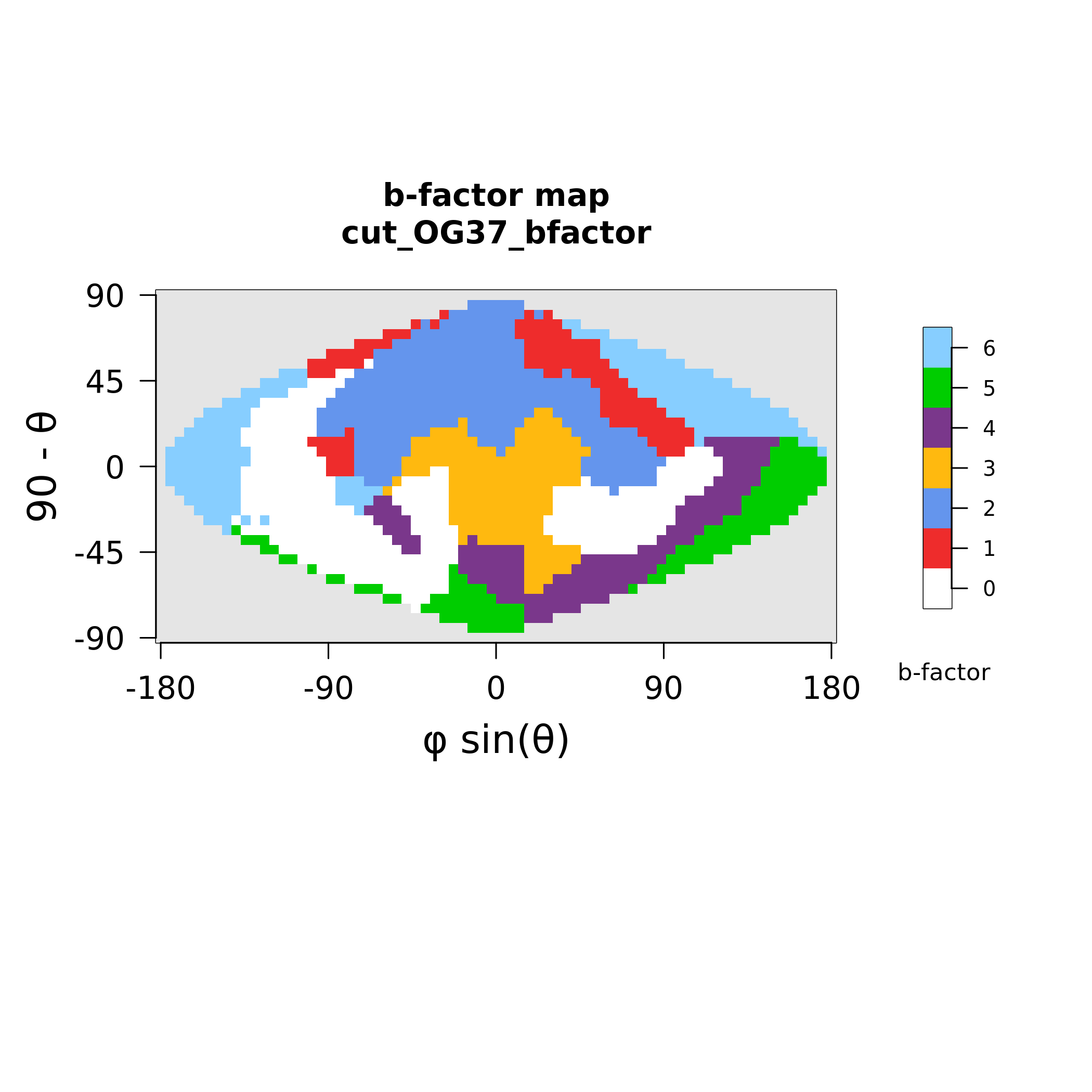

### MAX38_polymorphism.png

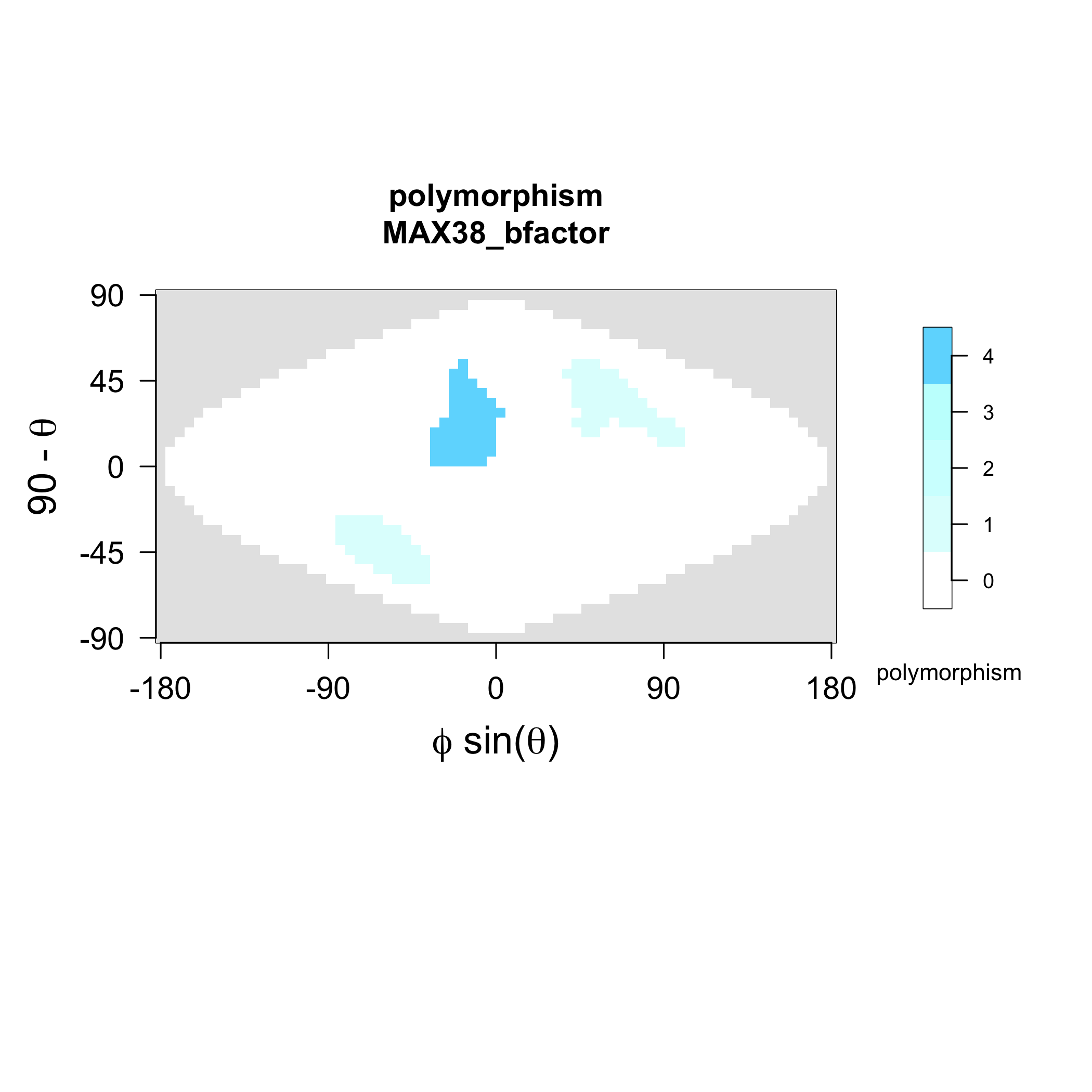

### MAX38_strands.png

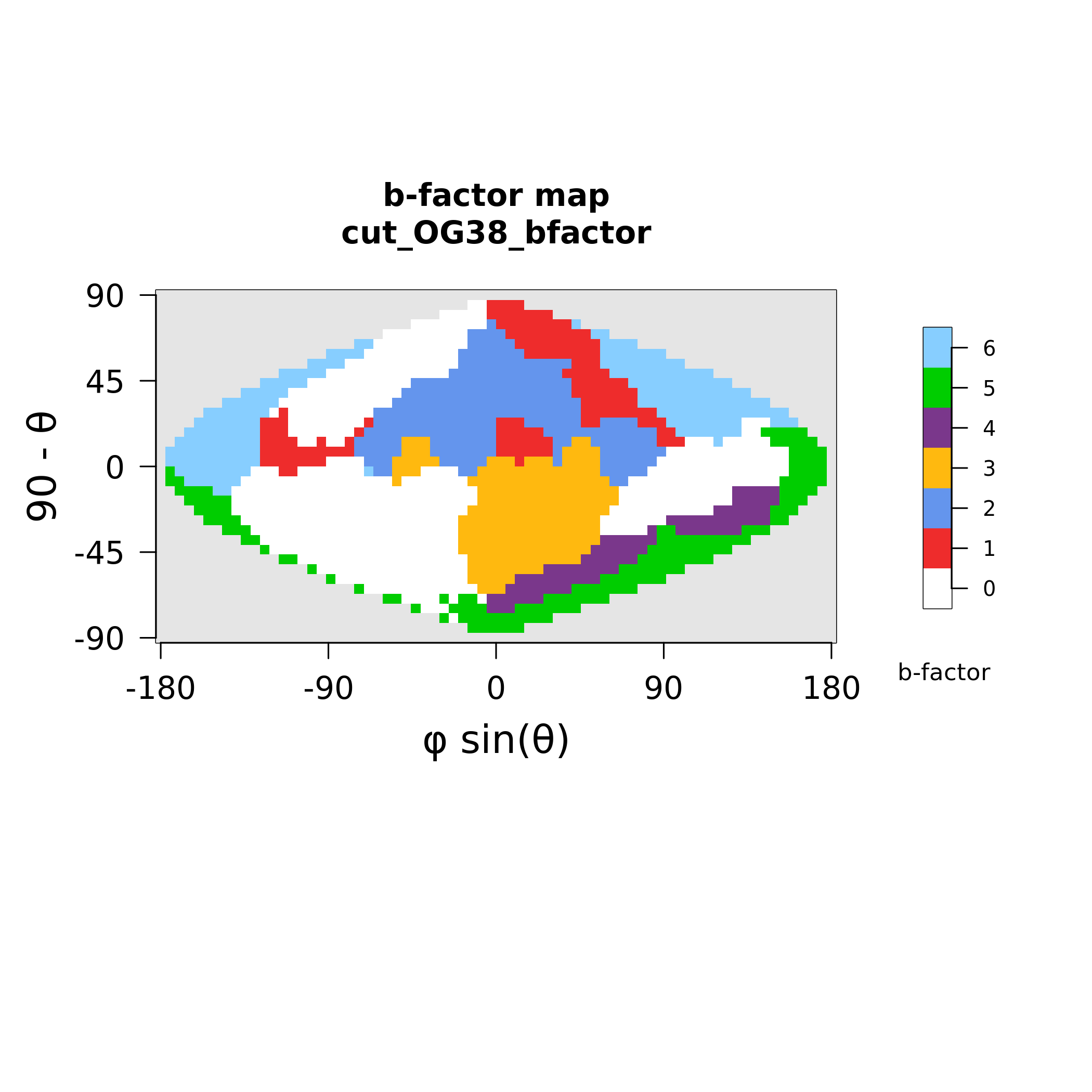

### MAX45_polymorphism.png

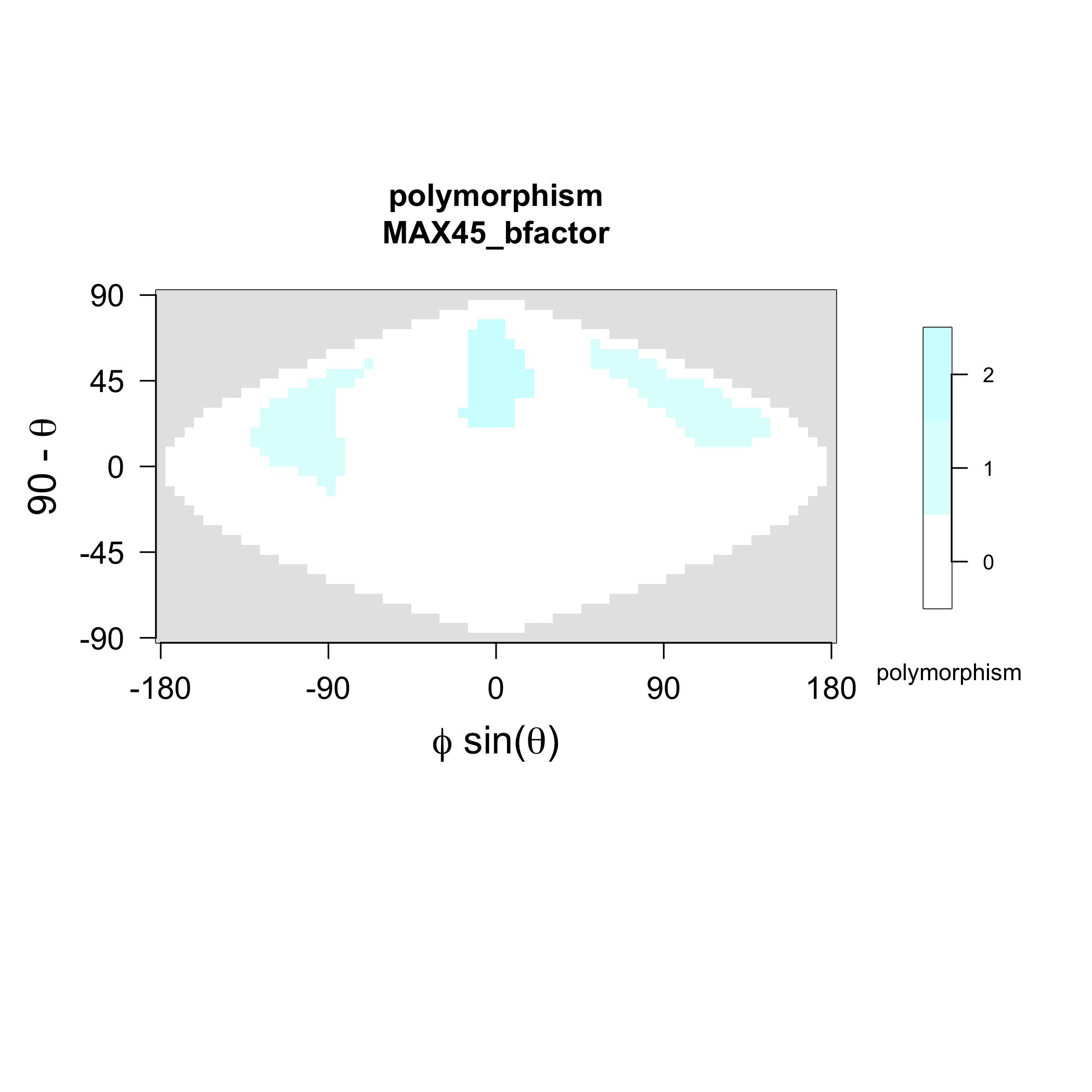

### MAX45_strands.png

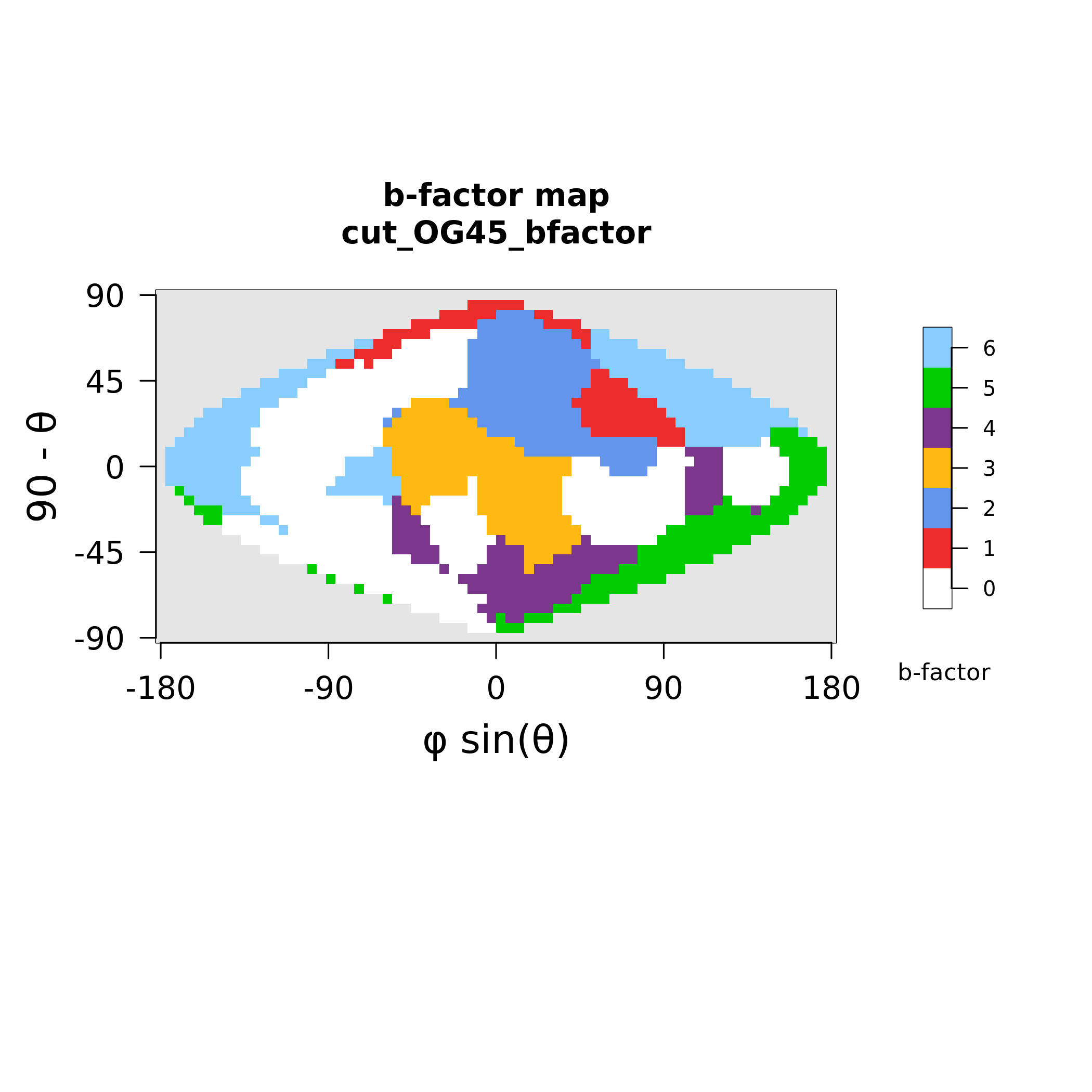

### MAX46_polymorphism.png

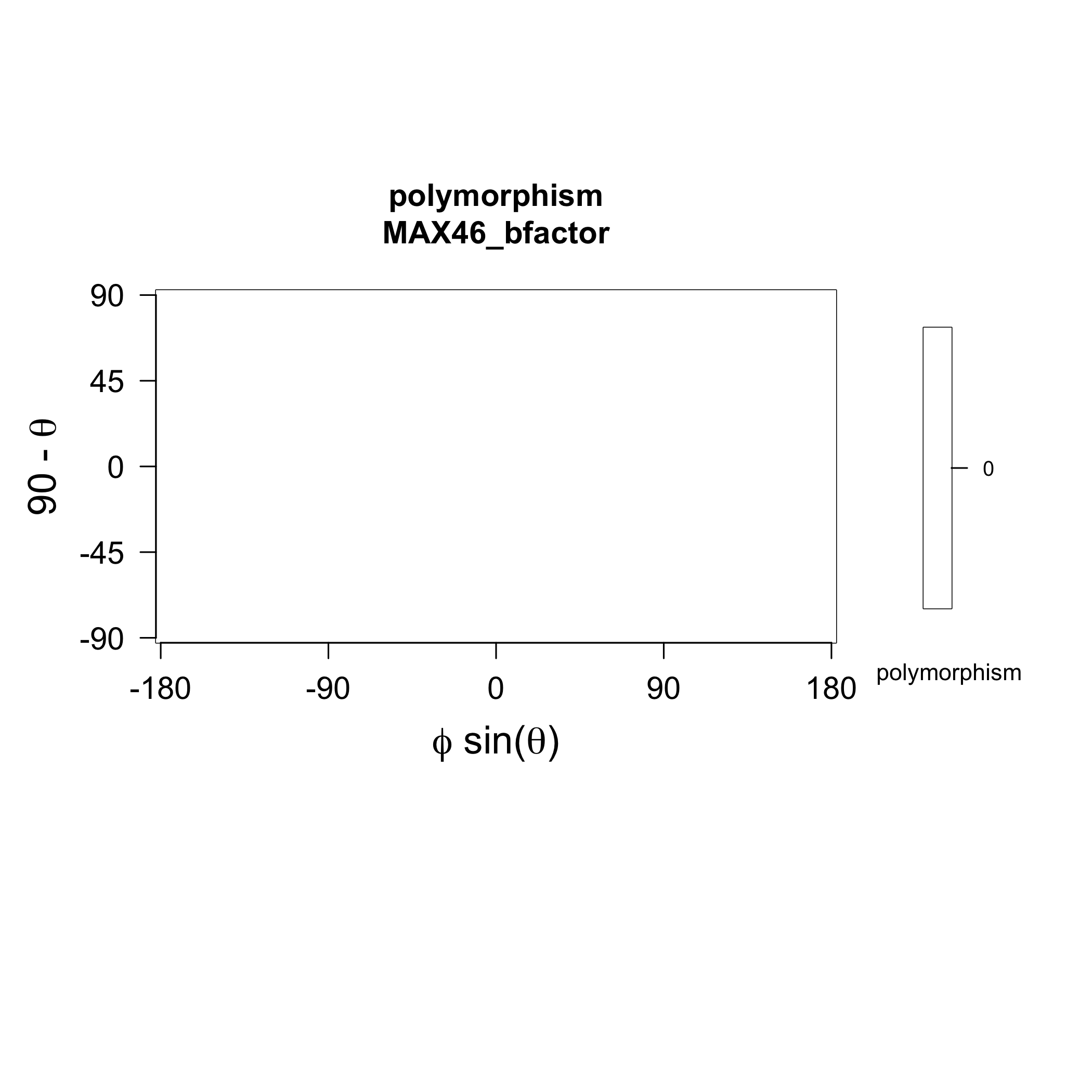

### MAX46_strands.png

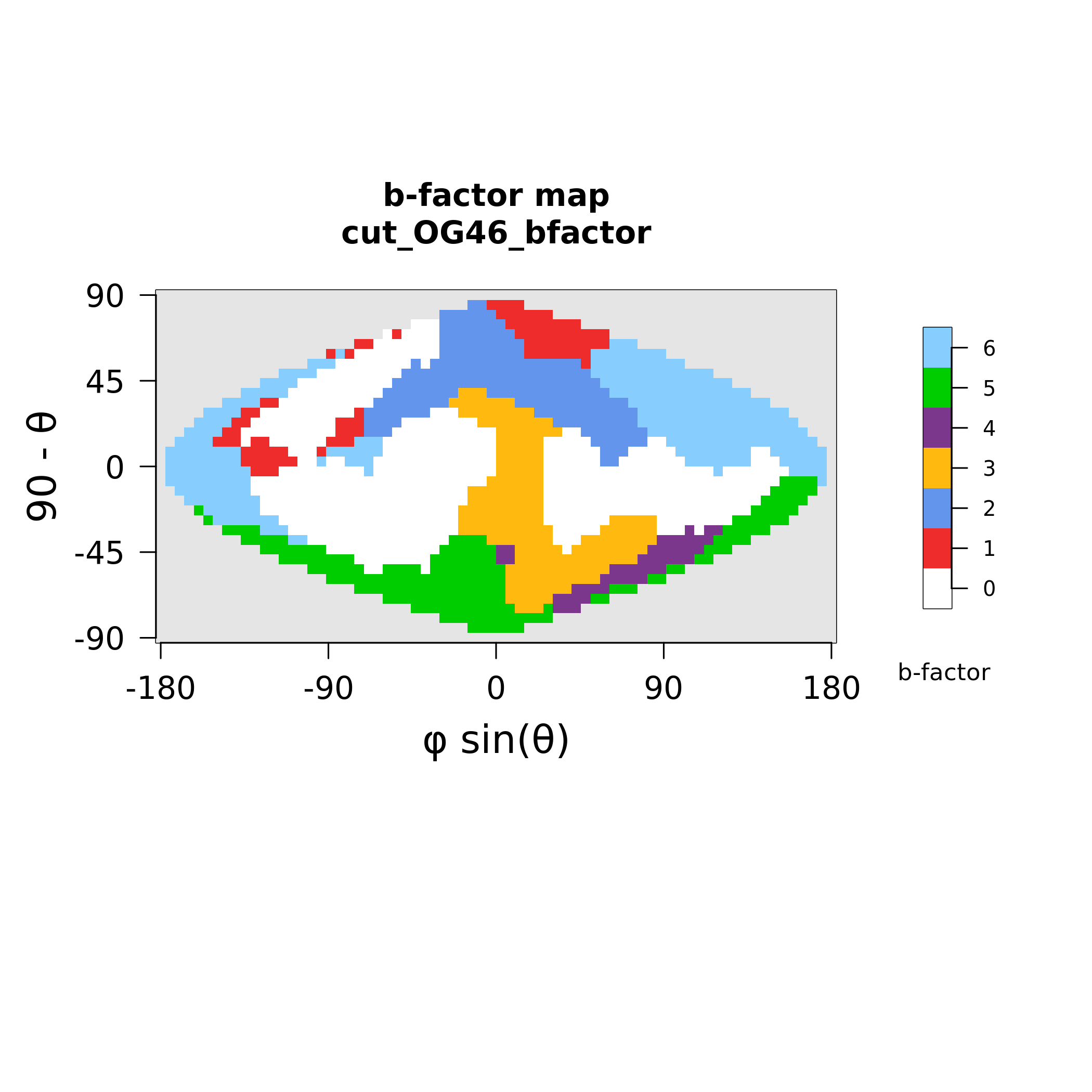

### MAX47_polymorphism.png

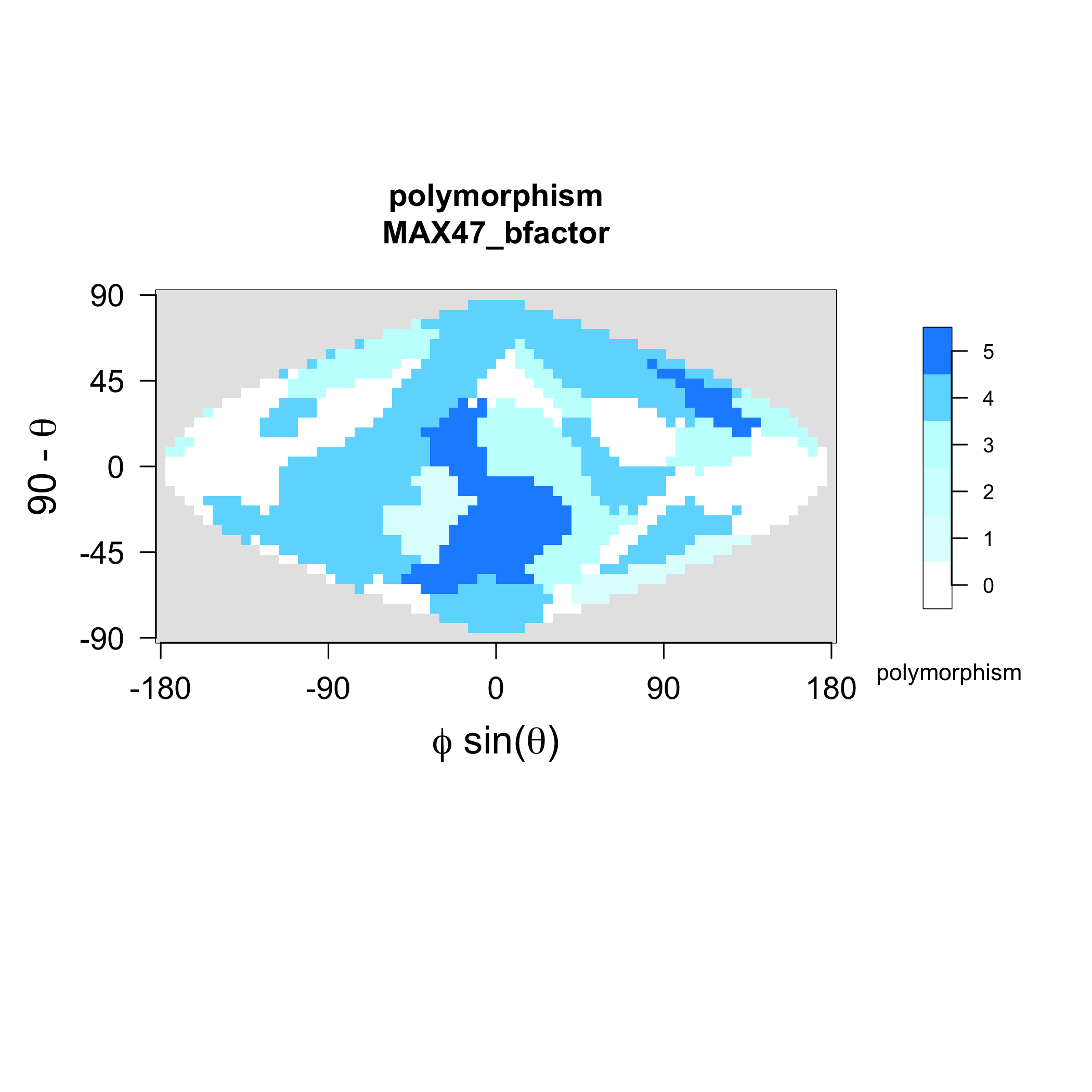

### MAX47_strands.png

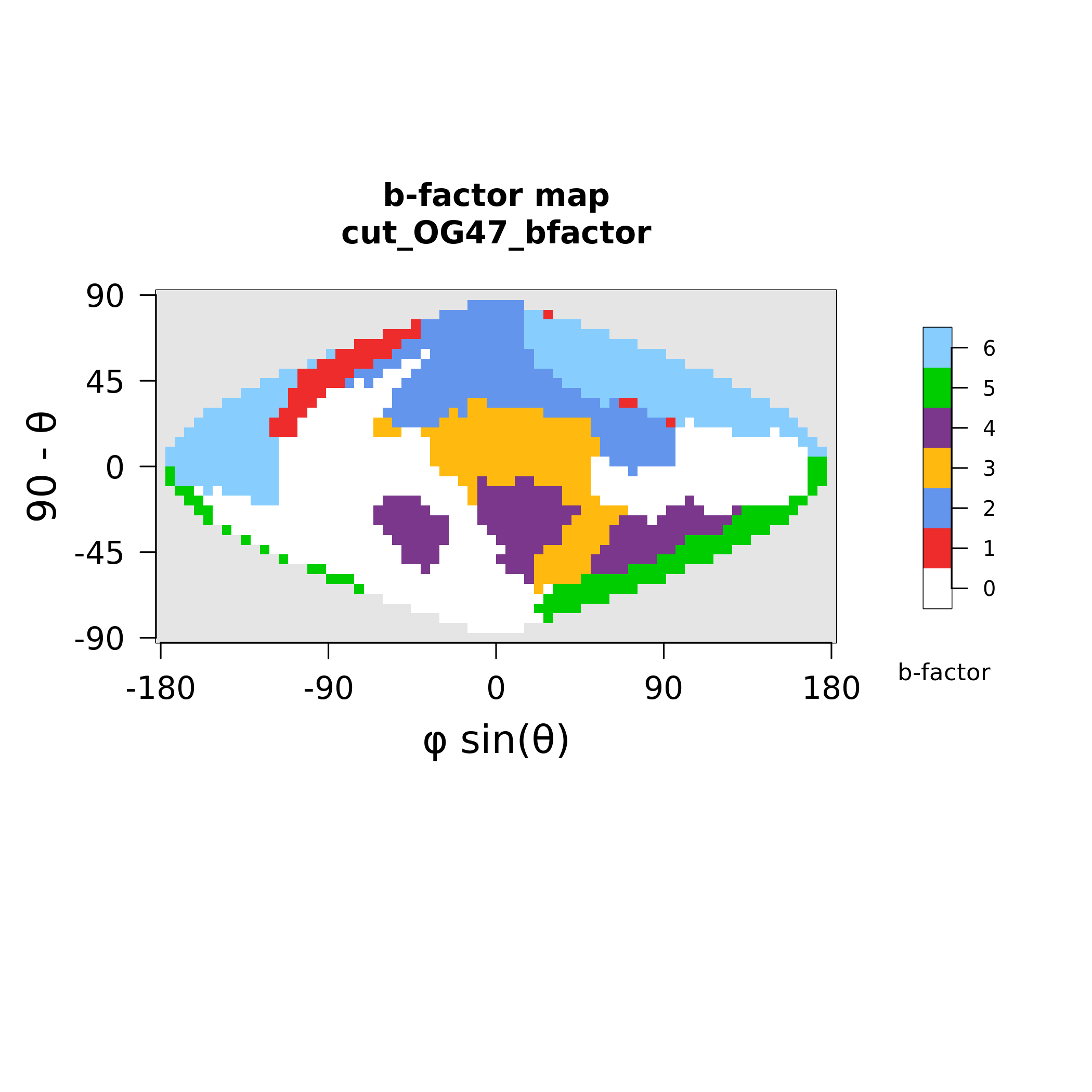

### MAX48_polymorphism.png

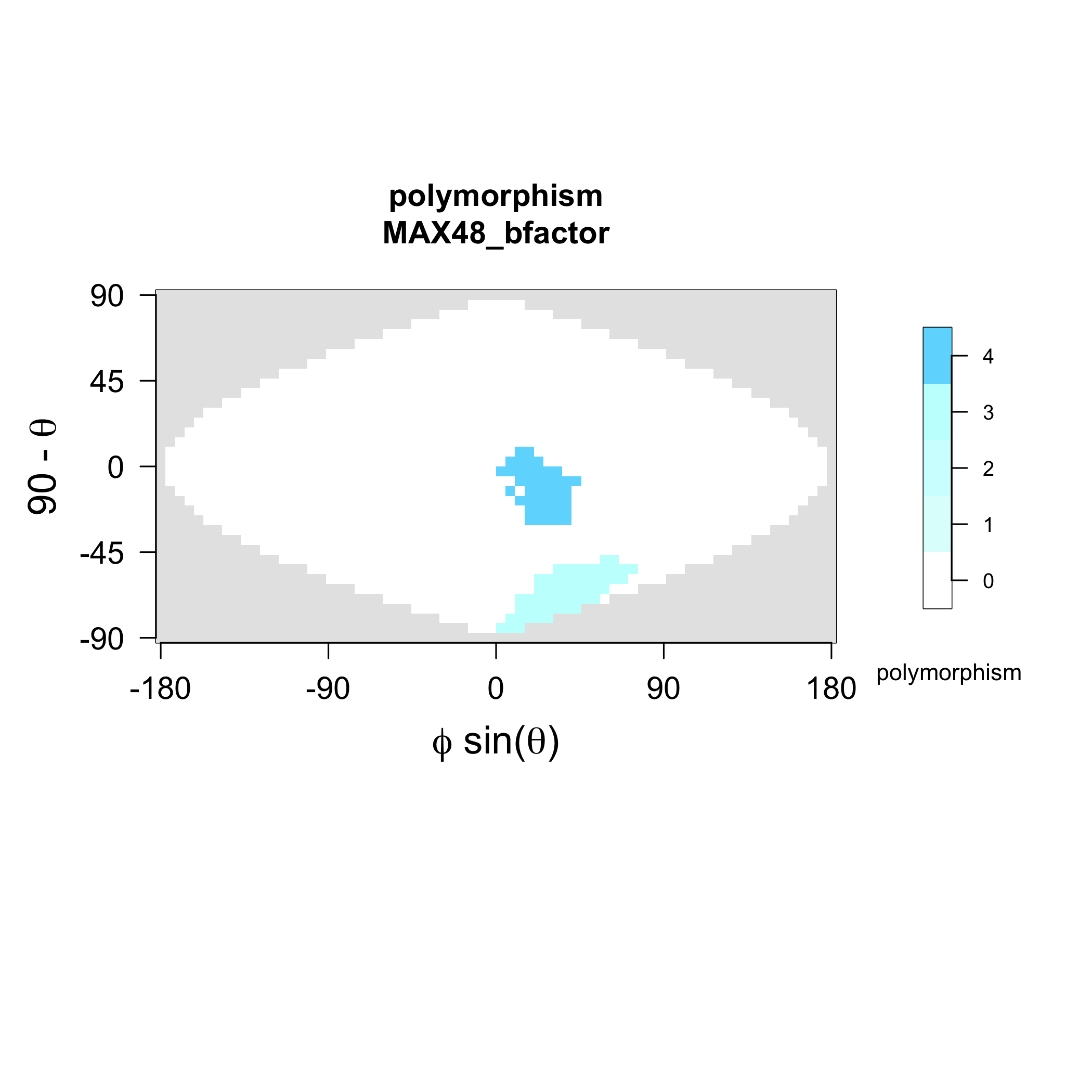

### MAX48_strands.png

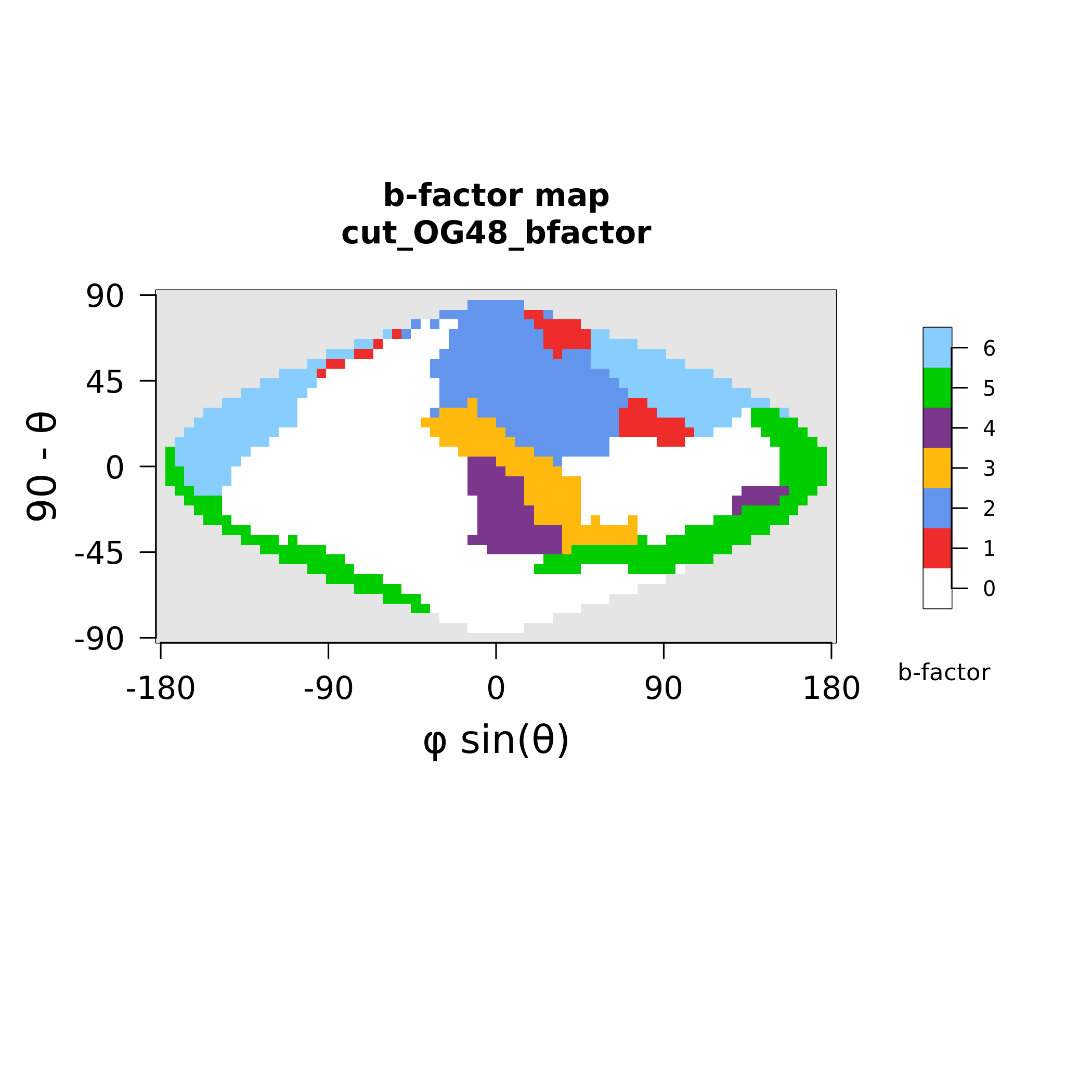

### MAX50_polymorphism.png

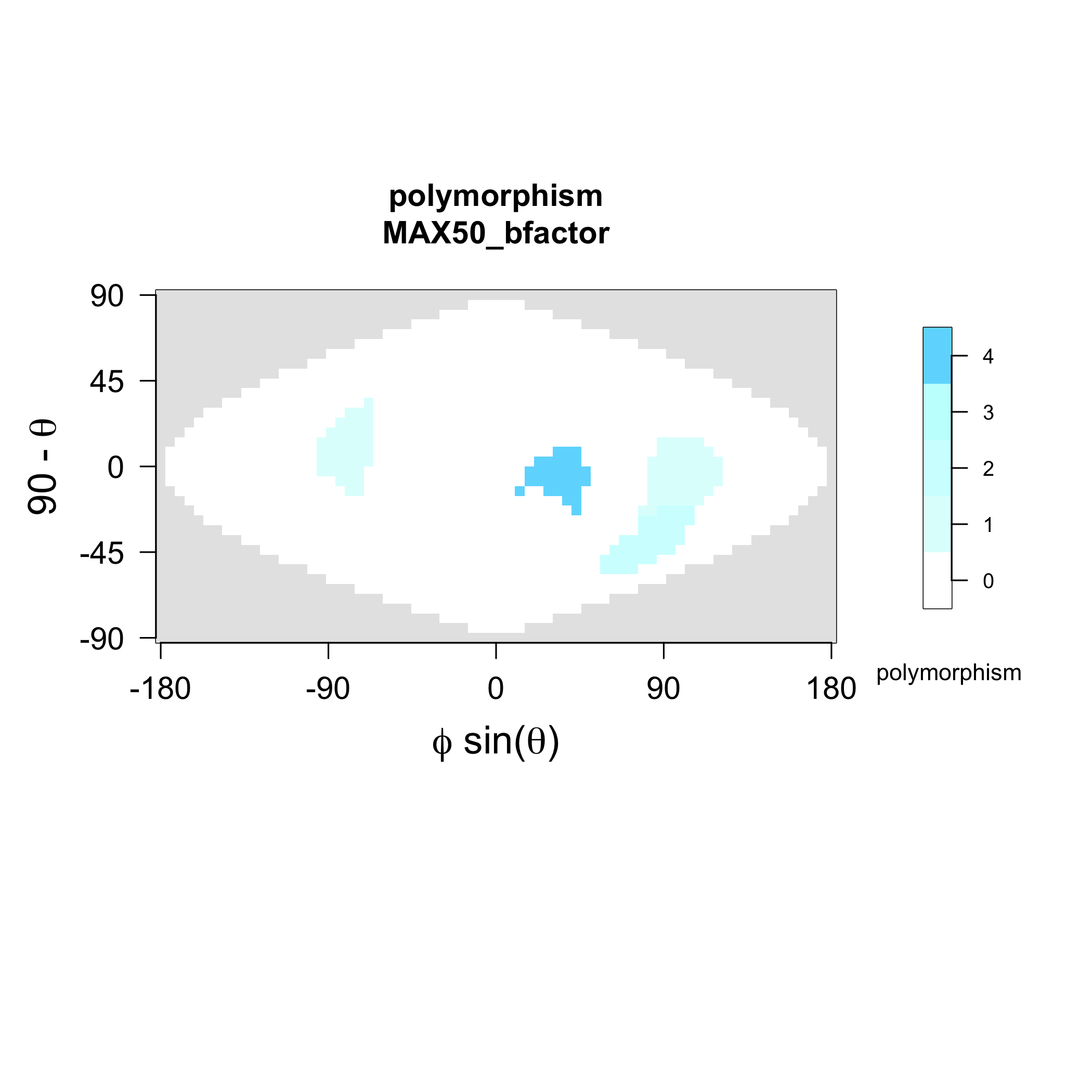

### MAX50_strands.png

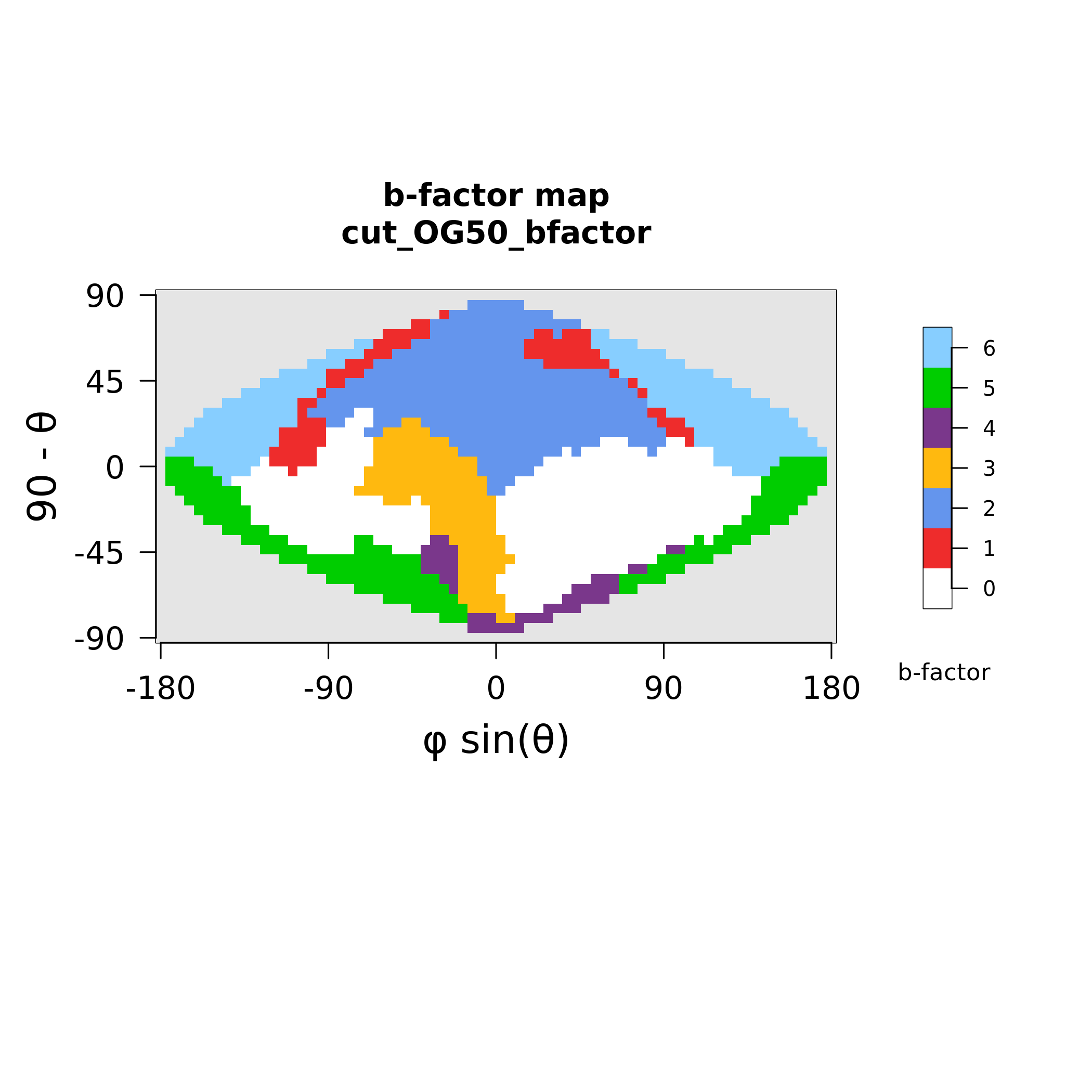

### MAX56_polymorphism.png

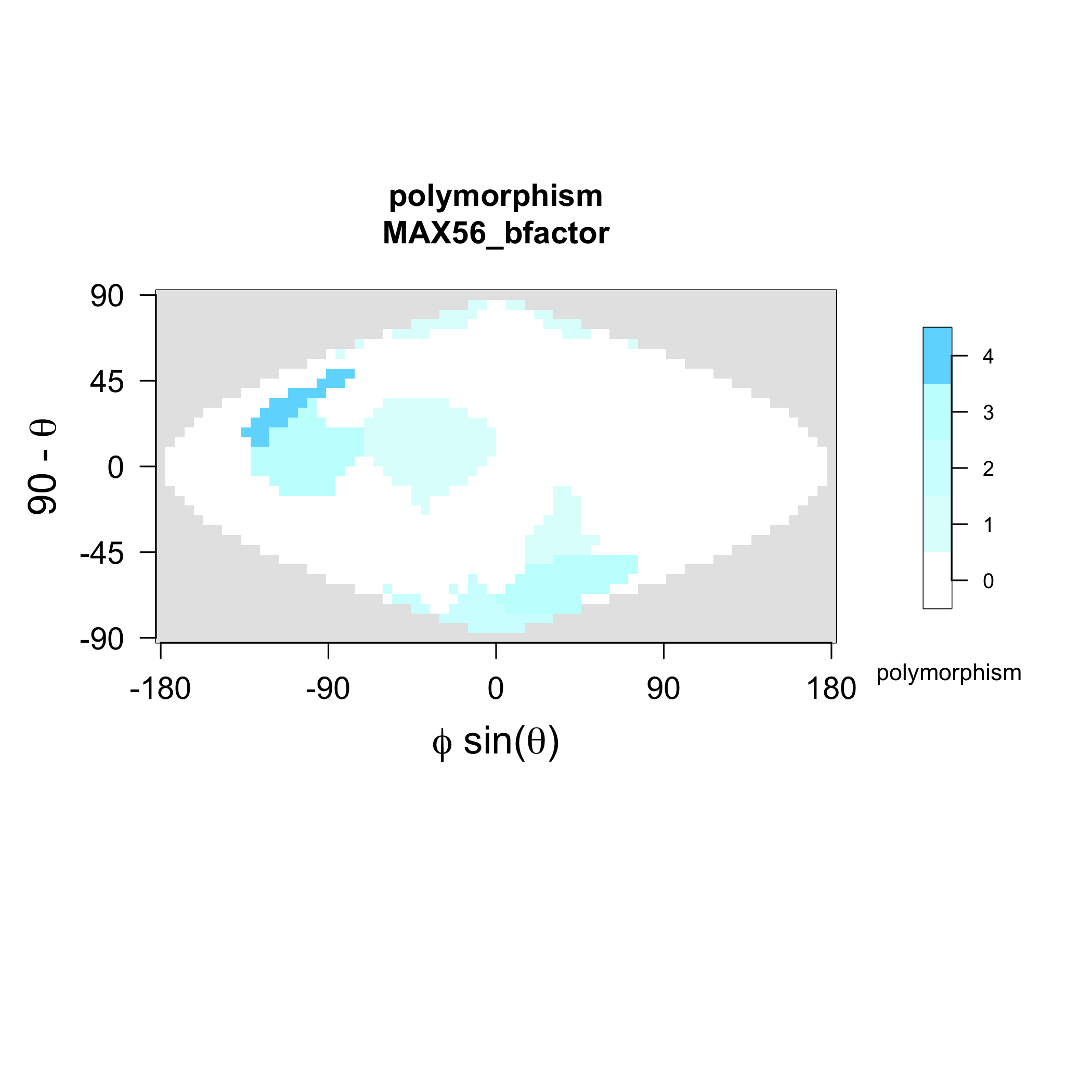

### MAX56_strands.png

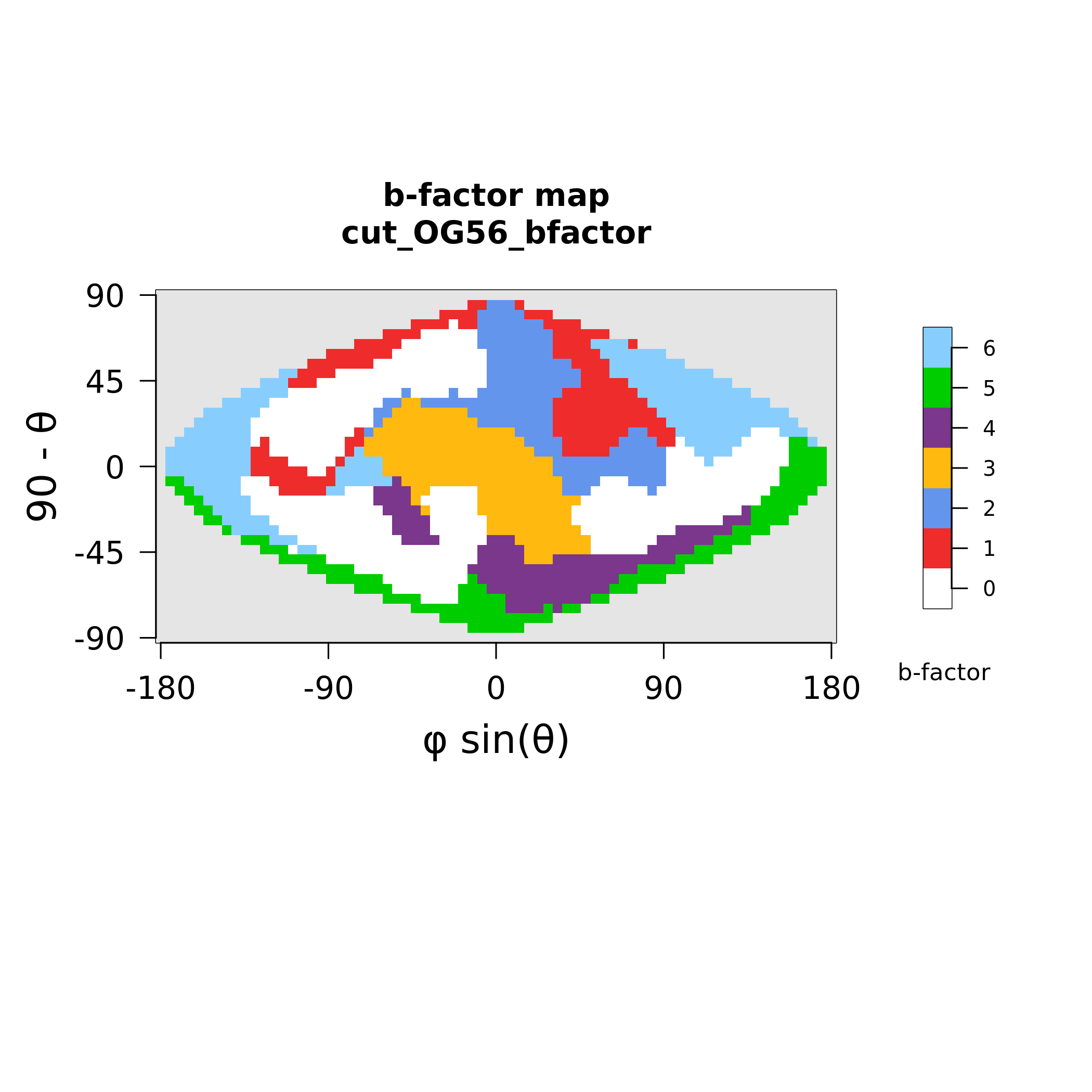

### MAX58_strands.png

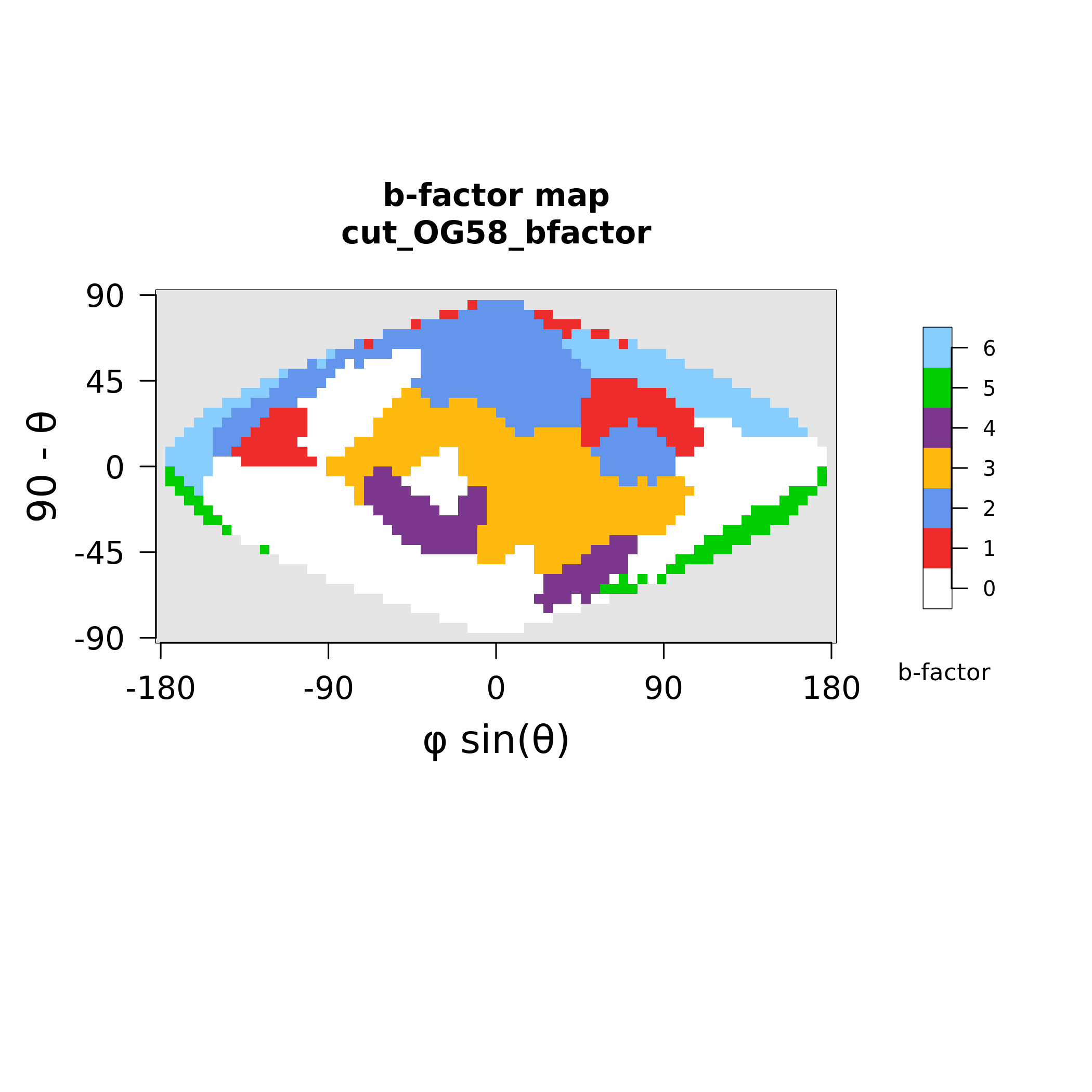

### MAX61_polymorphism.png

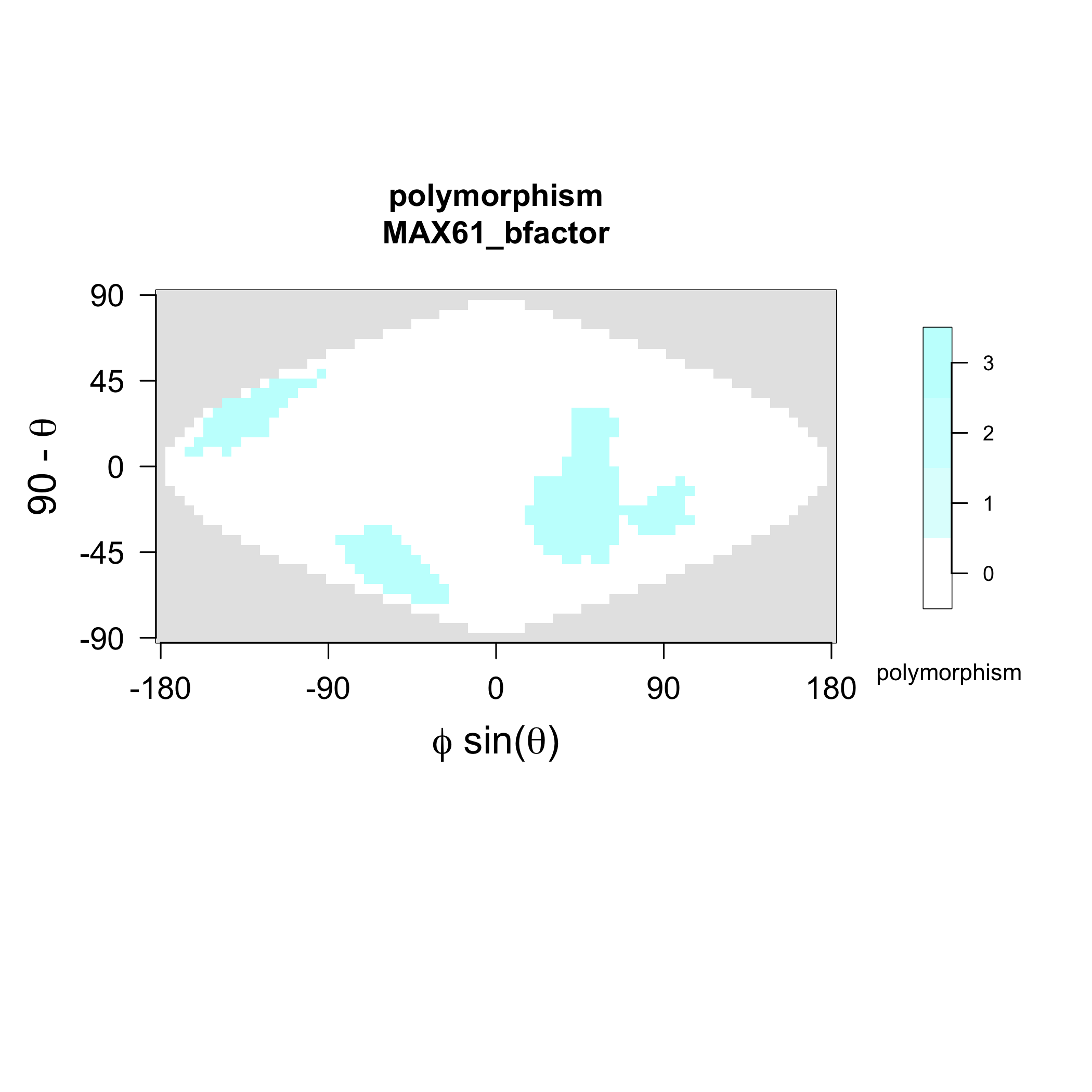

### MAX61_strands.png

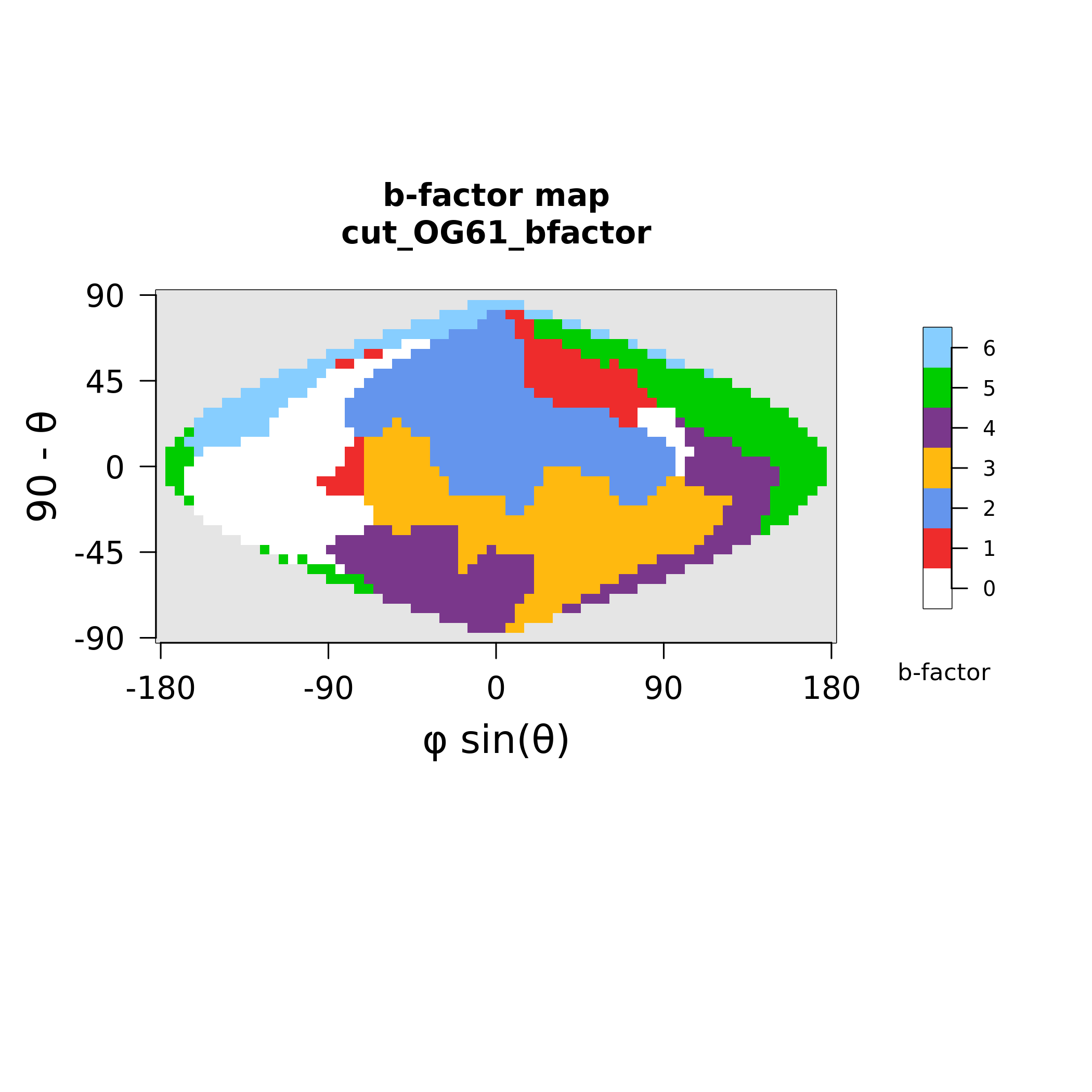

### MAX62_polymorphism.png

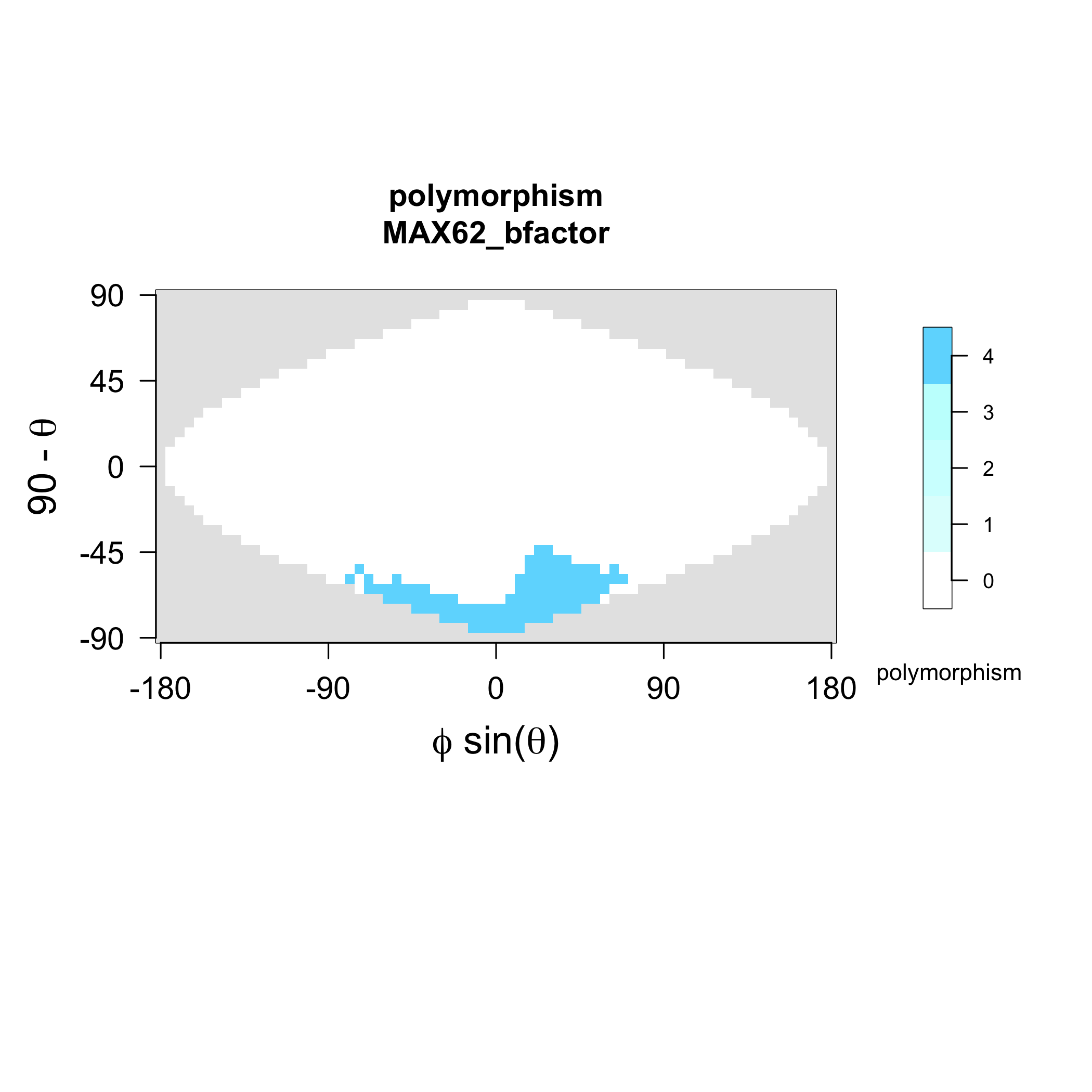

### MAX62_strands.png

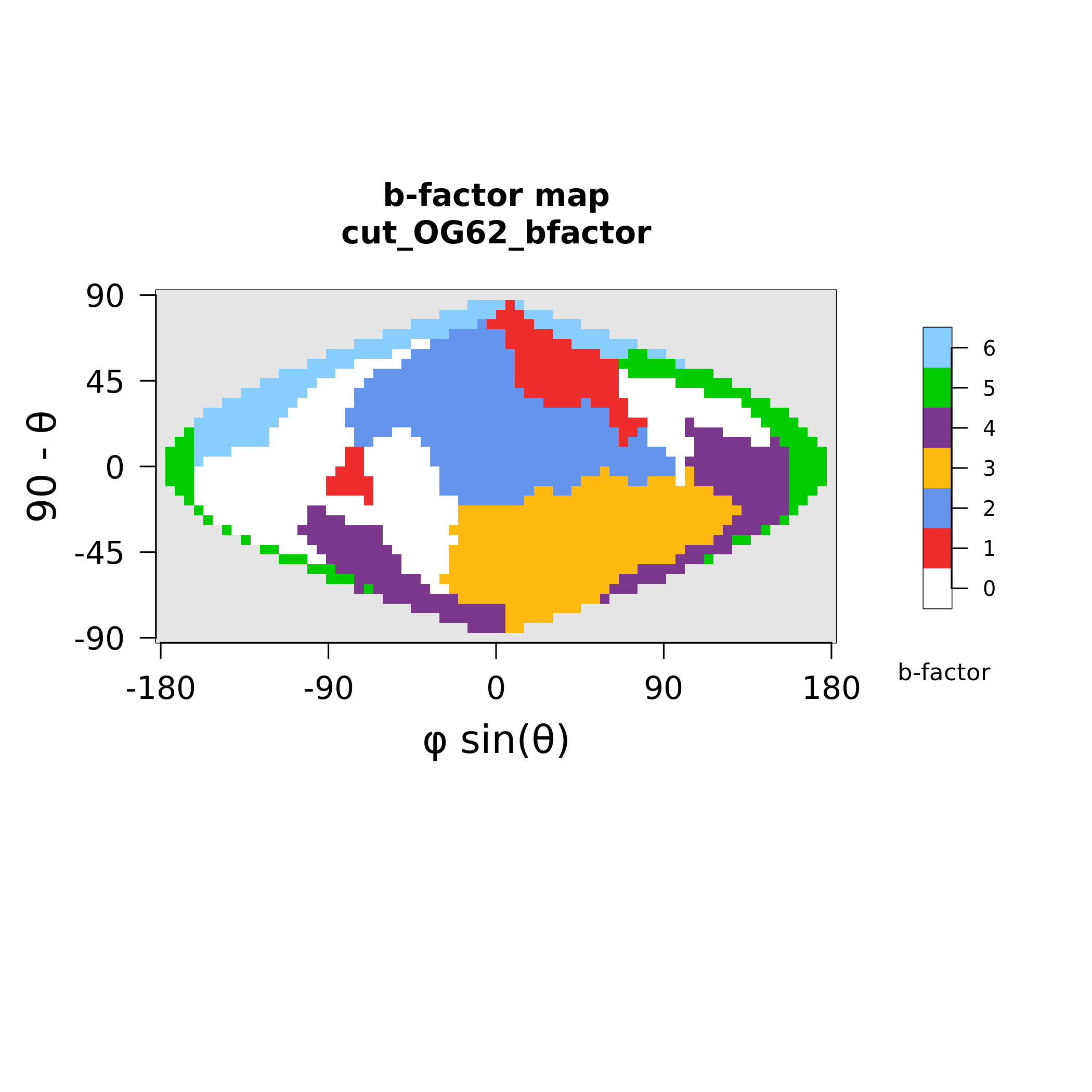

### MAX63_polymorphism.png

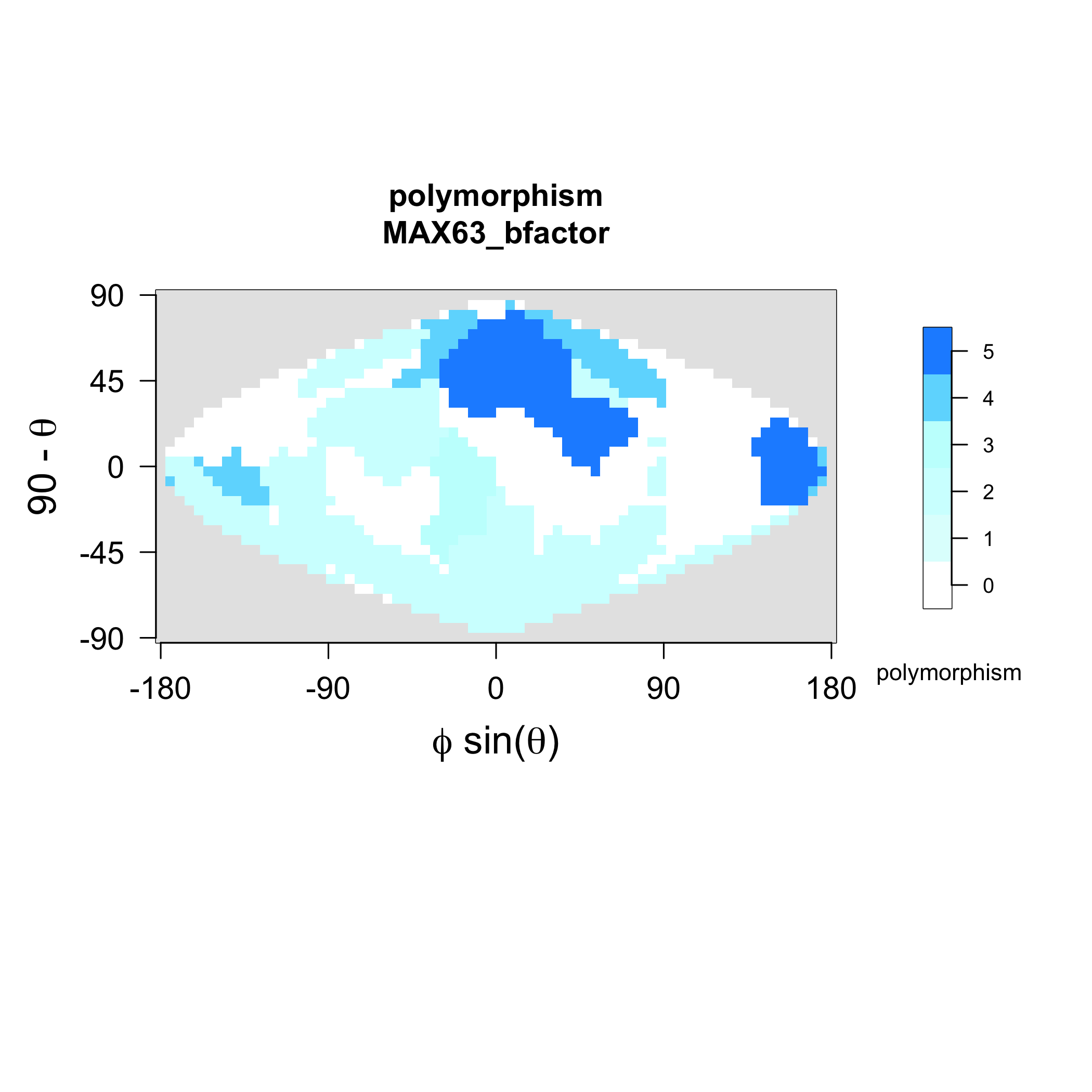

### MAX63_strands.png

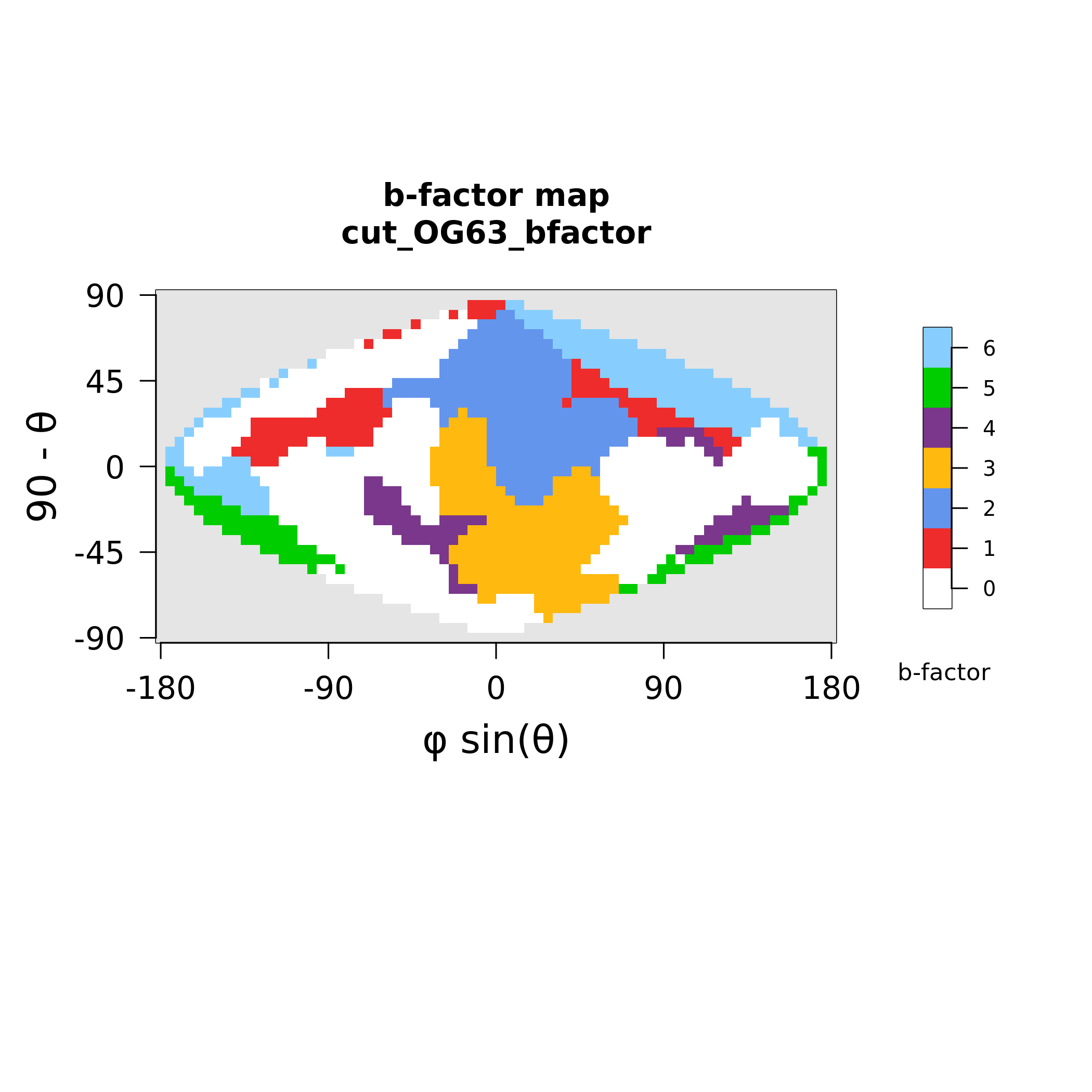

### MAX67_polymorphism.png

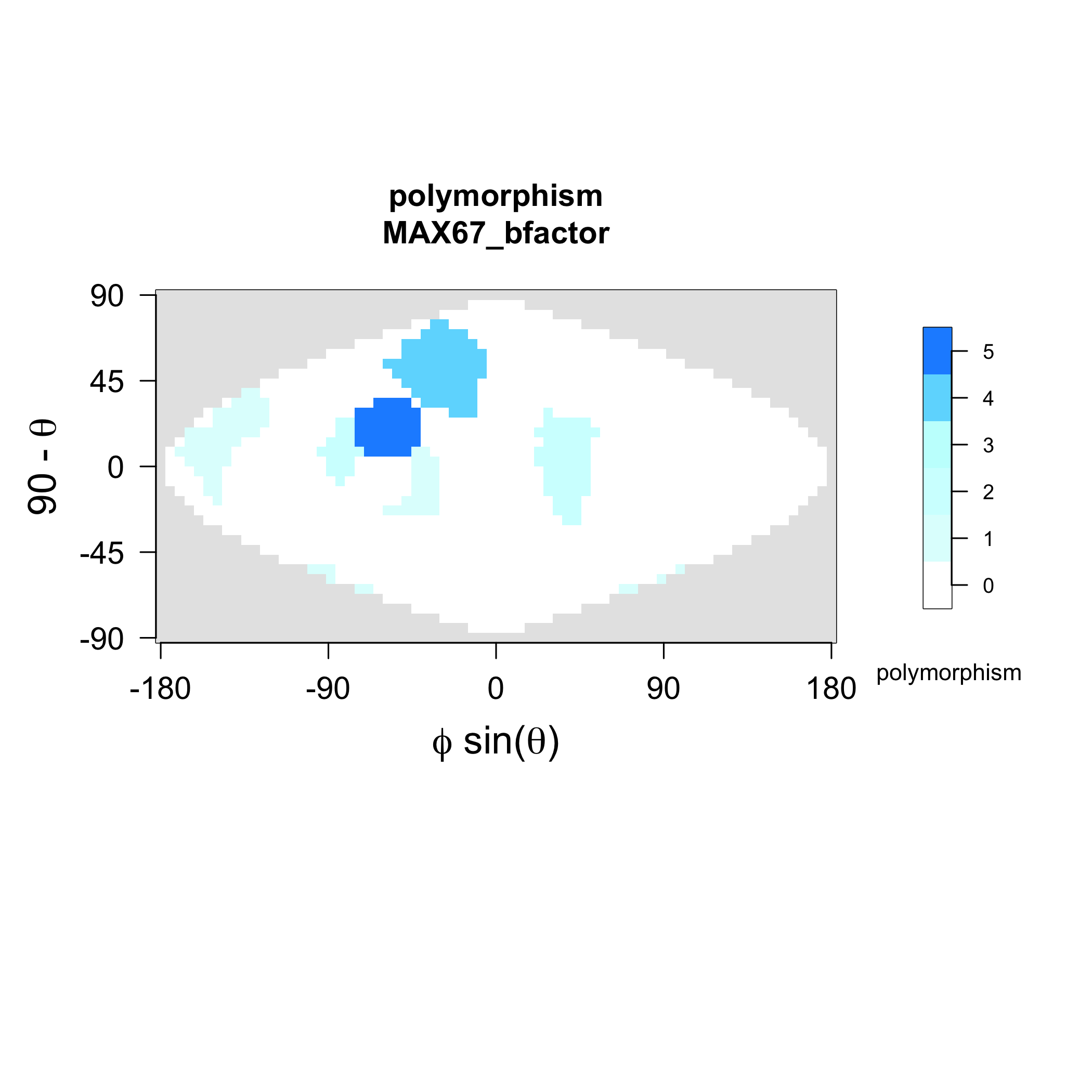

### MAX67_strands.png

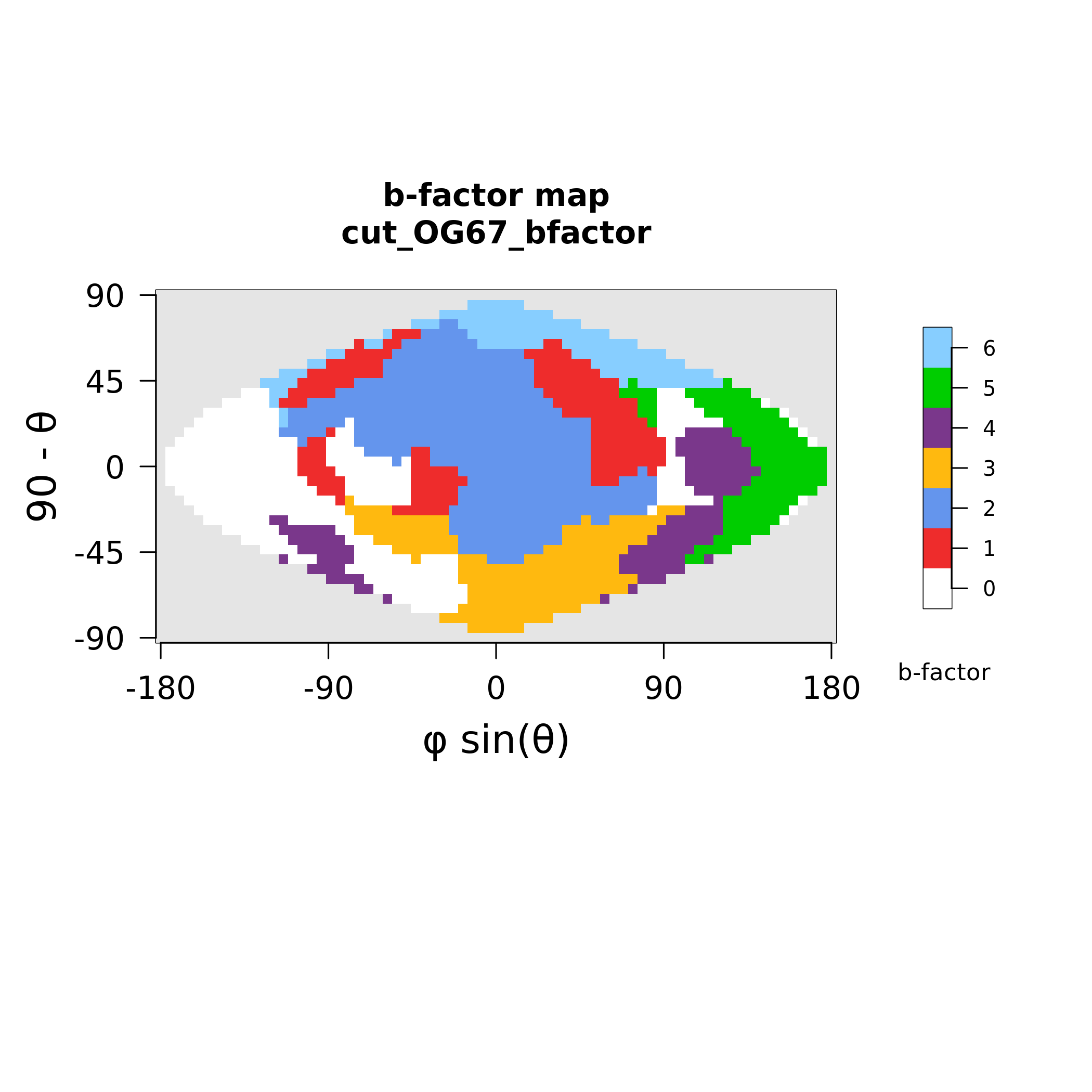

### MAX69_polymorphism.png

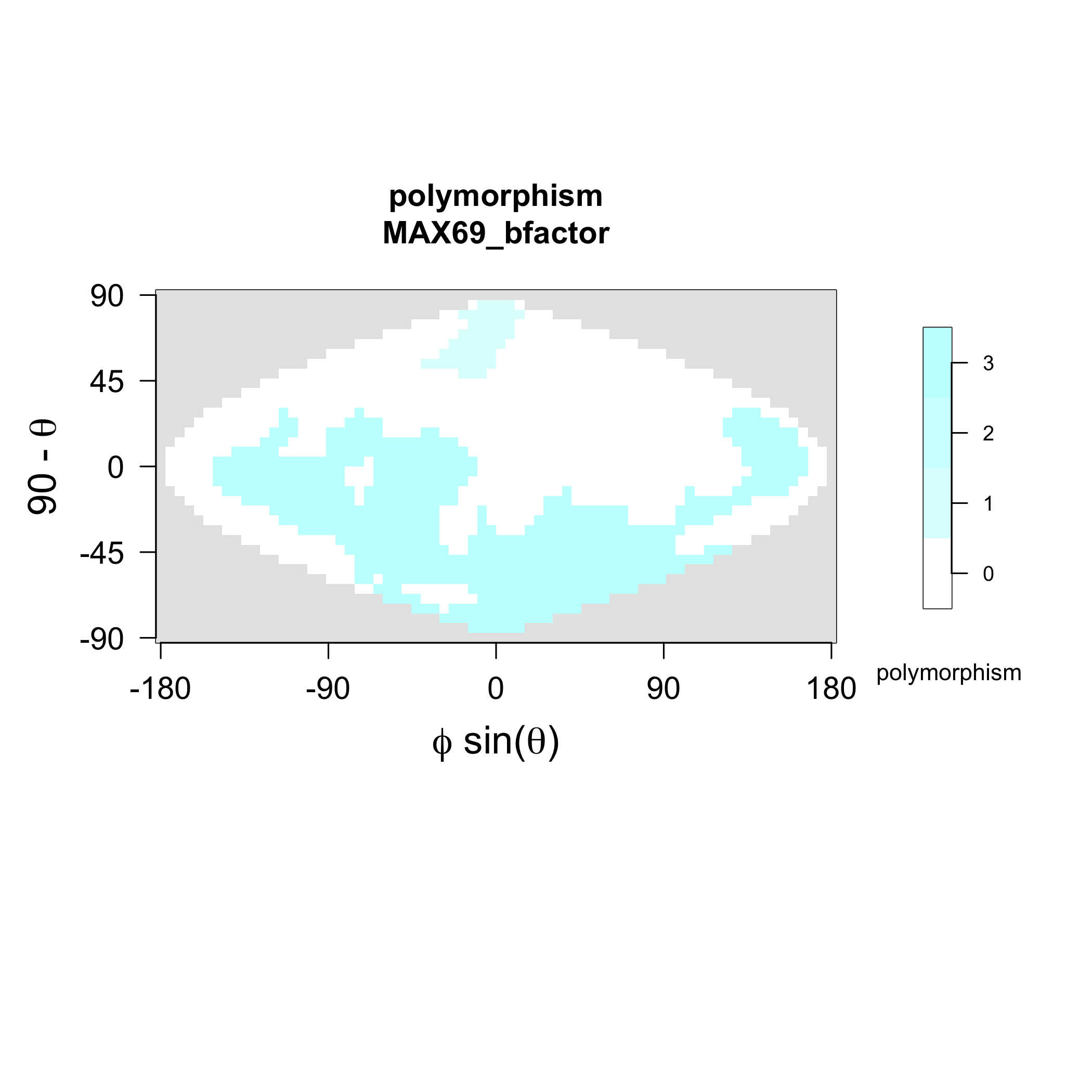

### MAX69_strands.png

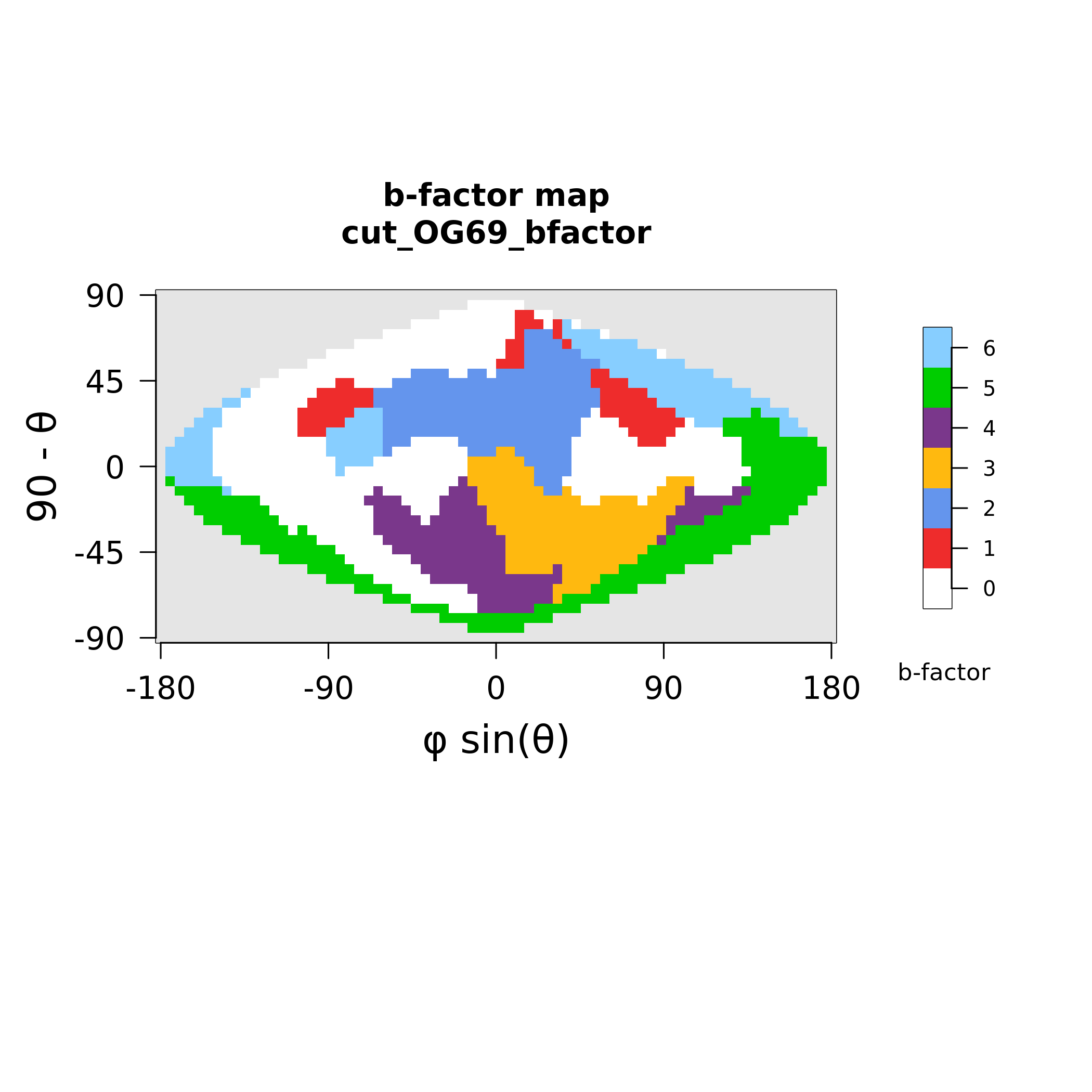

### MAX70_polymorphism.png

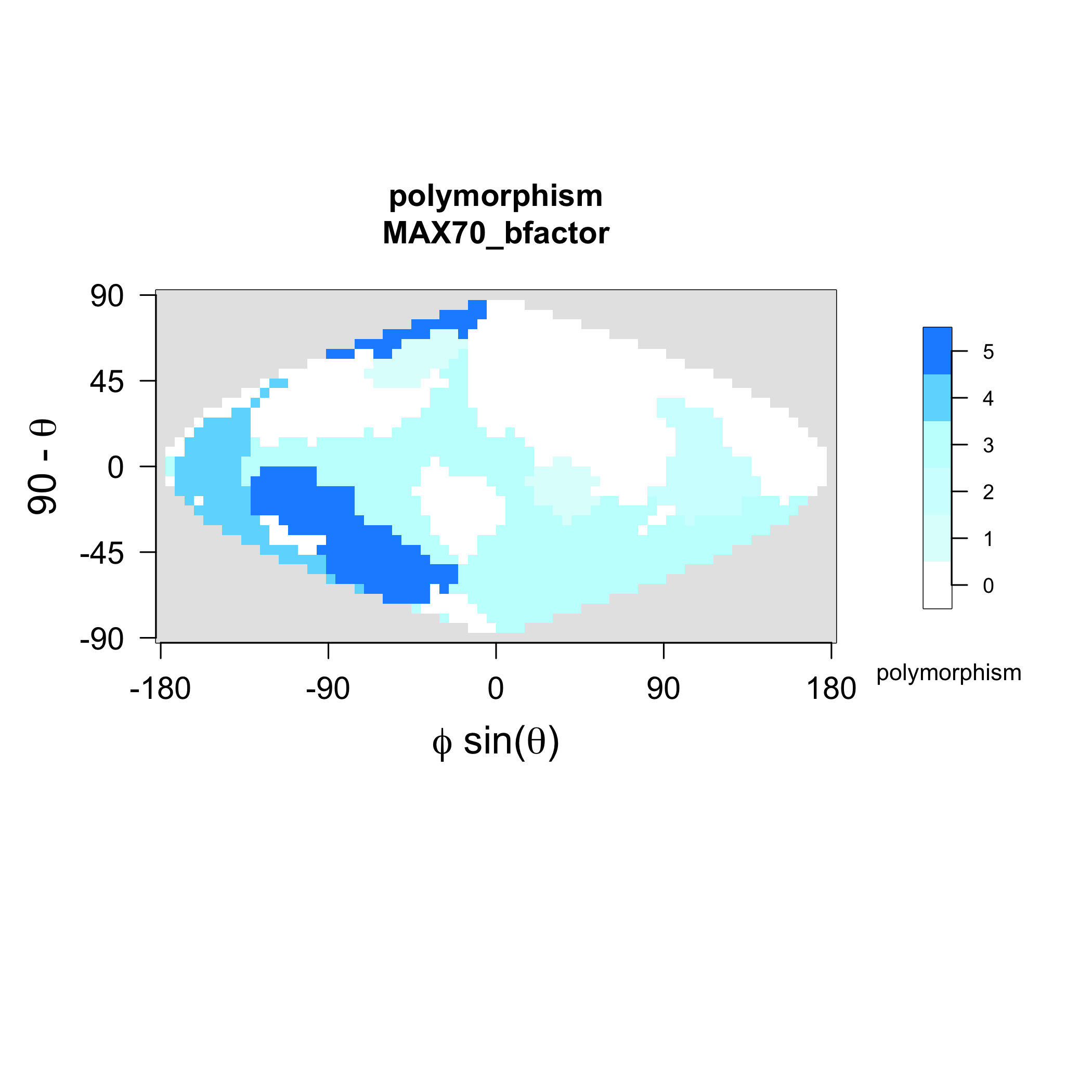

### MAX70_strands.png

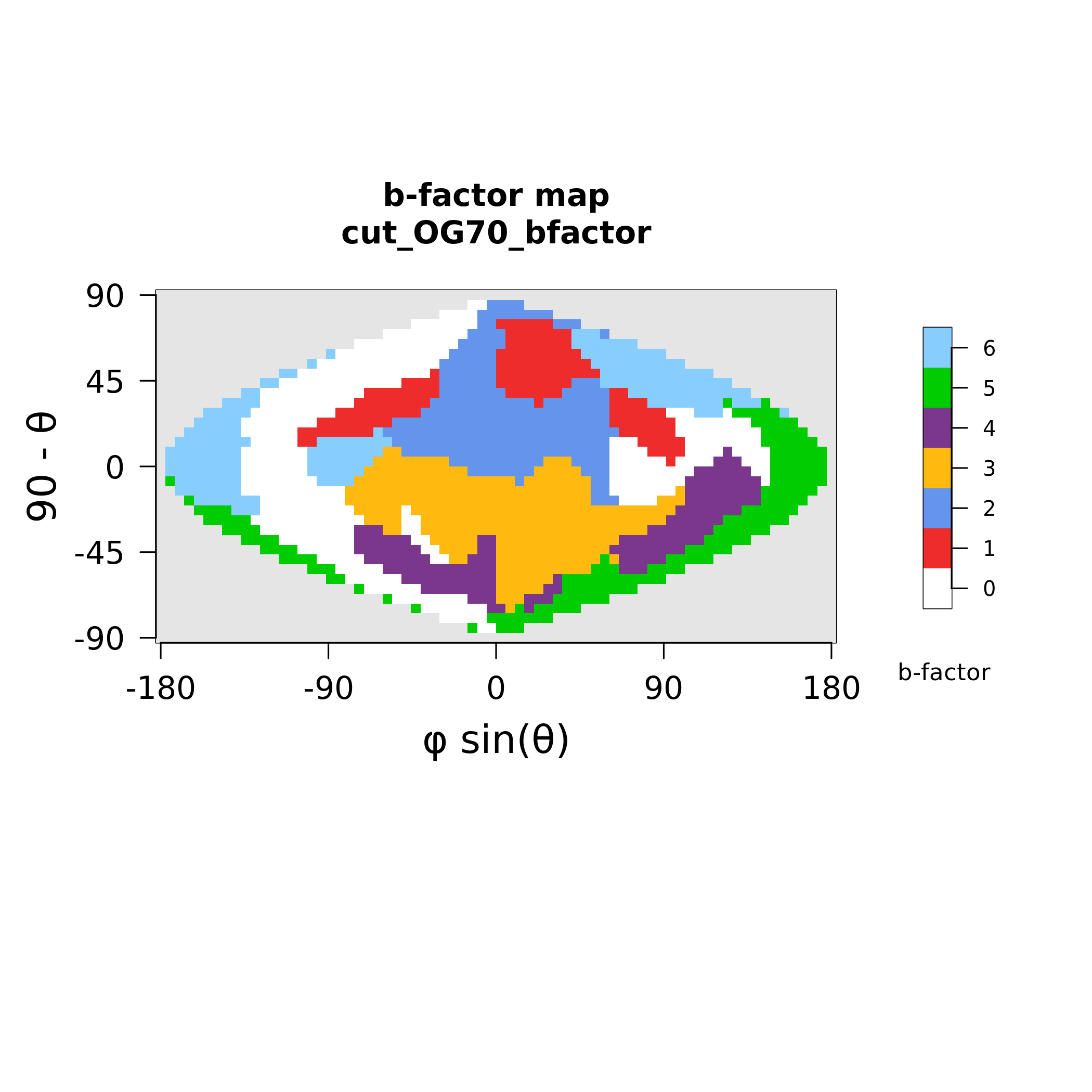

### MAX72_polymorphism.png

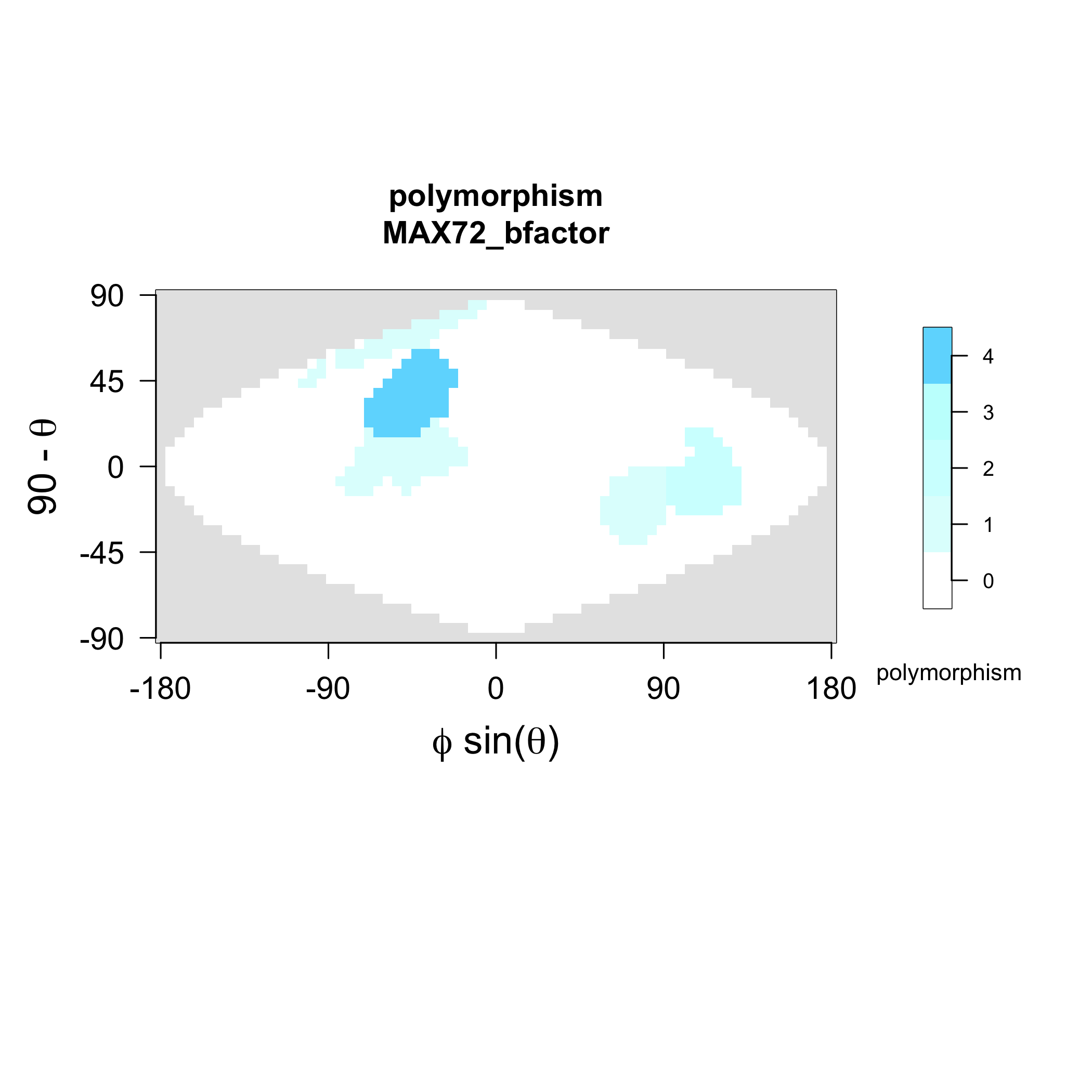

### MAX72_strands.png

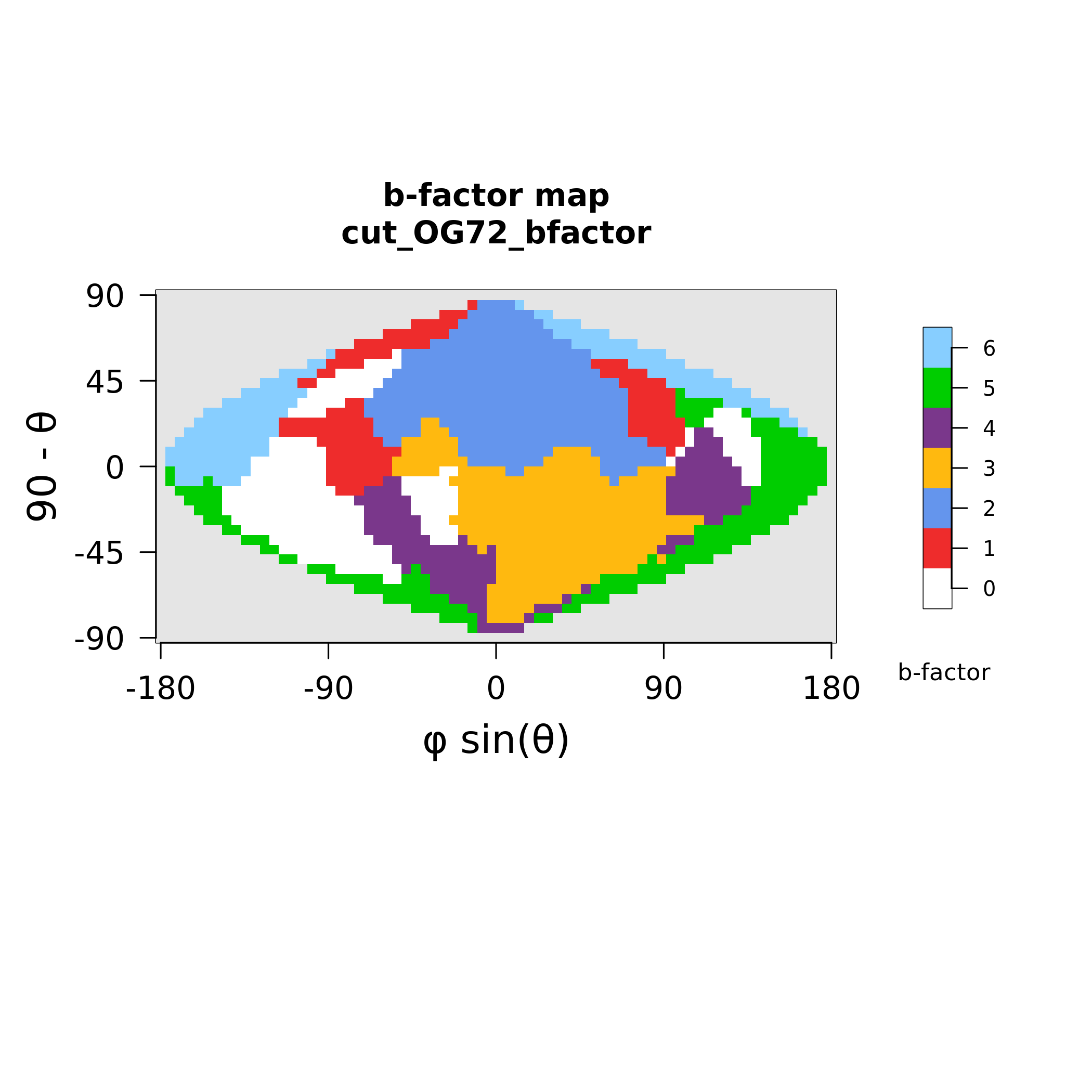

### MAX76_polymorphism.png

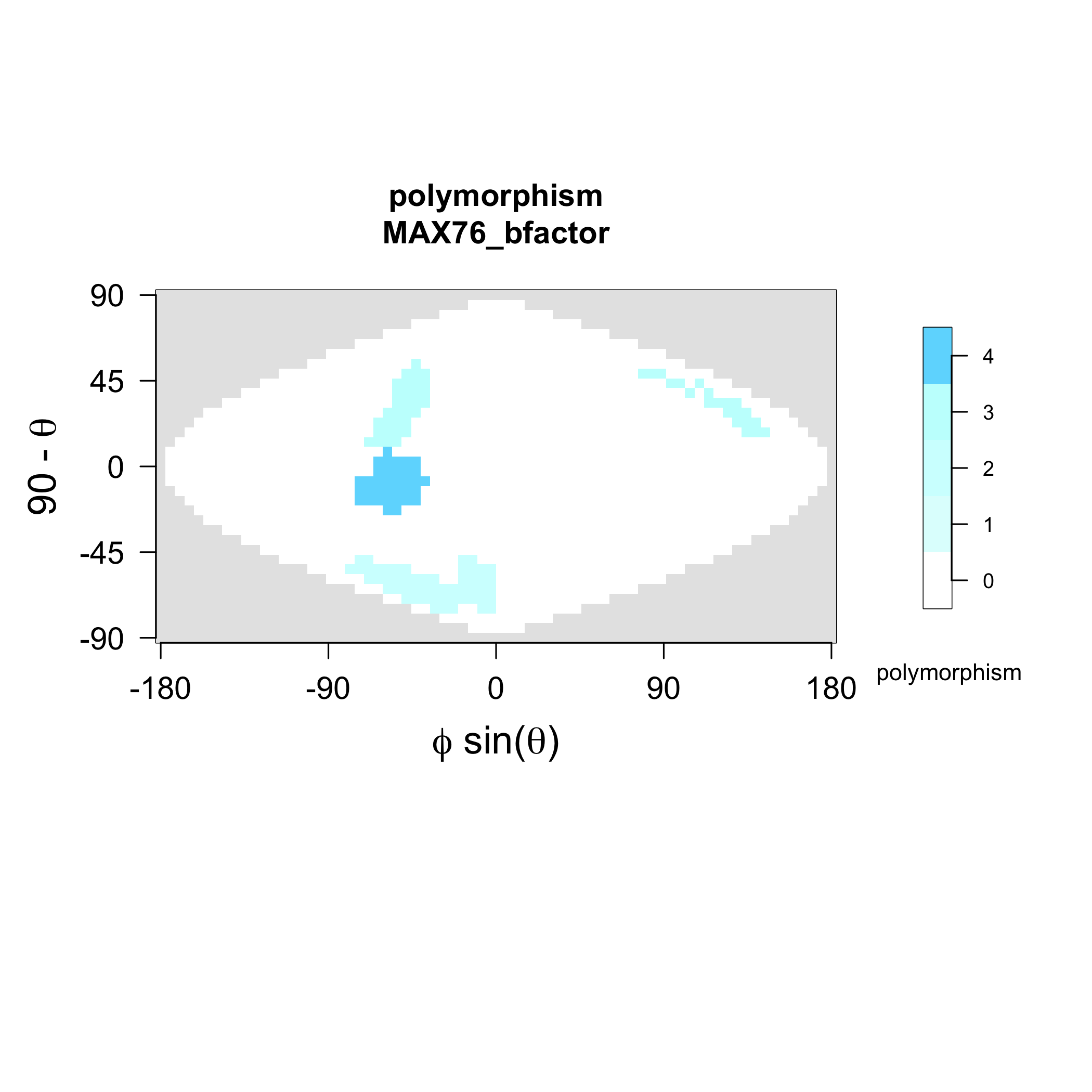

### MAX76_strands.png

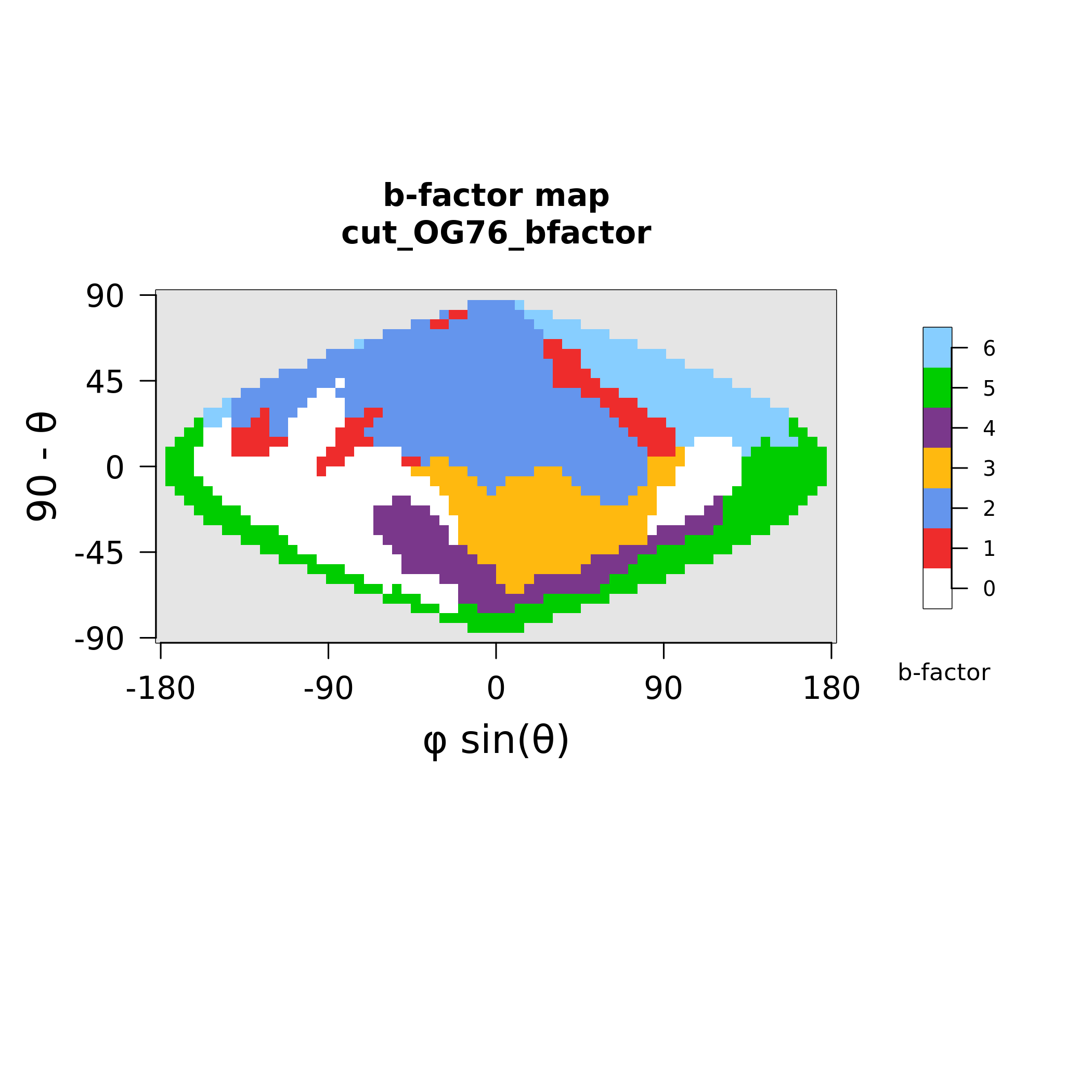

### MAX78_polymorphism.png

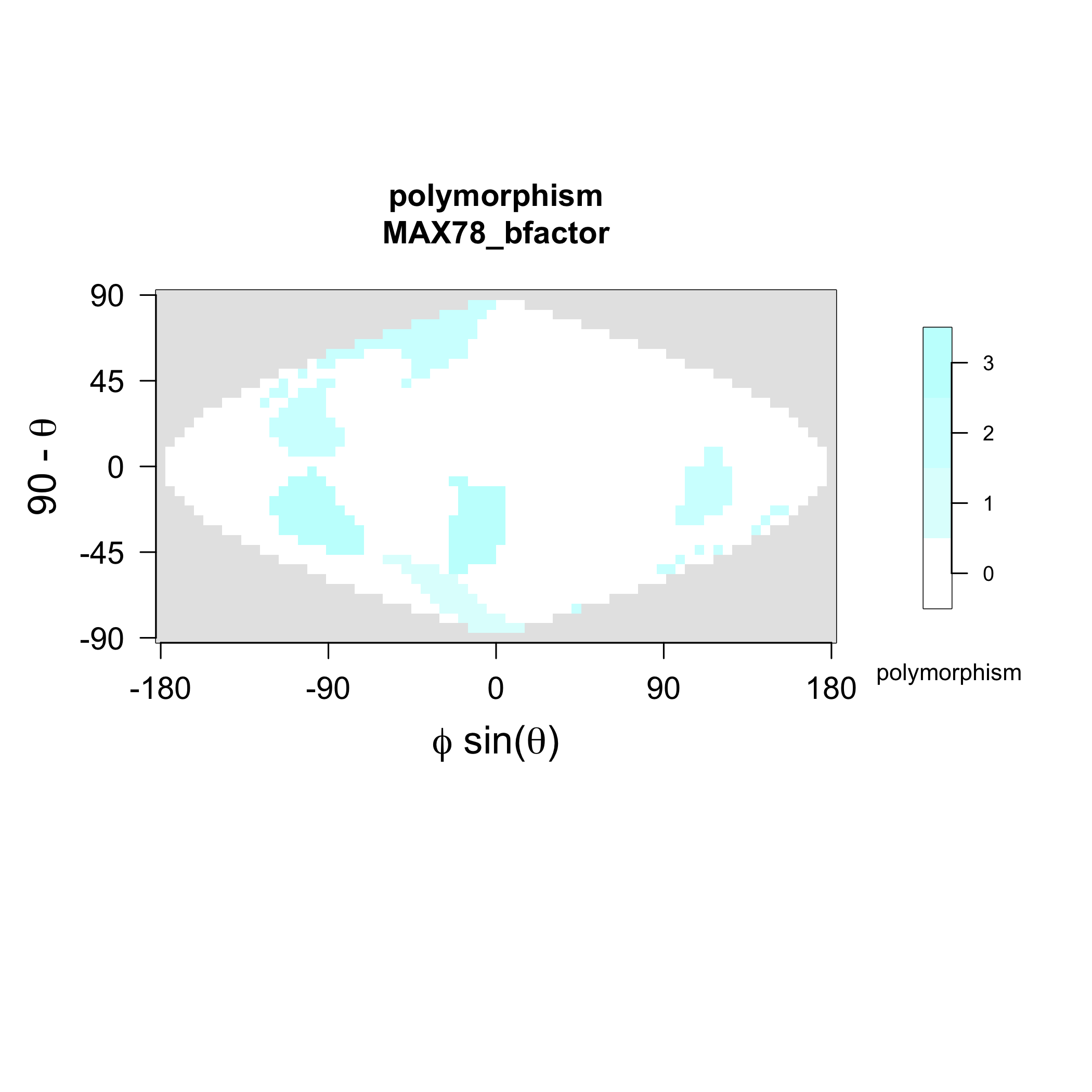

### MAX78_strands.png

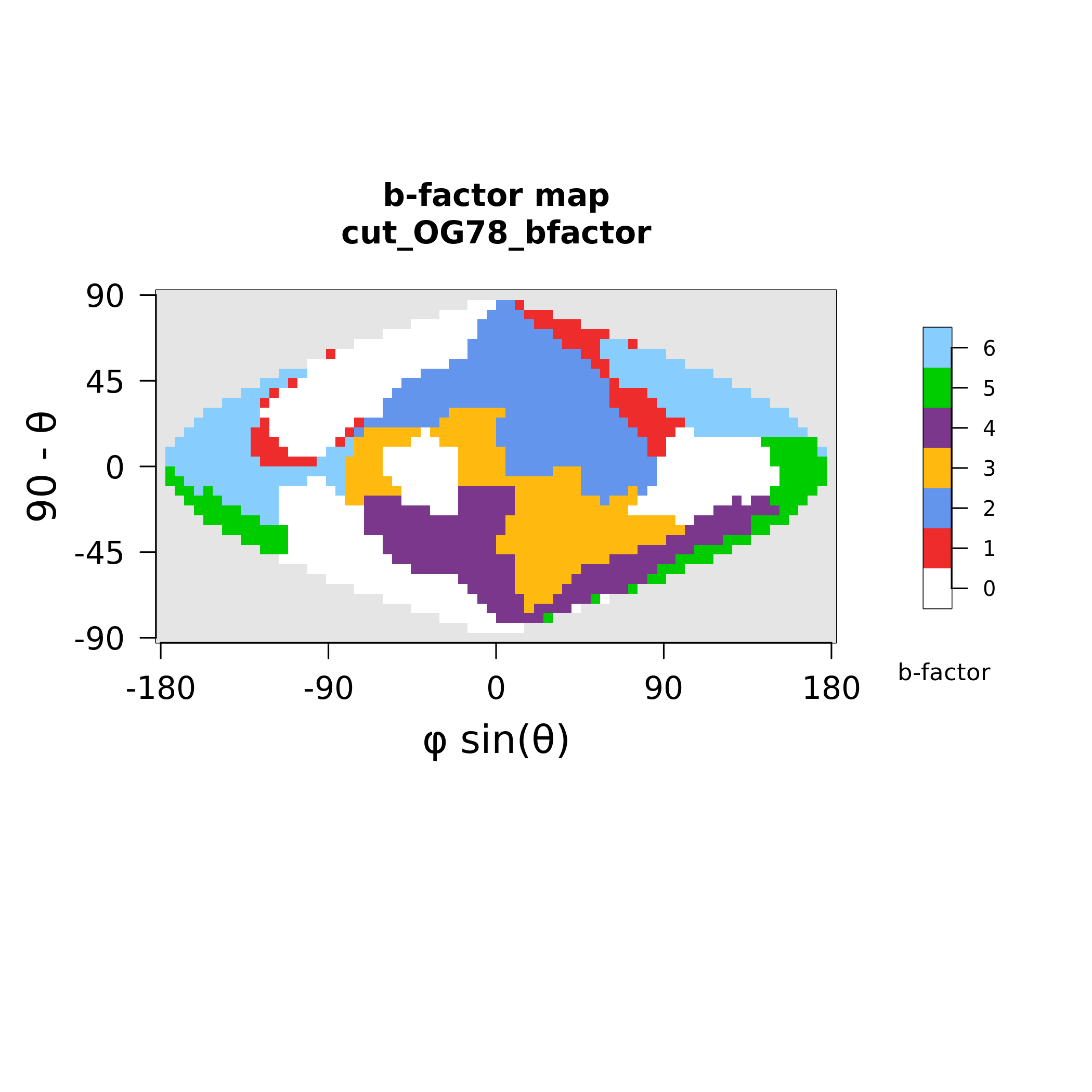

### MAX80_strands.png

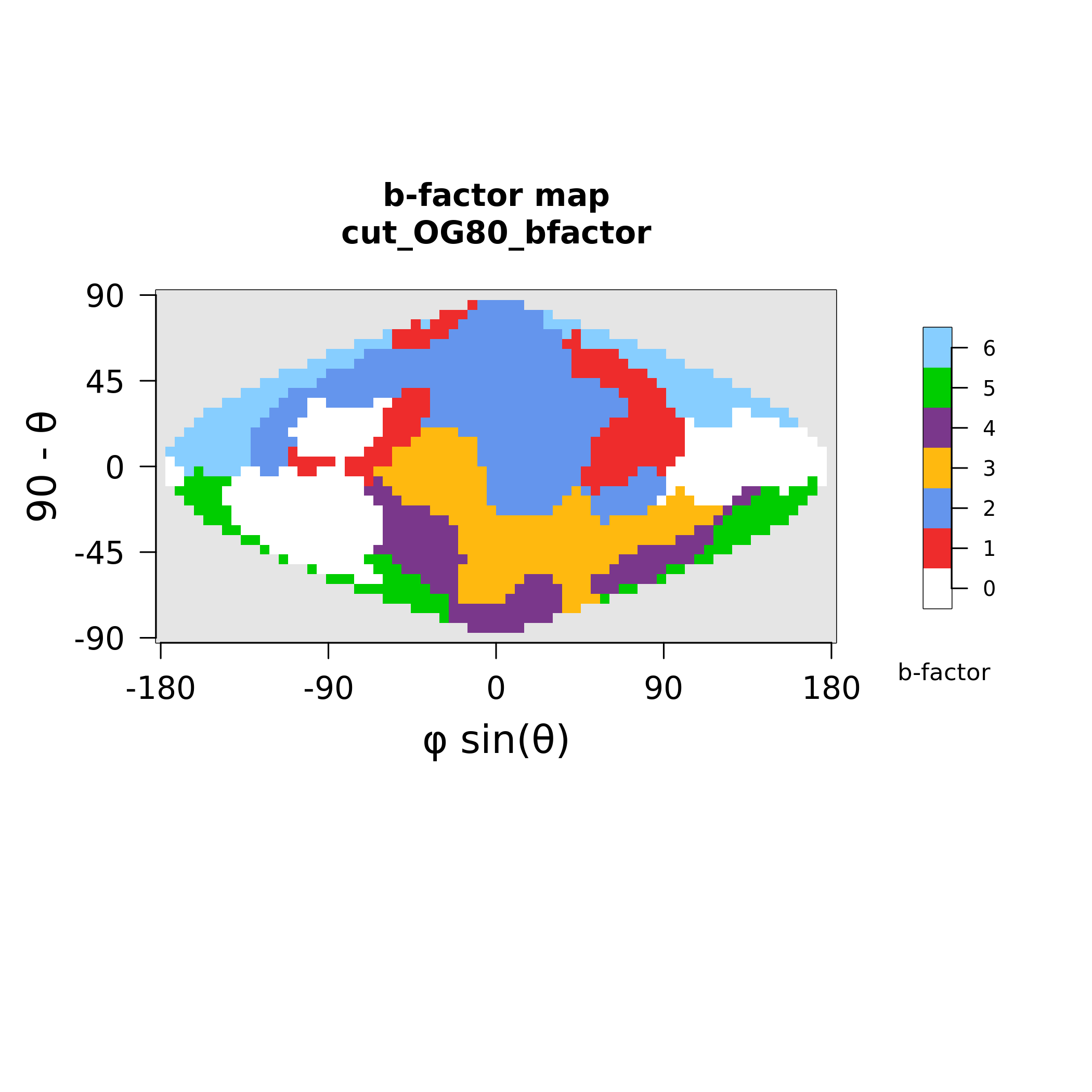

### MAX82_polymorphism.png

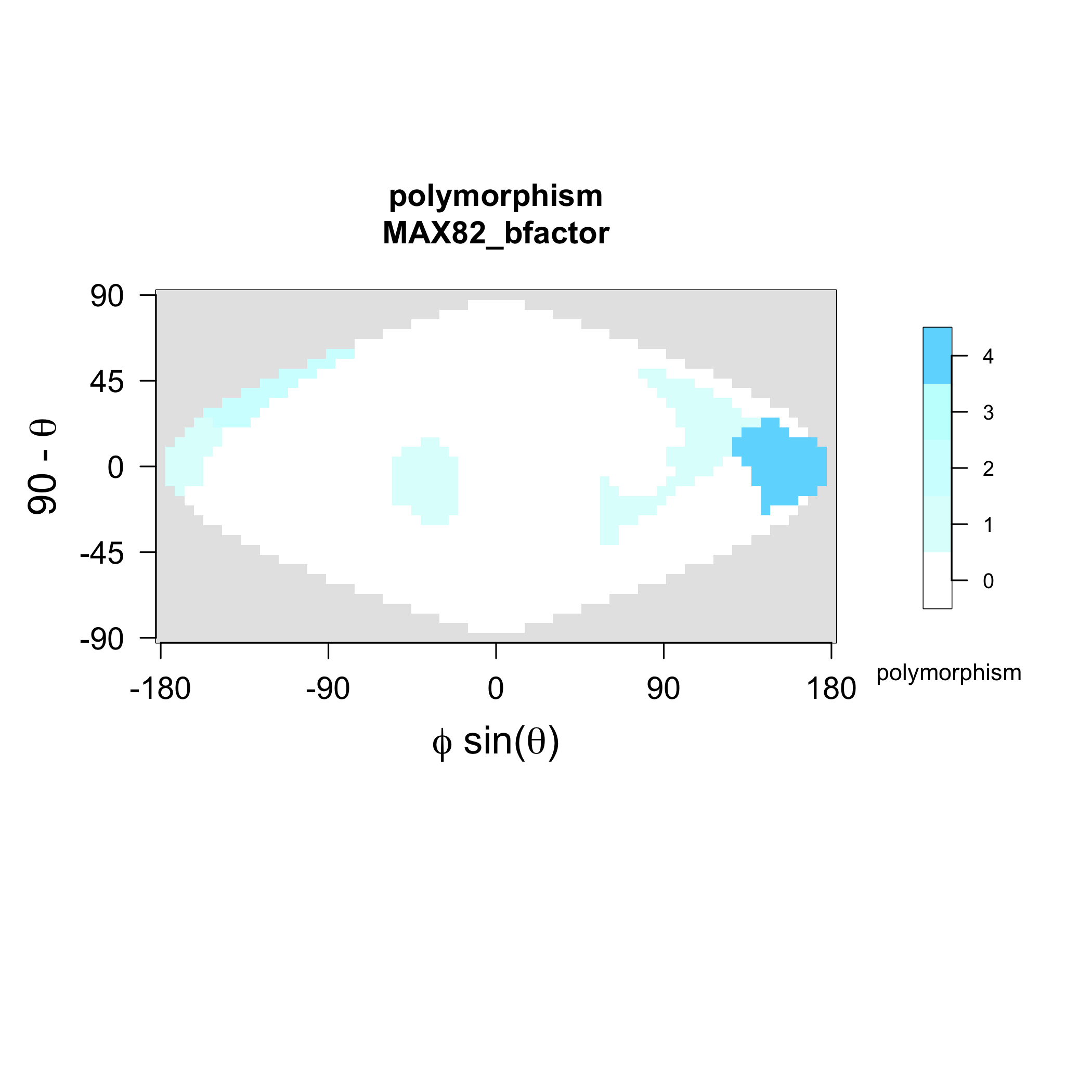

### MAX82_strands.png

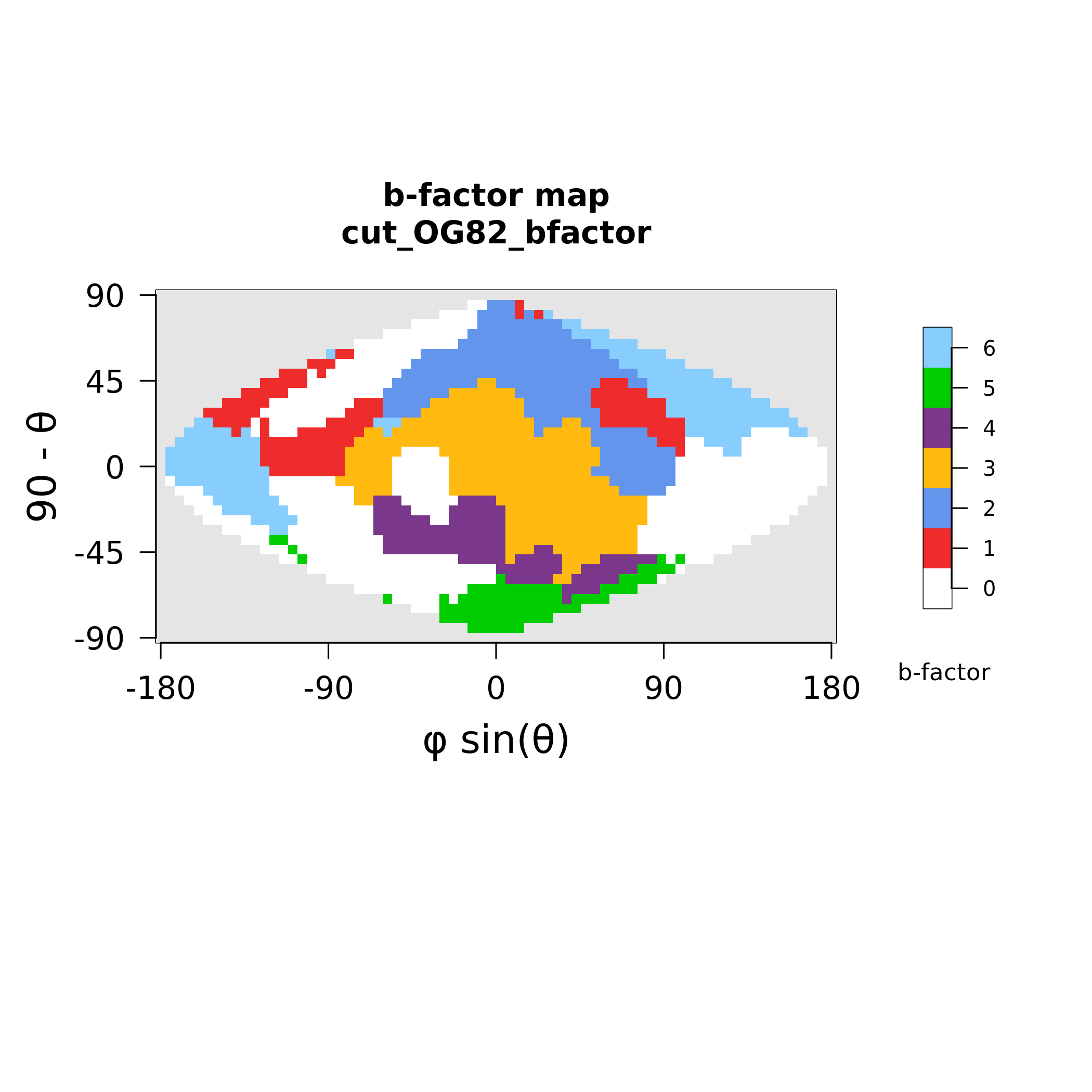

### MAX86_polymorphism.png

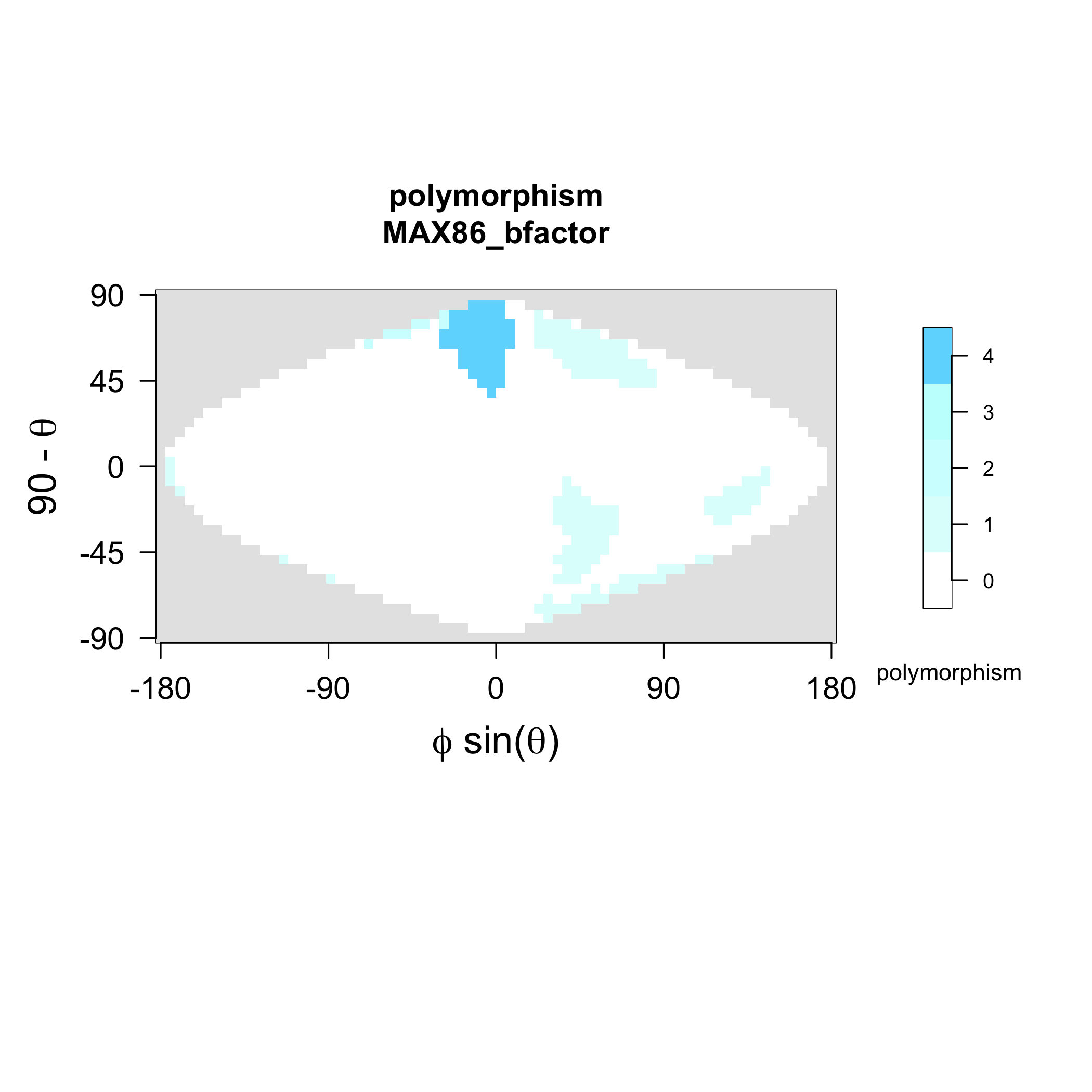

### MAX86_strands.png

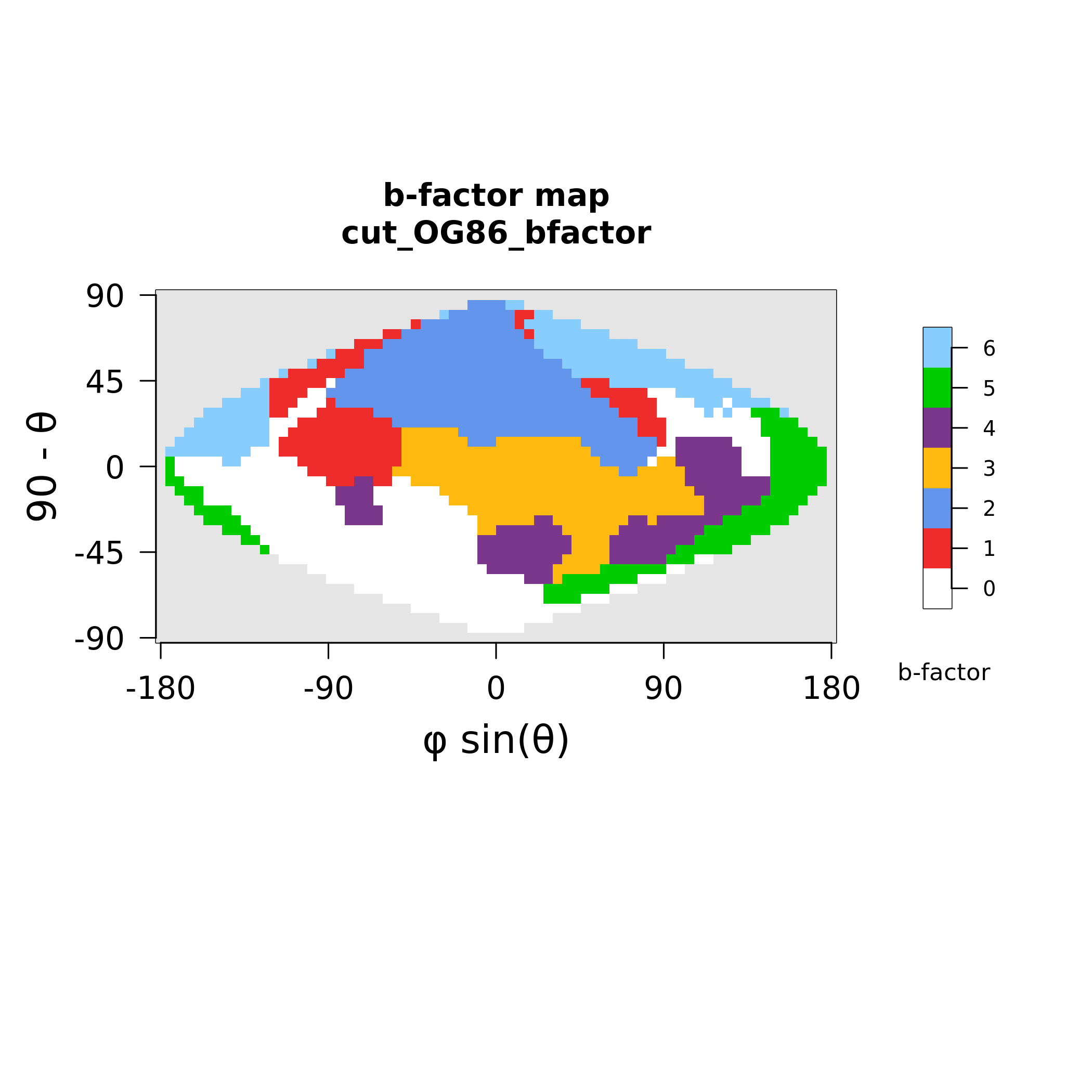

### MAX88_polymorphism.png

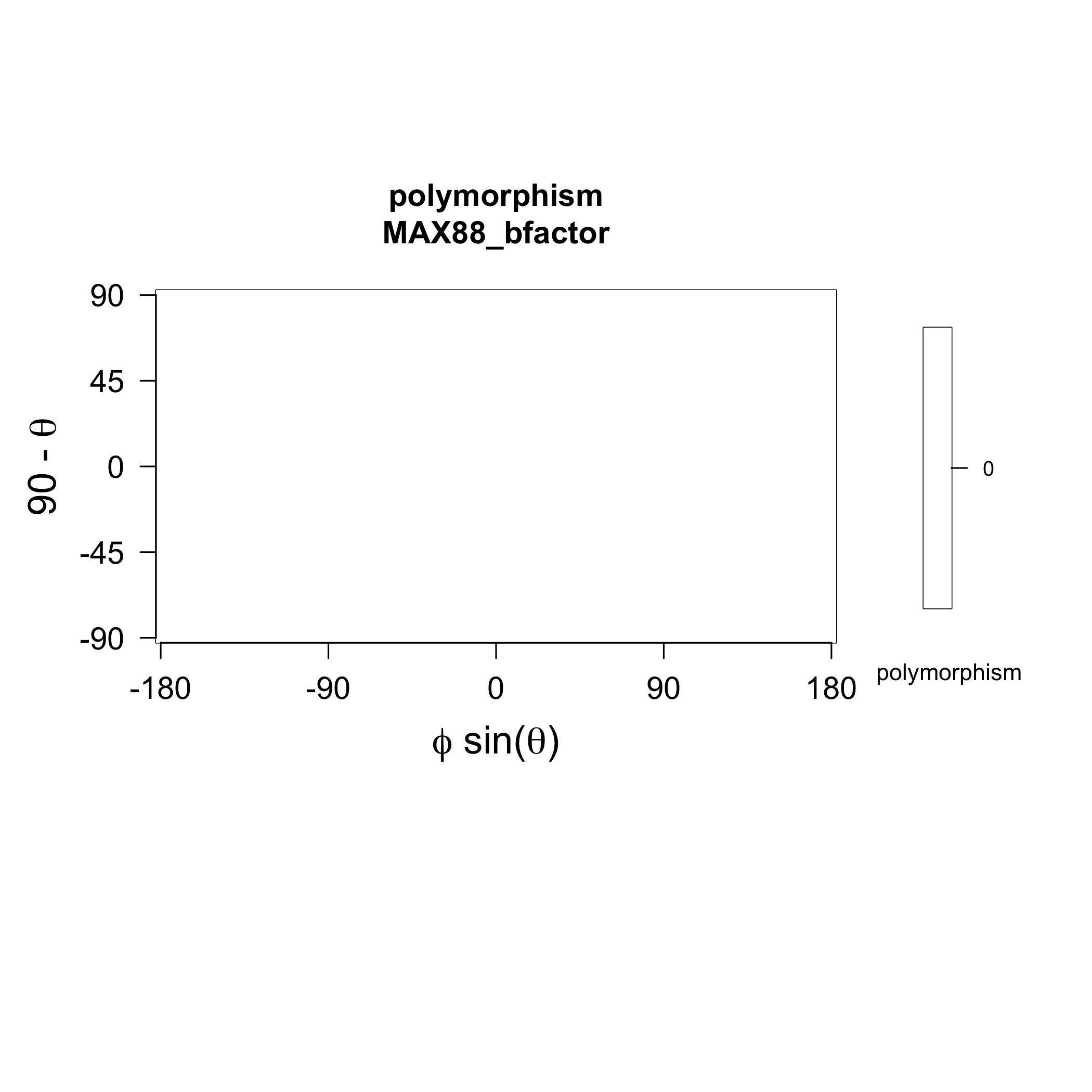

### MAX88_strands.png

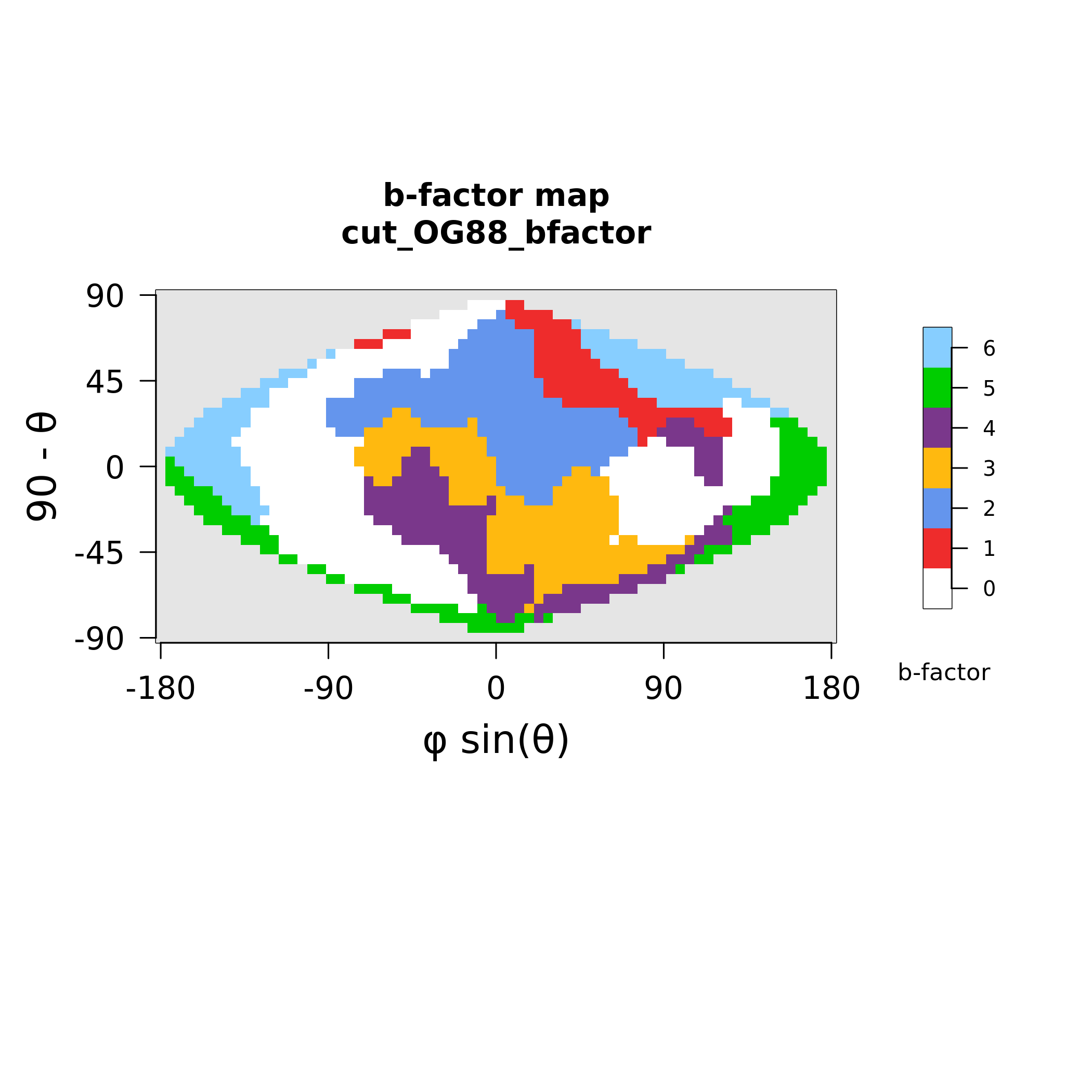

### MAX89_polymorphism.png

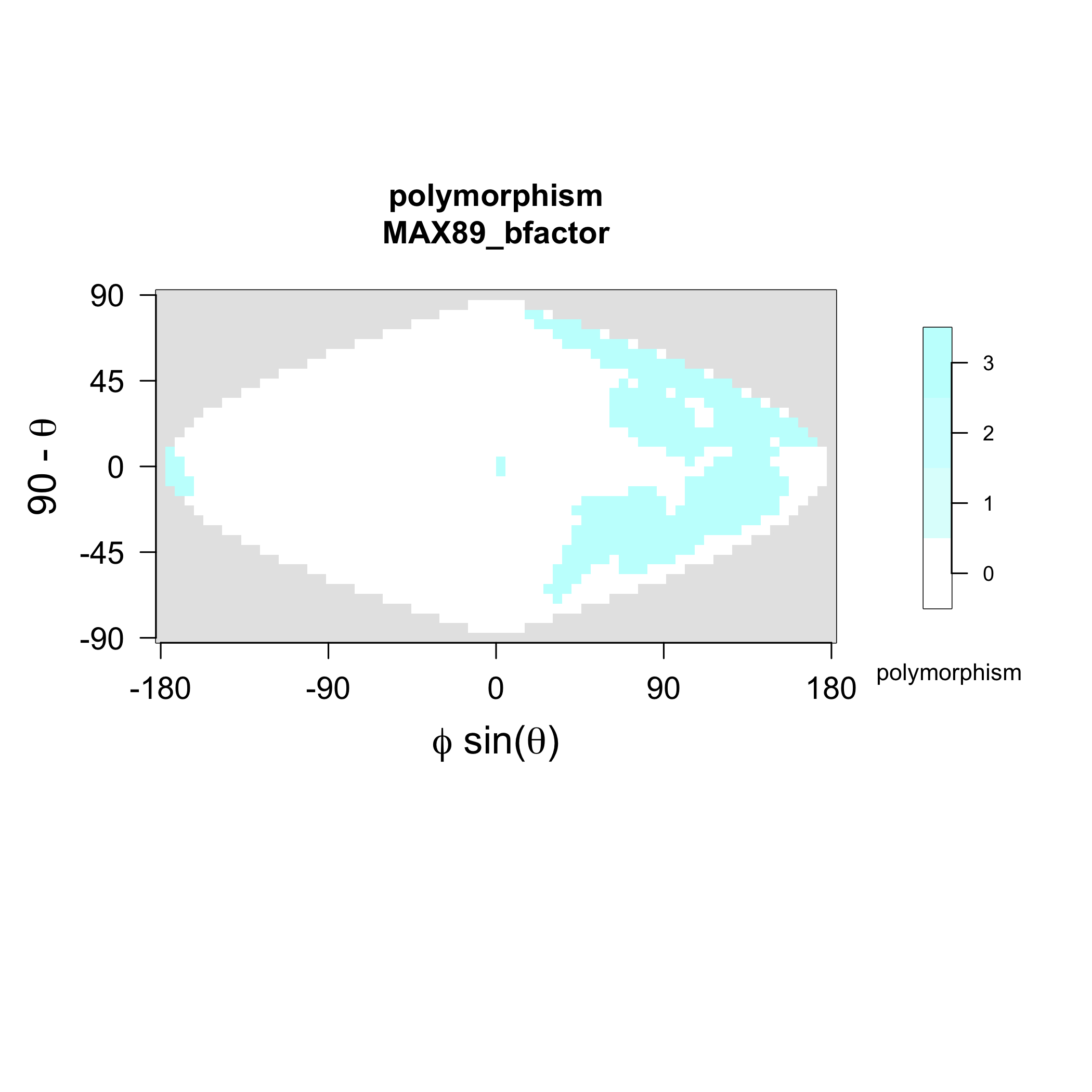

### MAX89_strands.png

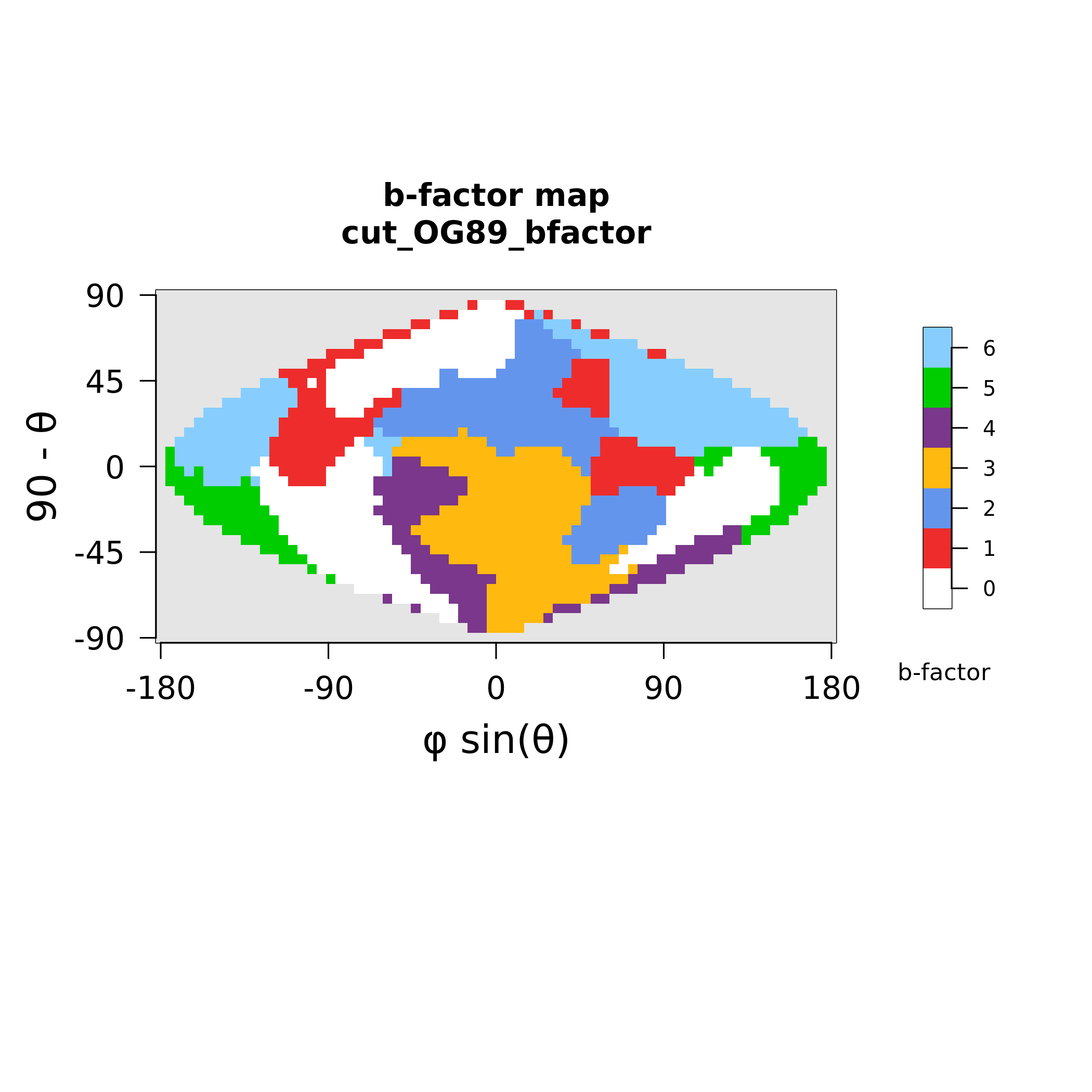

### MAX91_polymorphism.png

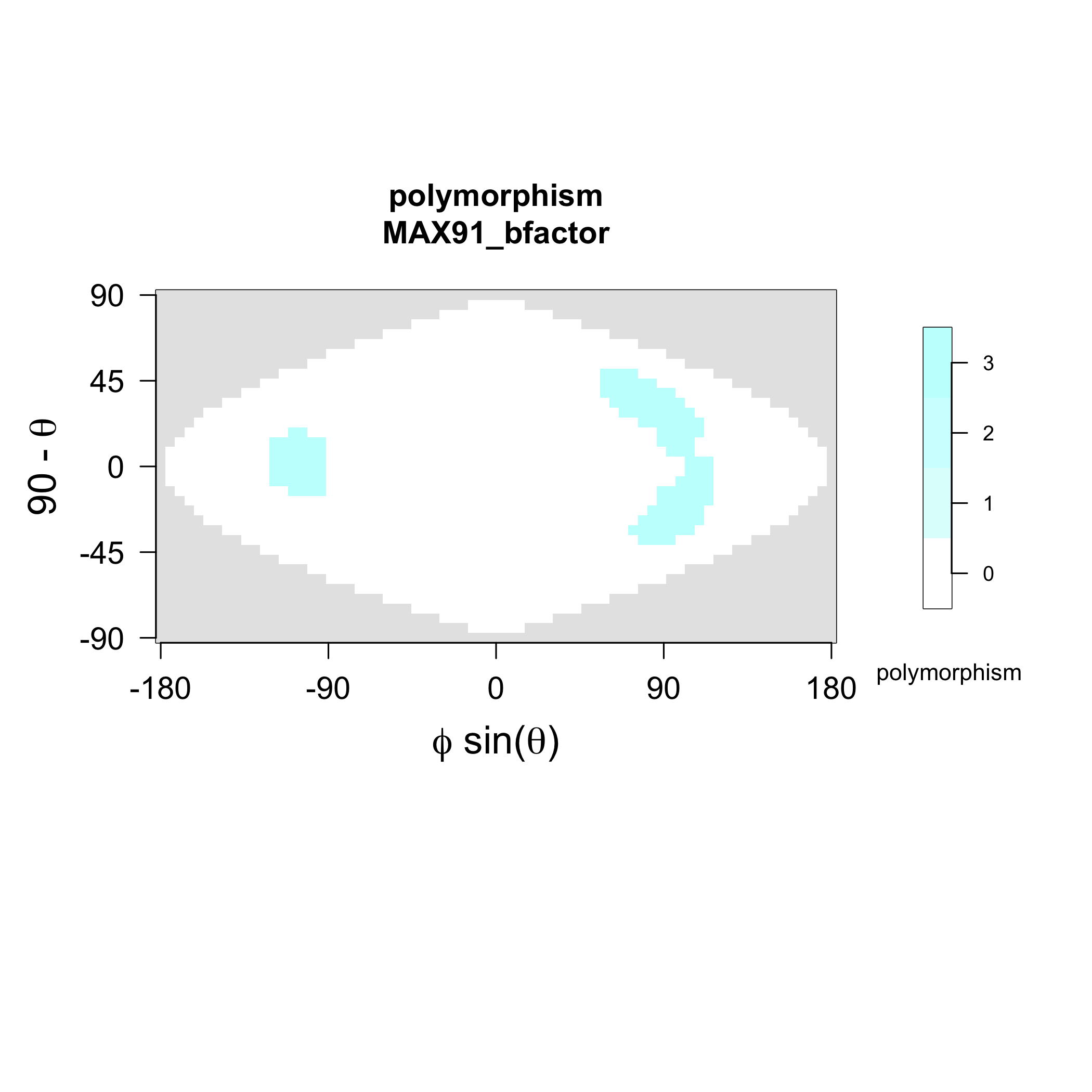

### MAX91_strands.png

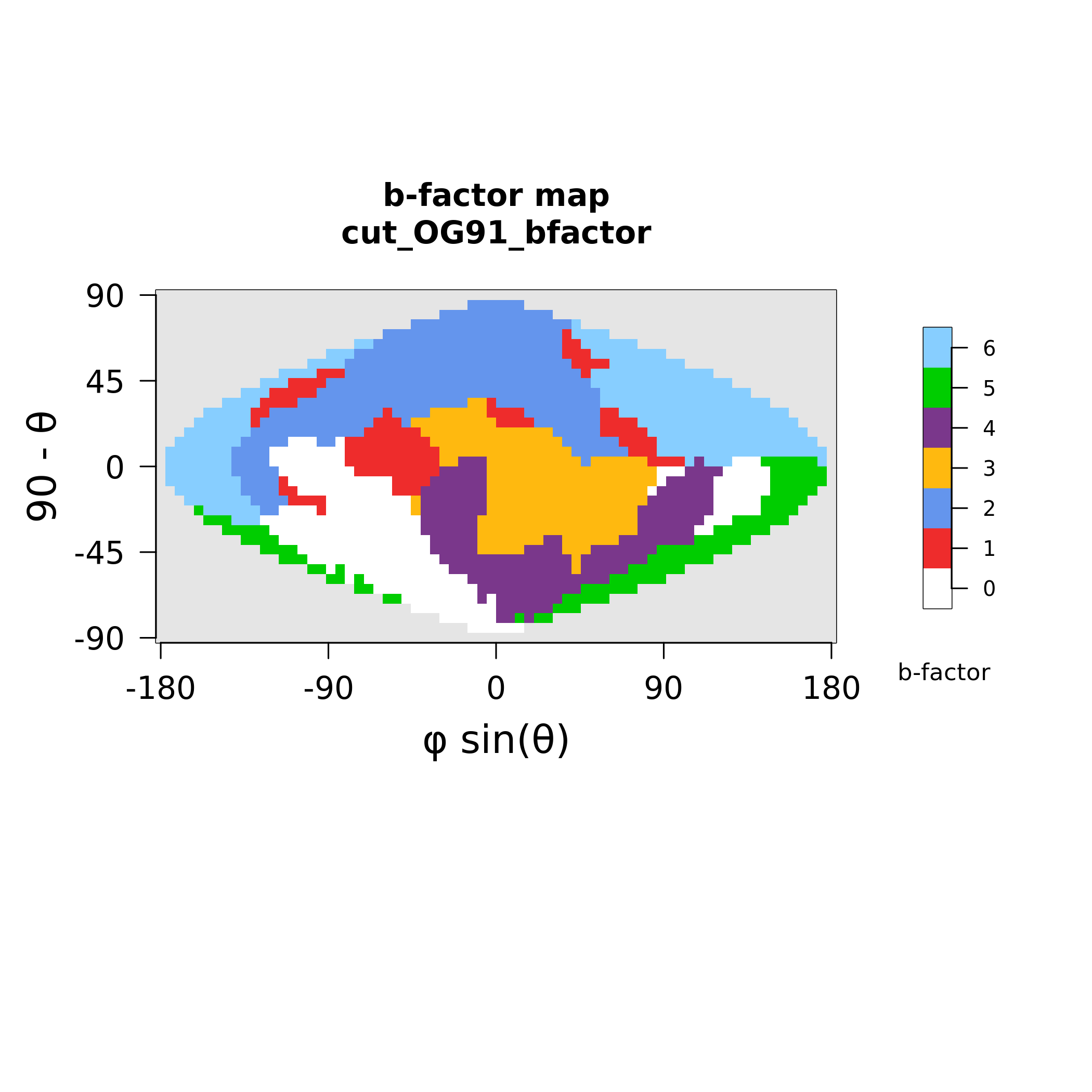

### MAX92_polymorphism.png

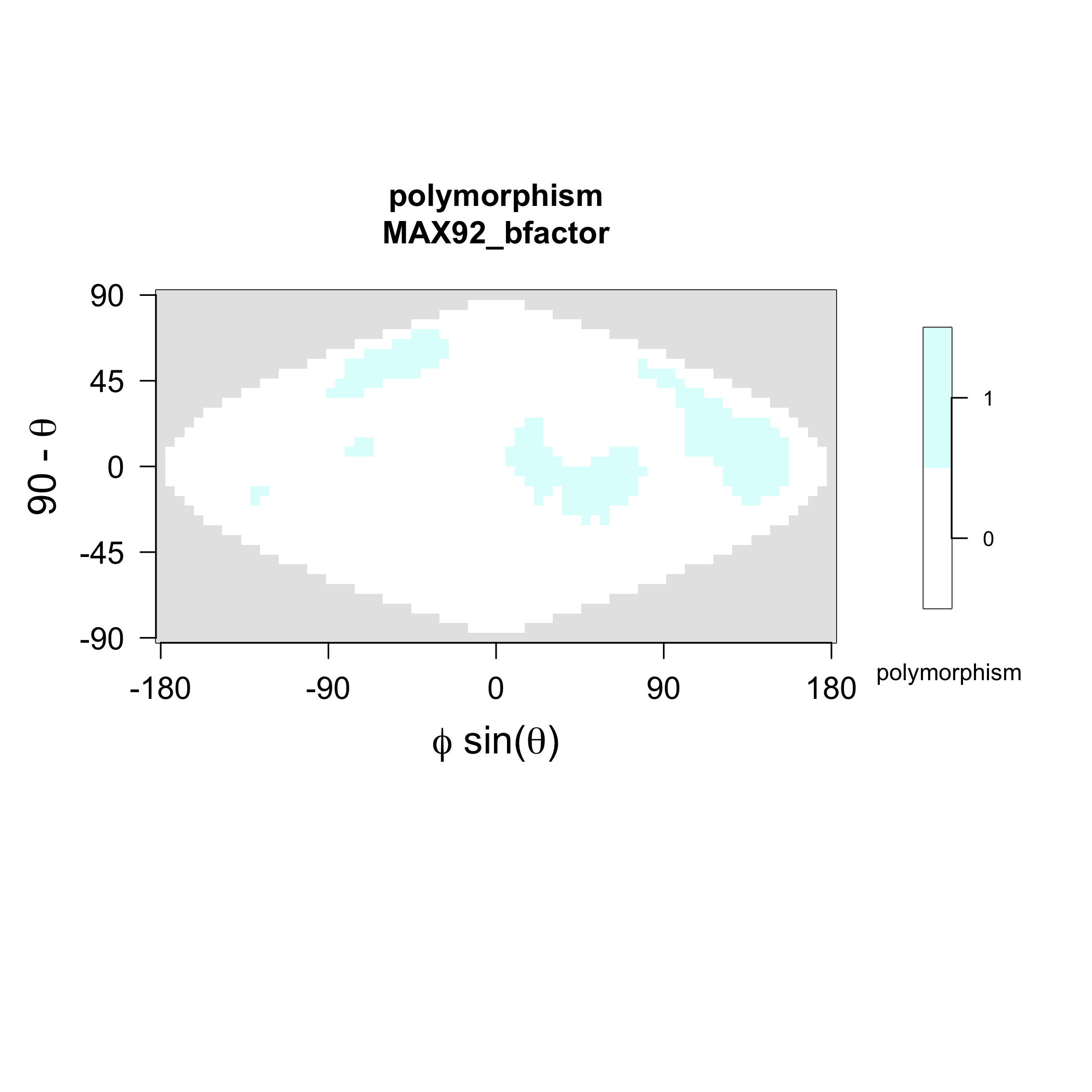

### MAX92_strands.png

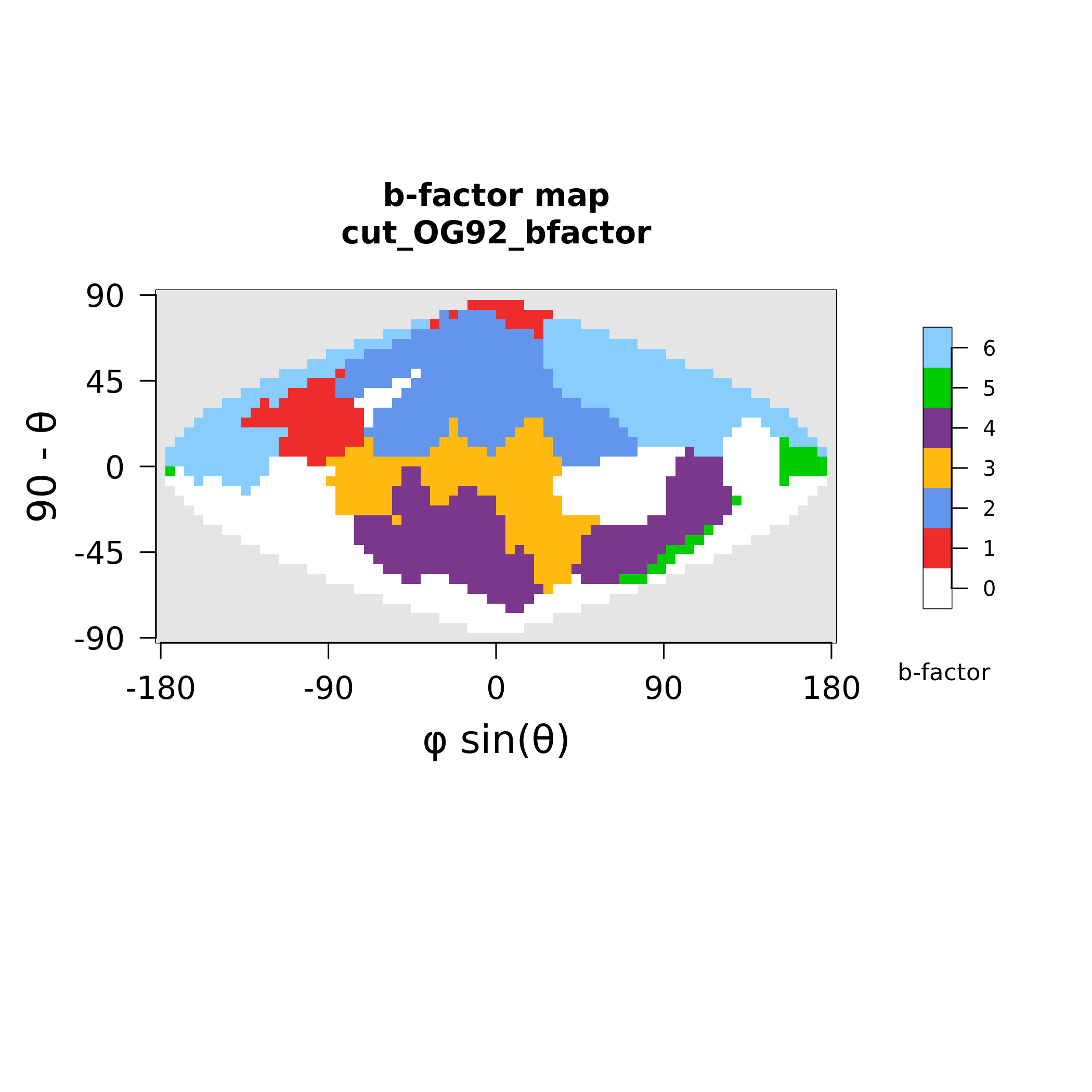

### MAX93_polymorphism.png

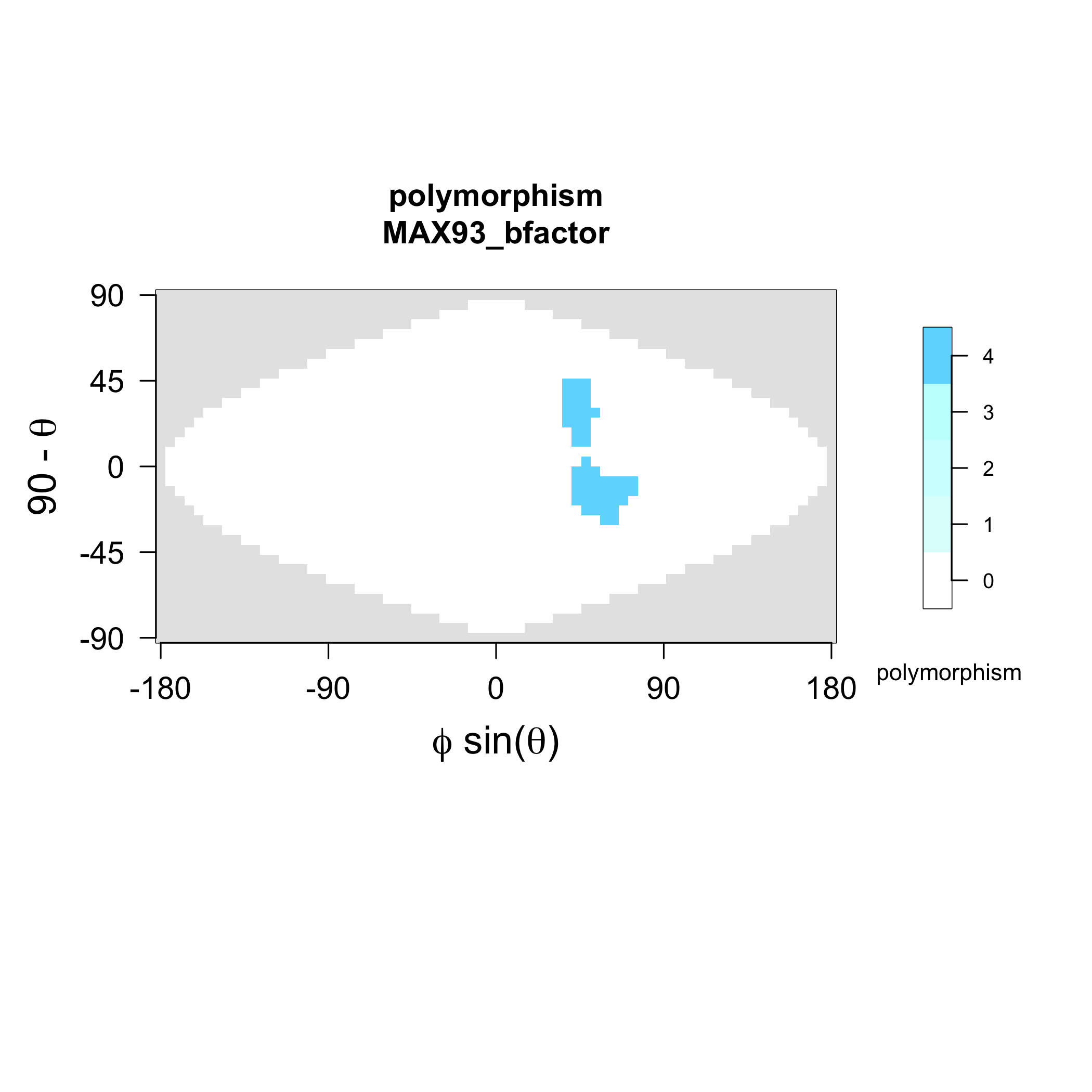

### MAX93_strands.png

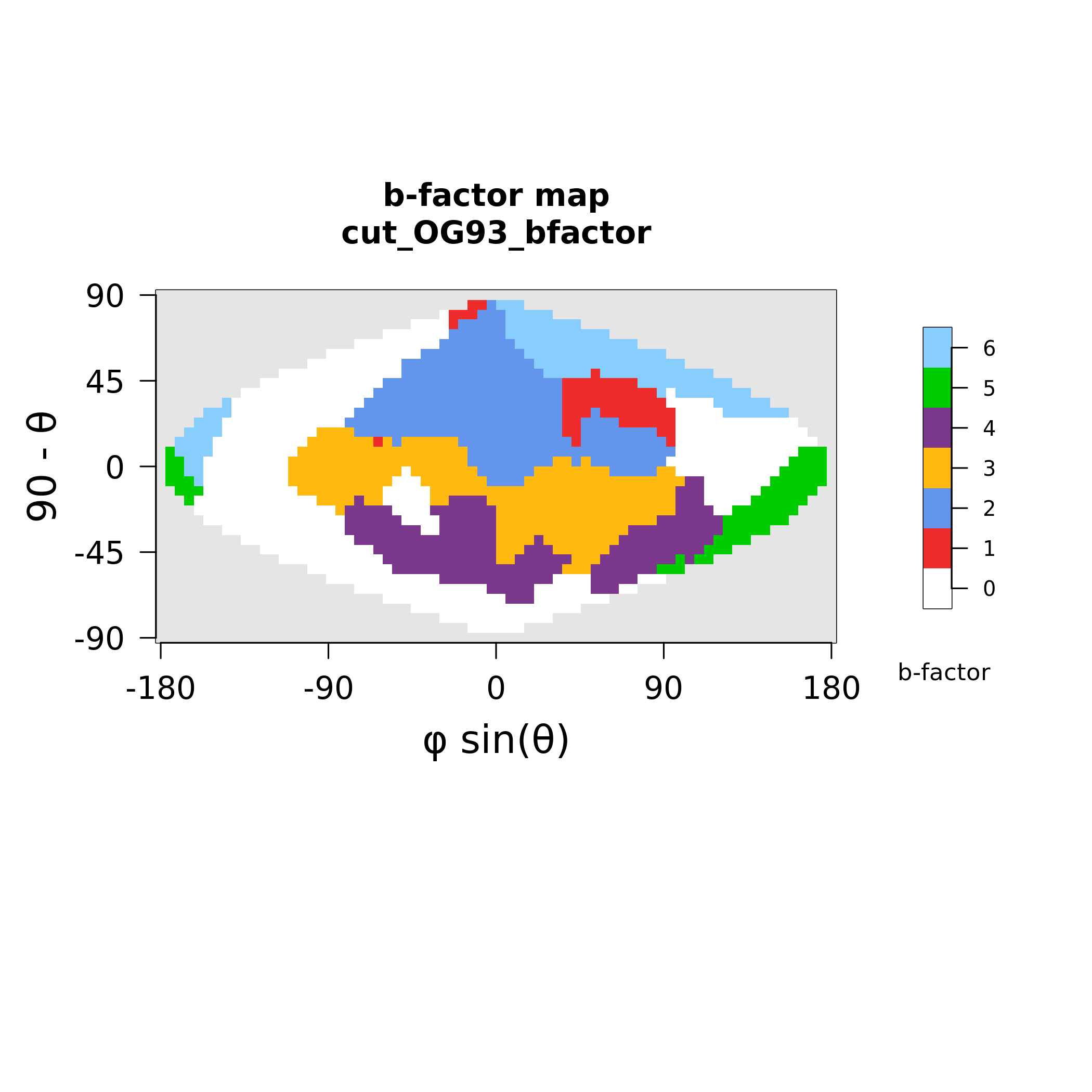
