## Supplementary figures and images for "The structural landscape and diversity of *Pyricularia oryzae* MAX effectors revisited"

### MAX01_electrostatics.png

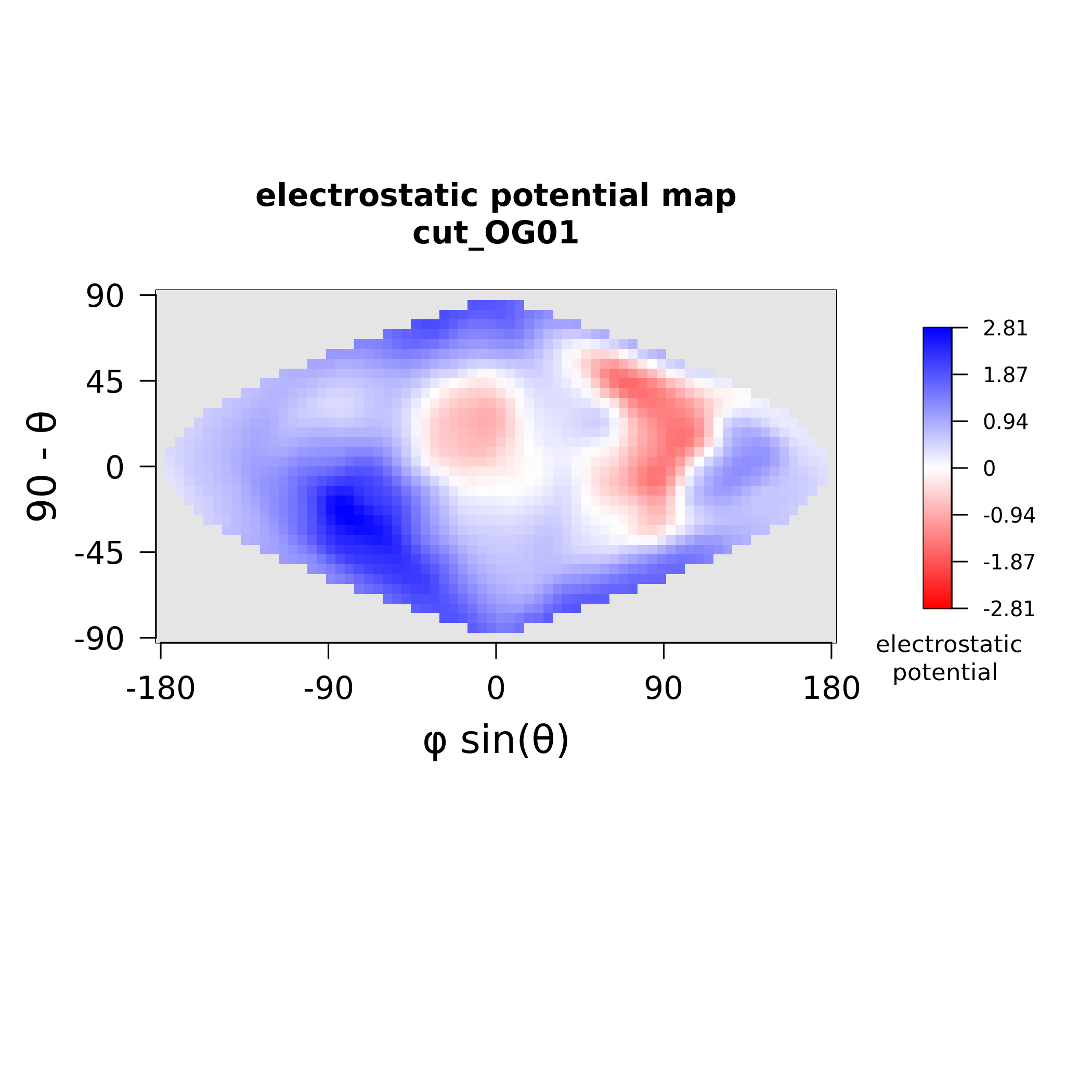

### MAX01_stickiness.png

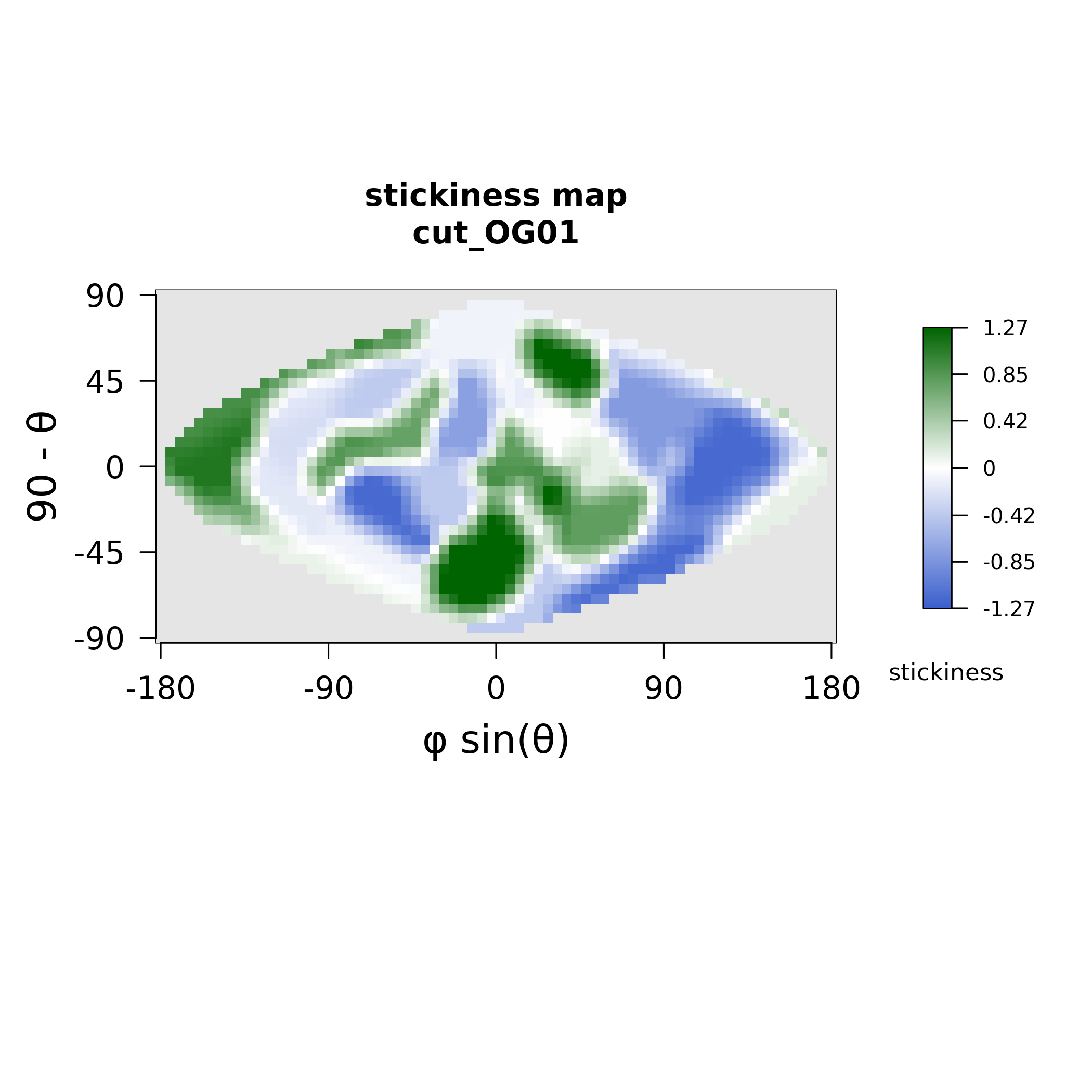

### MAX02_electrostatics.png

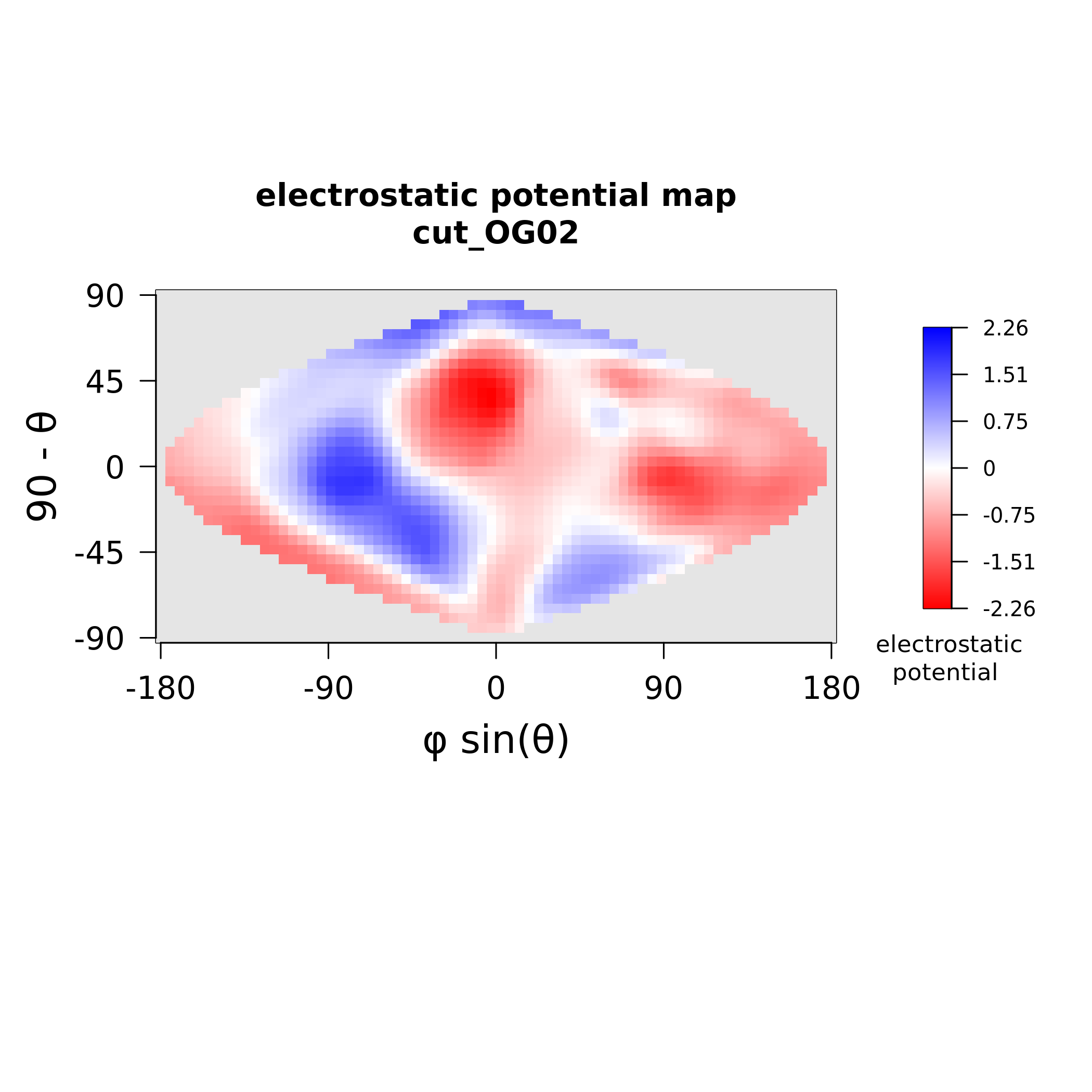

### MAX02_stickiness.png

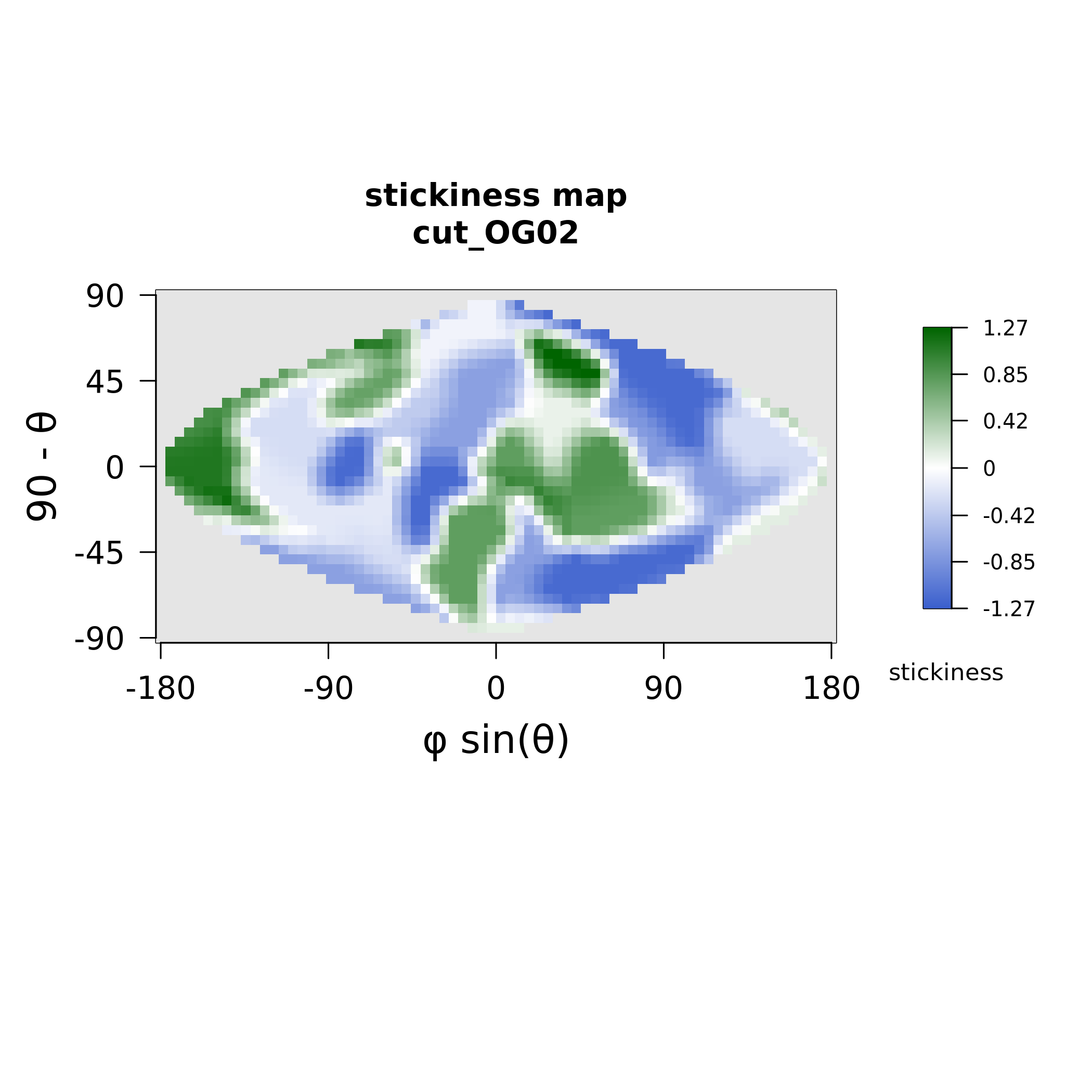

### MAX03_electrostatics.png

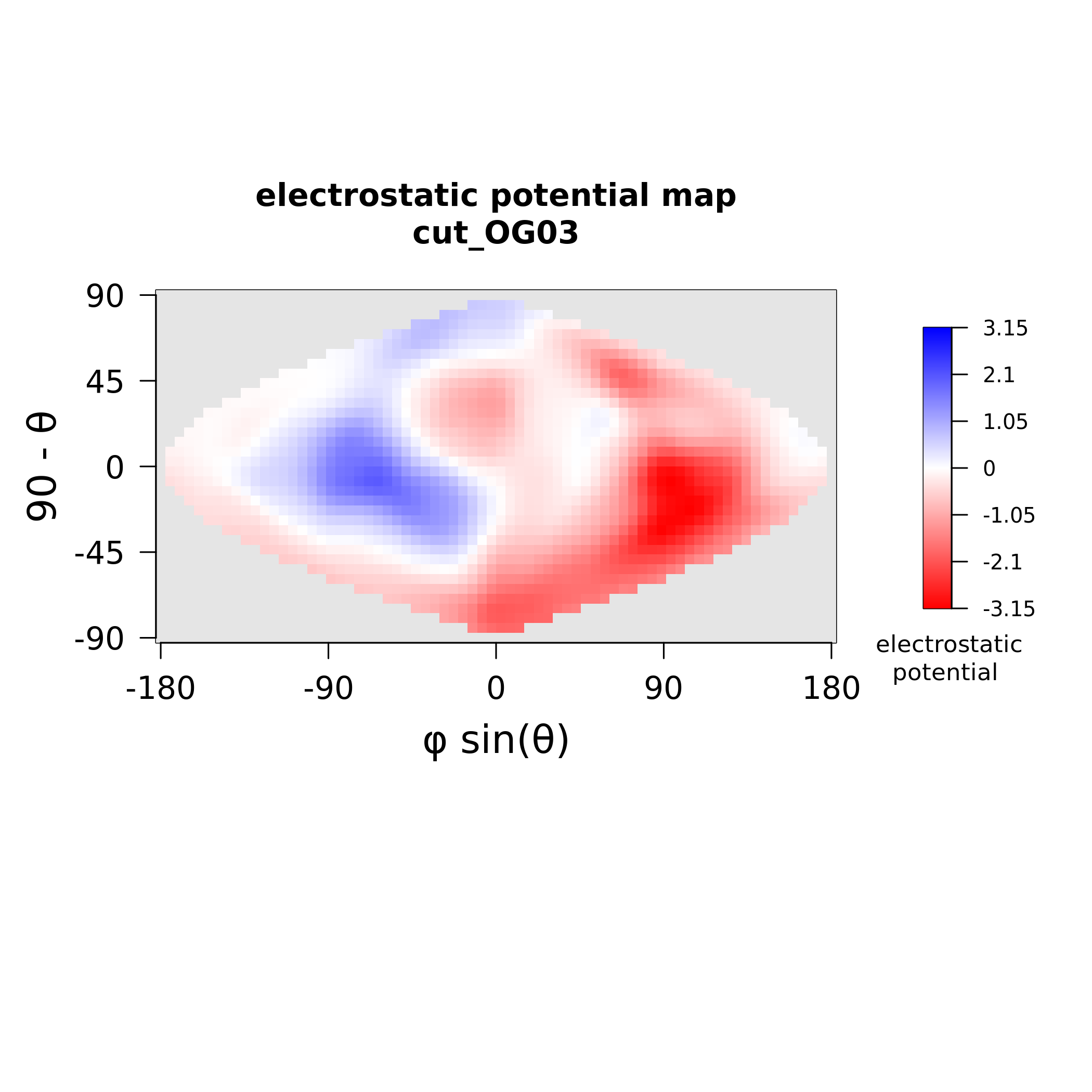

### MAX03_stickiness.png

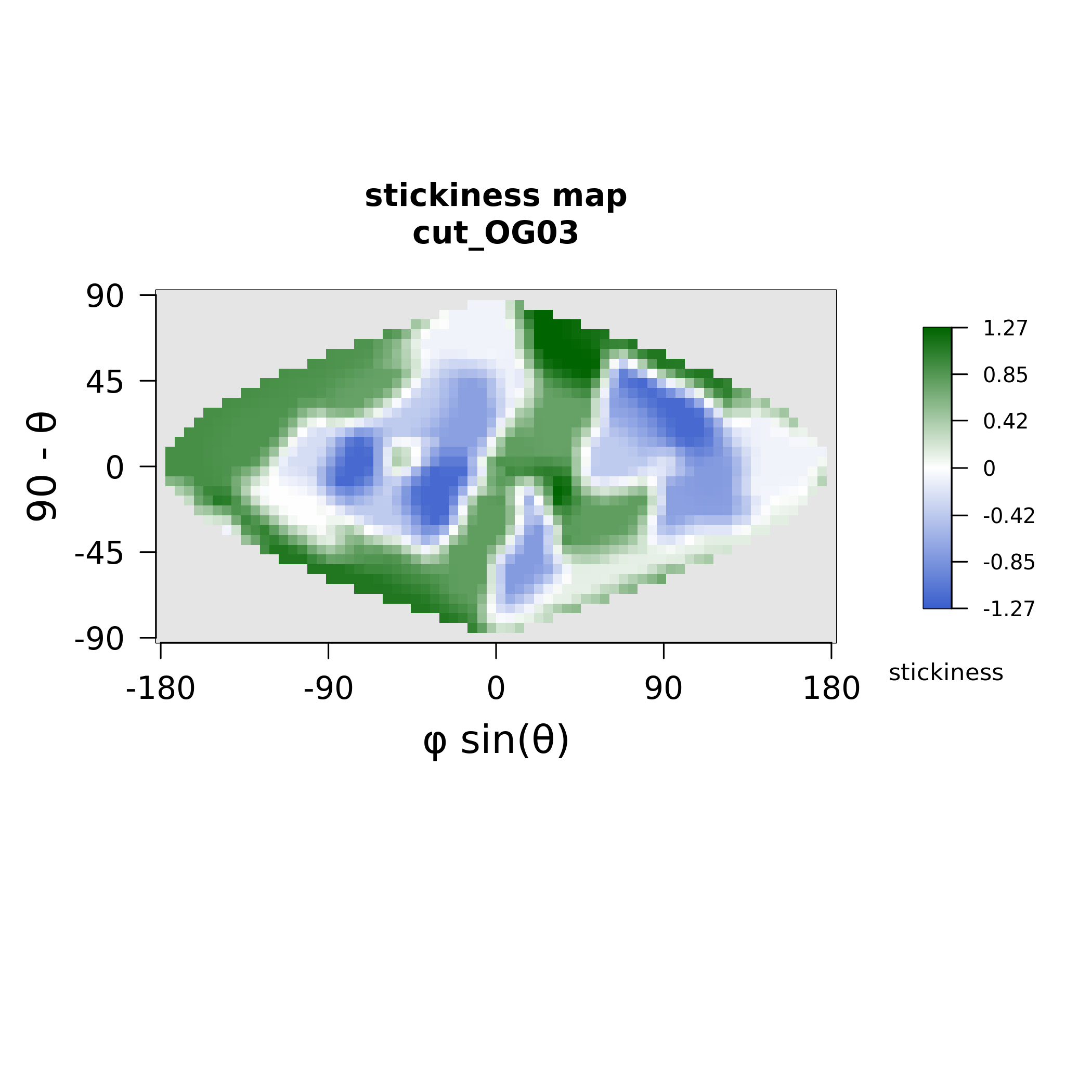

### MAX05_electrostatics.png

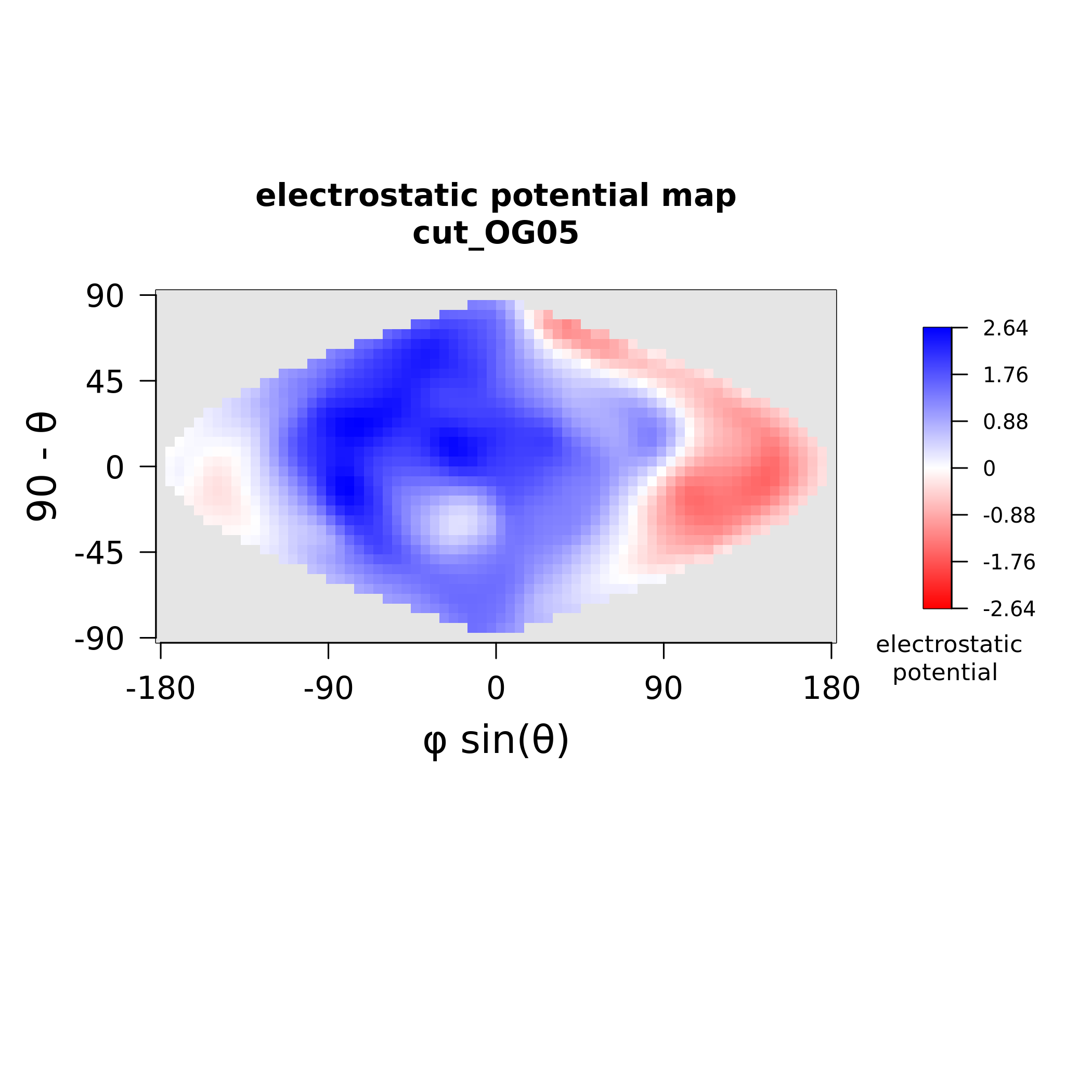

### MAX05_stickiness.png

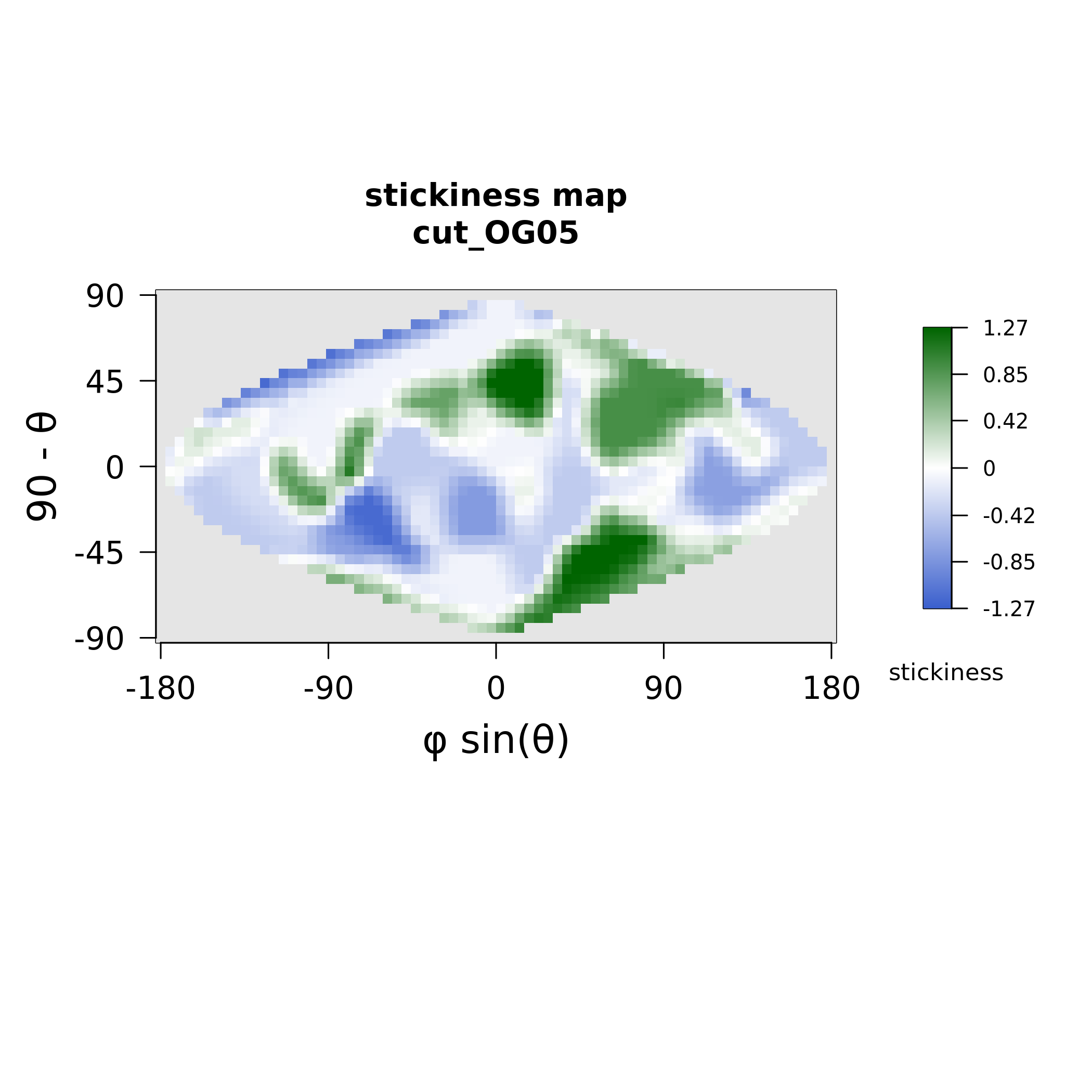

### MAX06_electrostatics.png

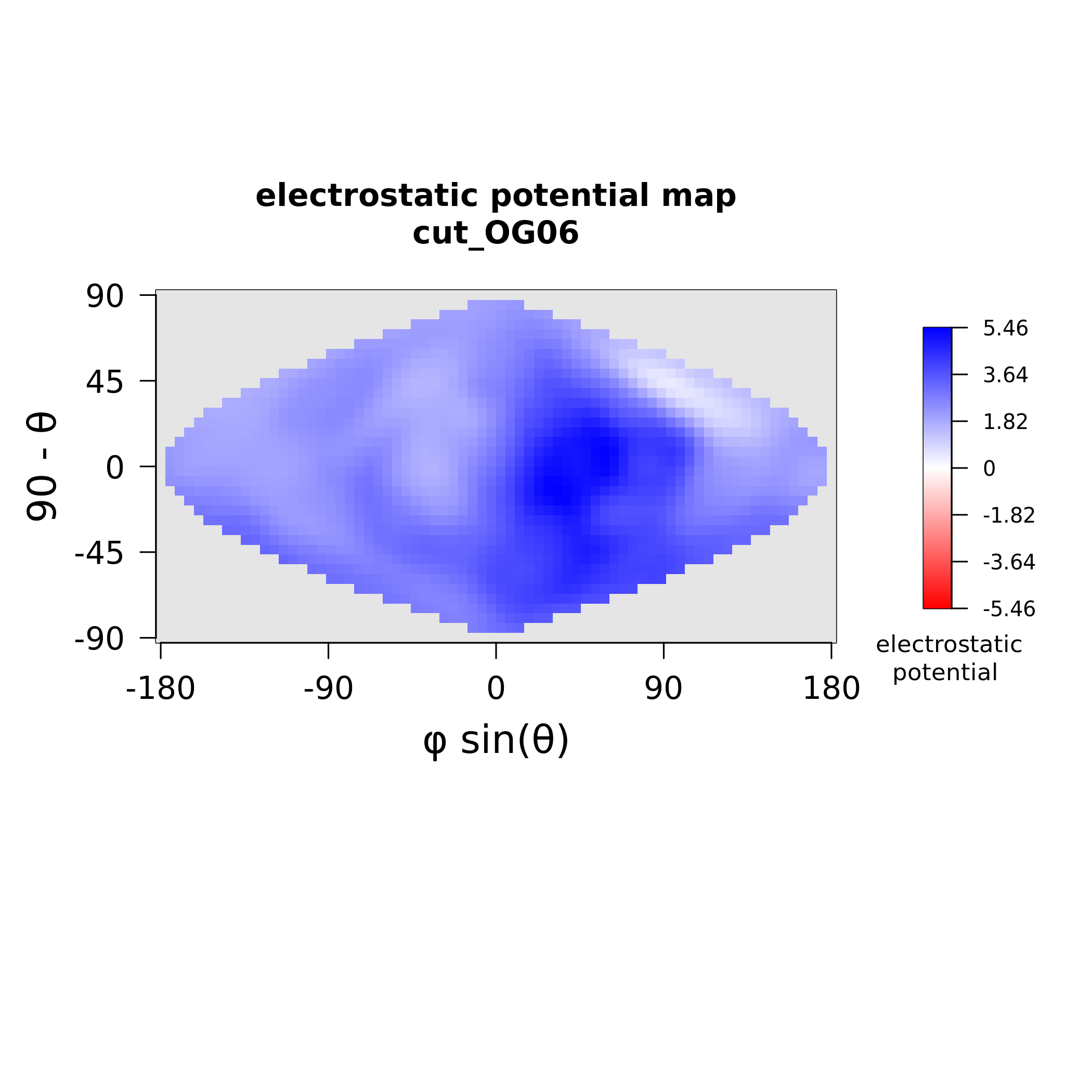

### MAX06_stickiness.png

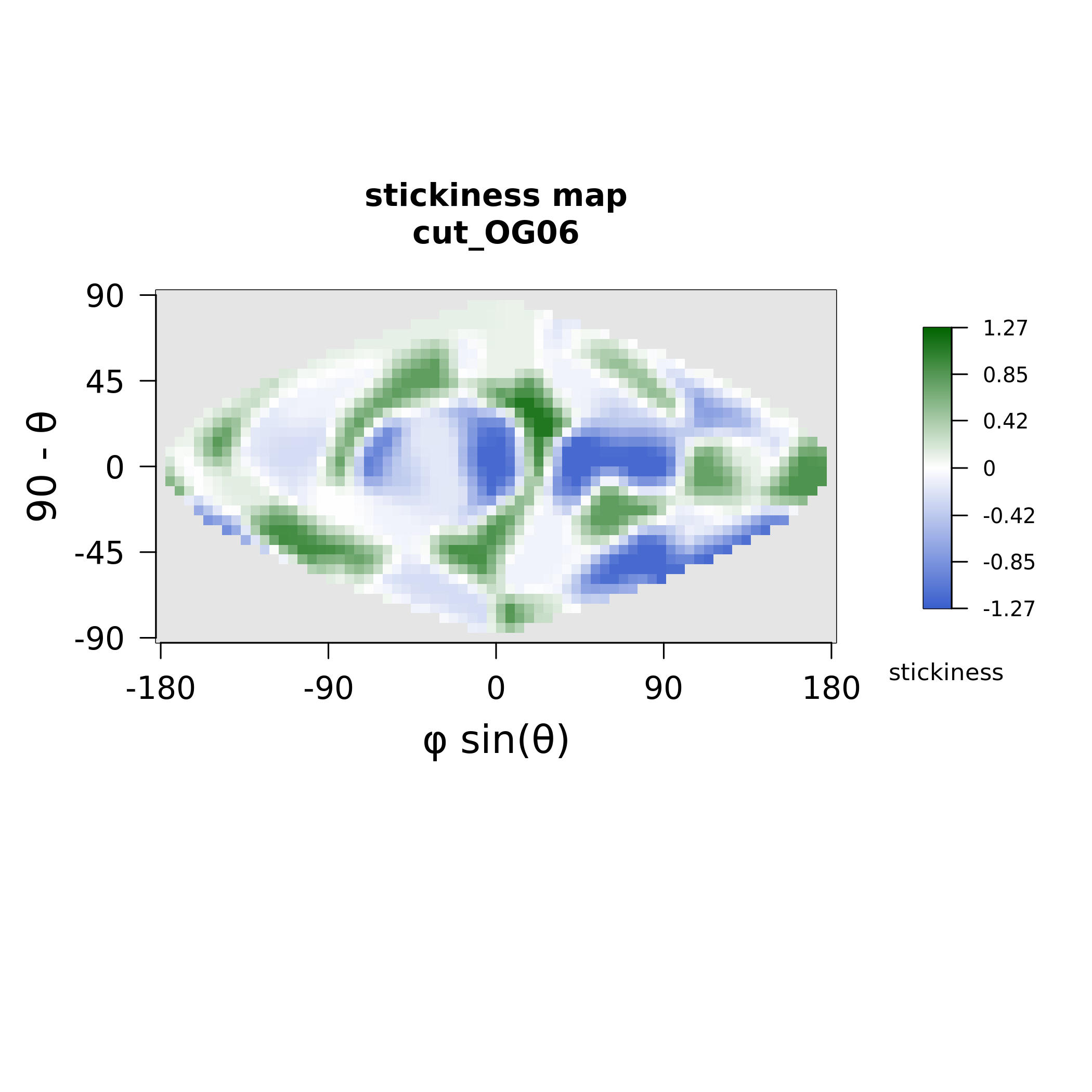

### MAX07_electrostatics.png

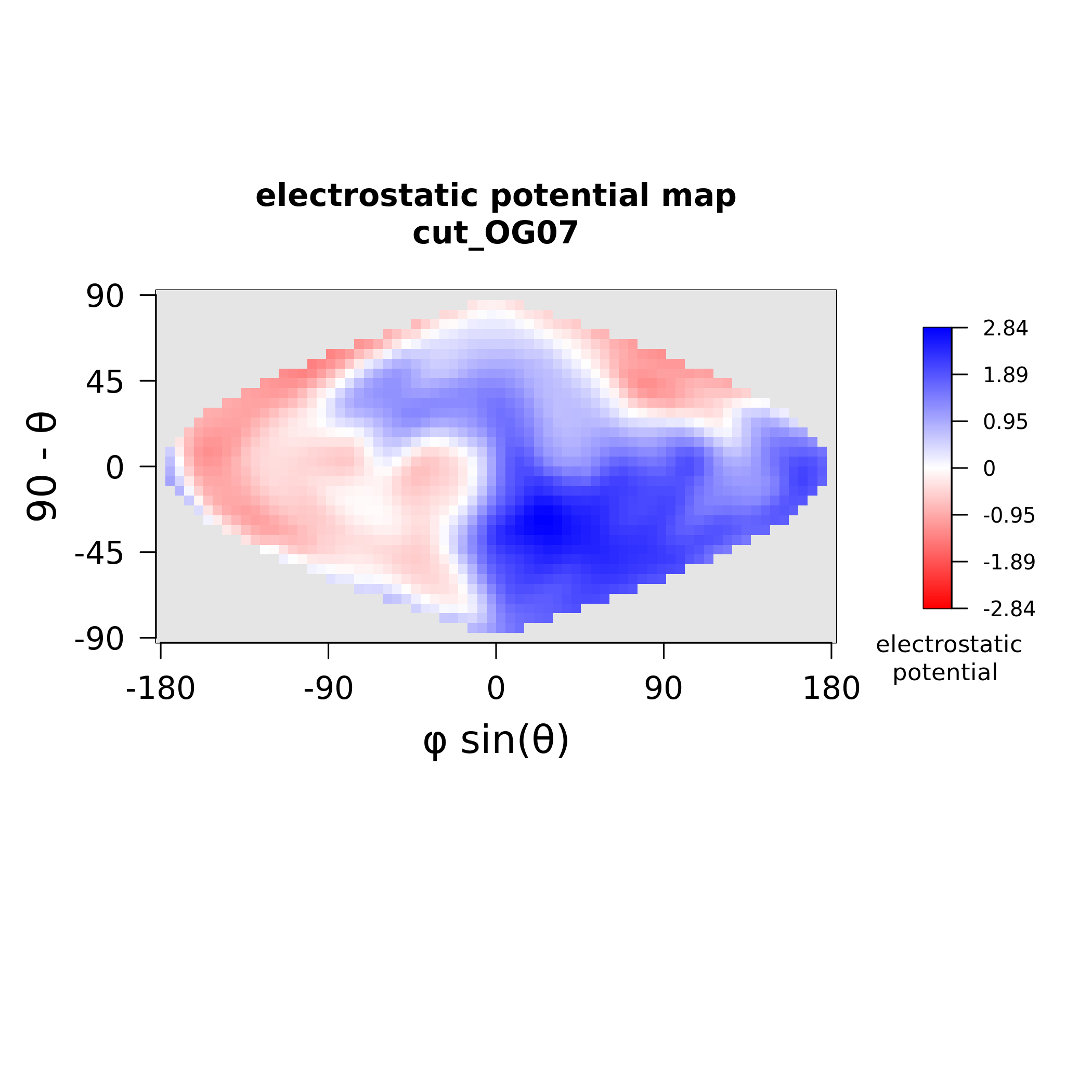

### MAX07_stickiness.png

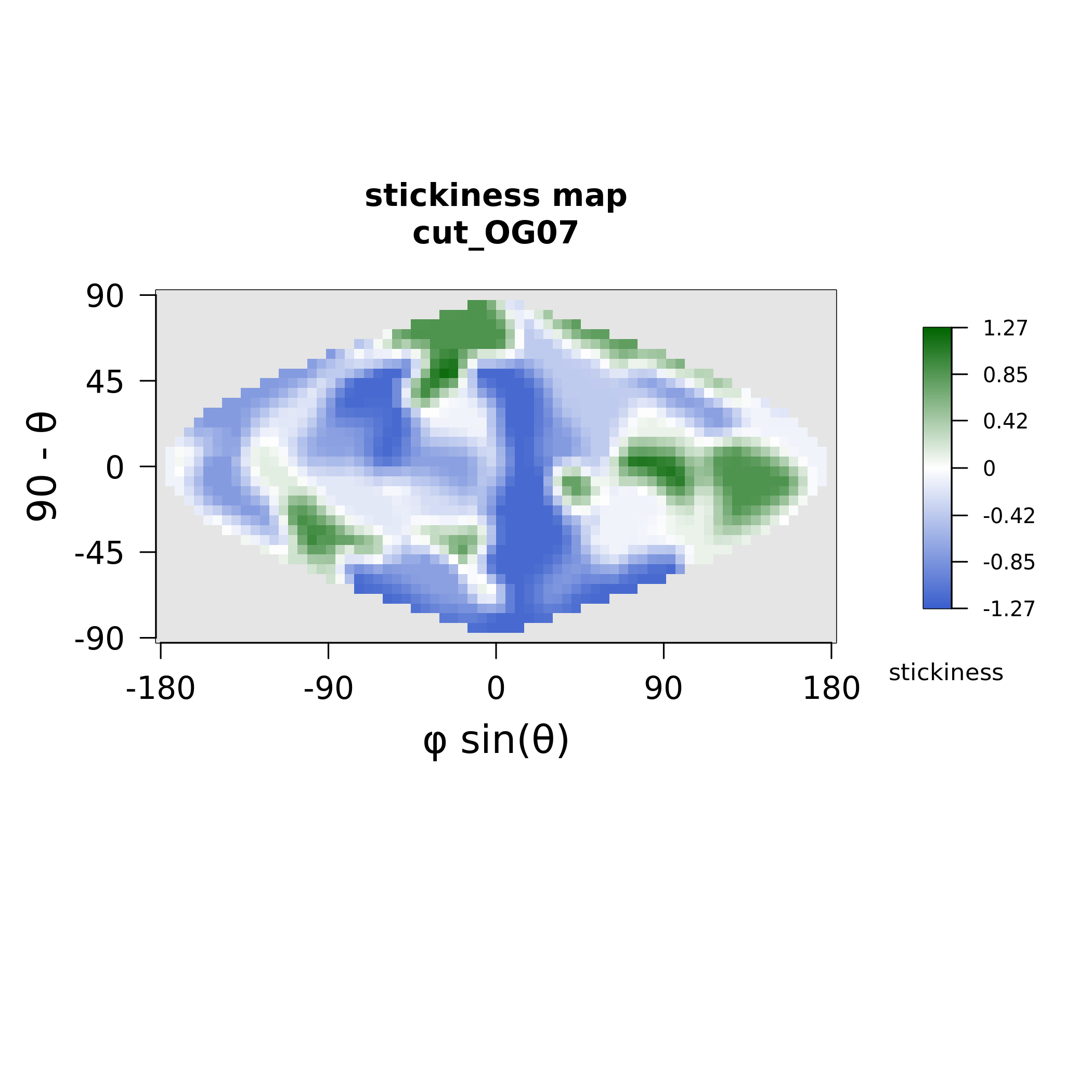

### MAX10_electrostatics.png

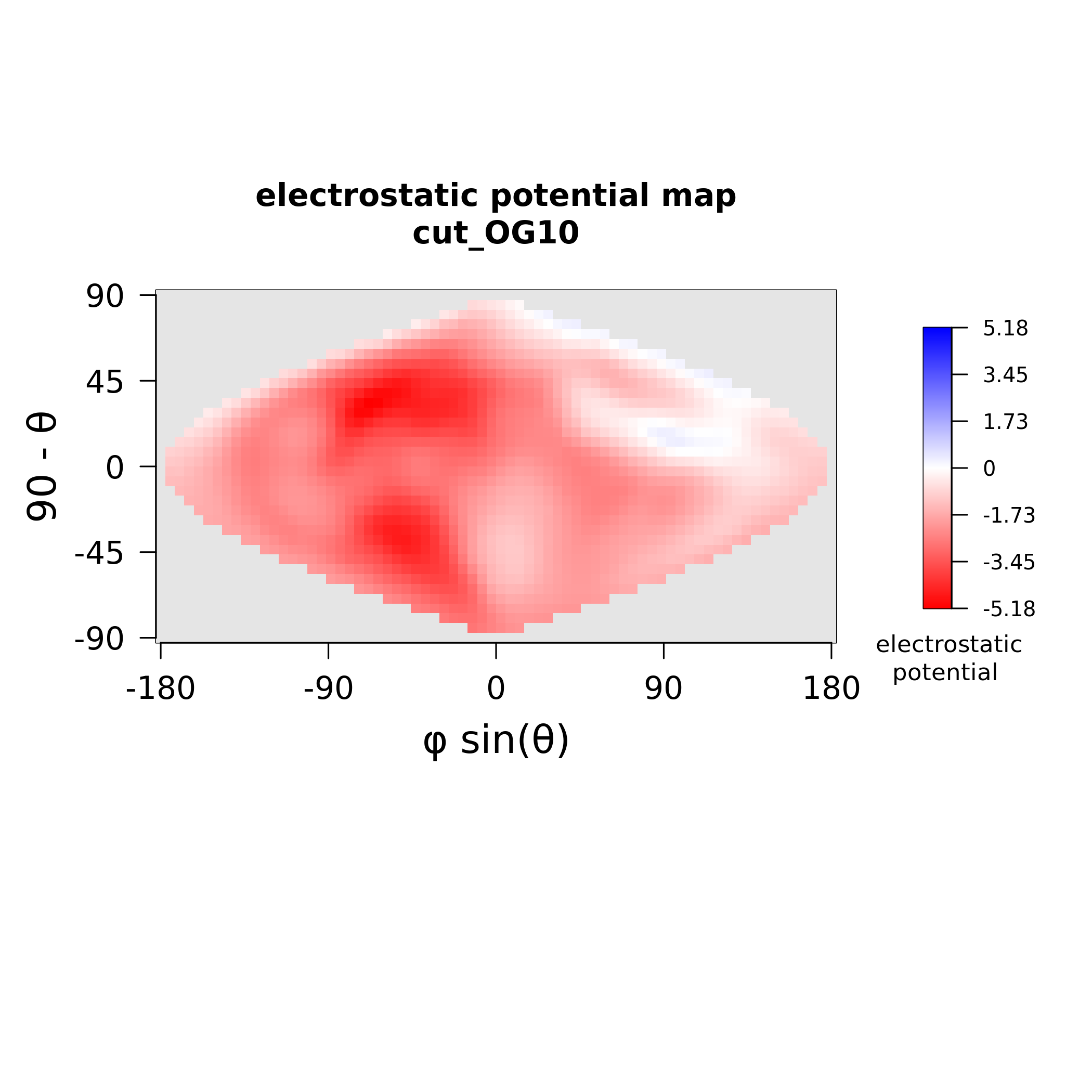

### MAX10_stickiness.png

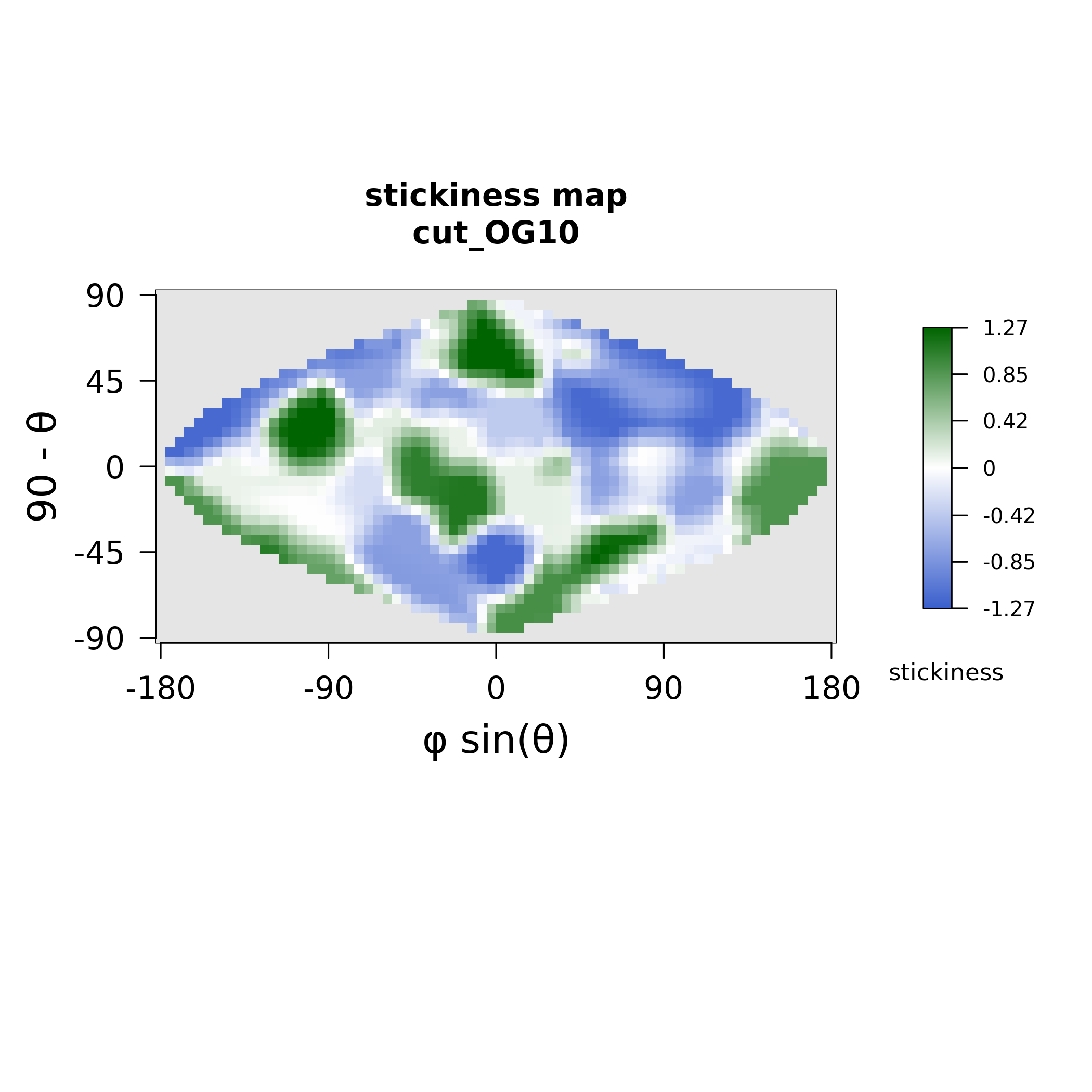

### MAX11_electrostatics.png

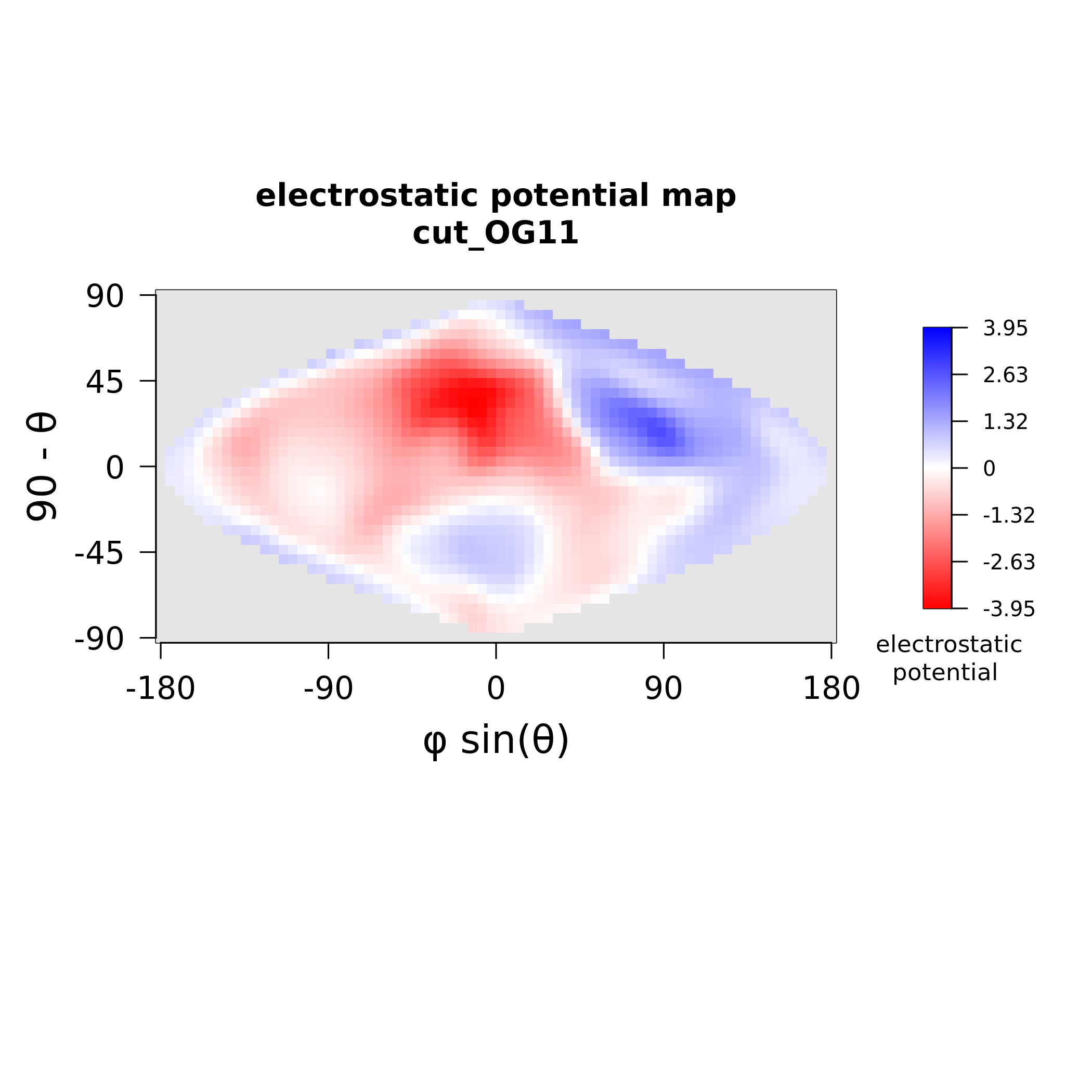

### MAX11_stickiness.png

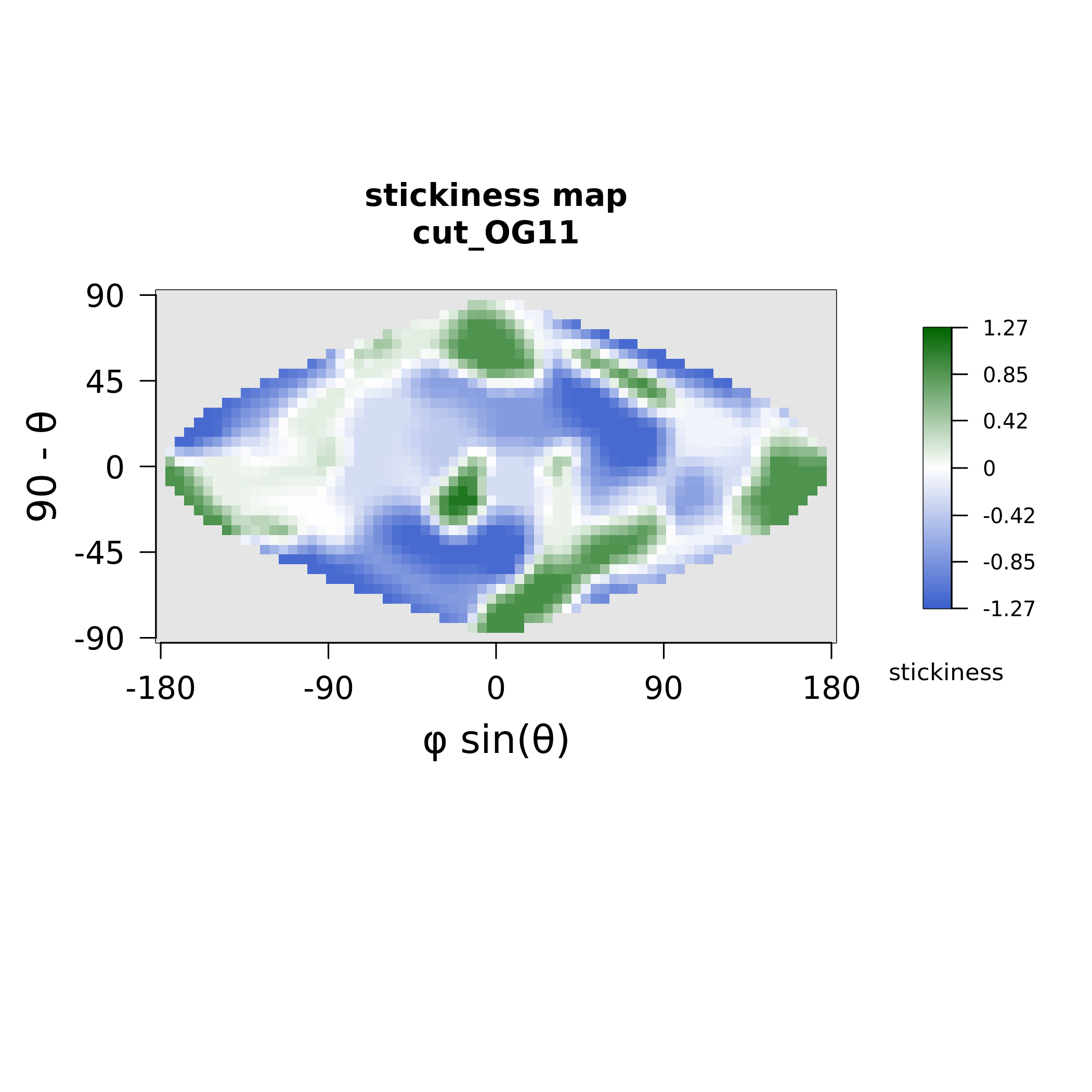

### MAX15_electrostatics.png

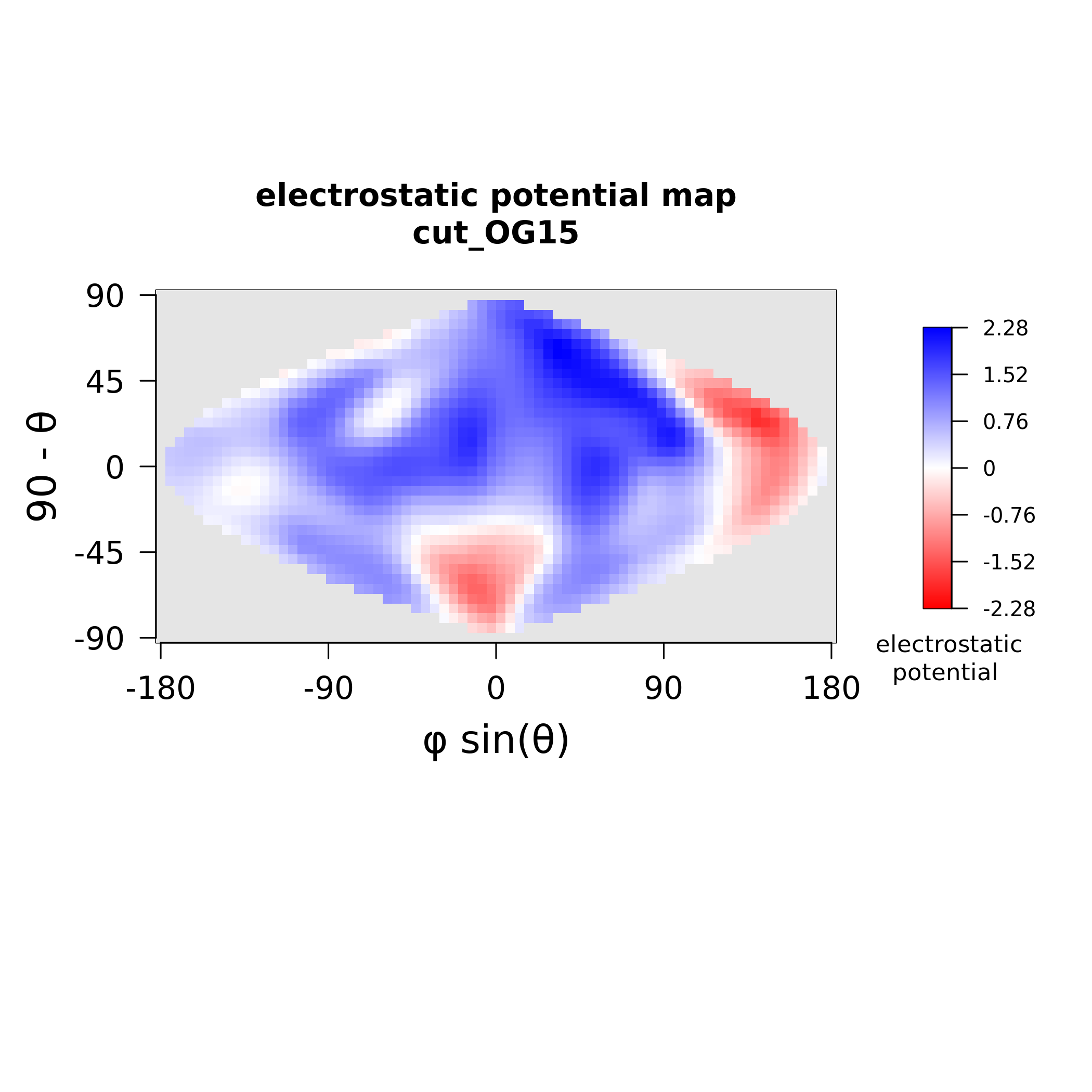

### MAX15_stickiness.png

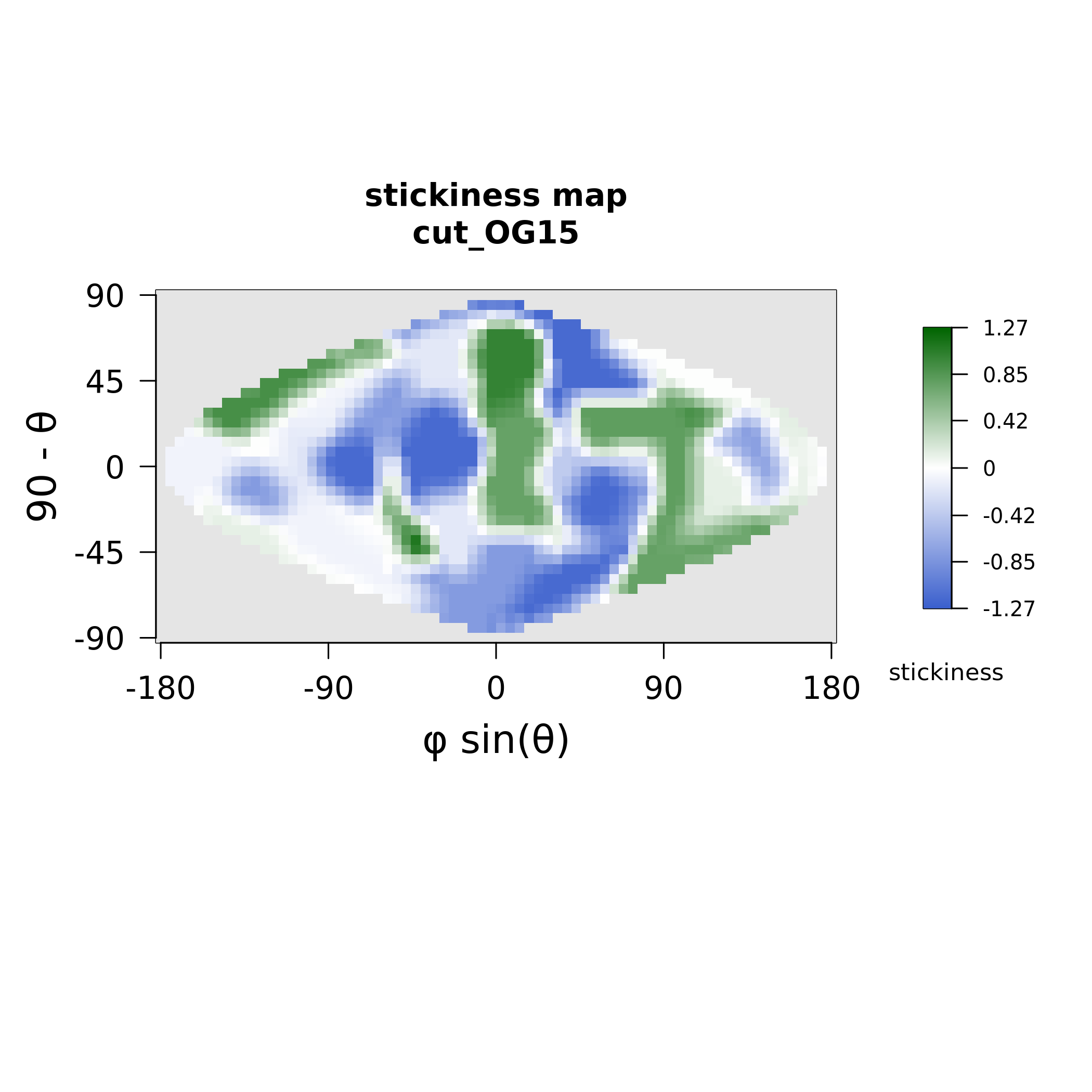

### MAX17_electrostatics.png

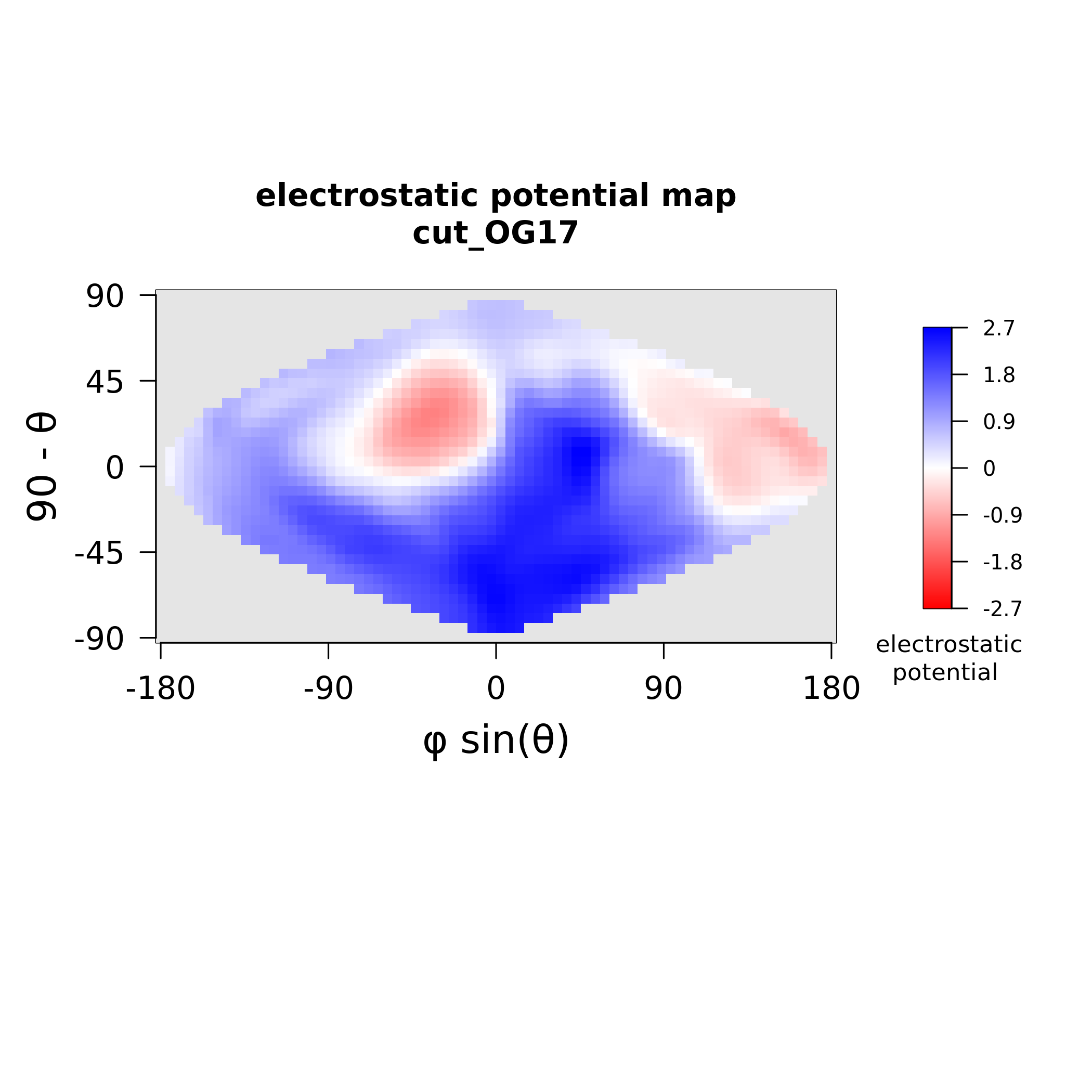

### MAX17_stickiness.png

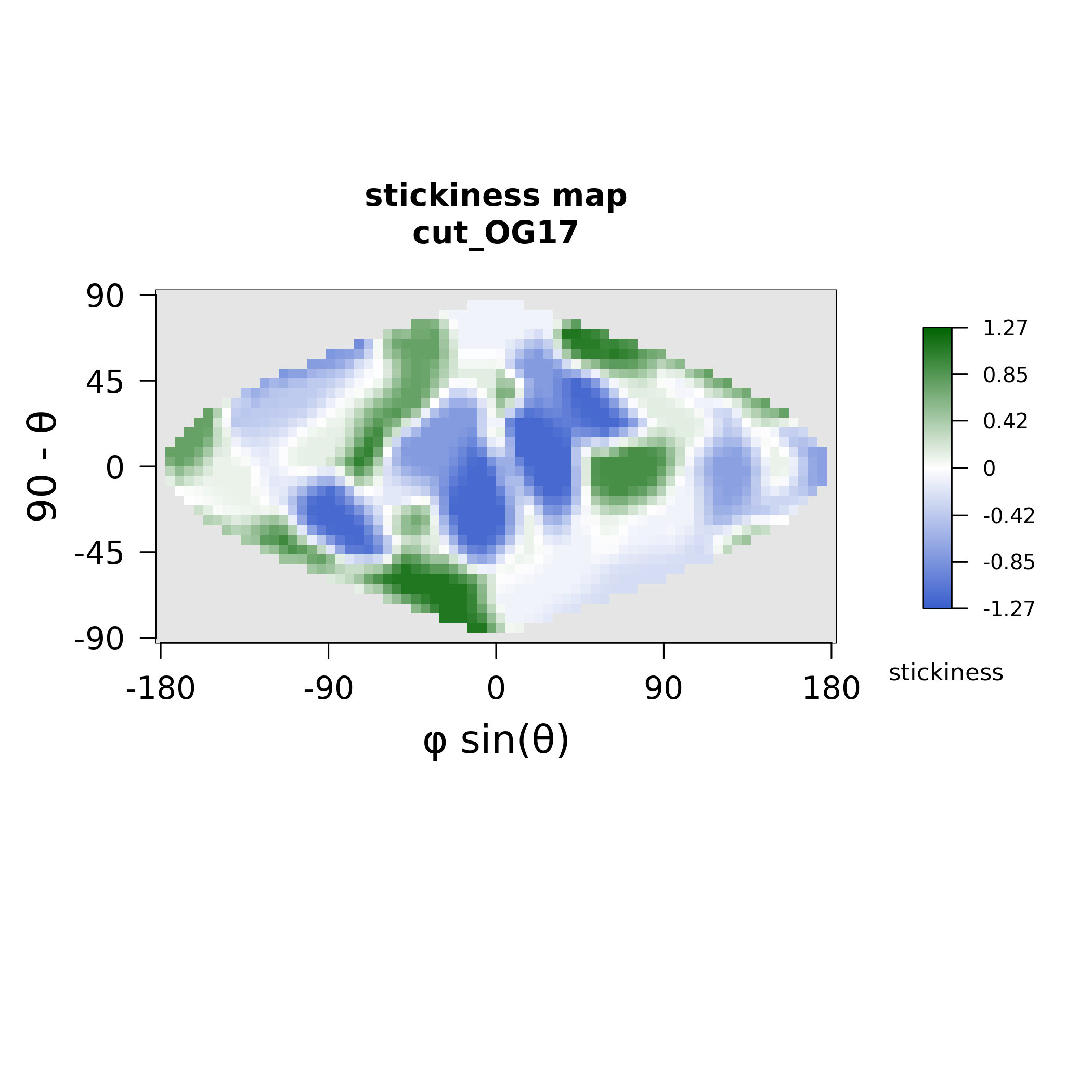

### MAX18_electrostatics.png

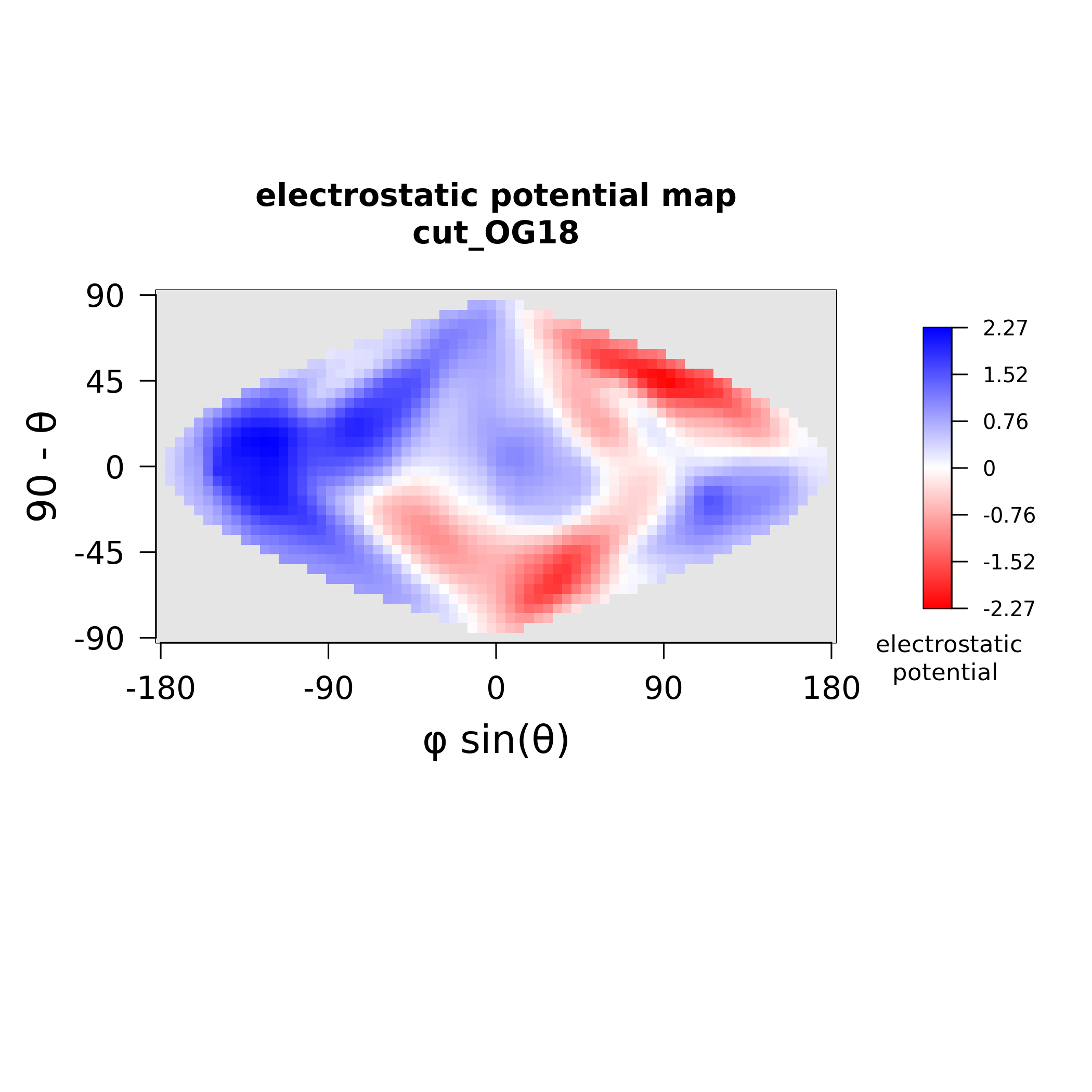

### MAX18_stickiness.png

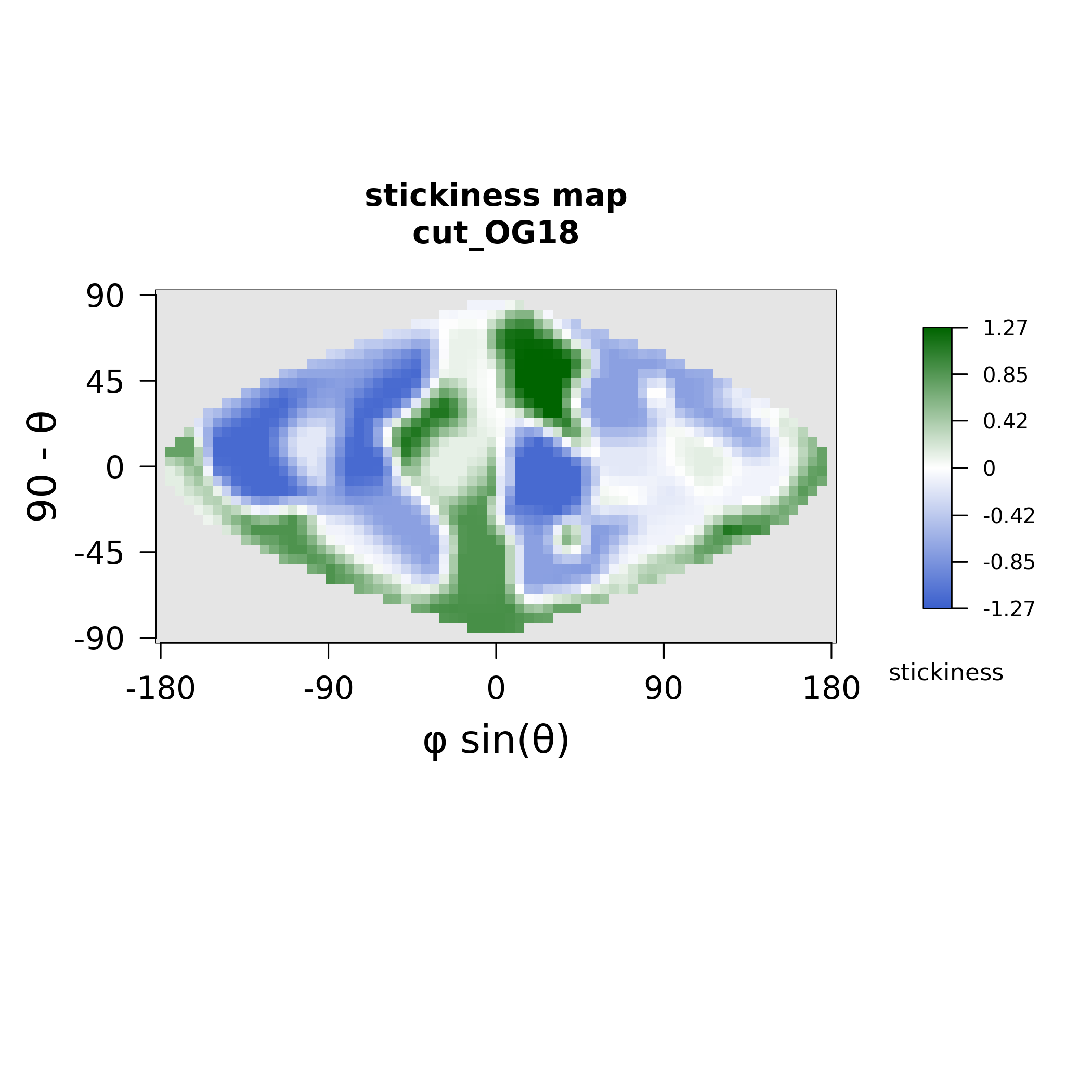

### MAX19_electrostatics.png

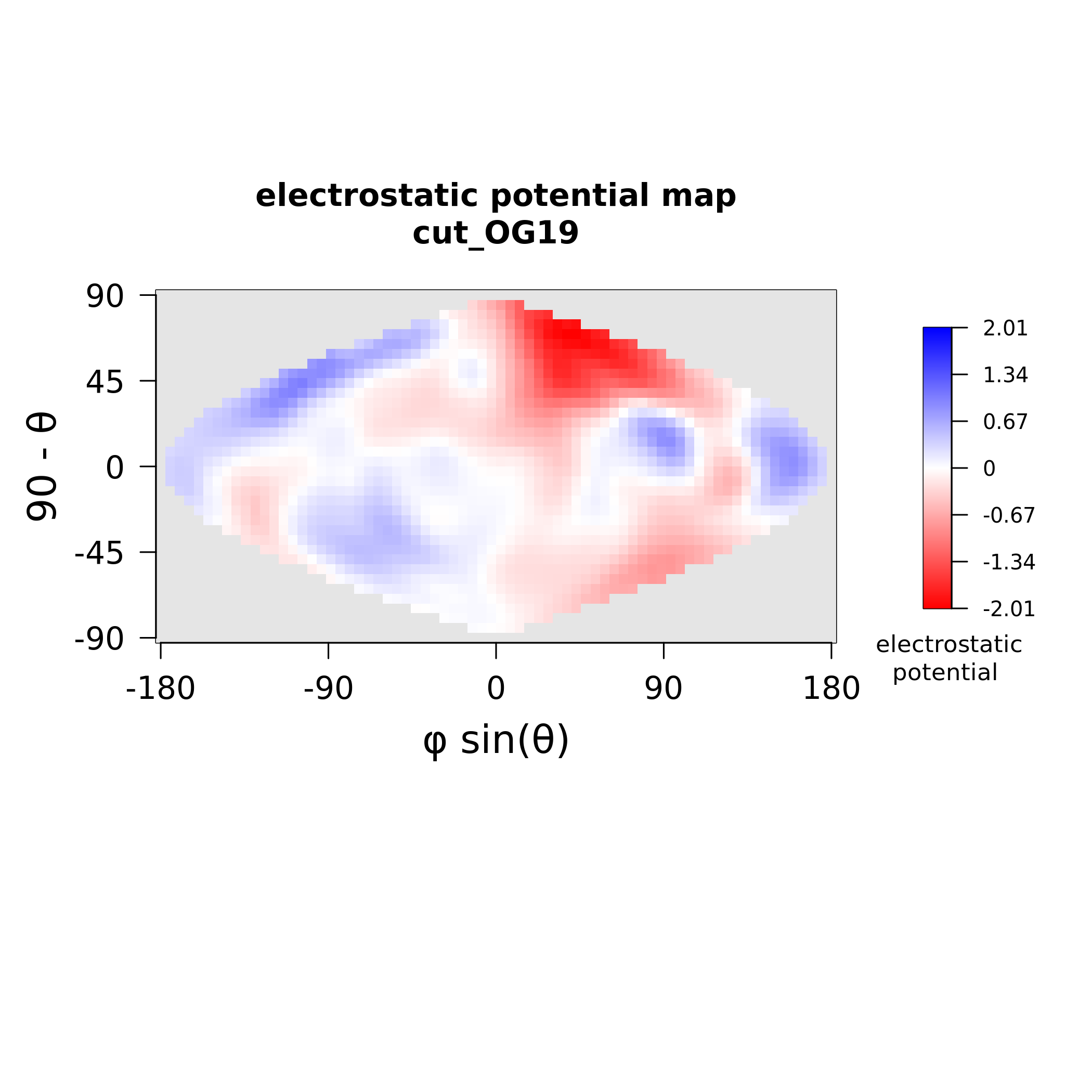

### MAX19_stickiness.png

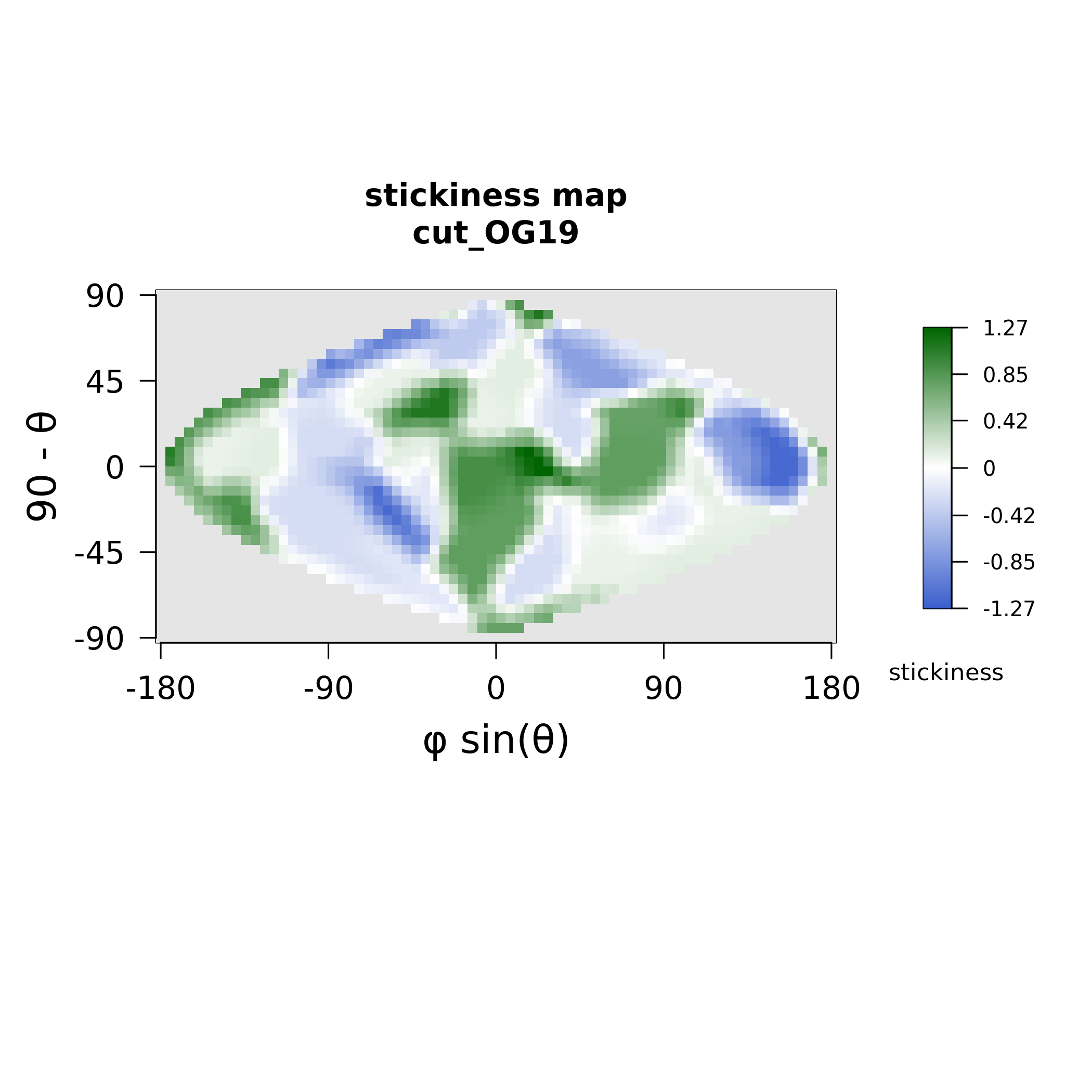

### MAX20_electrostatics.png

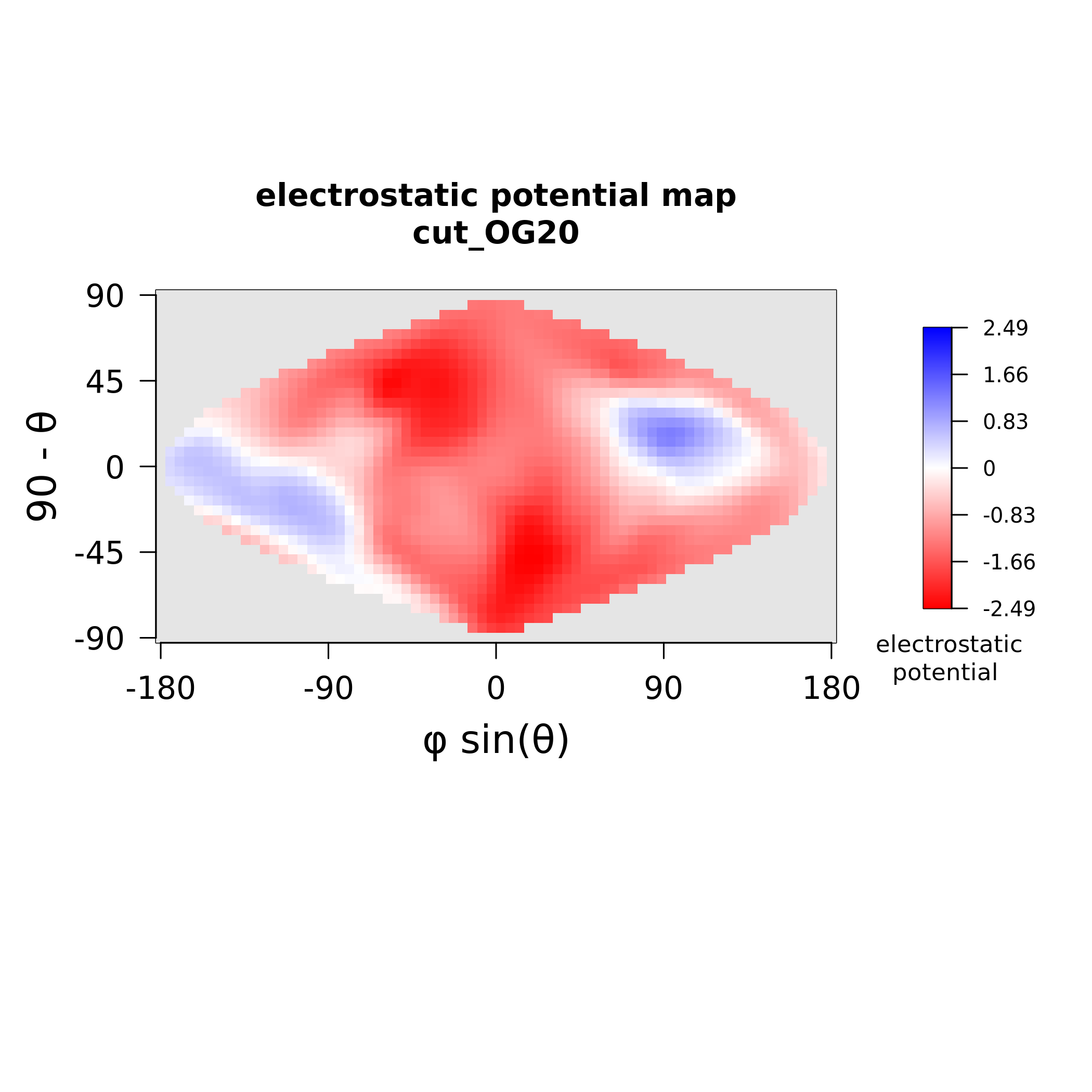

### MAX20_stickiness.png

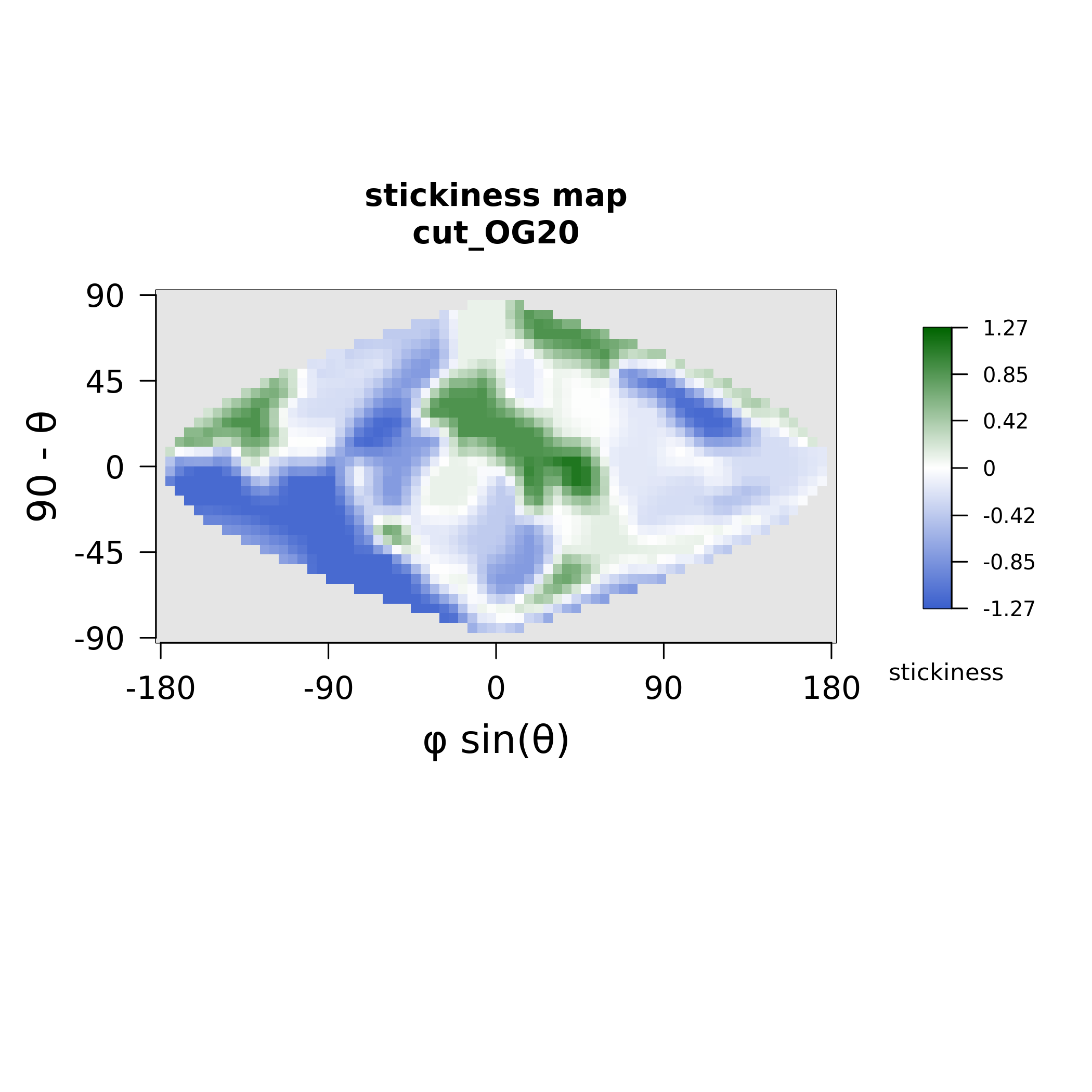

### MAX21_electrostatics.png

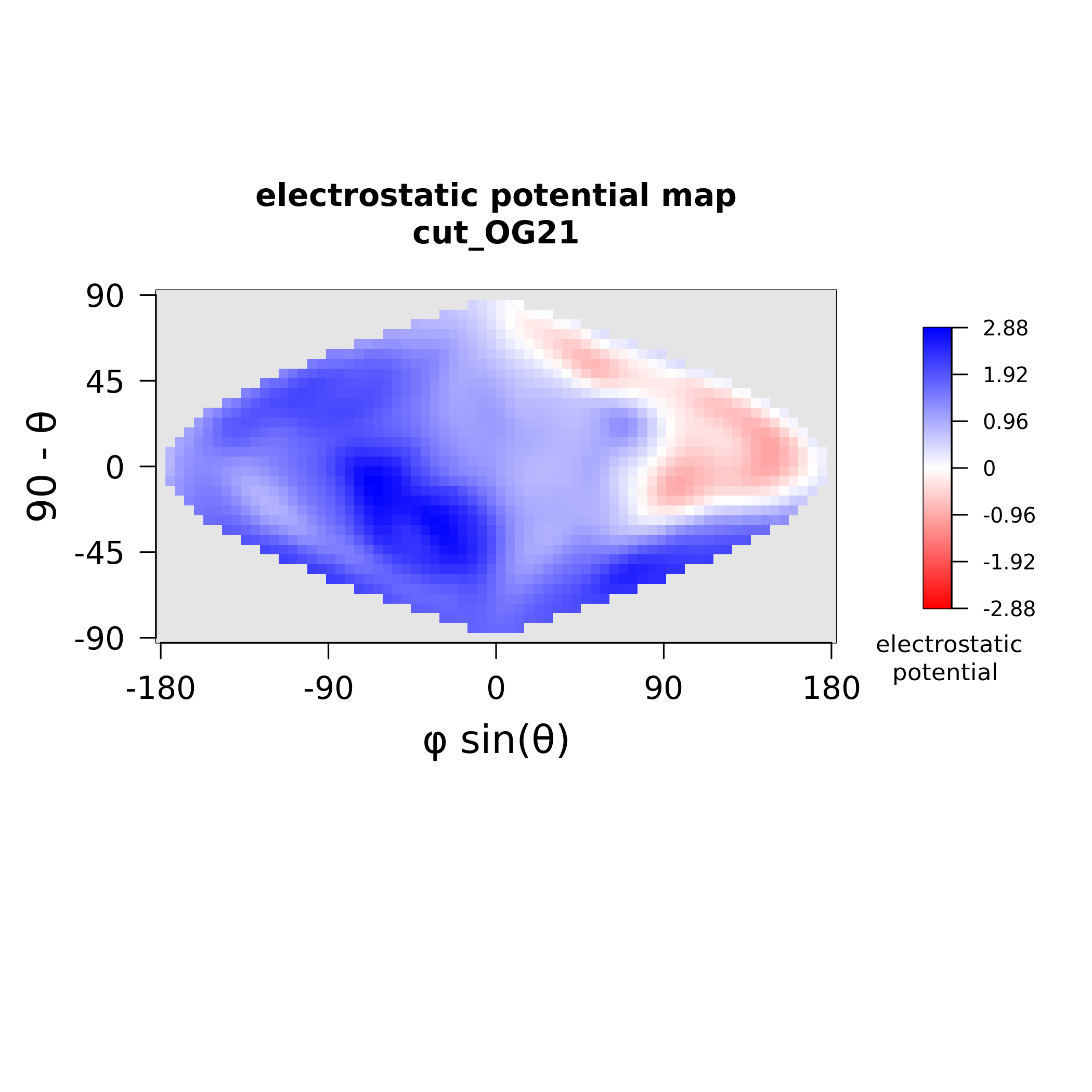

### MAX21_stickiness.png

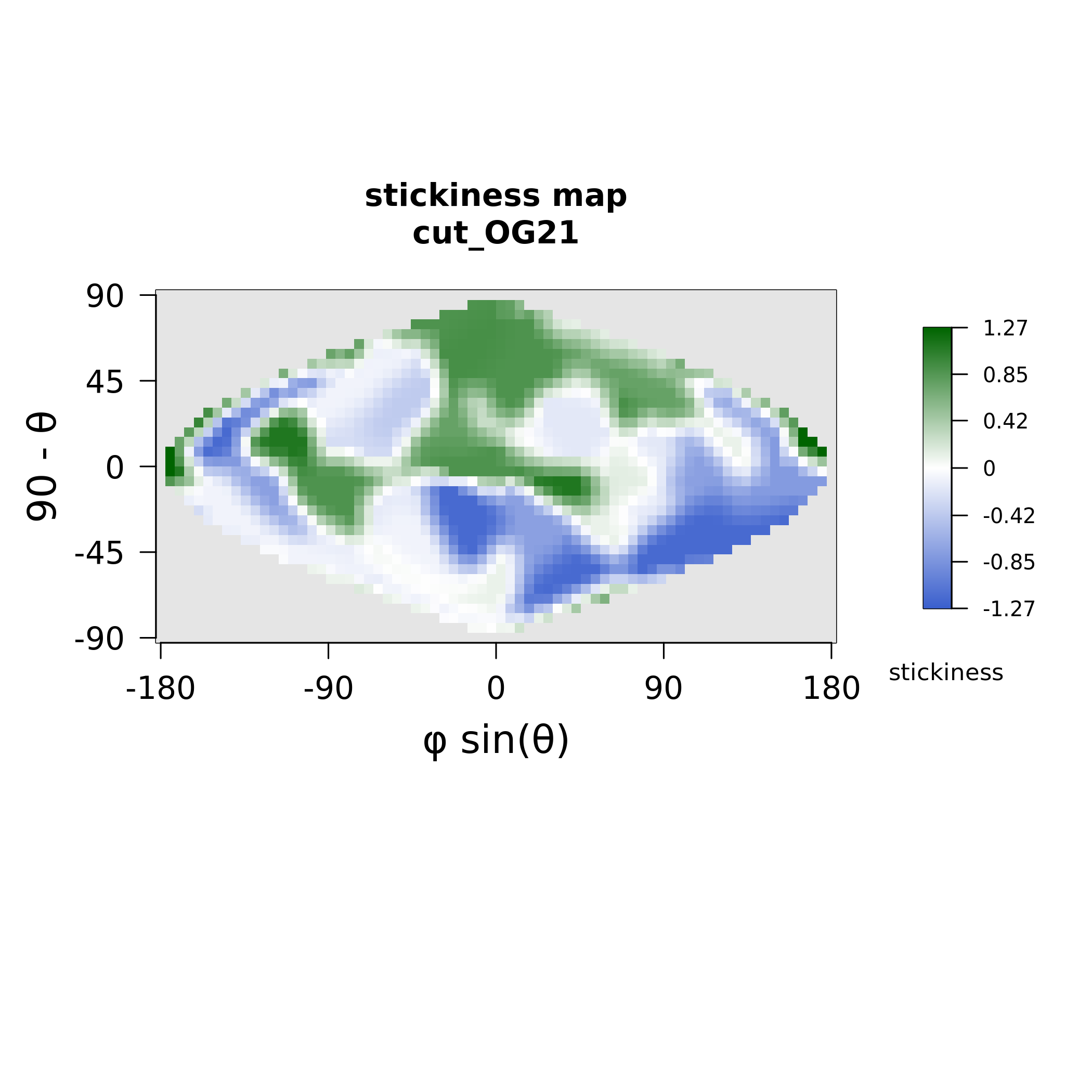

### MAX23_electrostatics.png

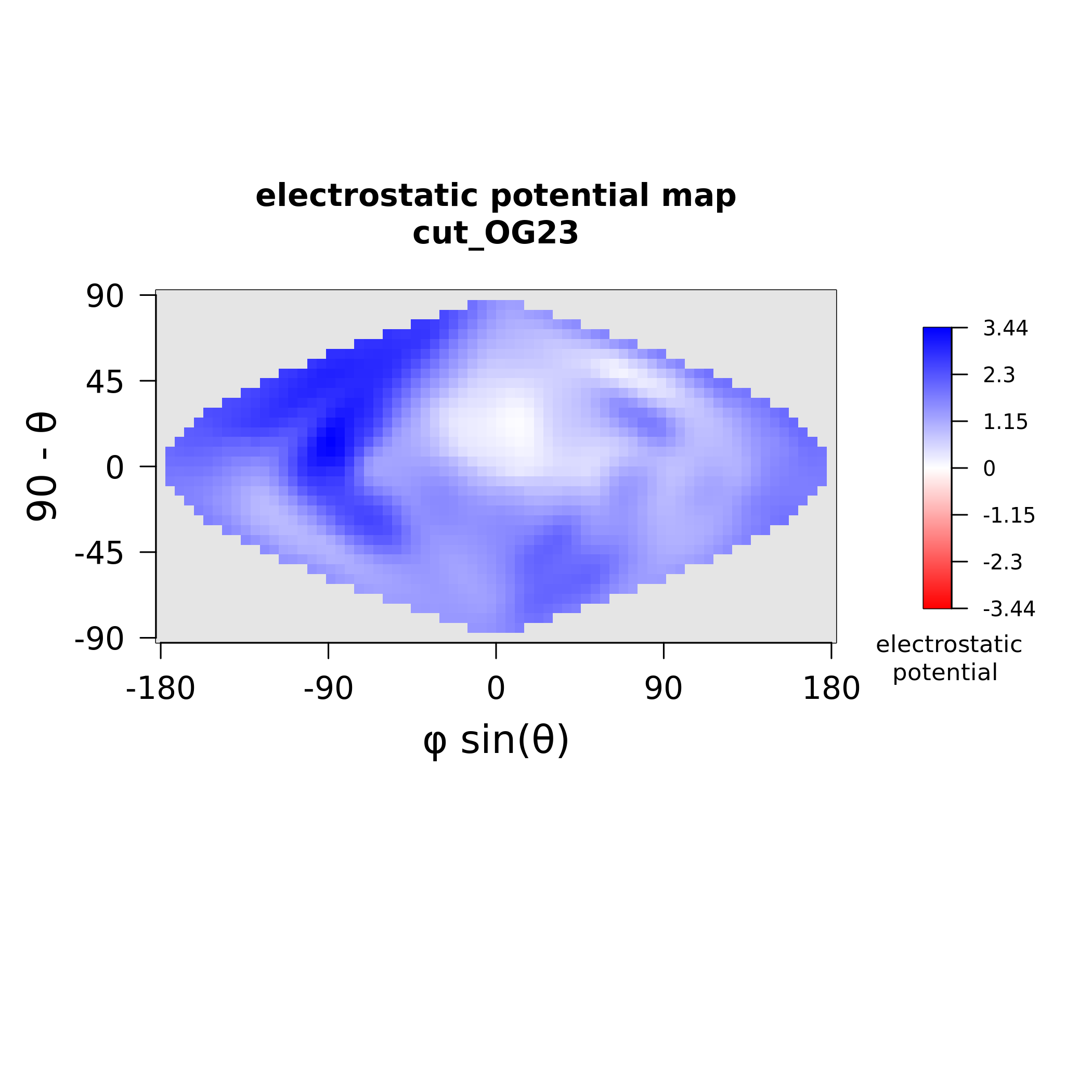

### MAX23_stickiness.png

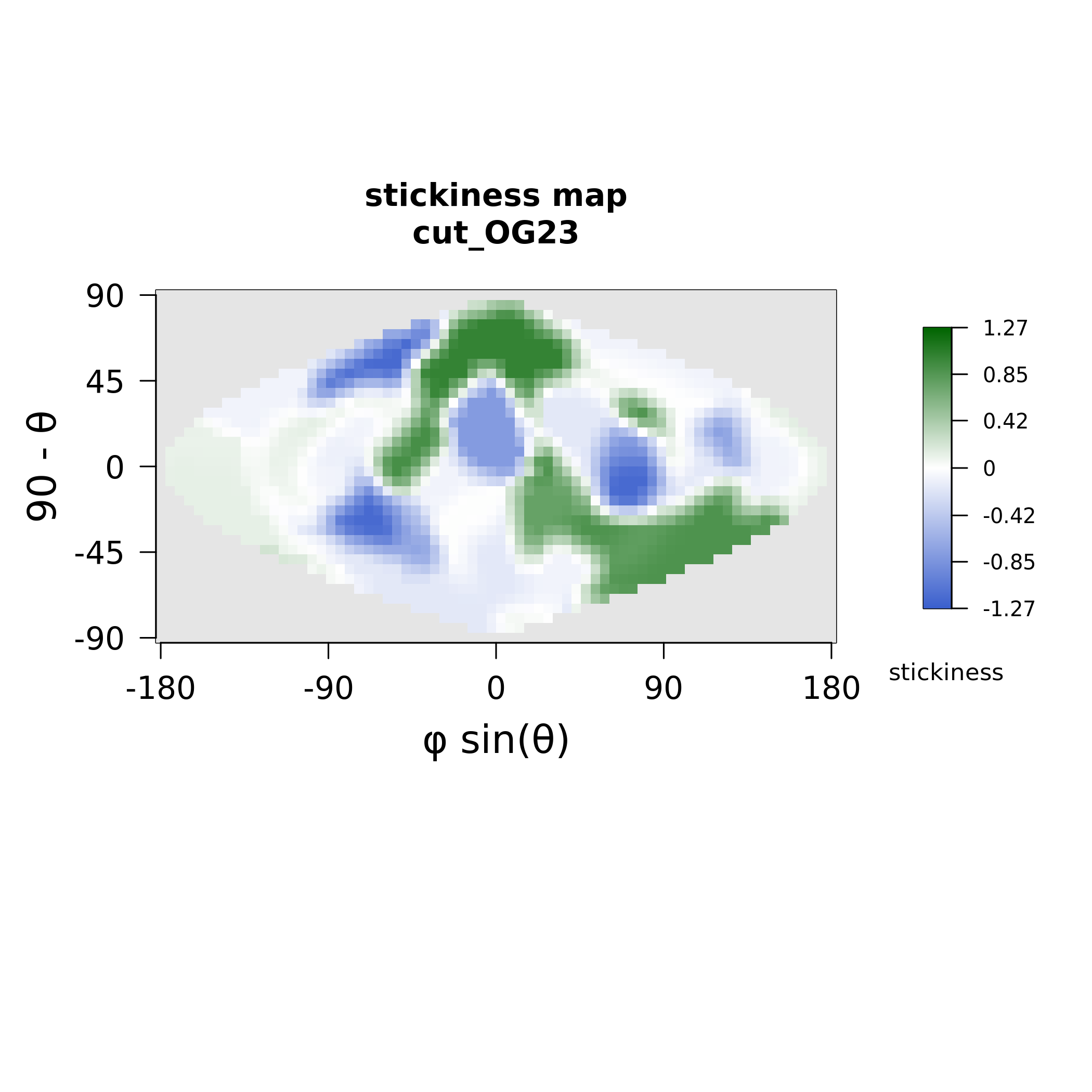

### MAX24_electrostatics.png

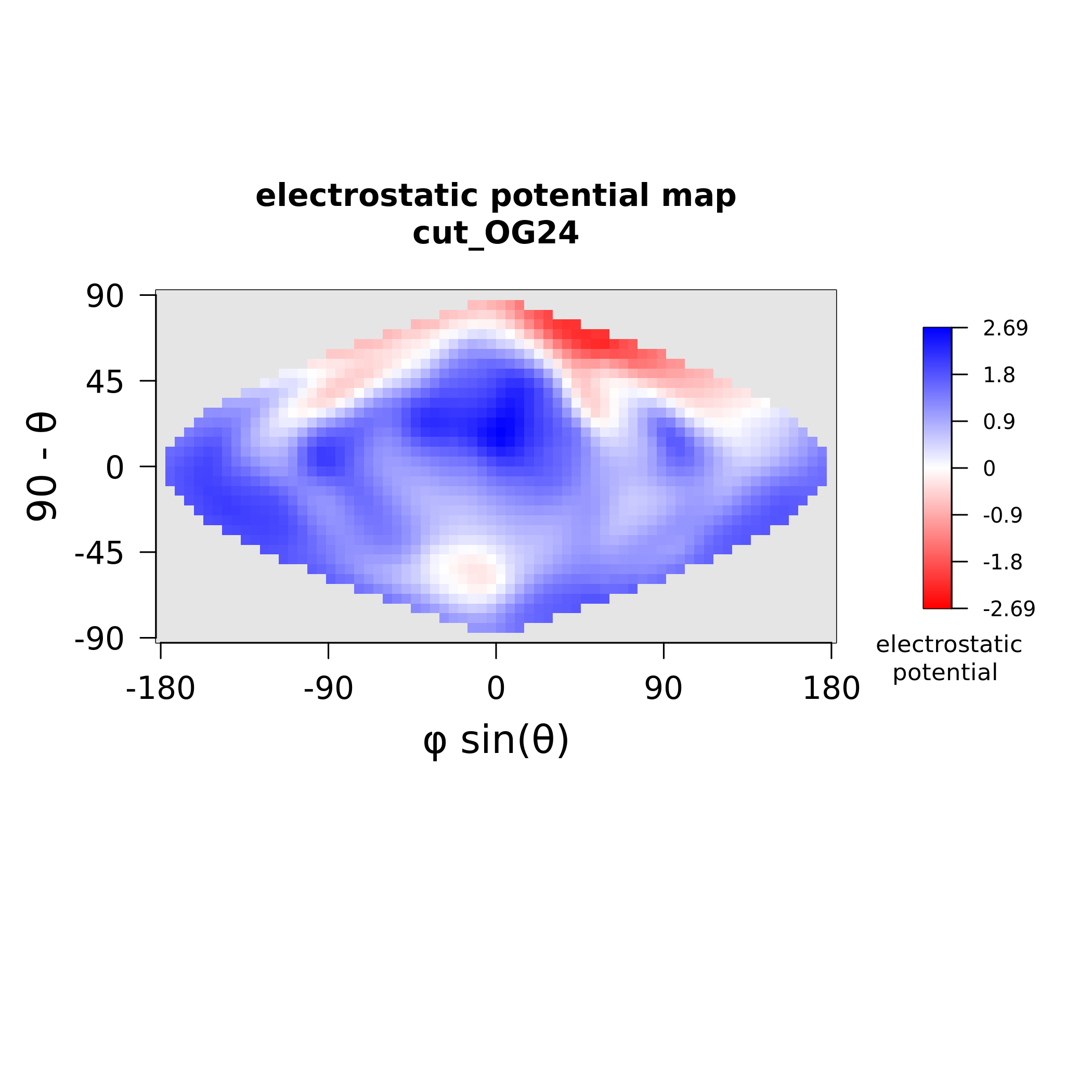

### MAX24_stickiness.png

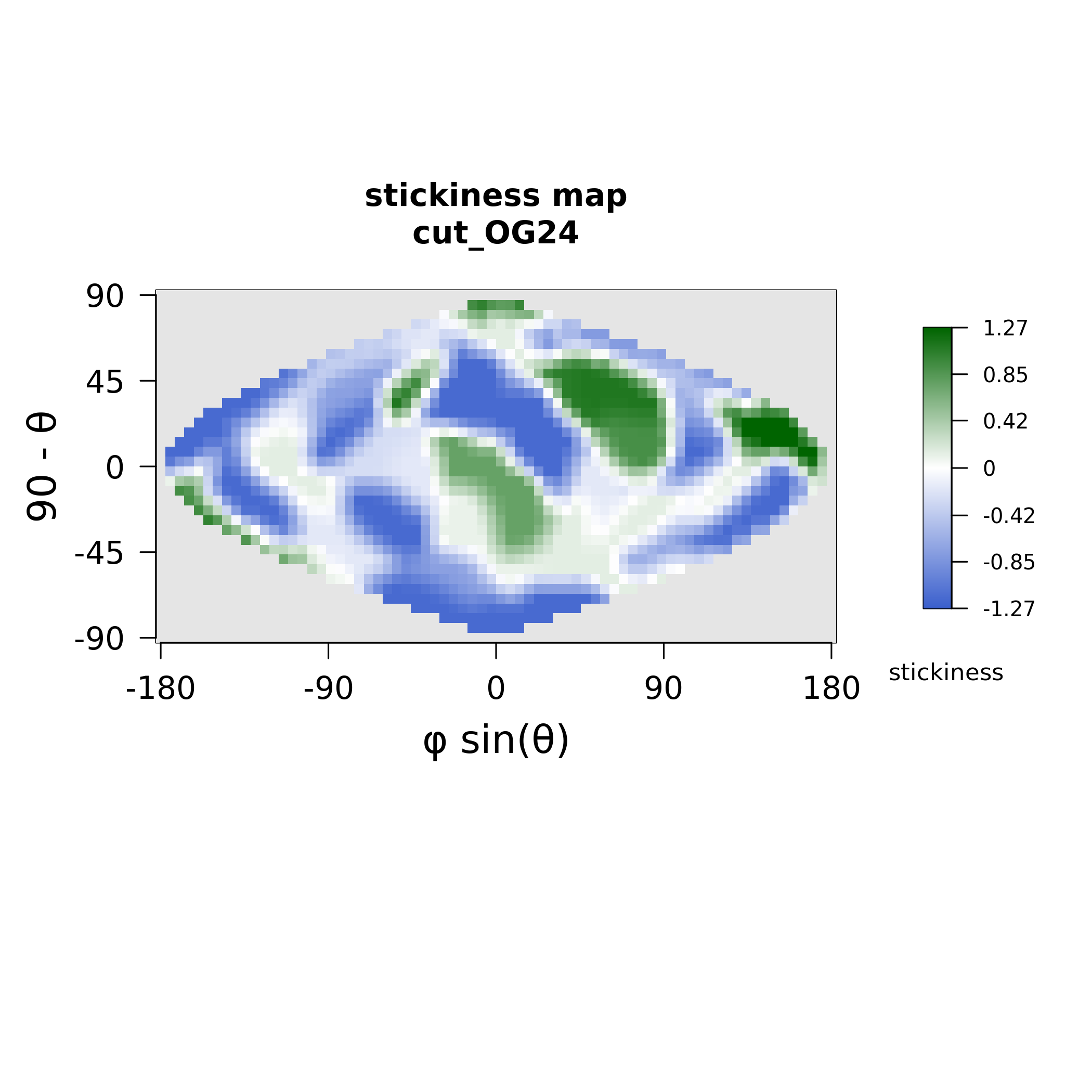

### MAX27_electrostatics.png

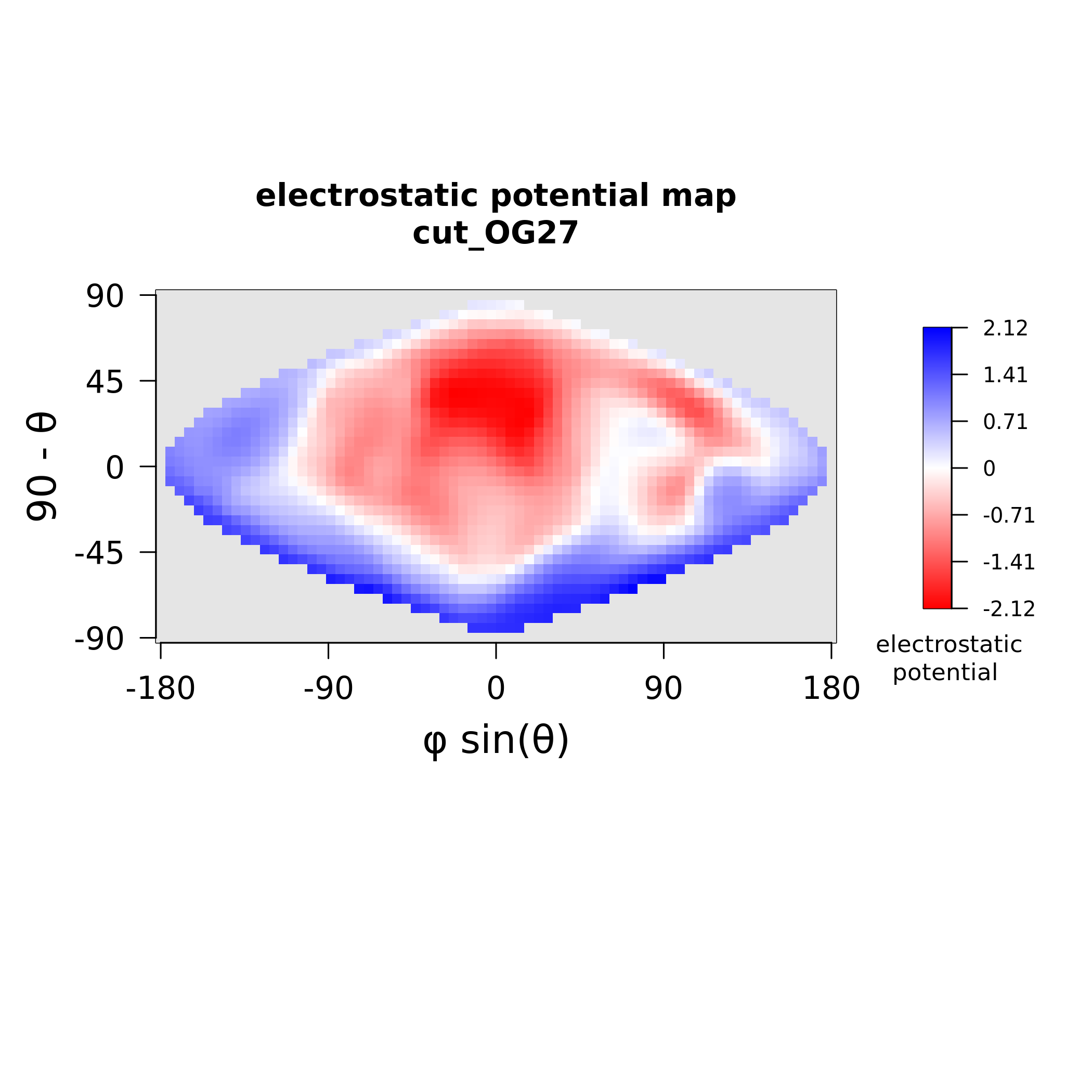

### MAX27_stickiness.png

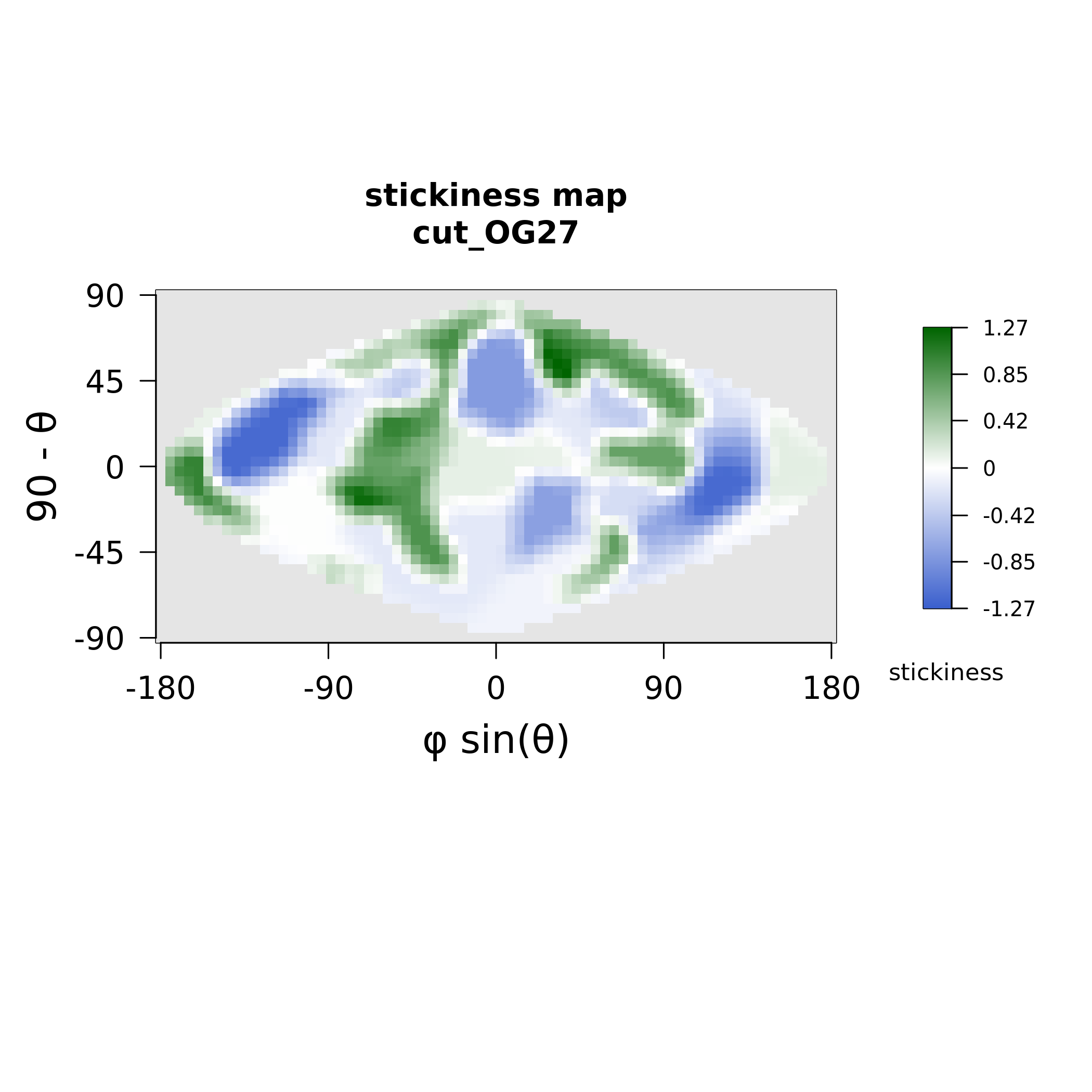

### MAX28_electrostatics.png

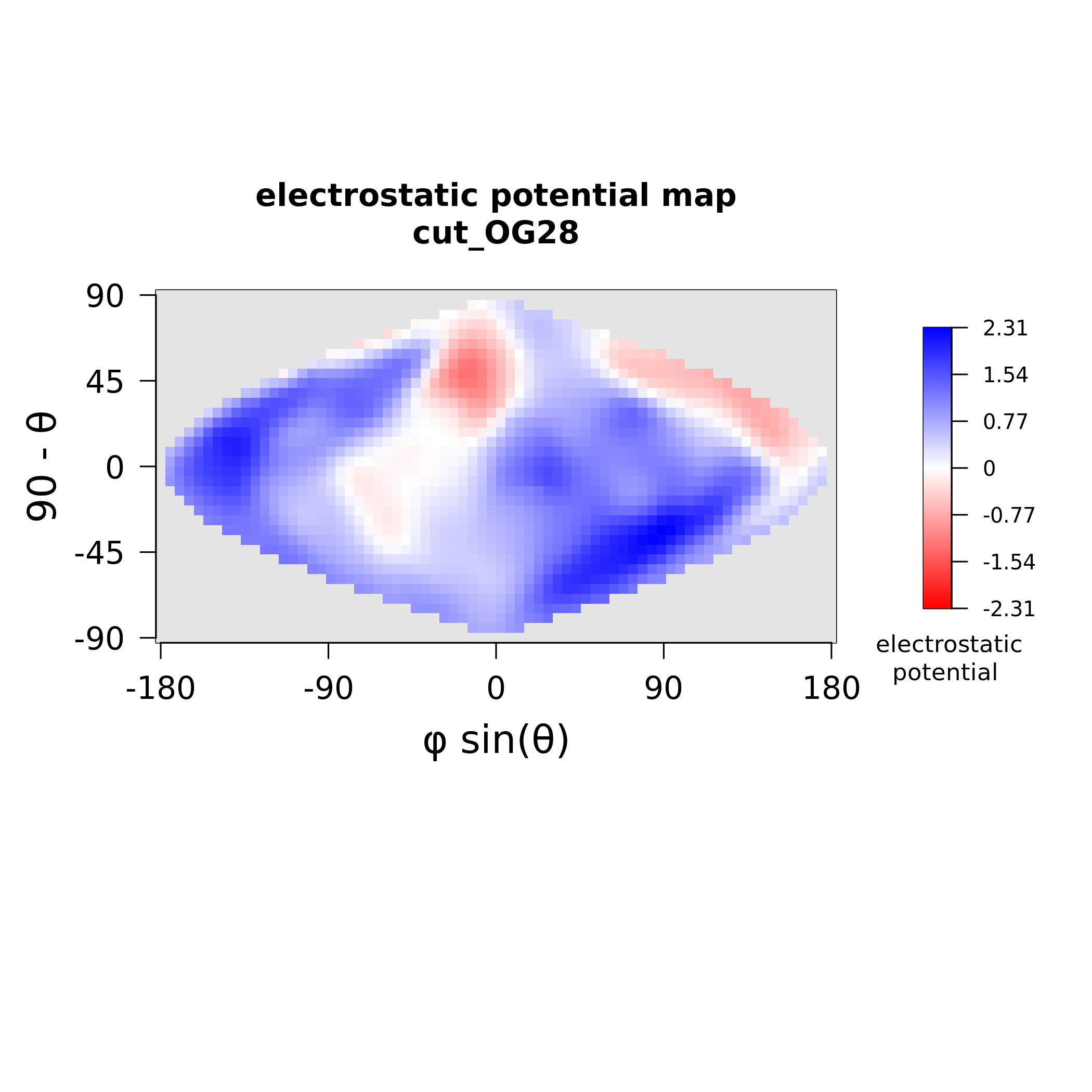

### MAX28_stickiness.png

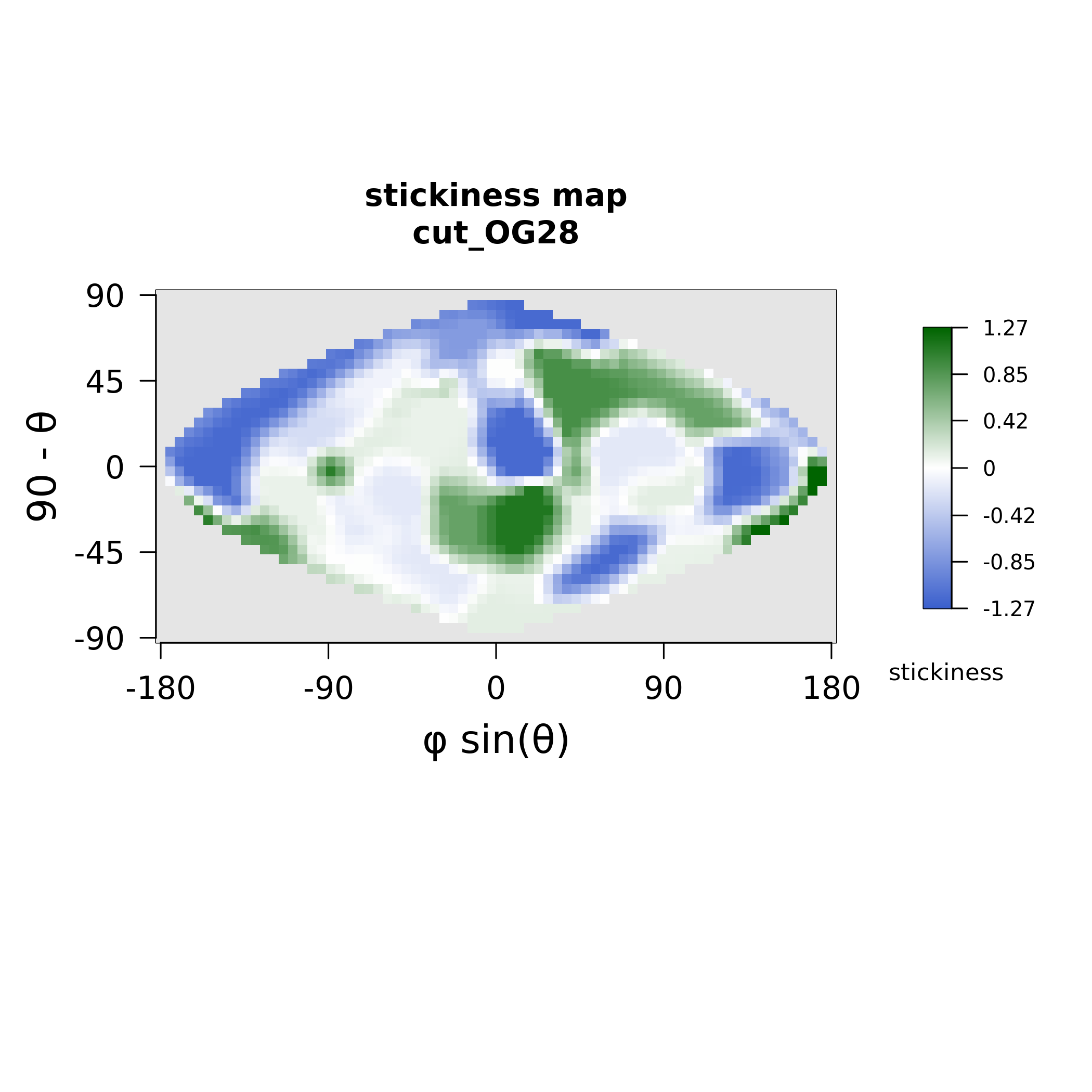

### MAX30_electrostatics.png

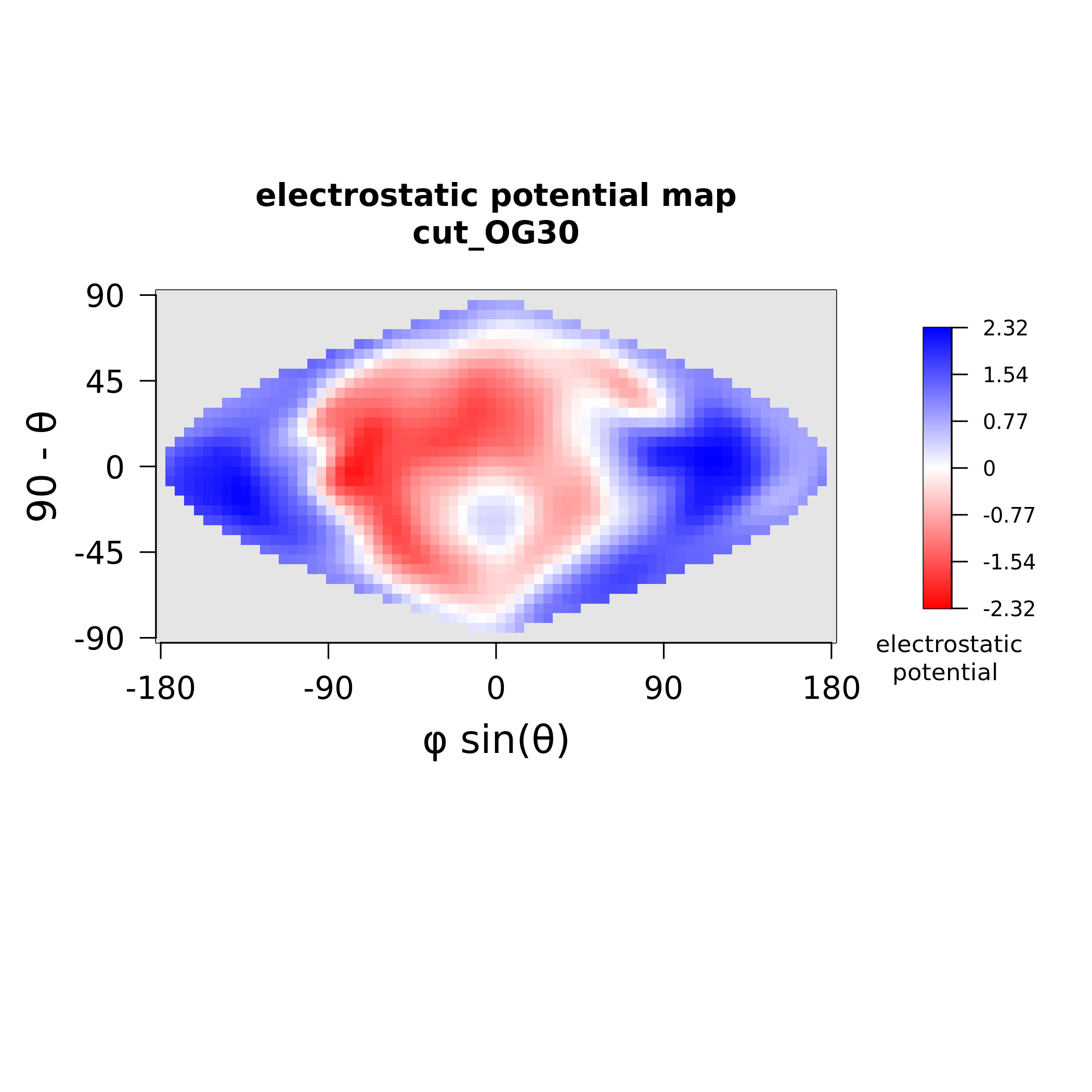

### MAX30_stickiness.png

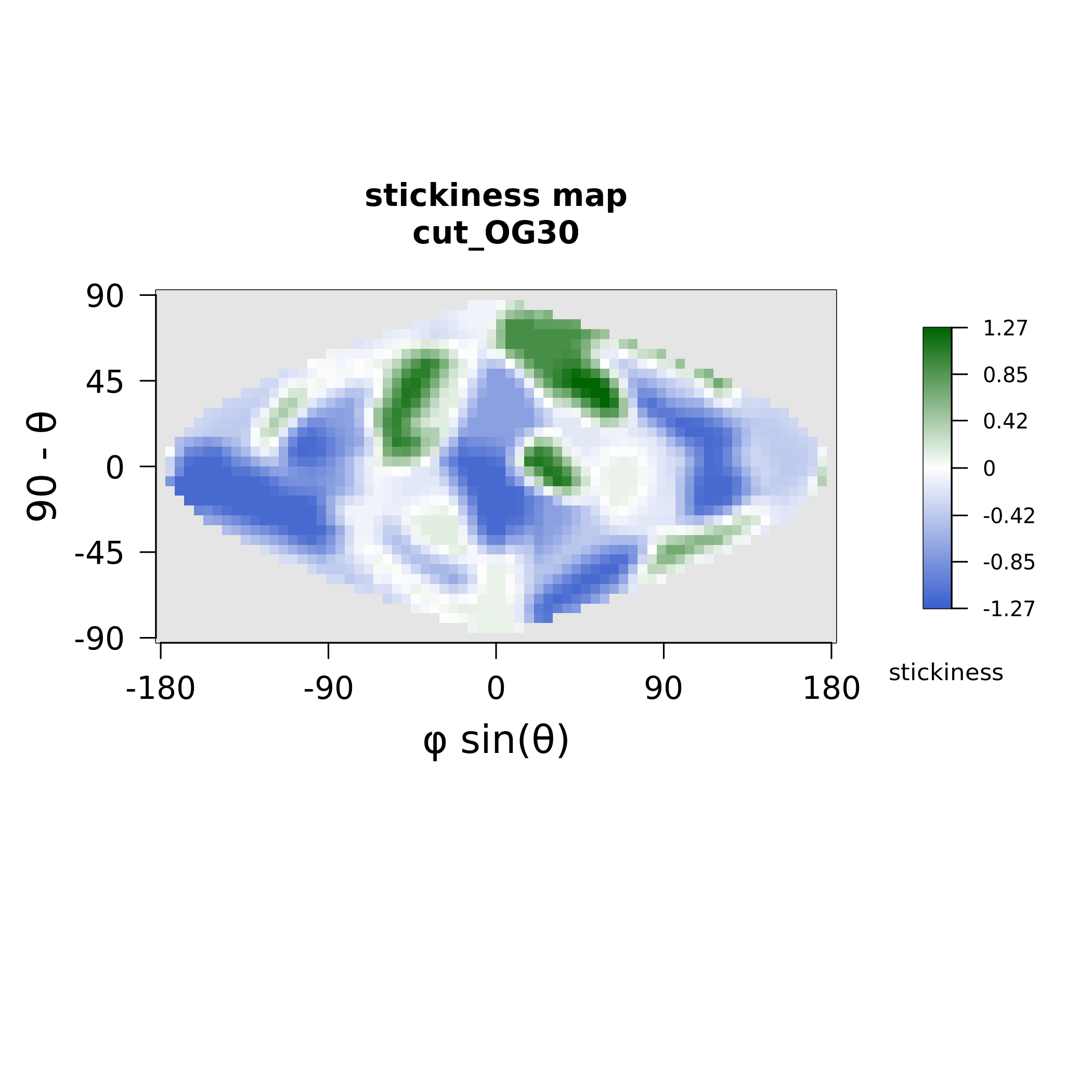

### MAX31_electrostatics.png

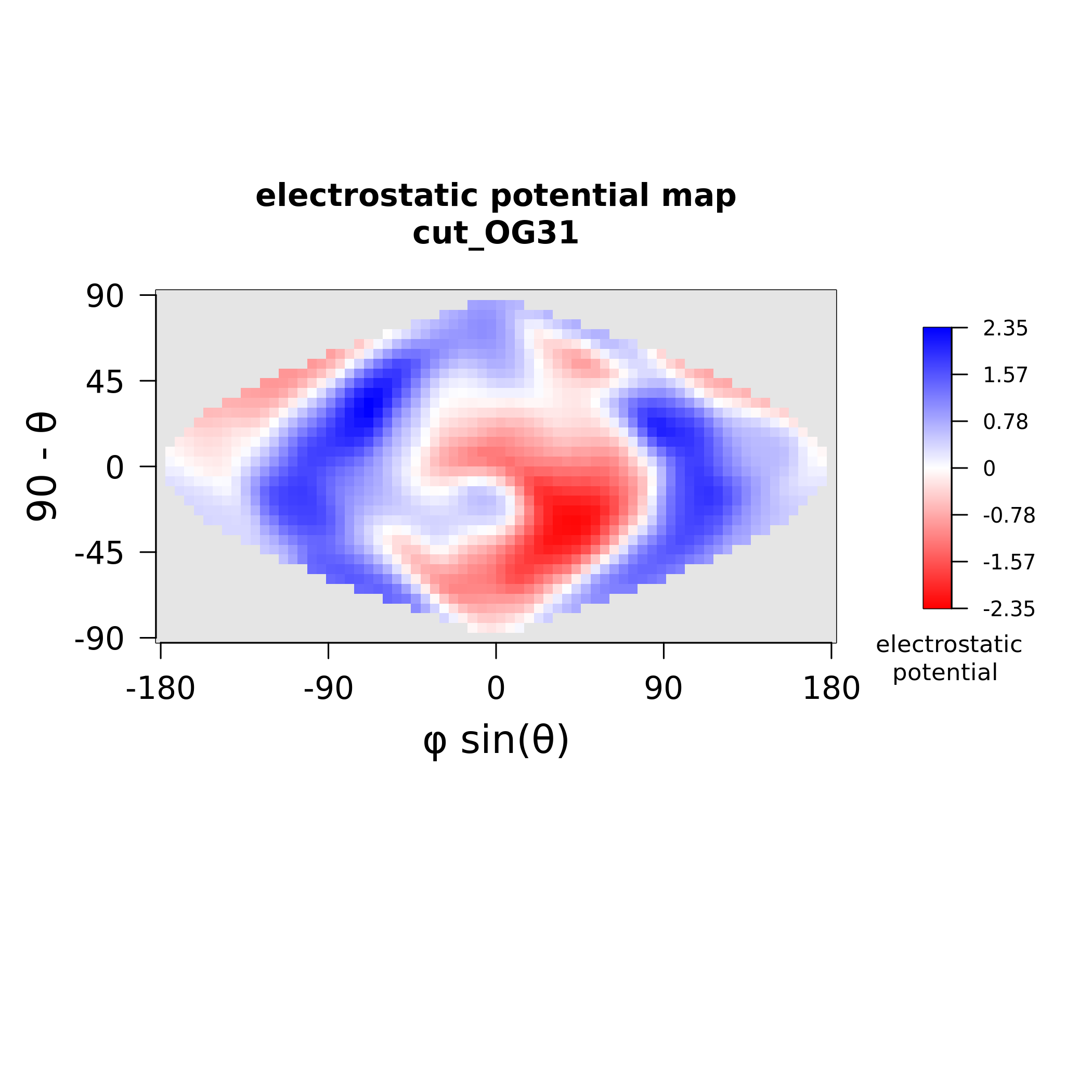

### MAX31_stickiness.png

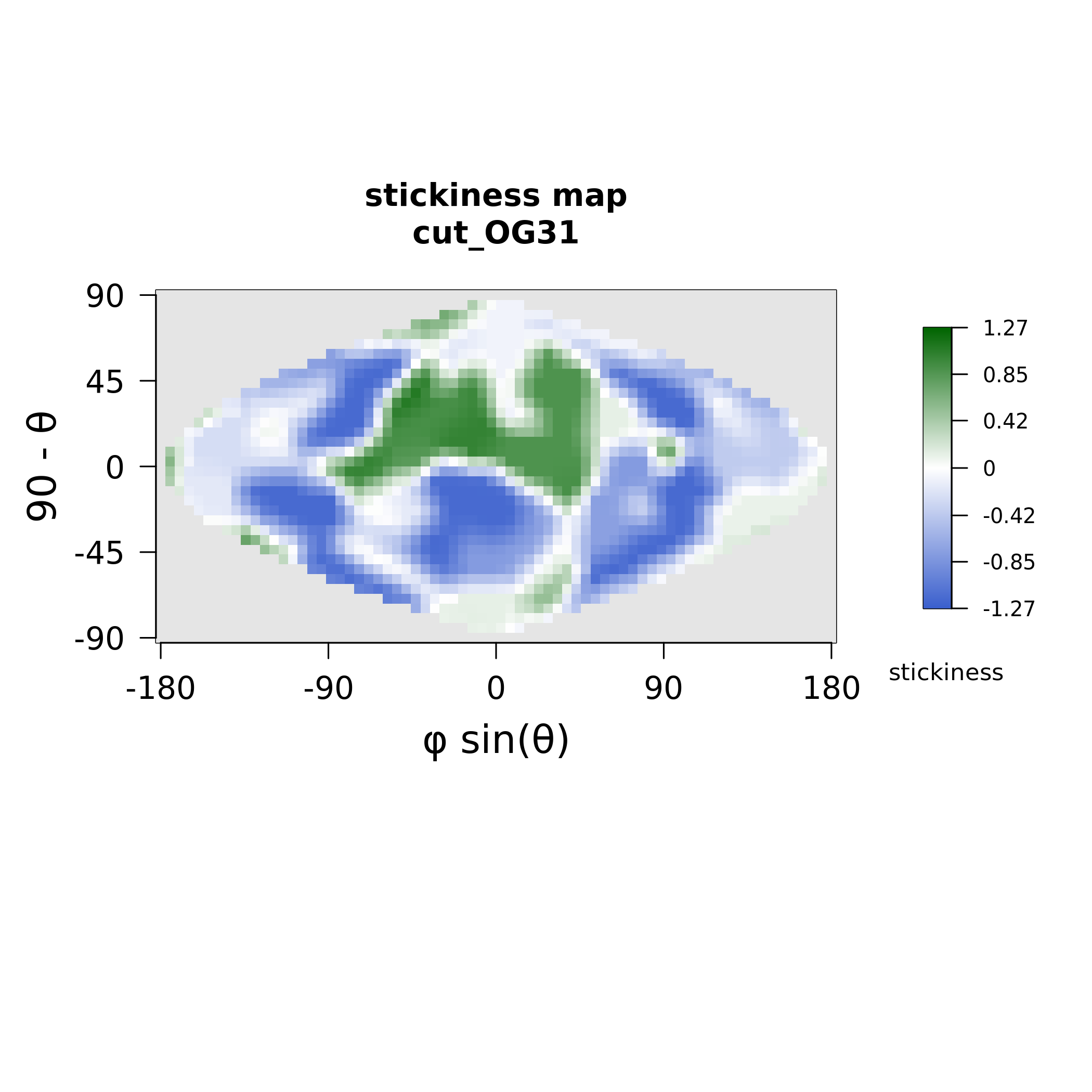

### MAX33_electrostatics.png

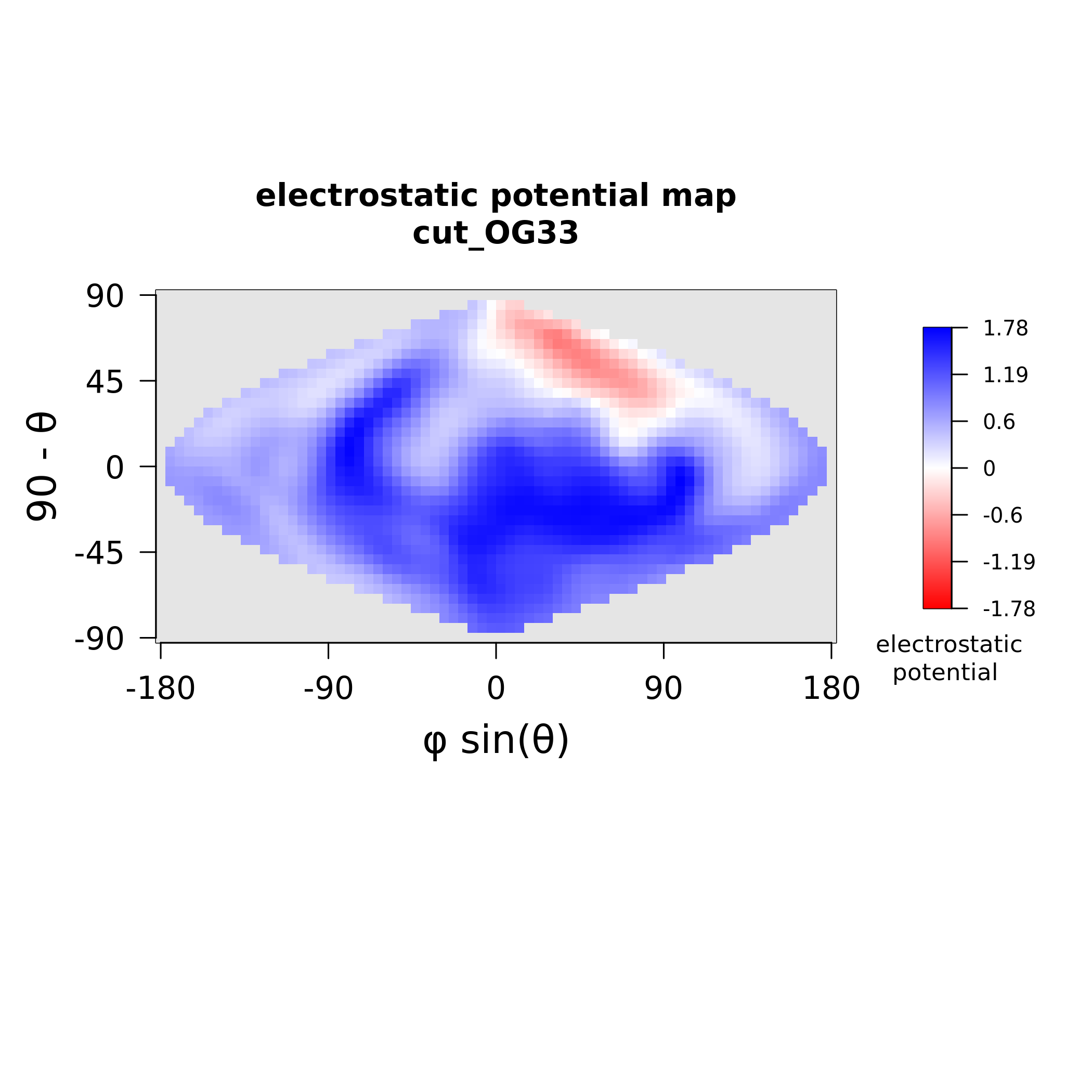

### MAX33_stickiness.png

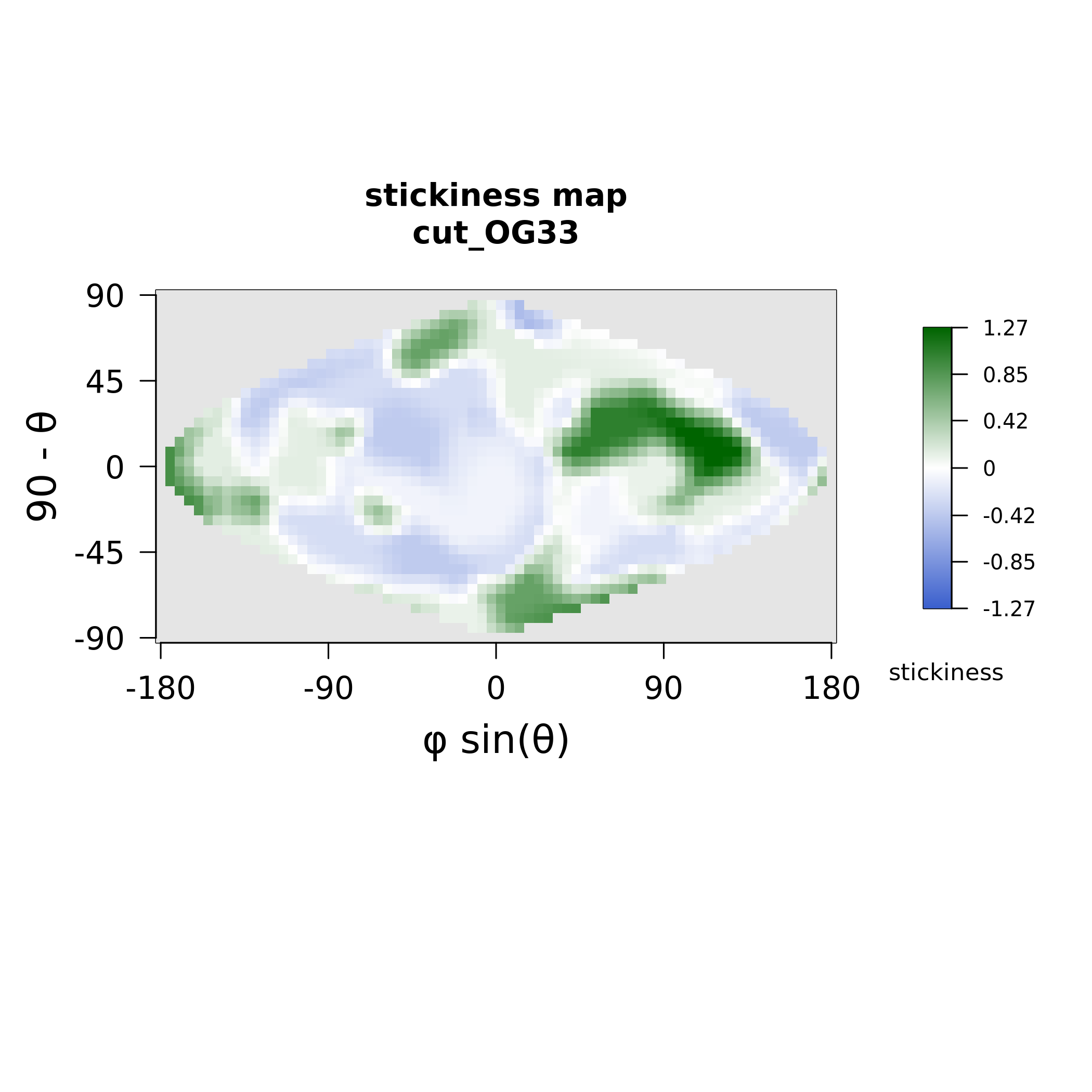

### MAX34_electrostatics.png

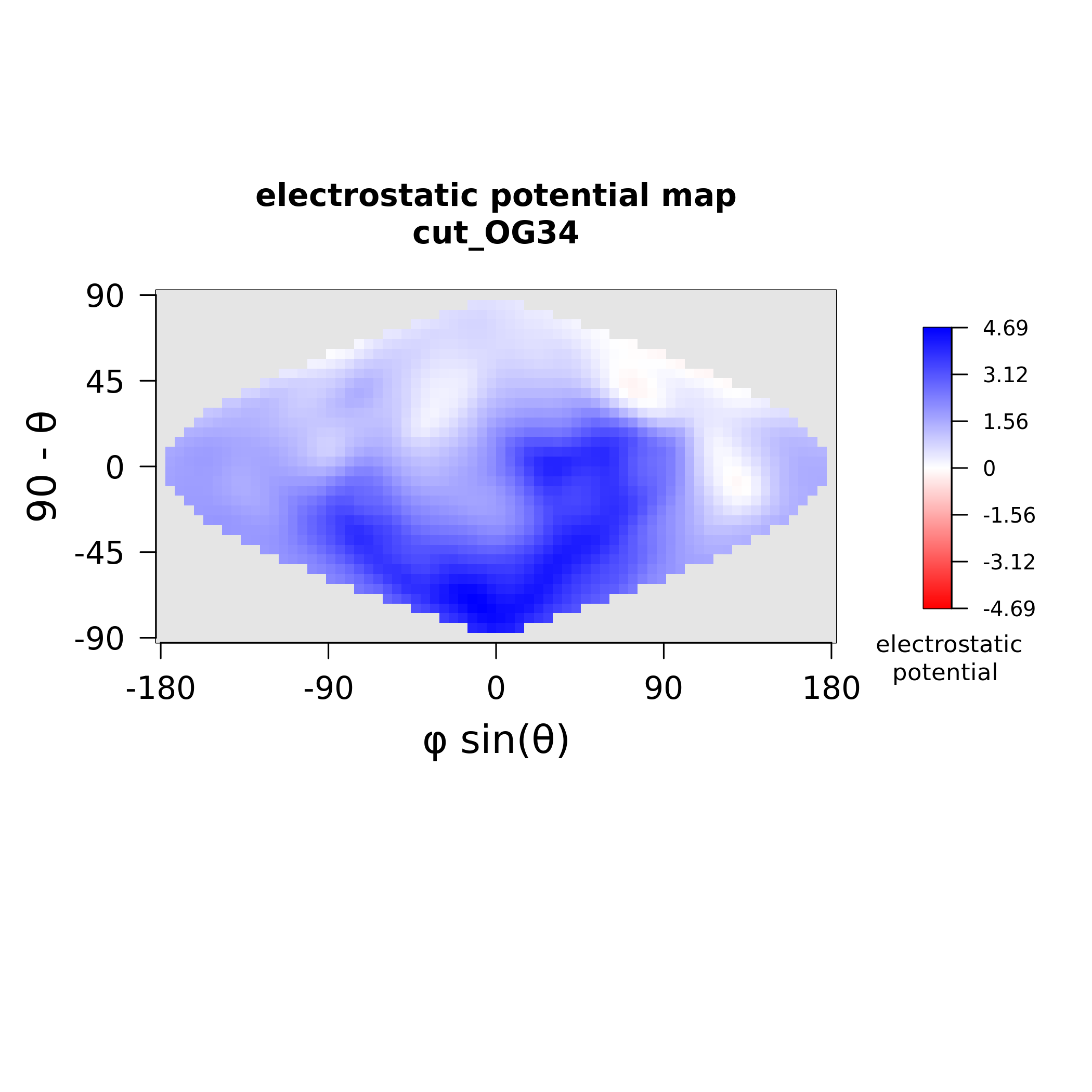

### MAX34_stickiness.png

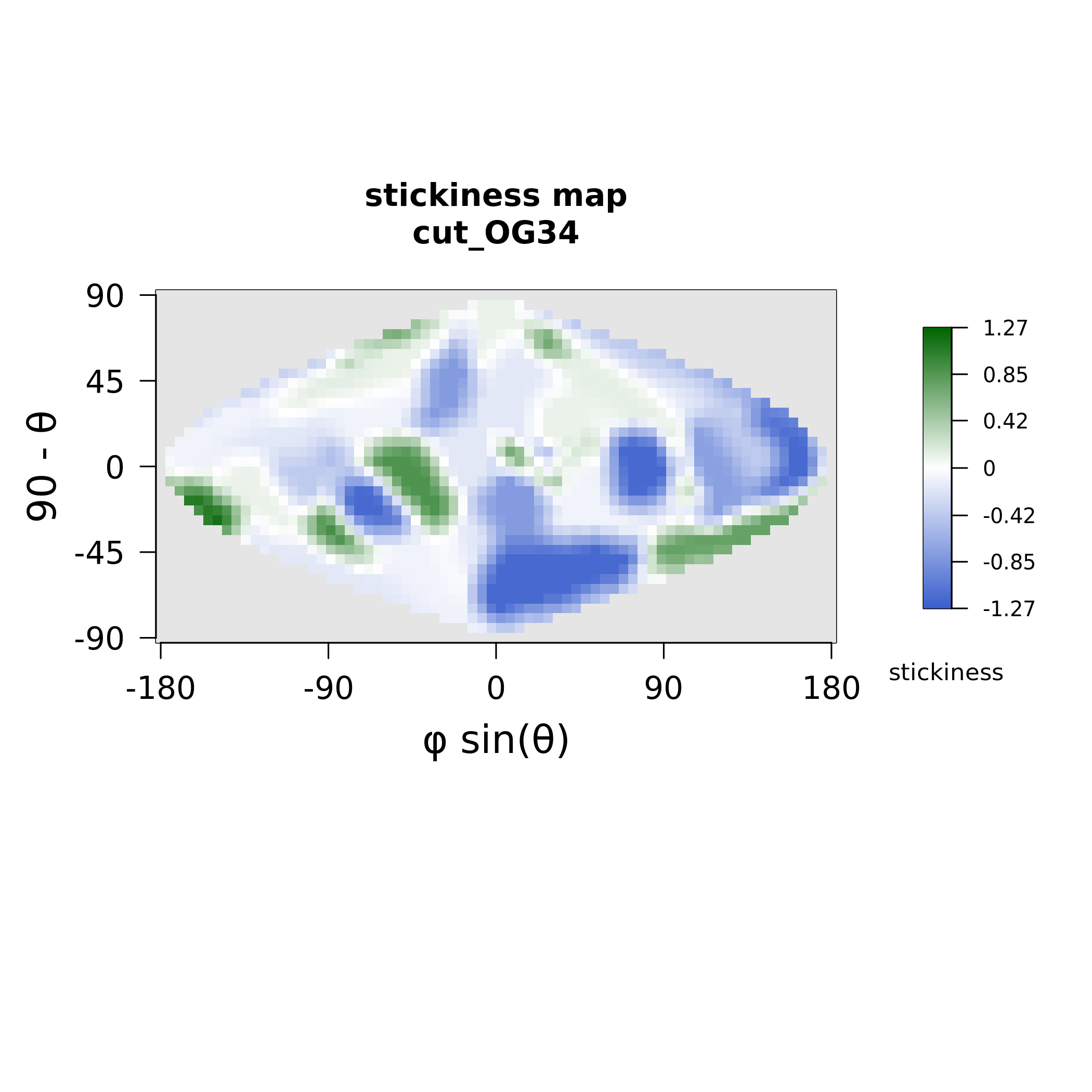

### MAX35_electrostatics.png

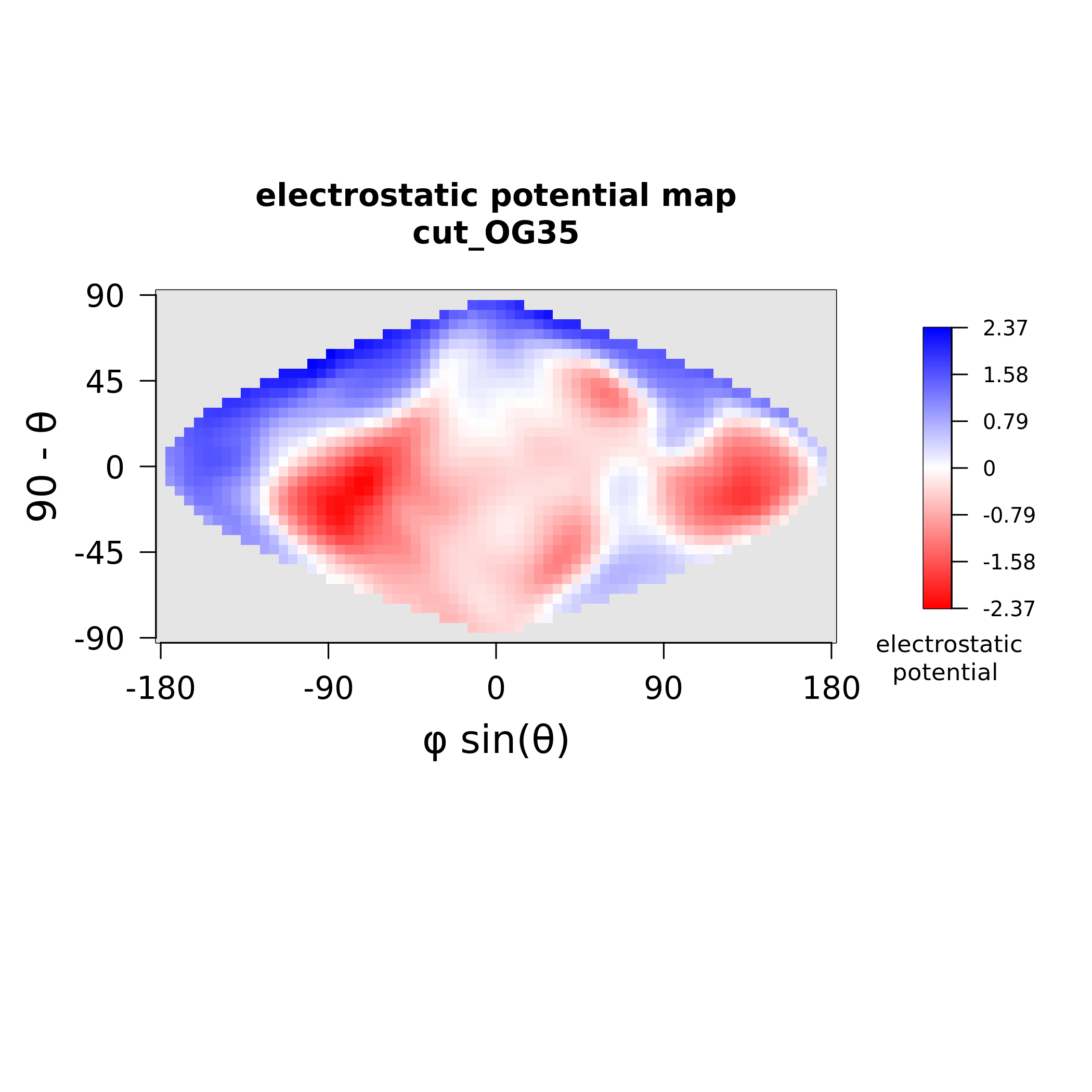

### MAX35_stickiness.png

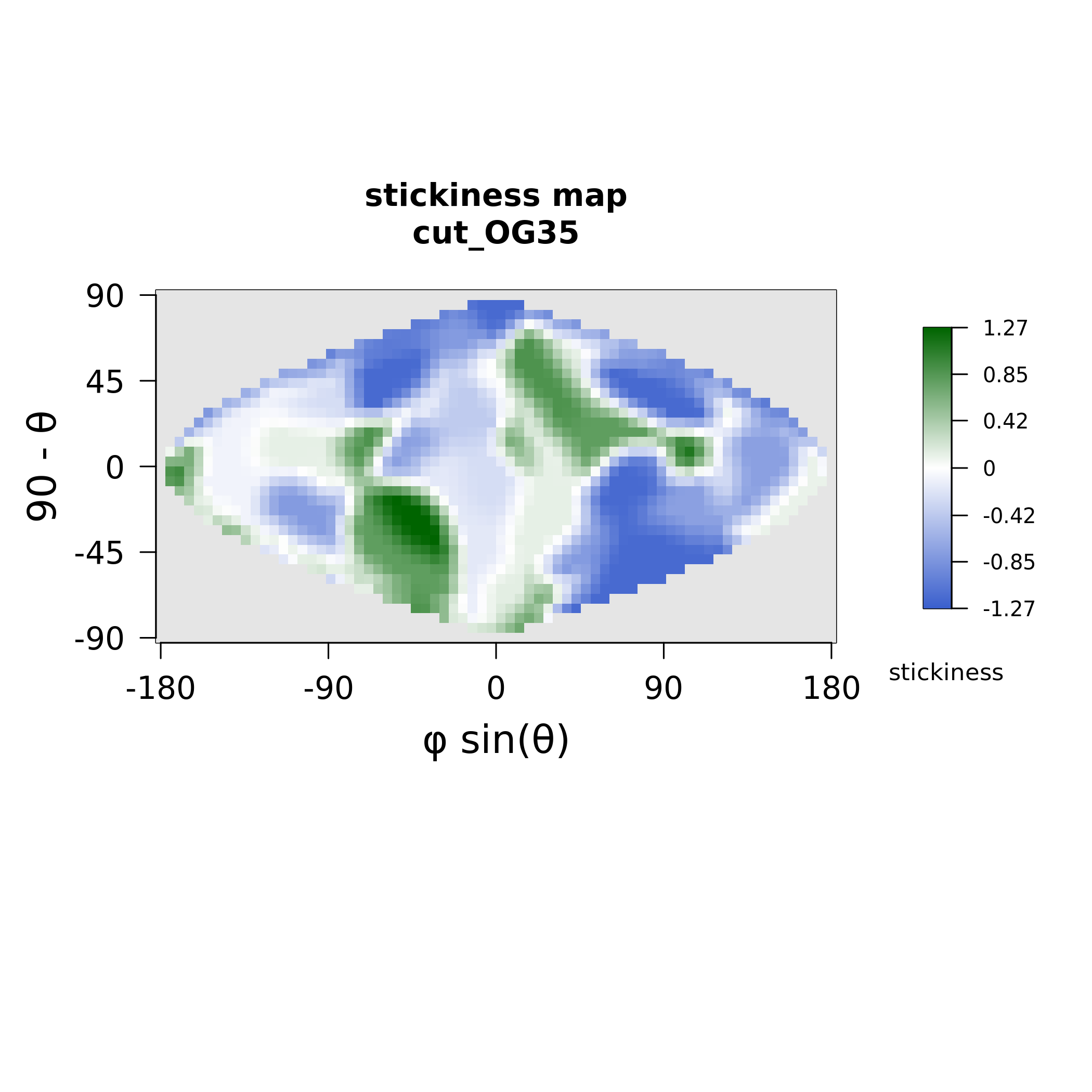

### MAX37_electrostatics.png

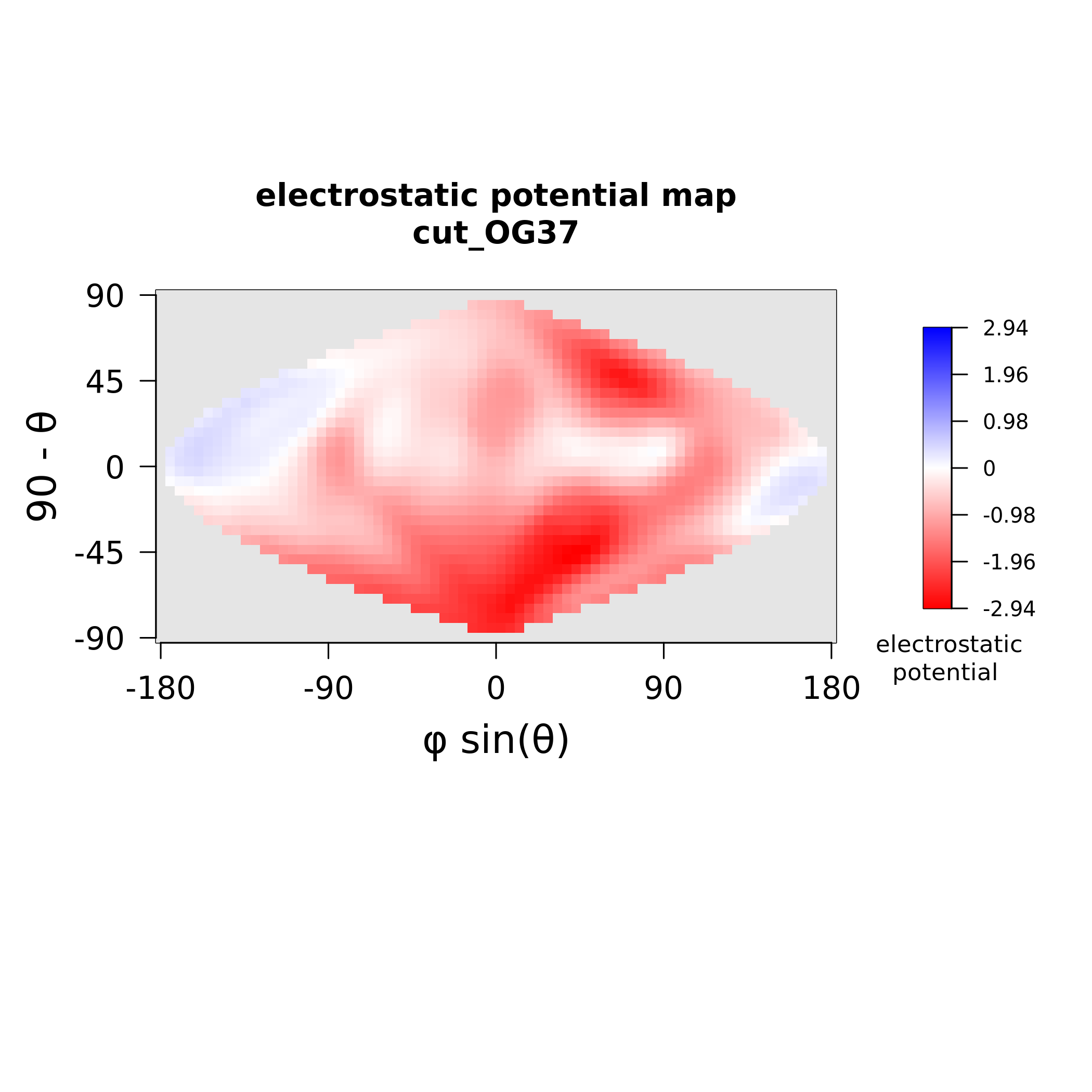

### MAX37_stickiness.png

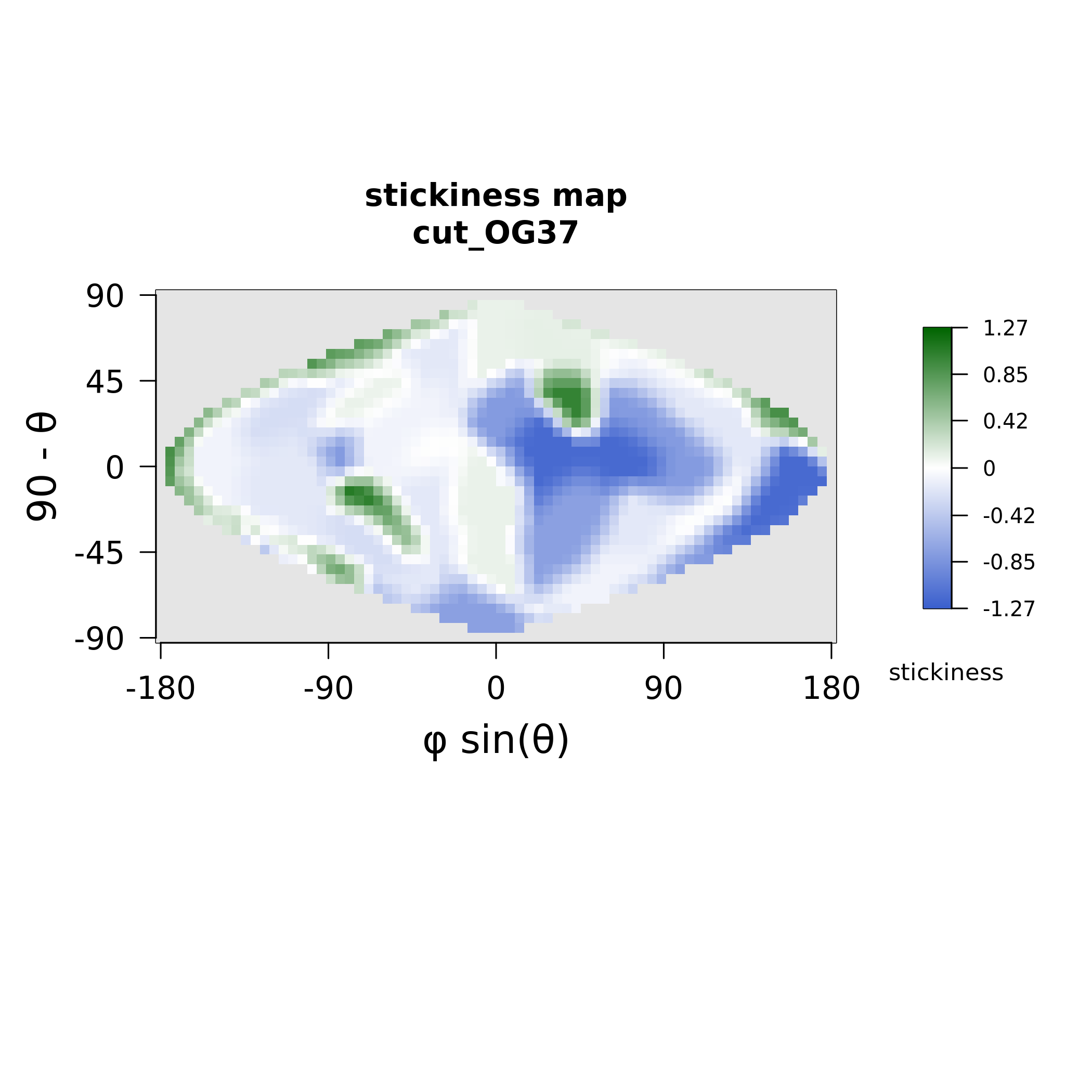

### MAX38_electrostatics.png

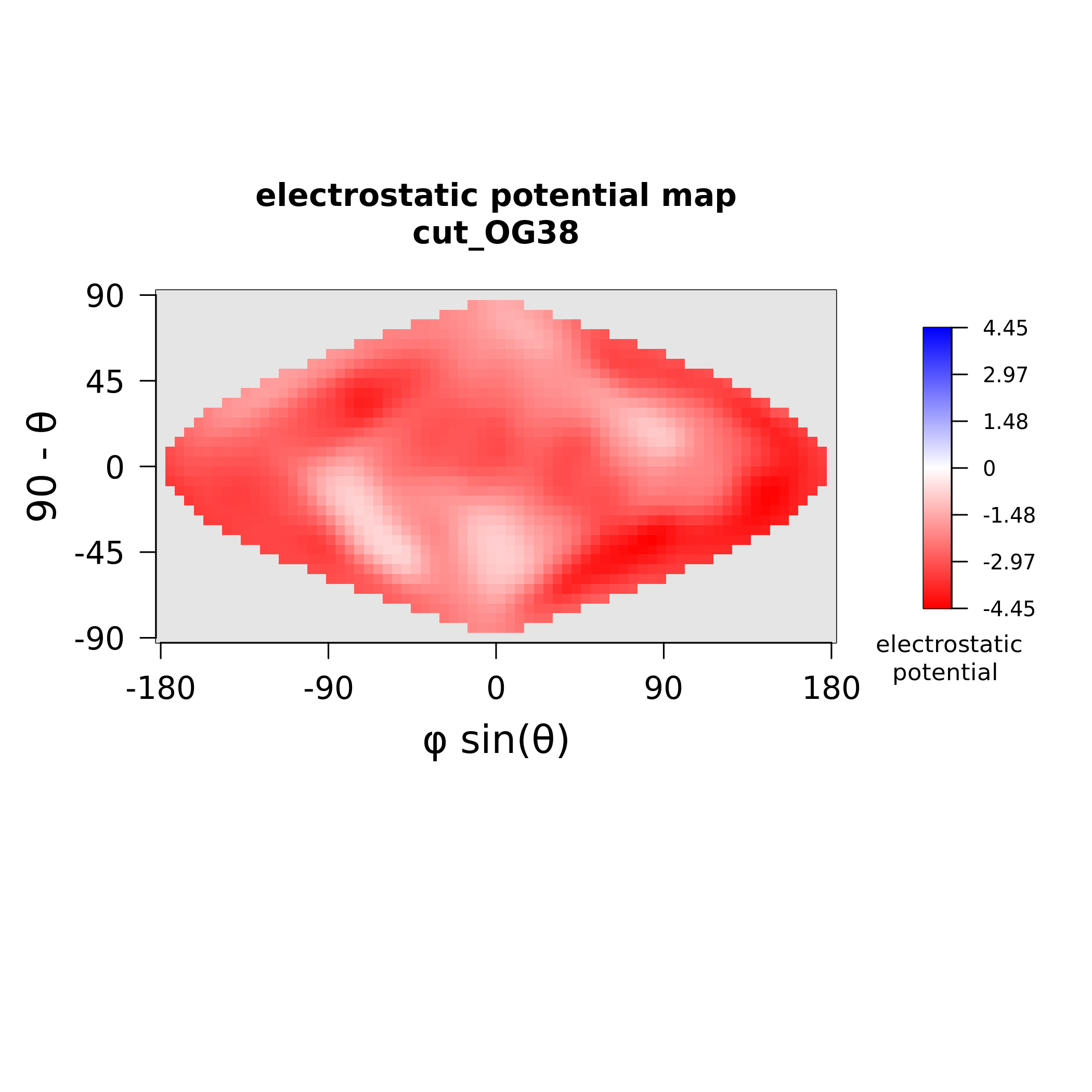

### MAX38_stickiness.png

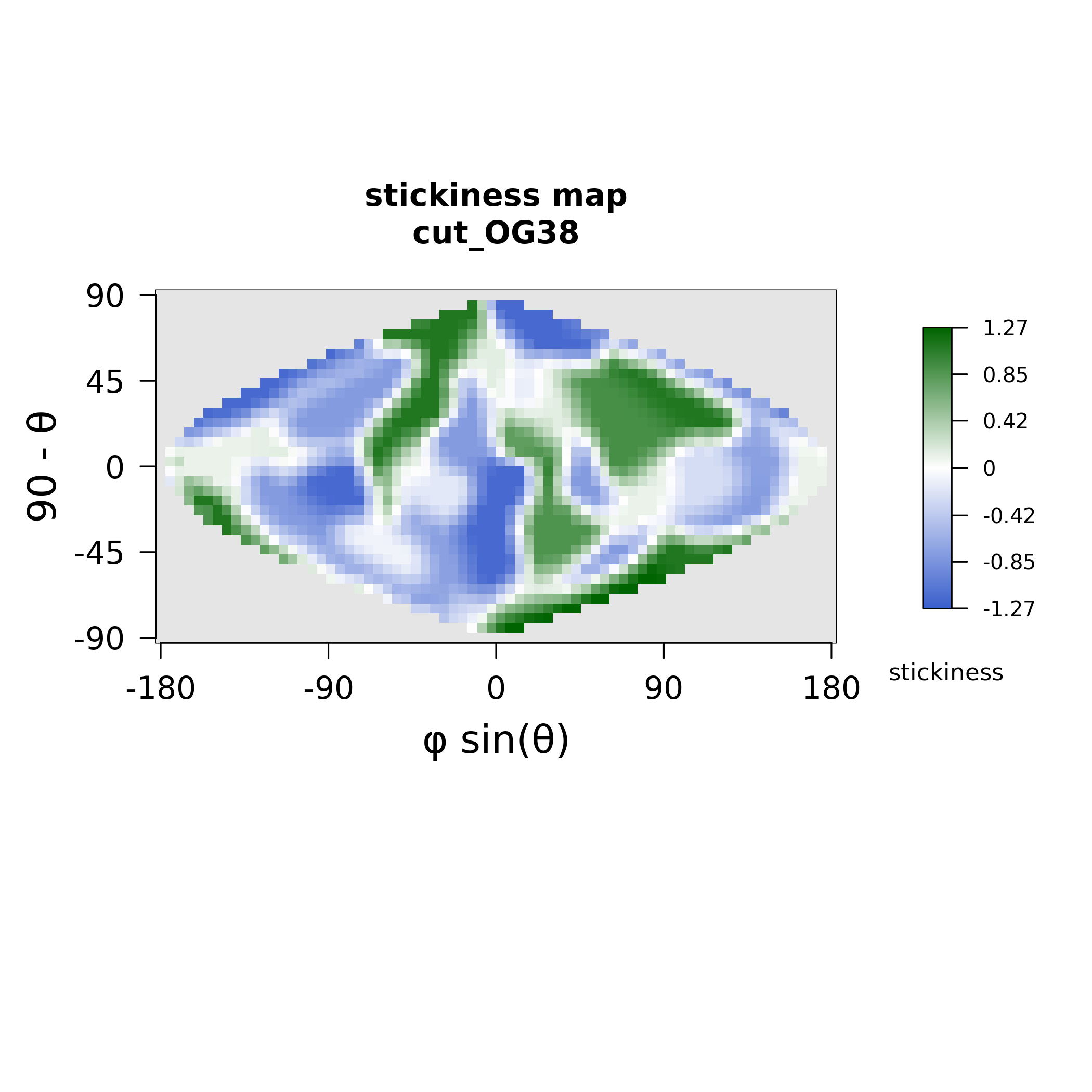

### MAX45_electrostatics.png

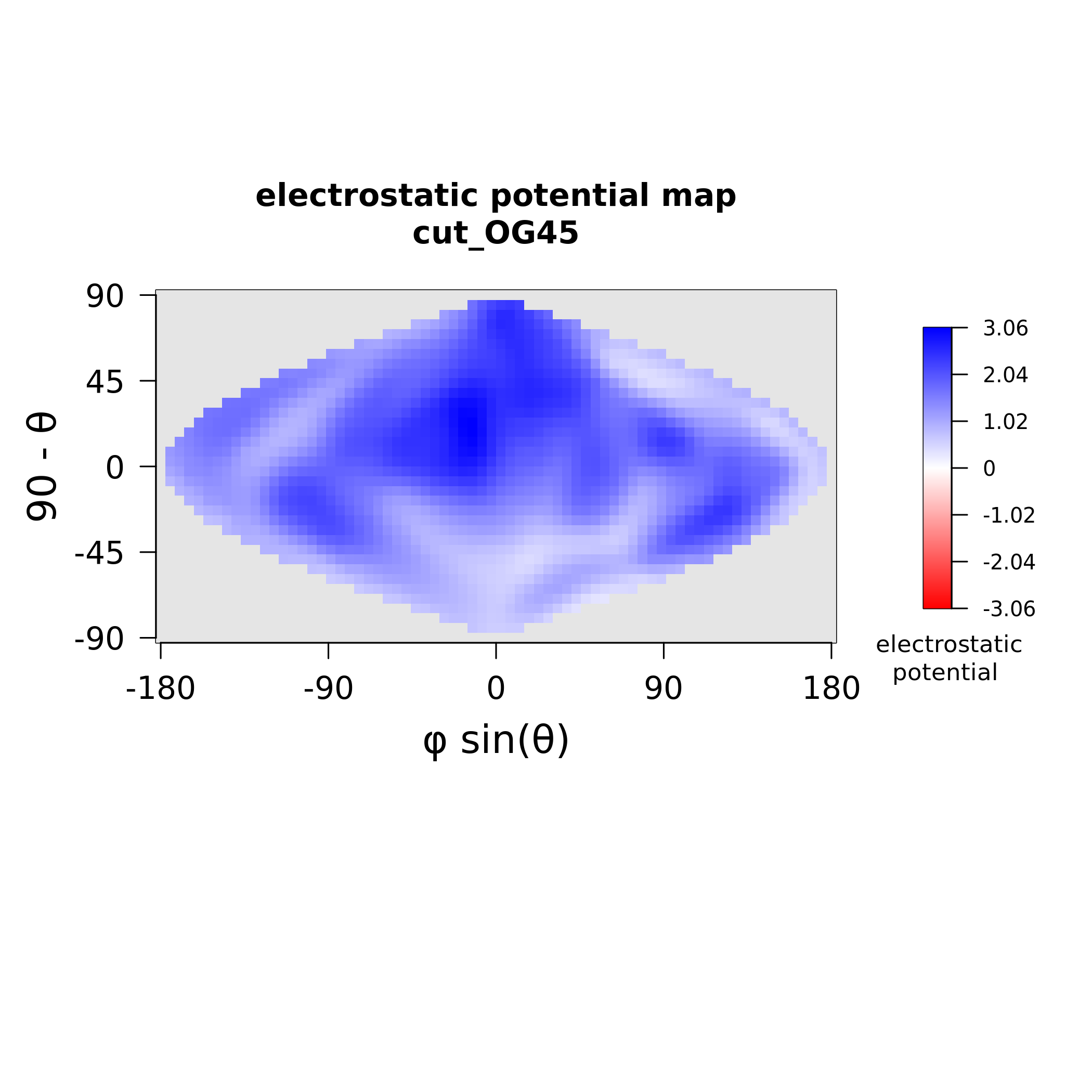

### MAX45_stickiness.png

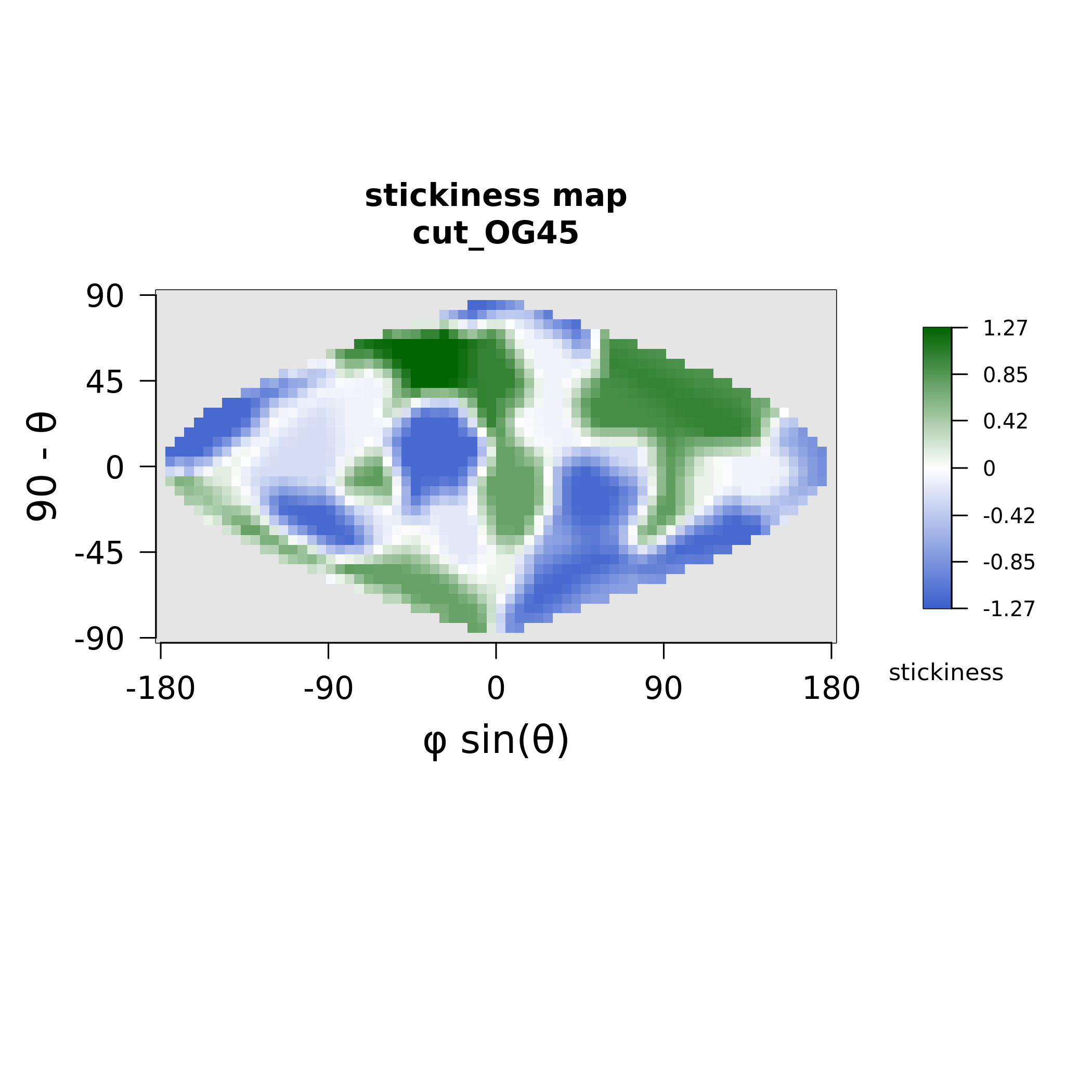

### MAX46_electrostatics.png

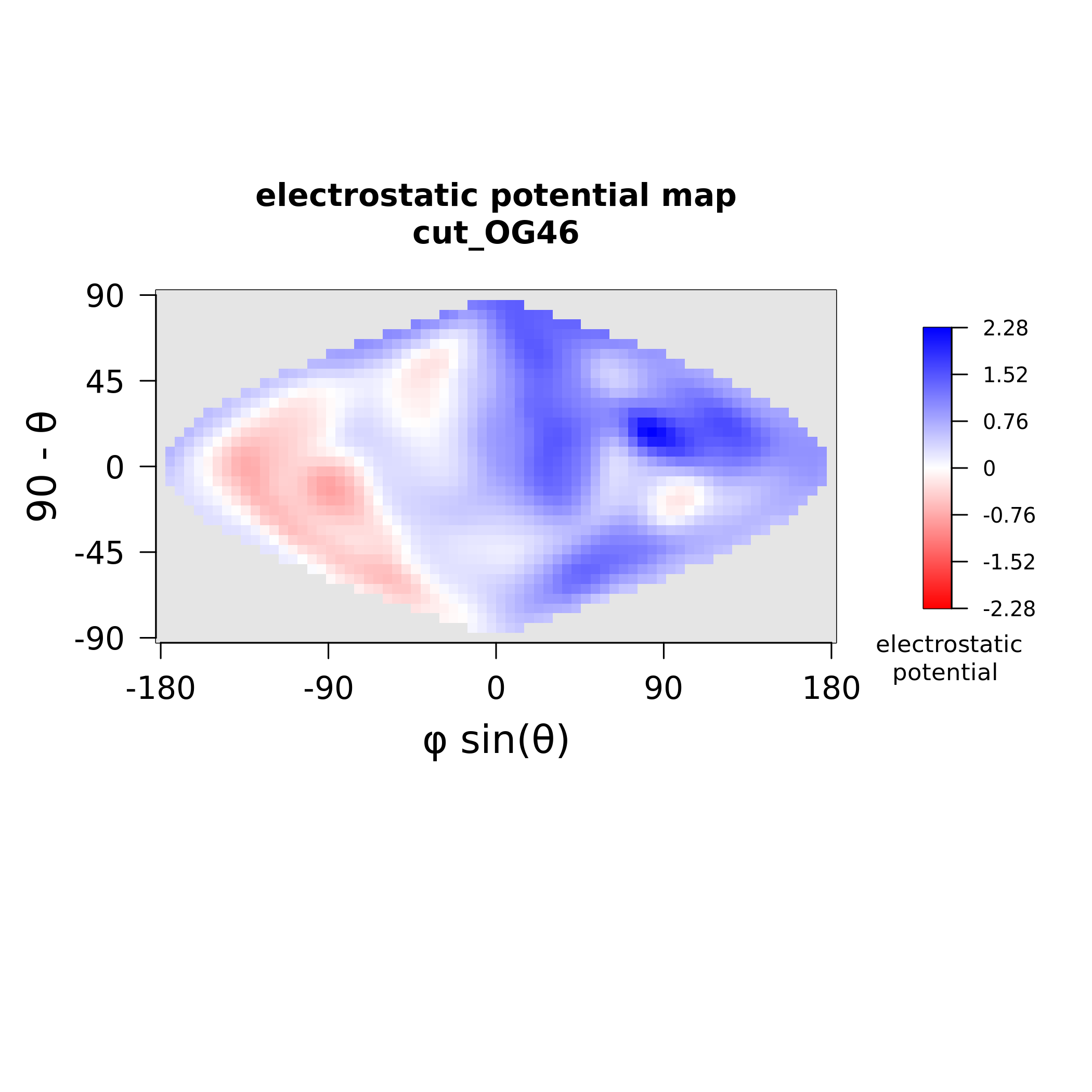

### MAX46_stickiness.png

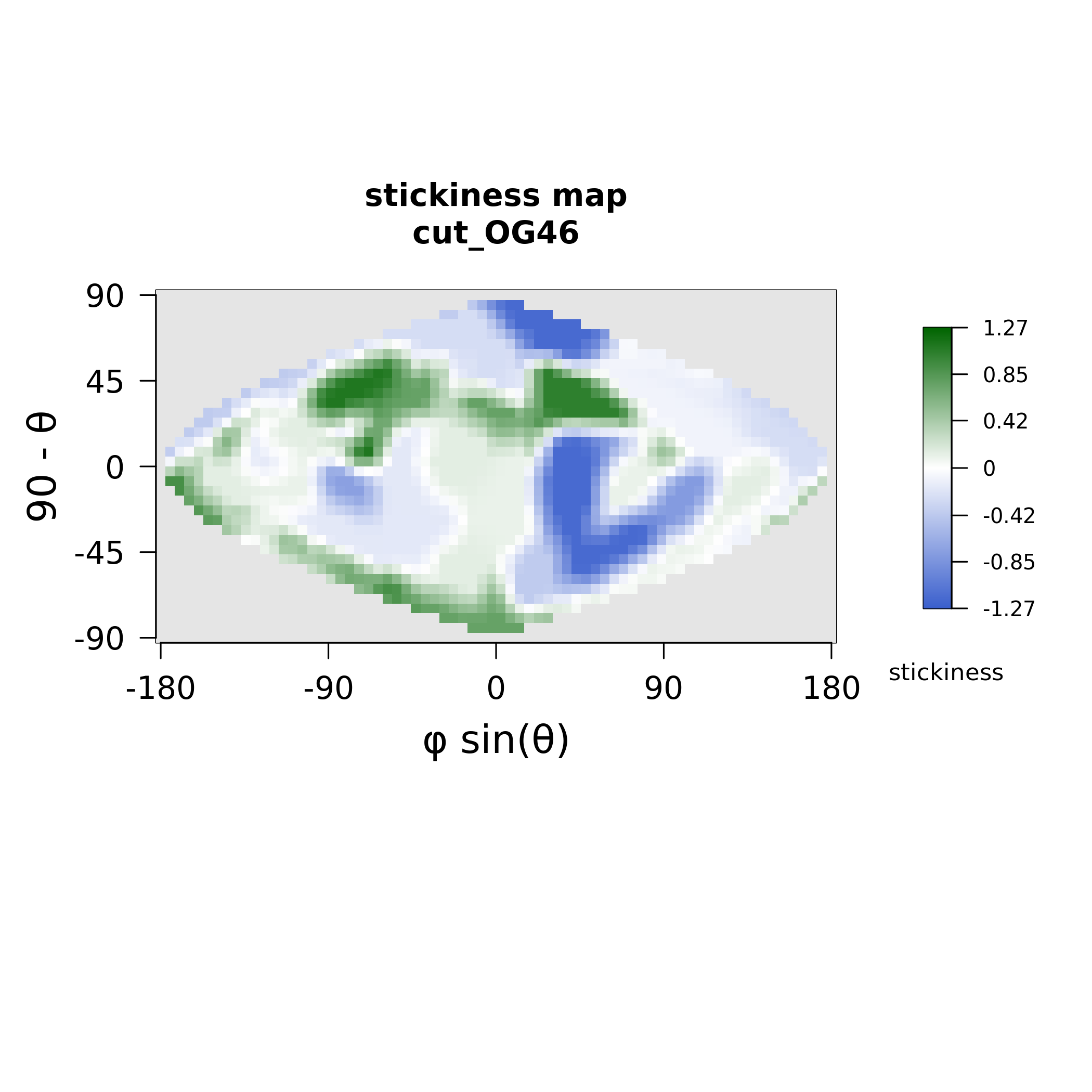

### MAX47_electrostatics.png

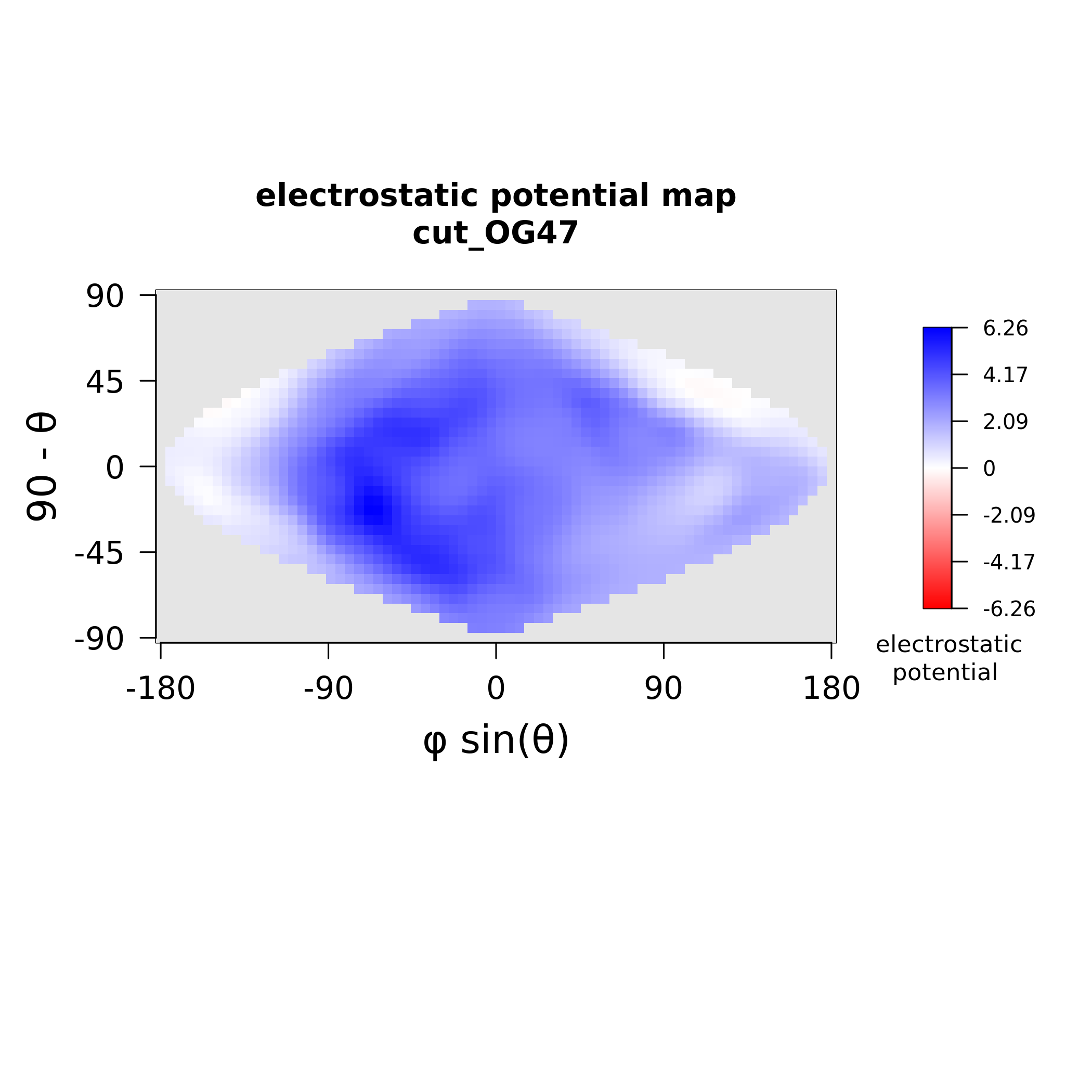

### MAX47_stickiness.png

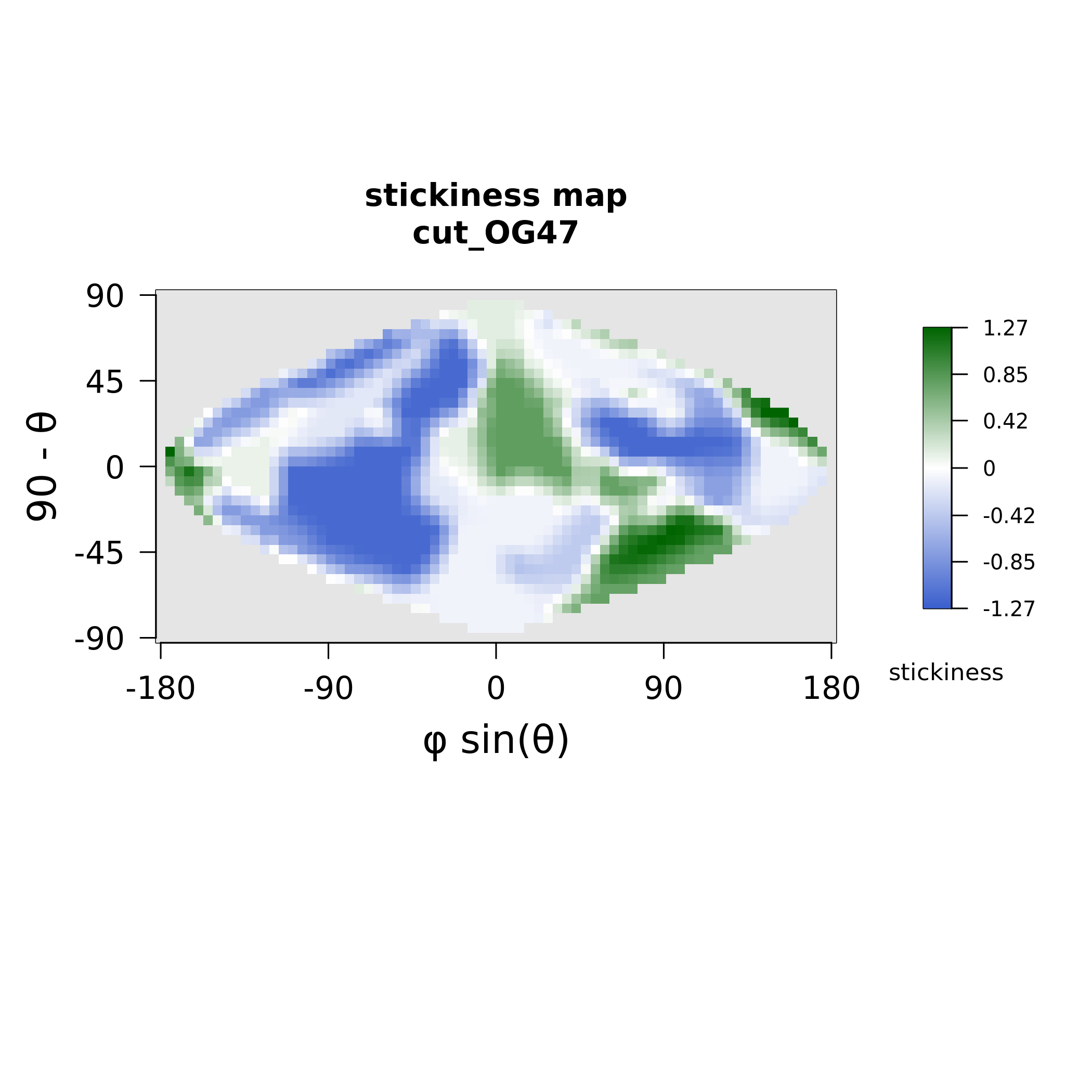

### MAX48_electrostatics.png

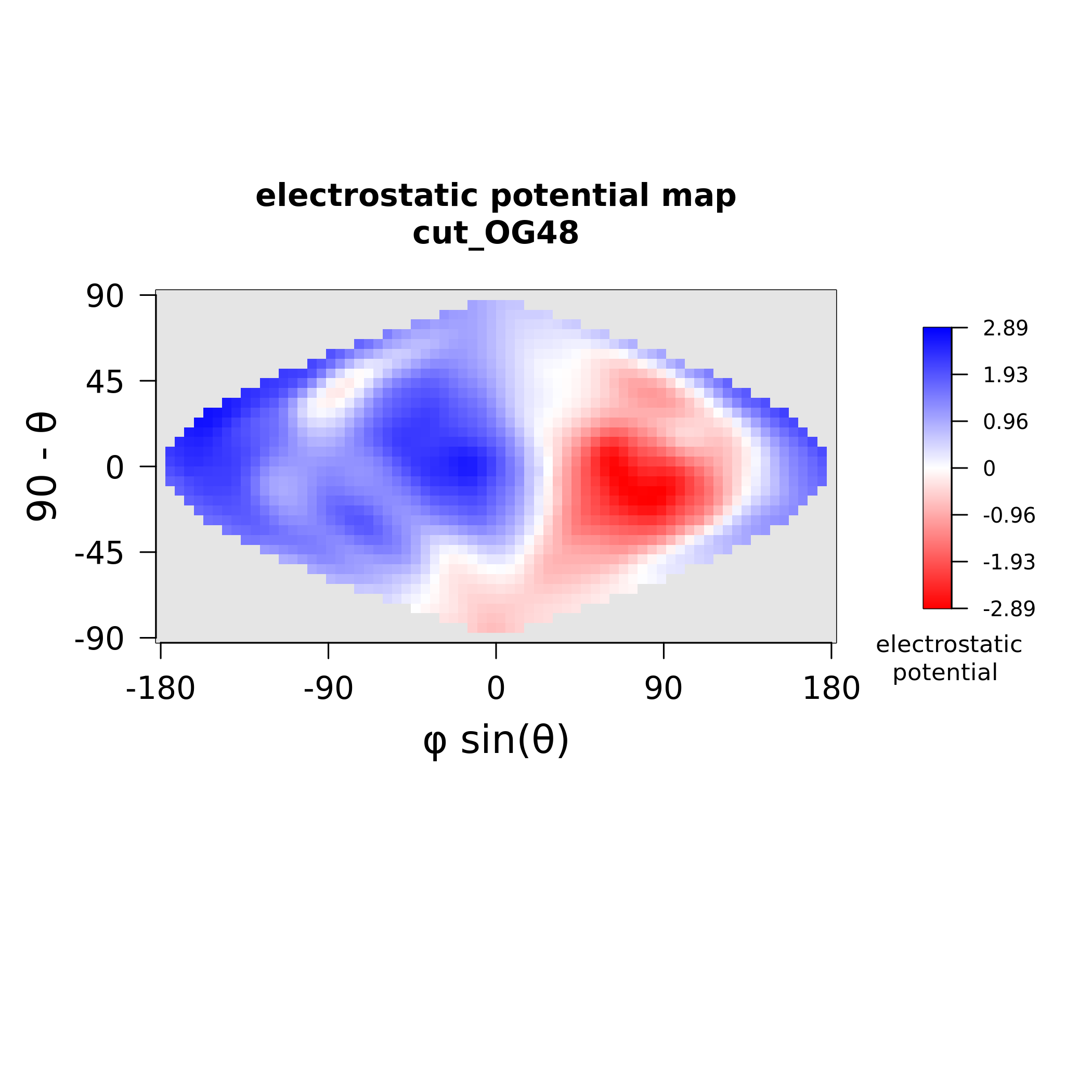

### MAX48_stickiness.png

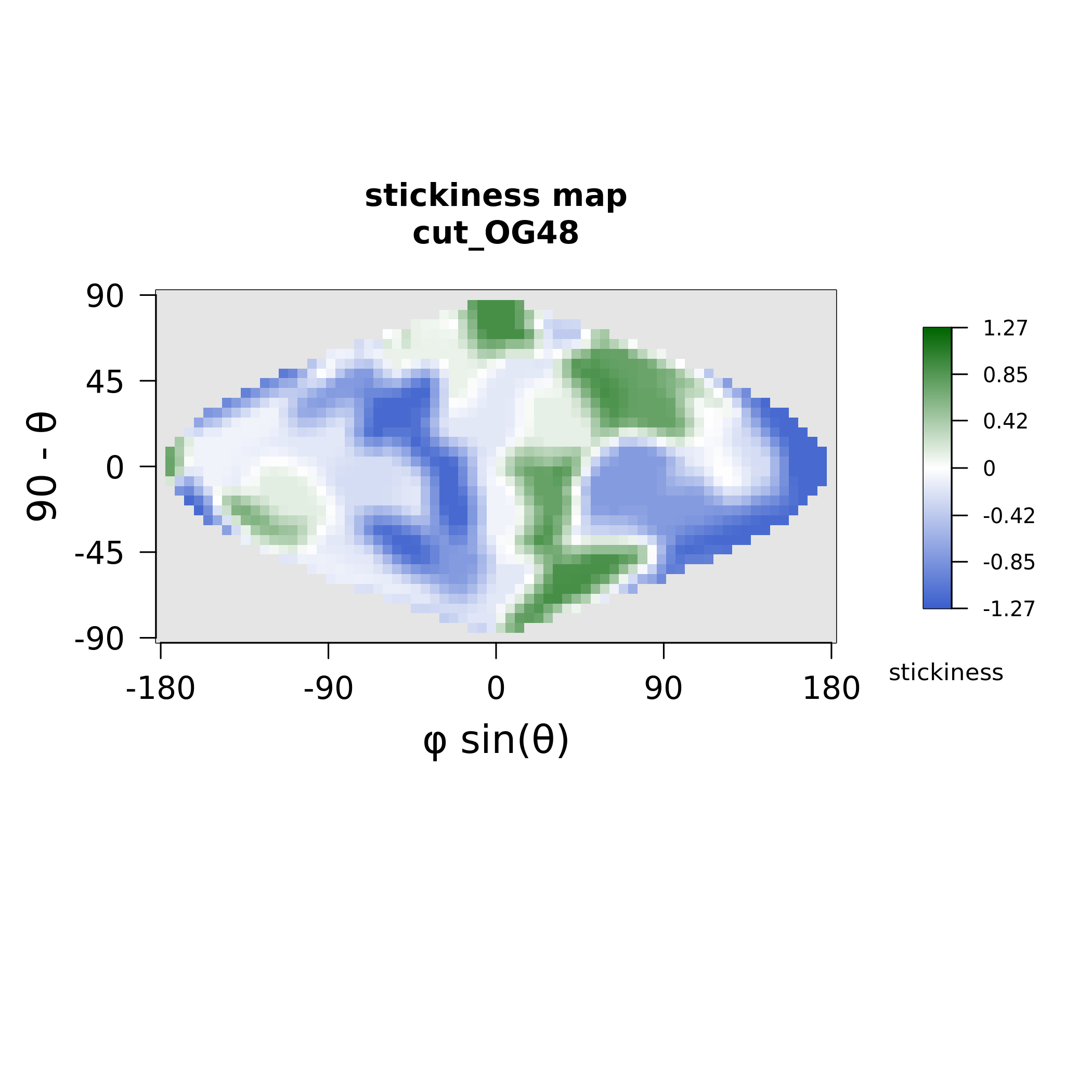

### MAX50_electrostatics.png

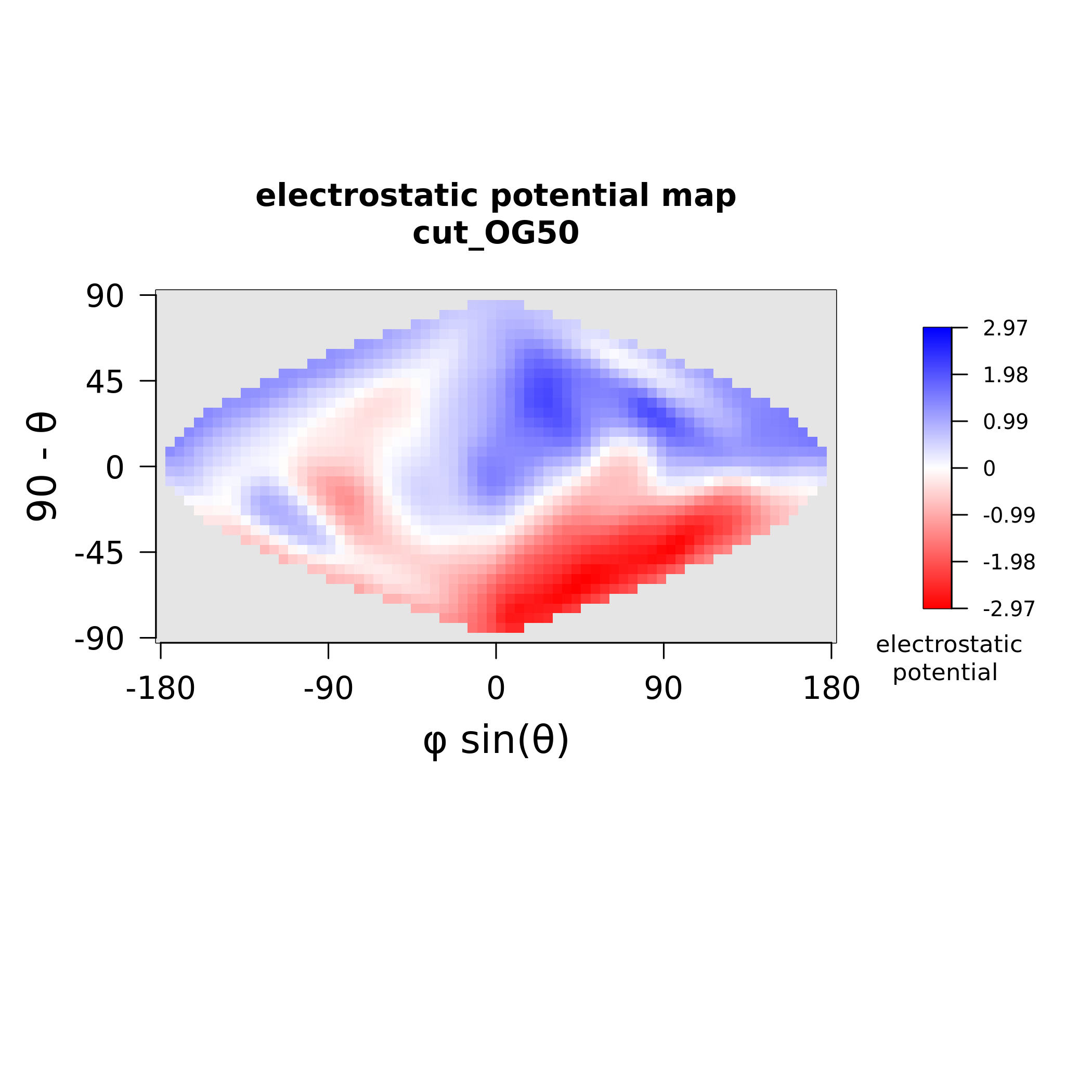

### MAX50_stickiness.png

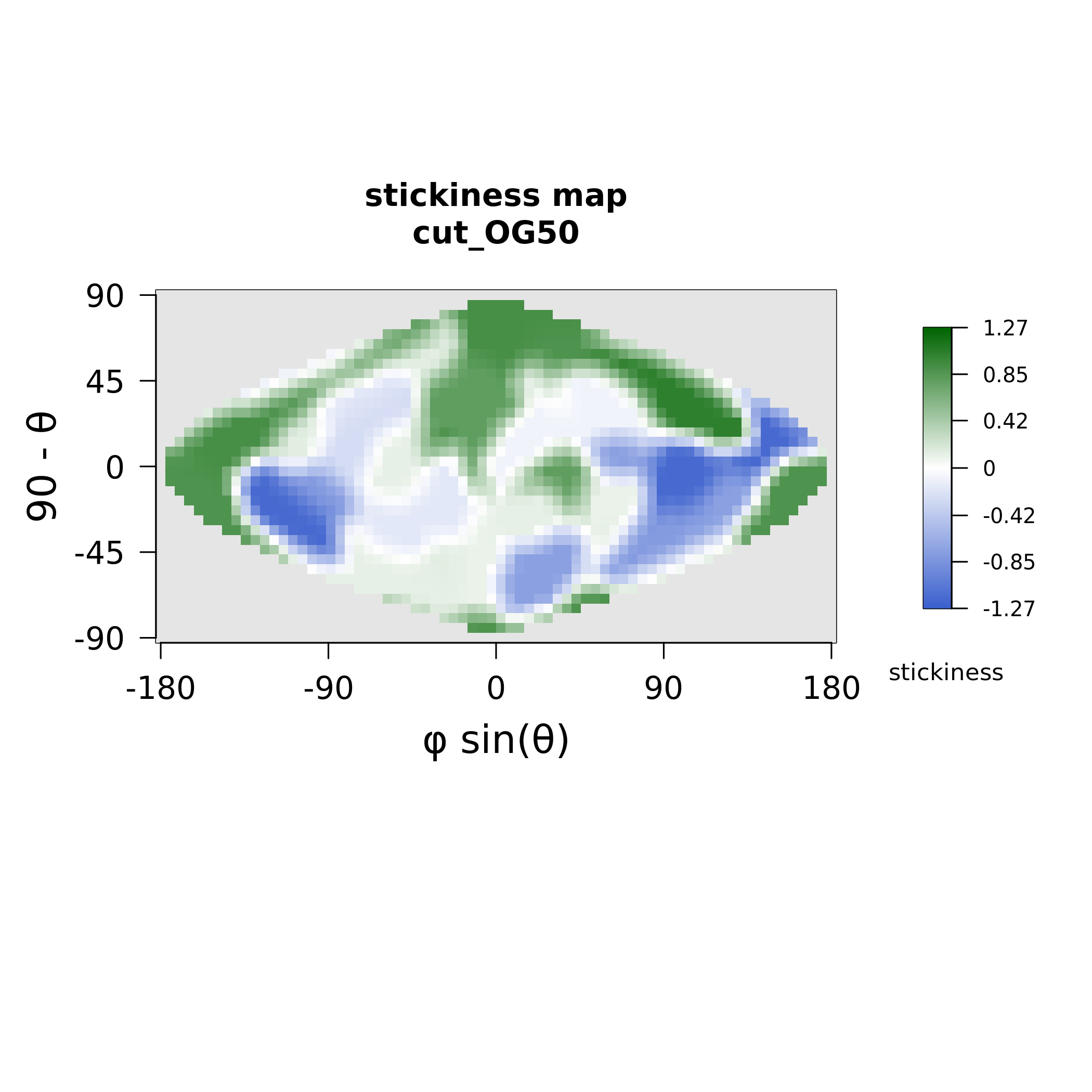

### MAX56_electrostatics.png

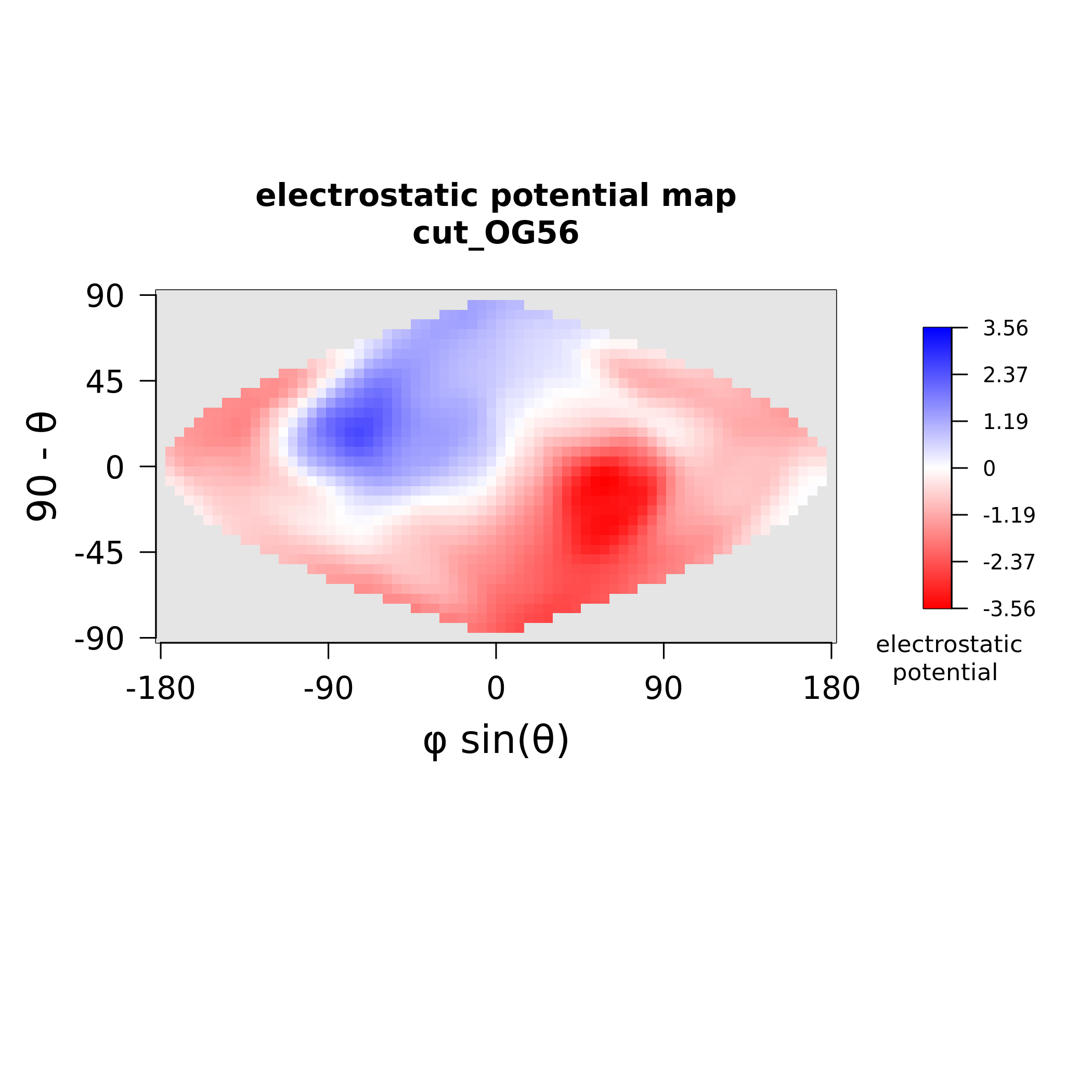

### MAX56_stickiness.png

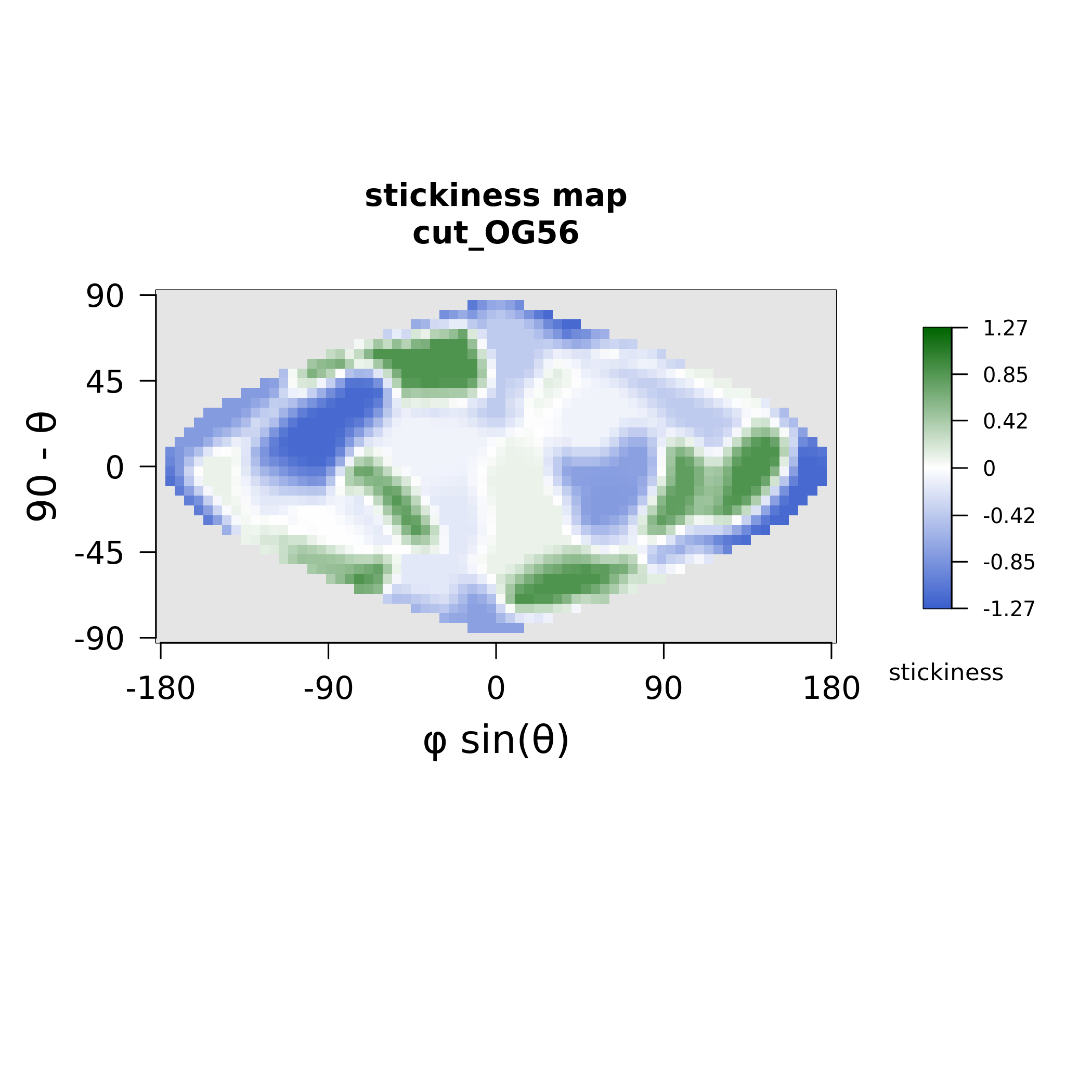

### MAX58_electrostatics.png

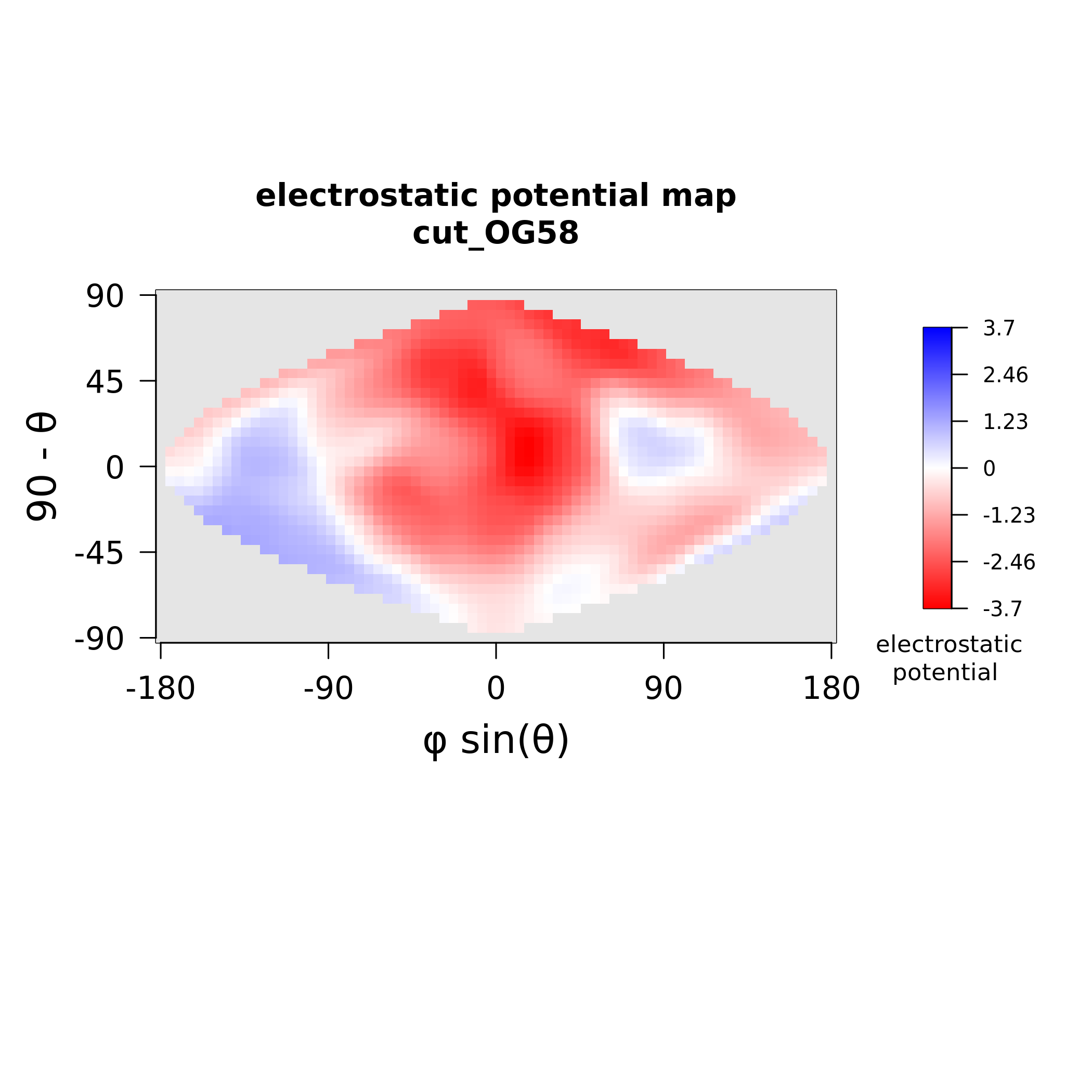

### MAX58_polymorphism.png

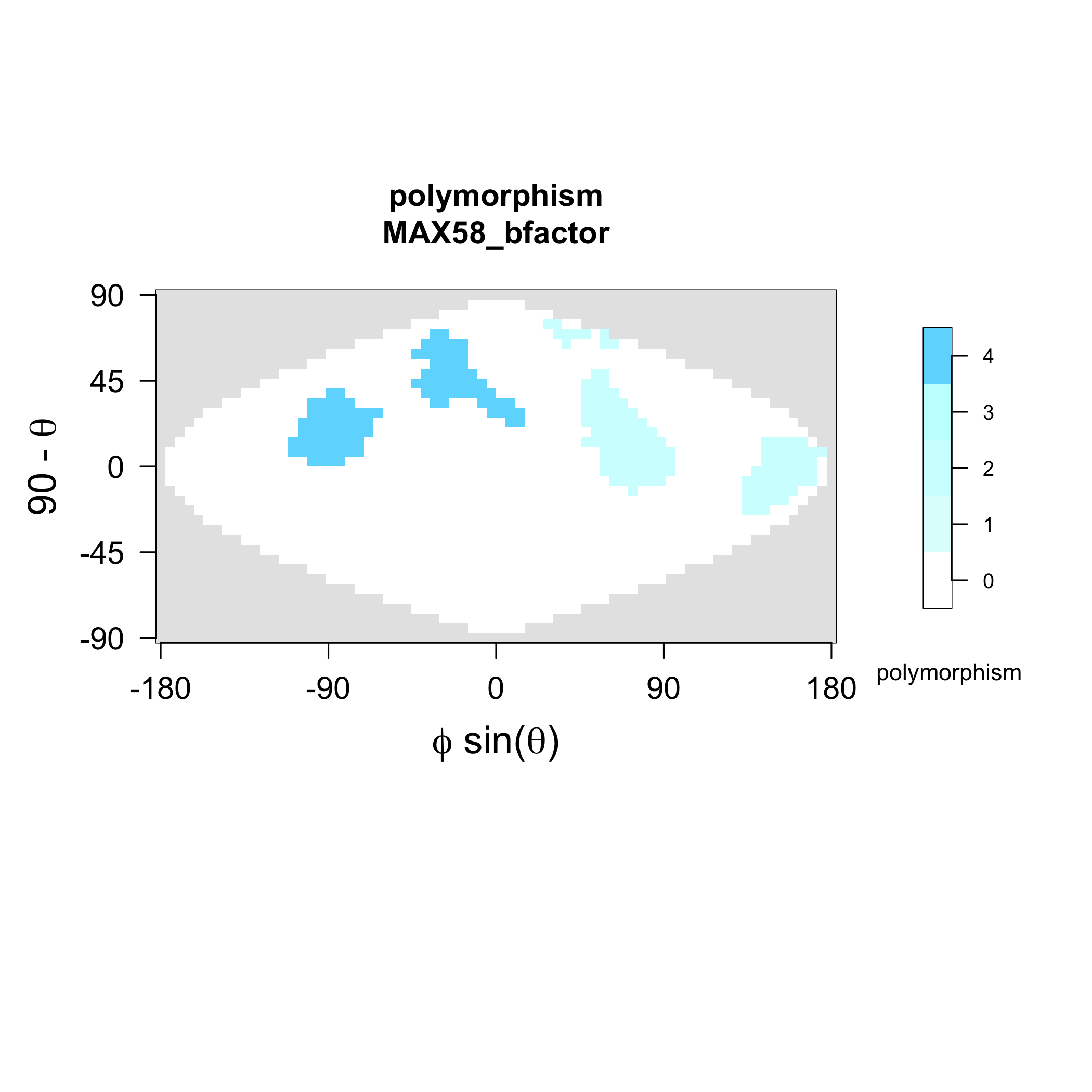

### MAX58_stickiness.png

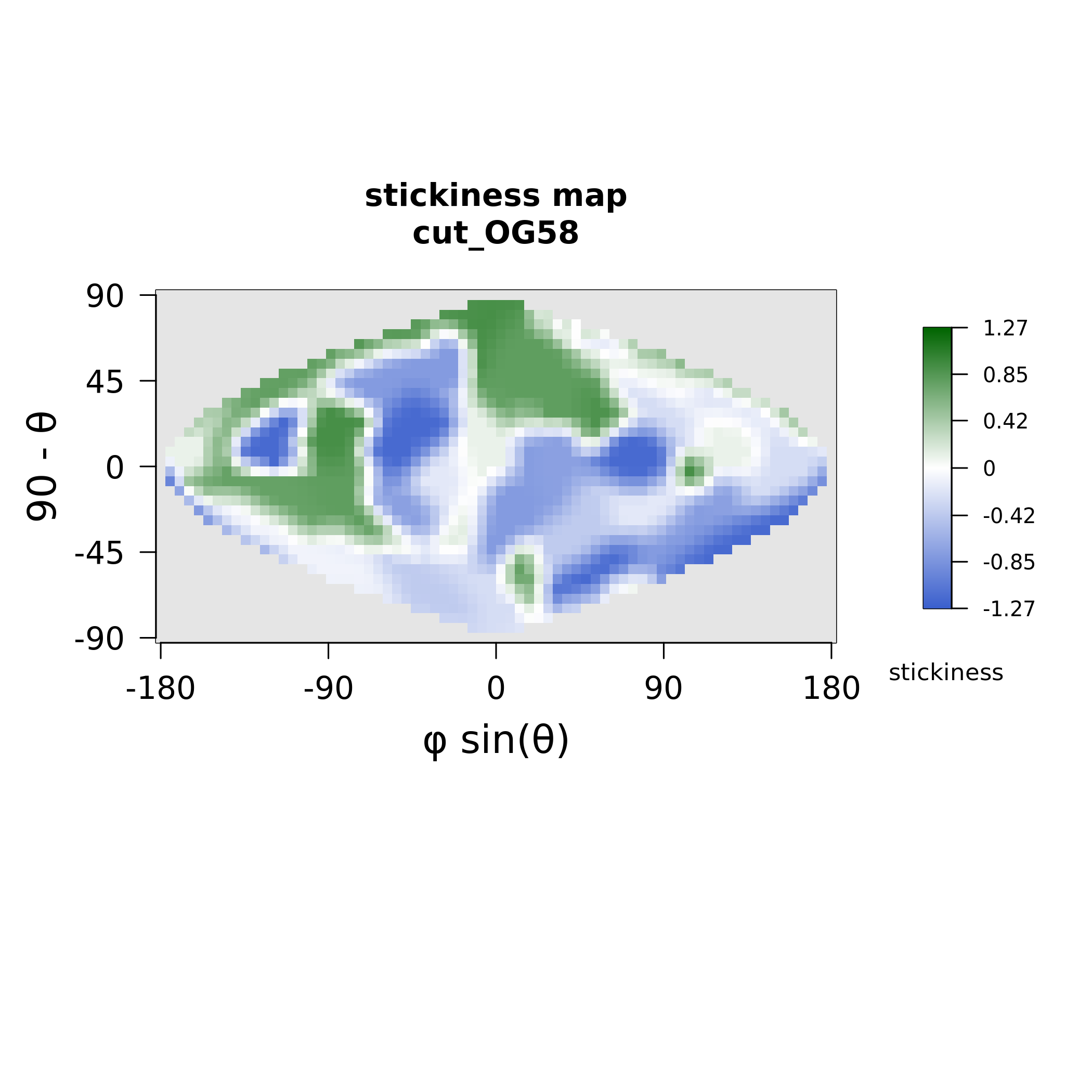

### MAX61_electrostatics.png

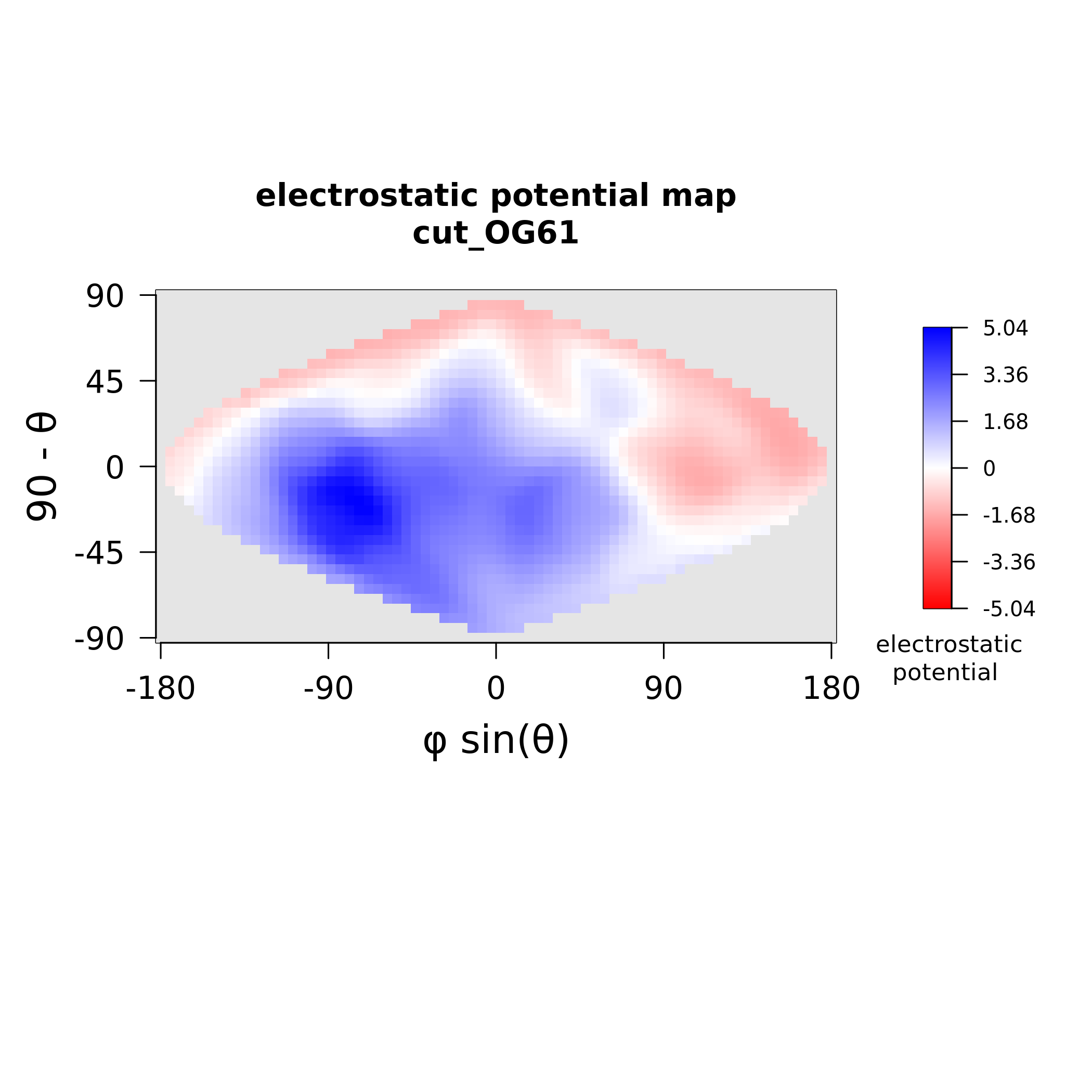

### MAX61_stickiness.png

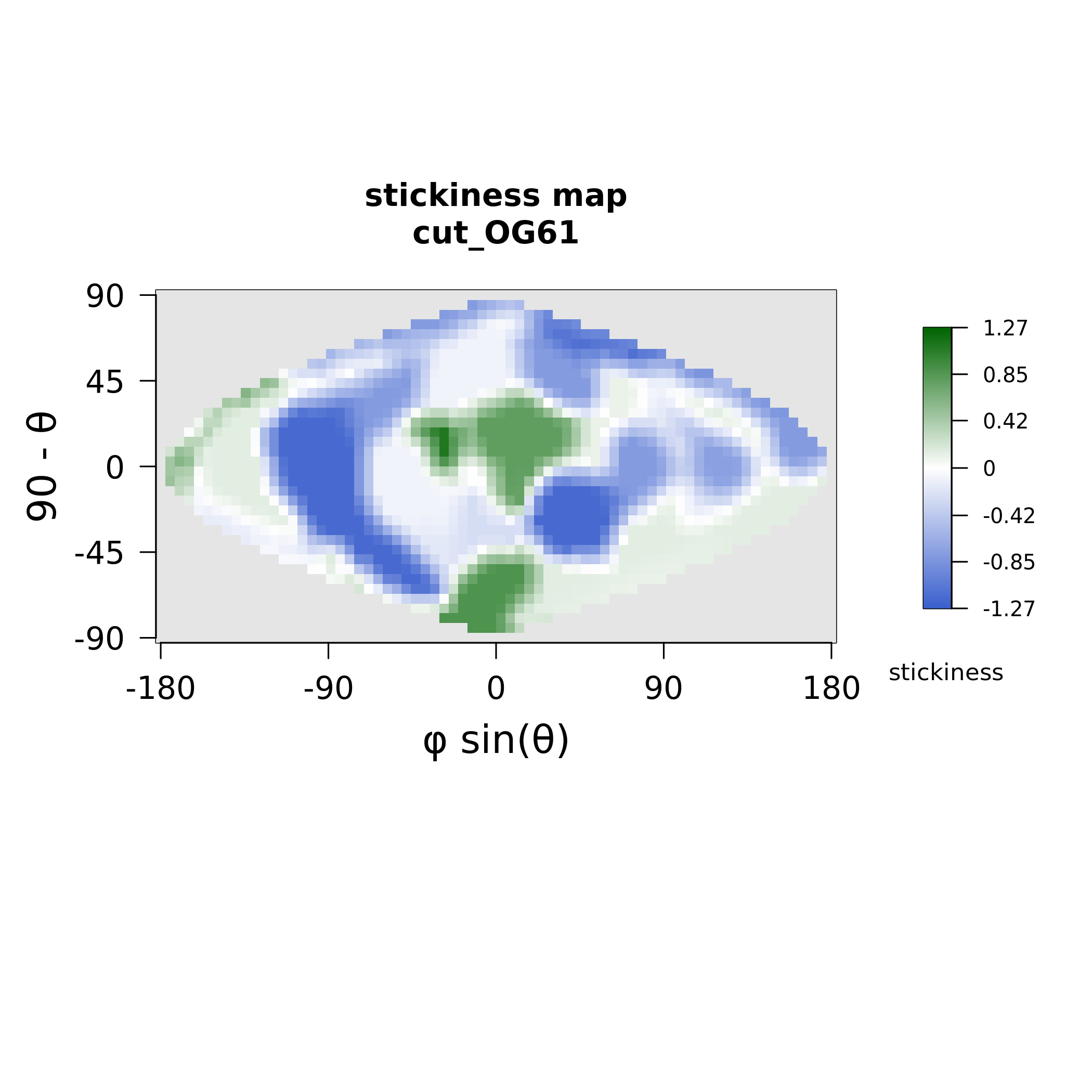

### MAX62_electrostatics.png

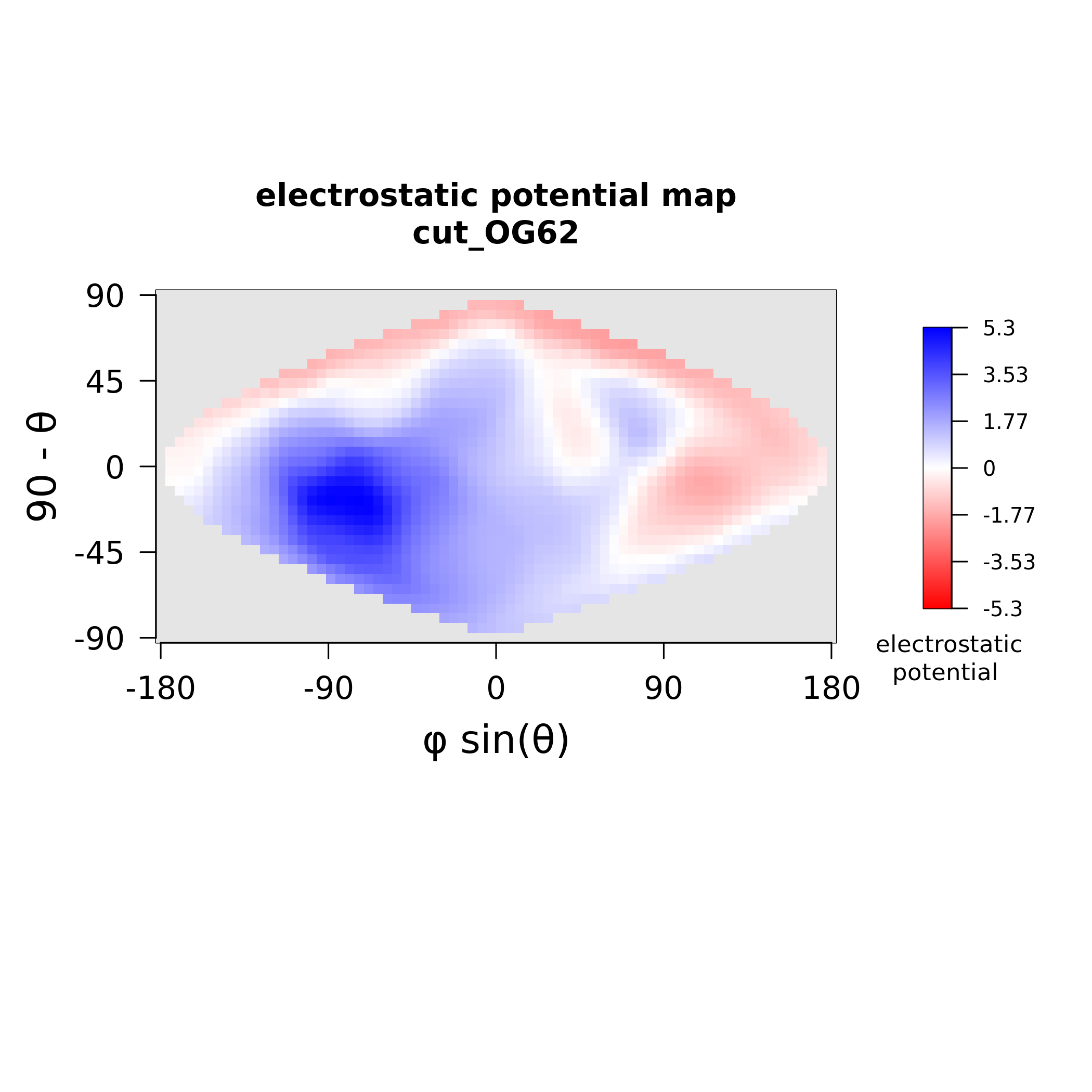

### MAX62_stickiness.png

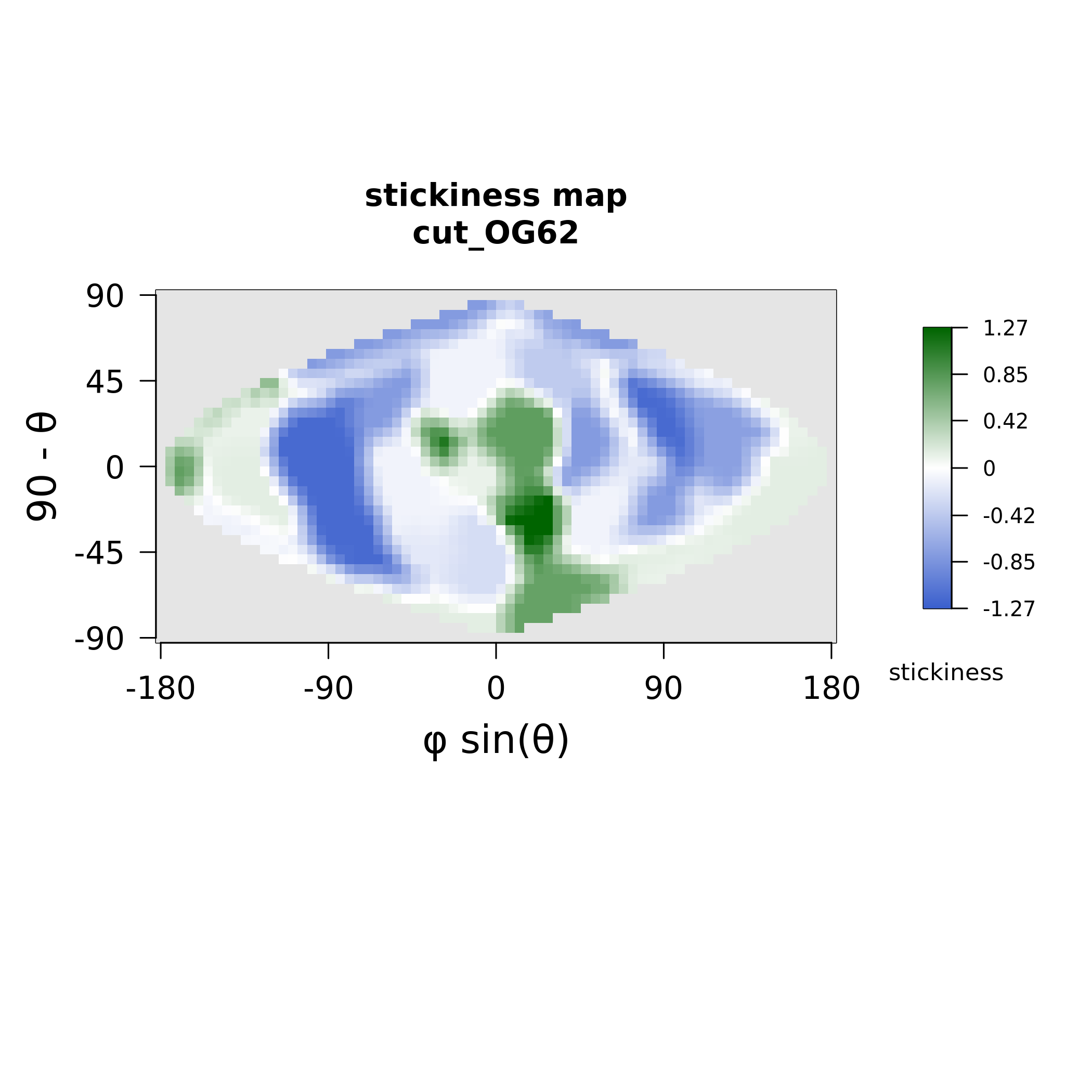

### MAX63_electrostatics.png

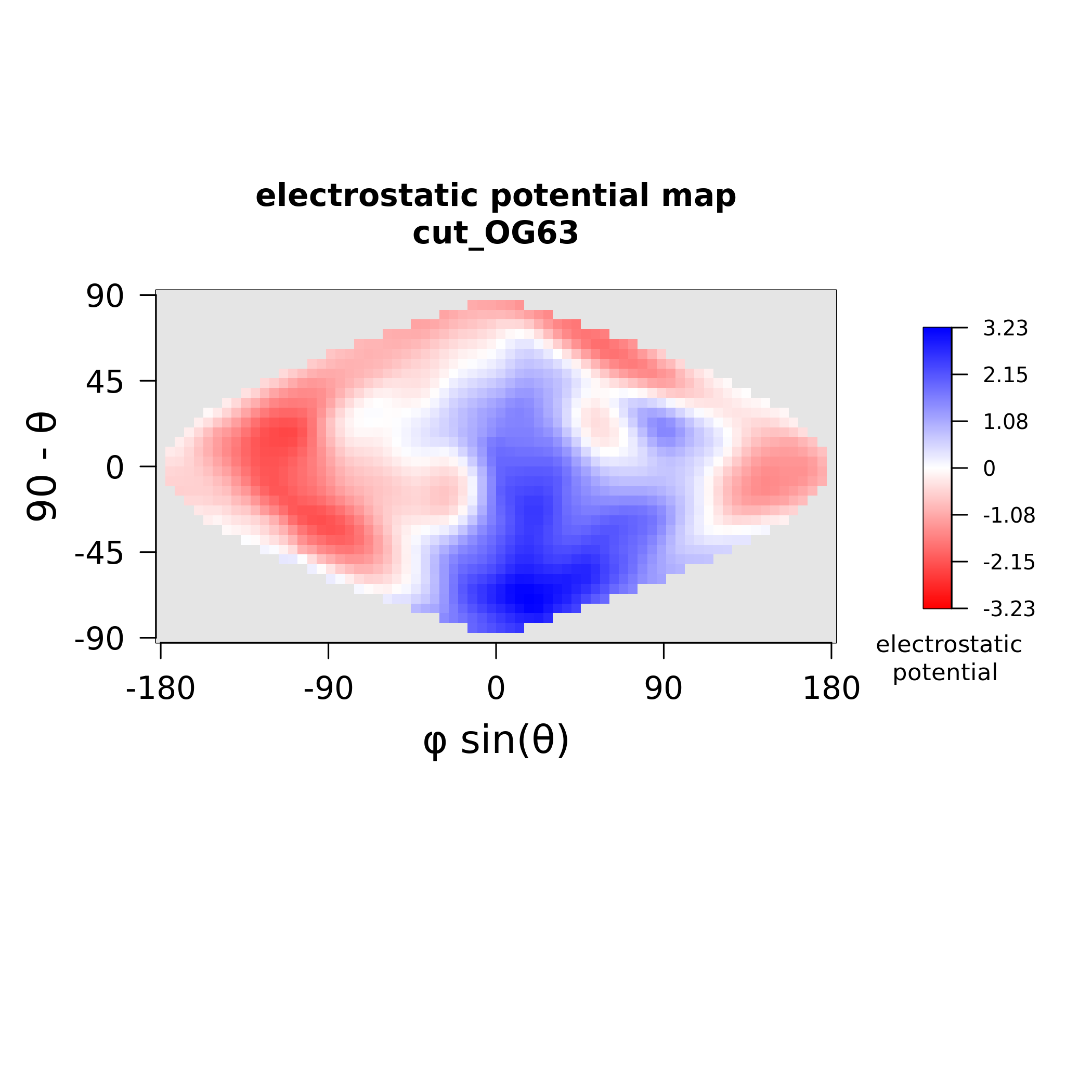

### MAX63_stickiness.png

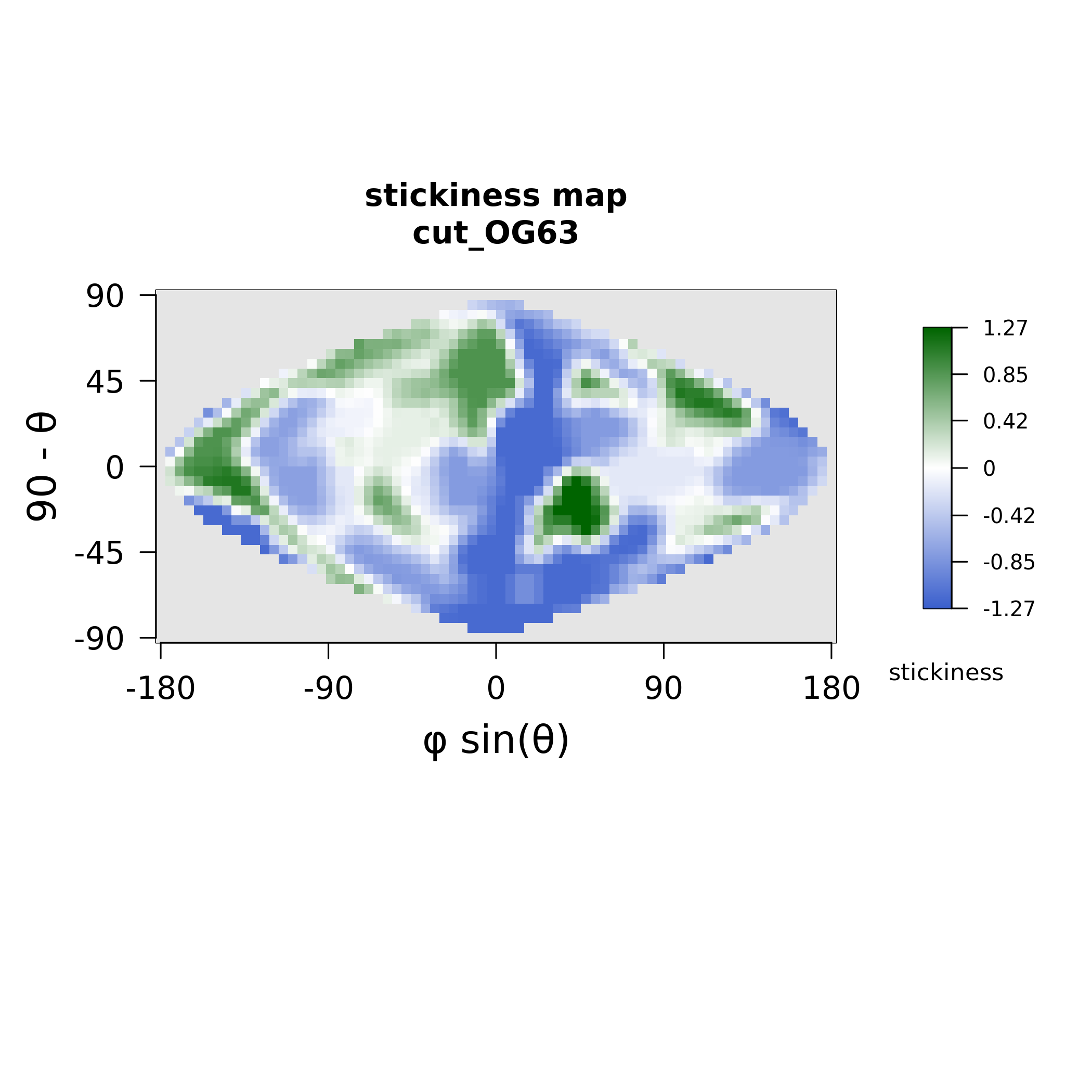

### MAX67_electrostatics.png

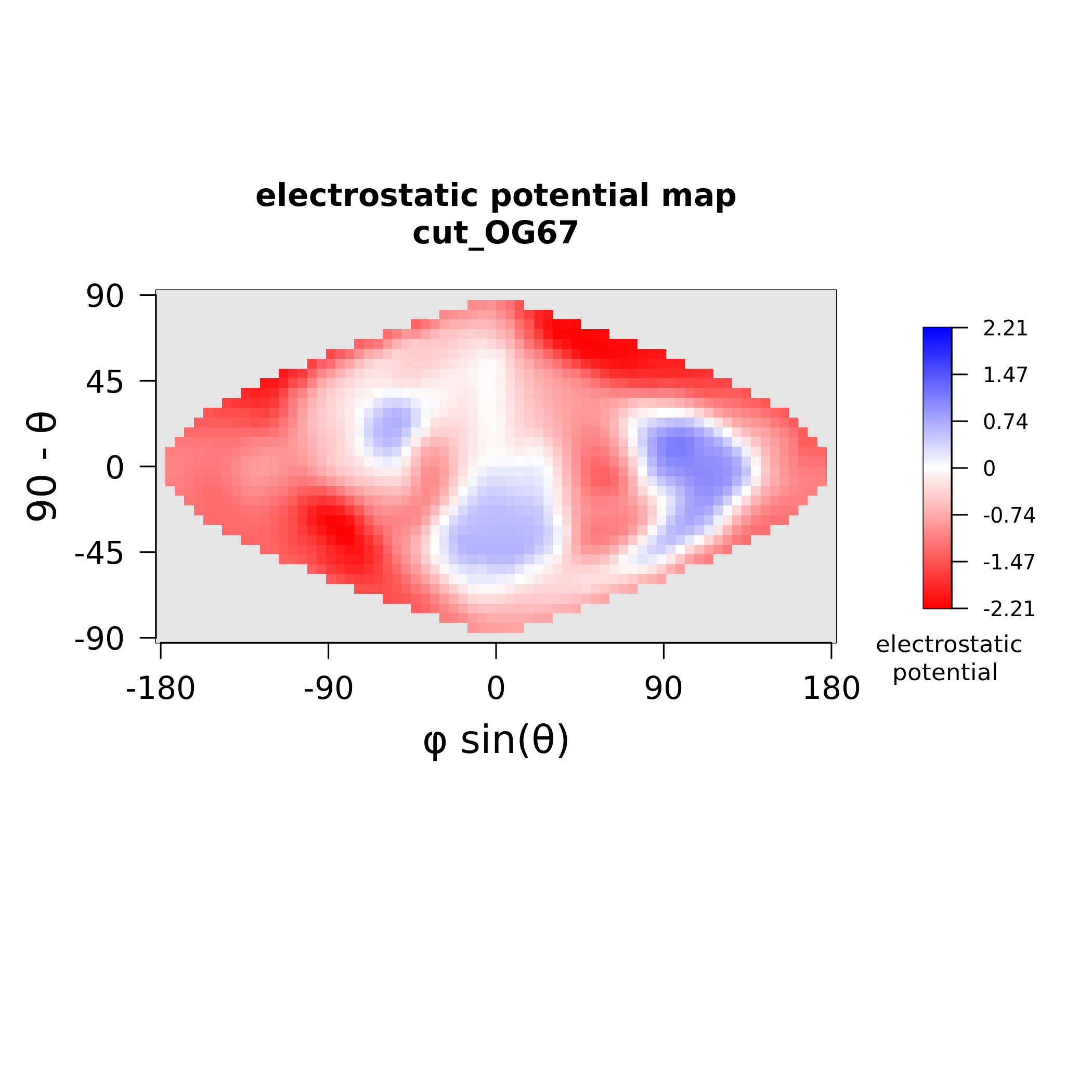

### MAX67_stickiness.png

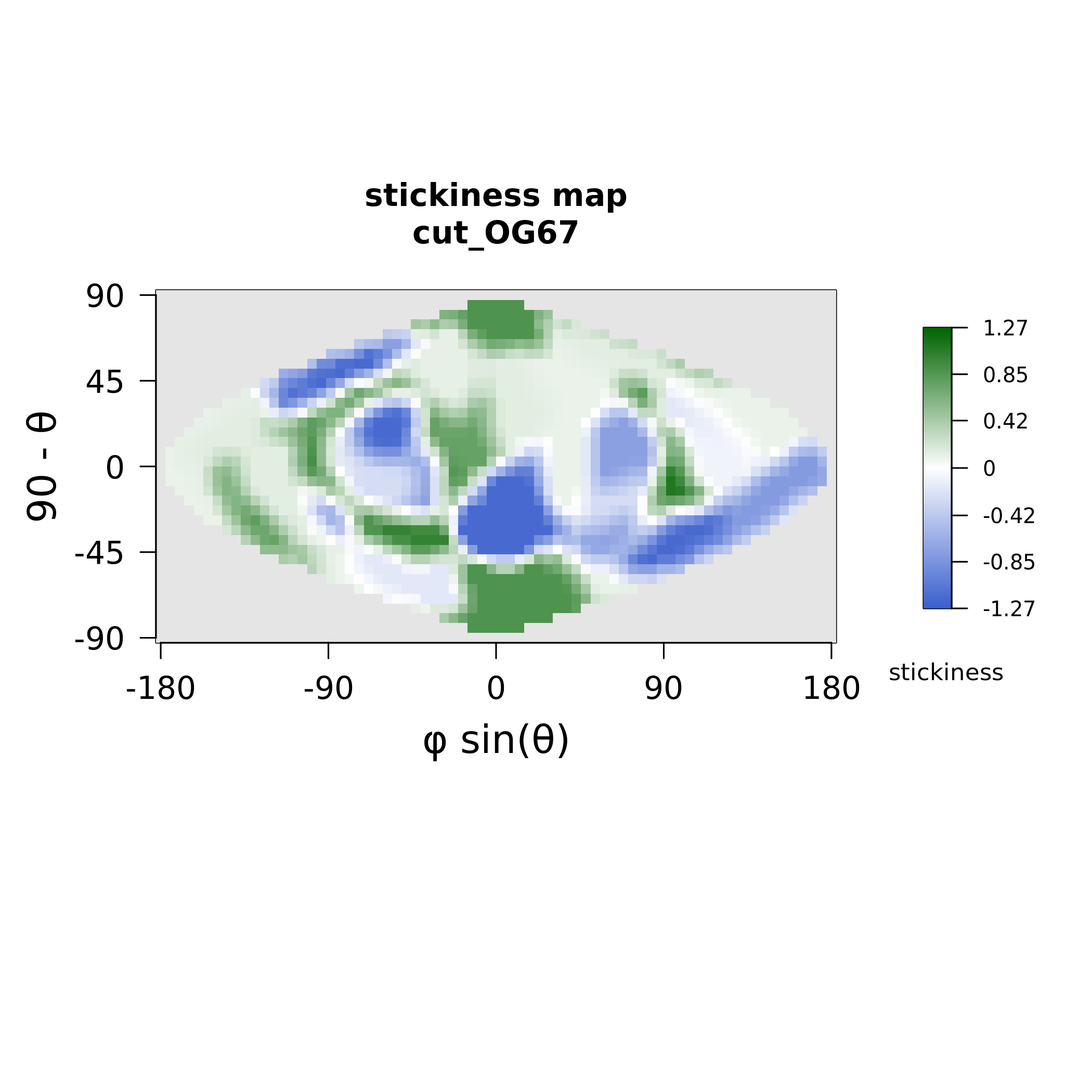

### MAX69_electrostatics.png

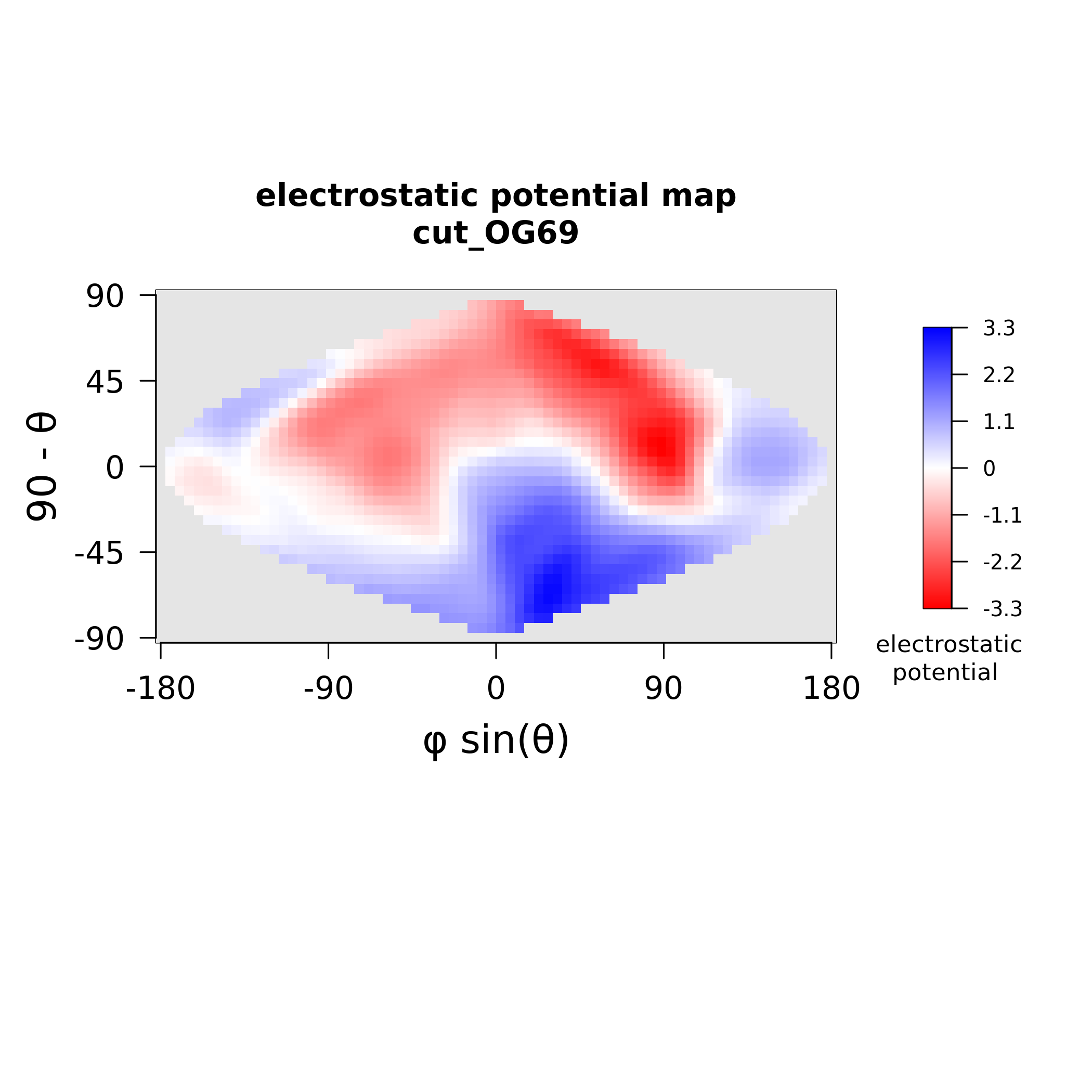

### MAX69_stickiness.png

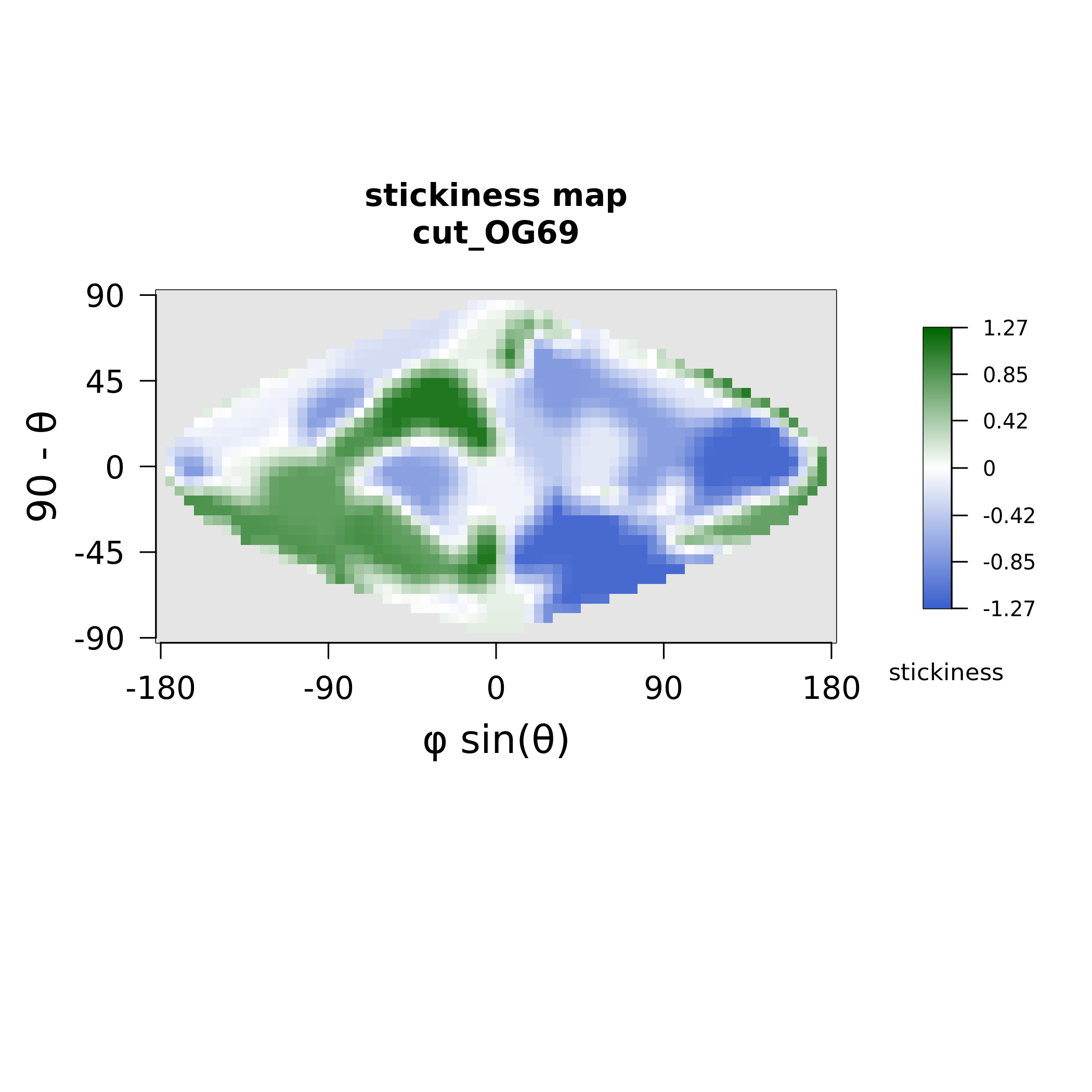

### MAX70_electrostatics.png

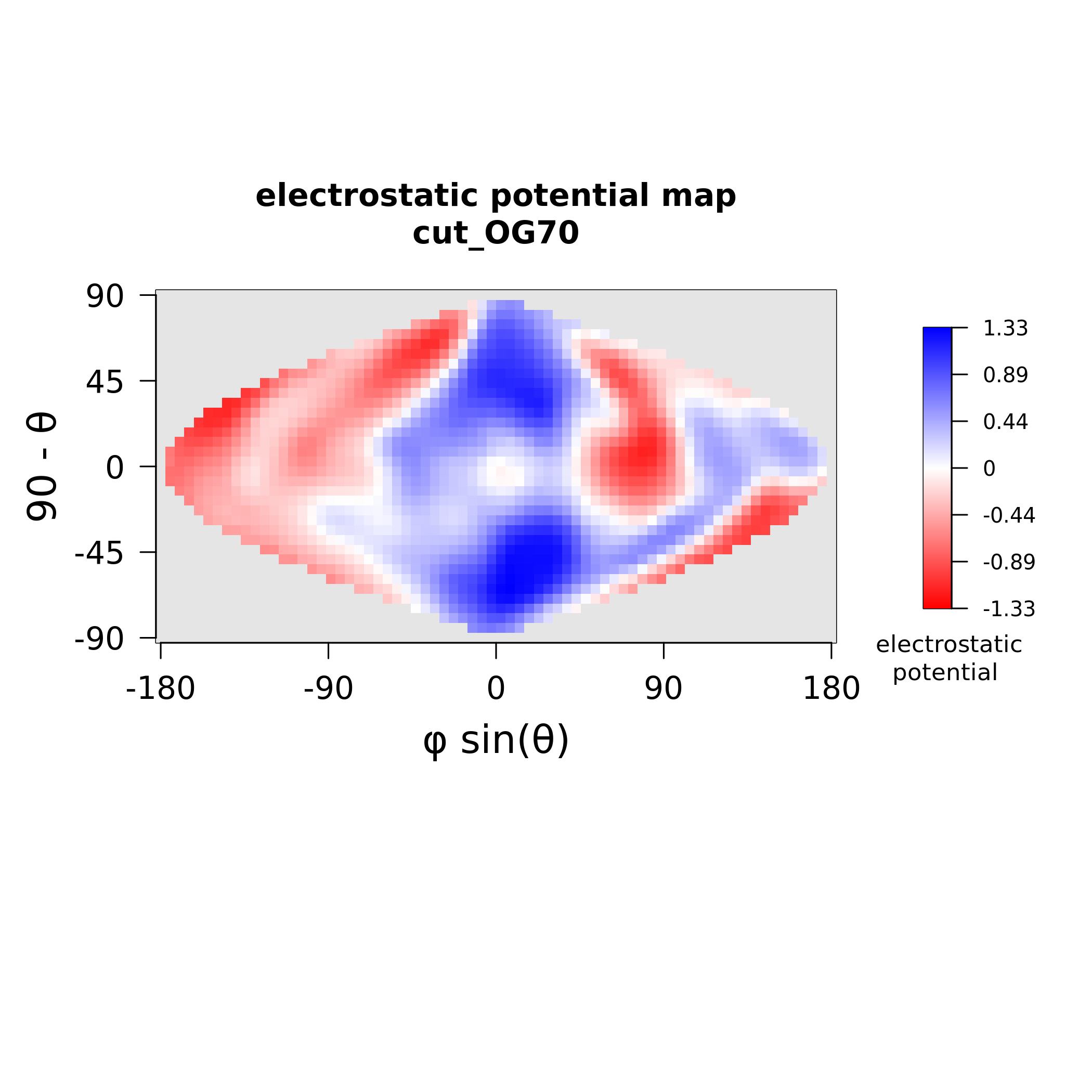

### MAX70_stickiness.png

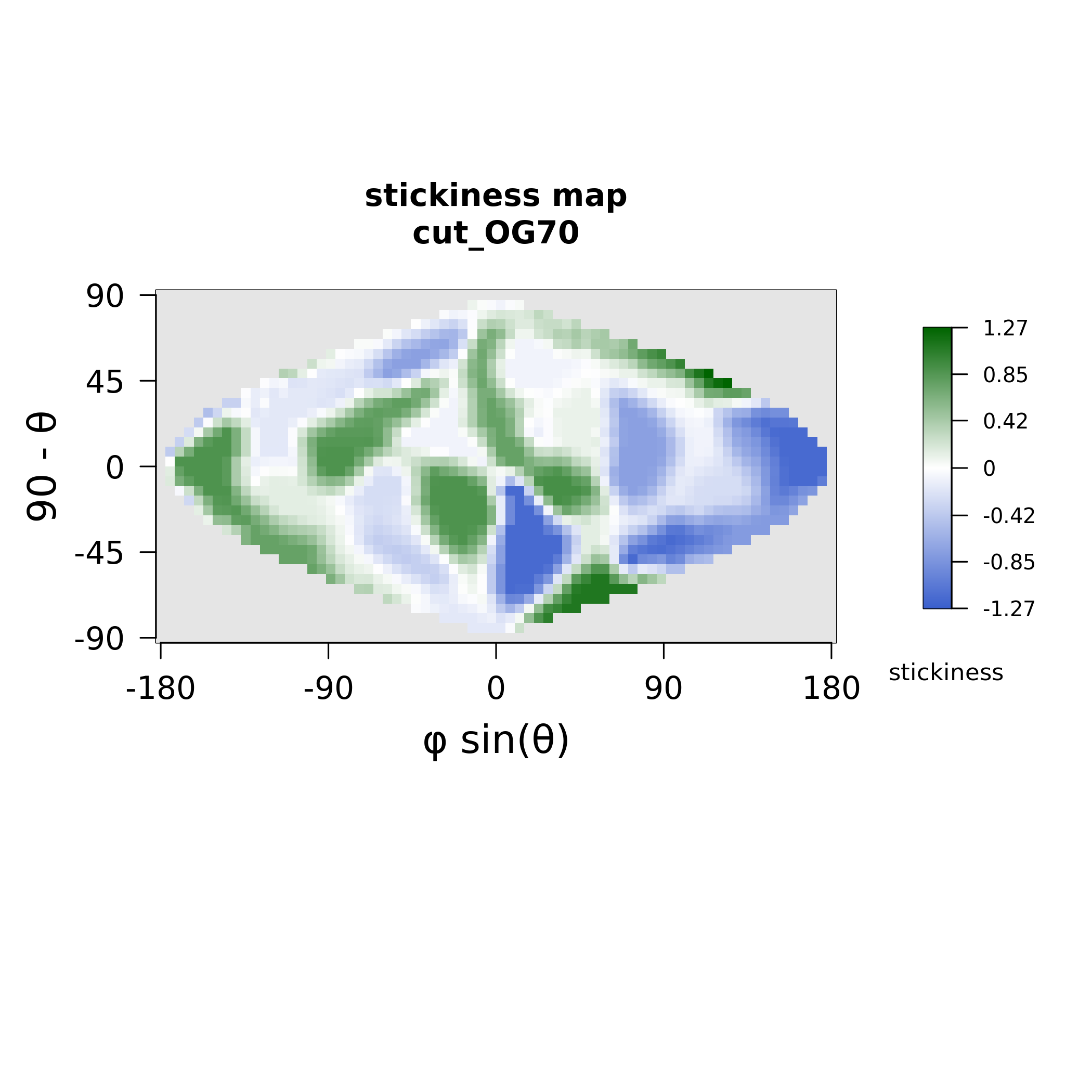

### MAX72_electrostatics.png

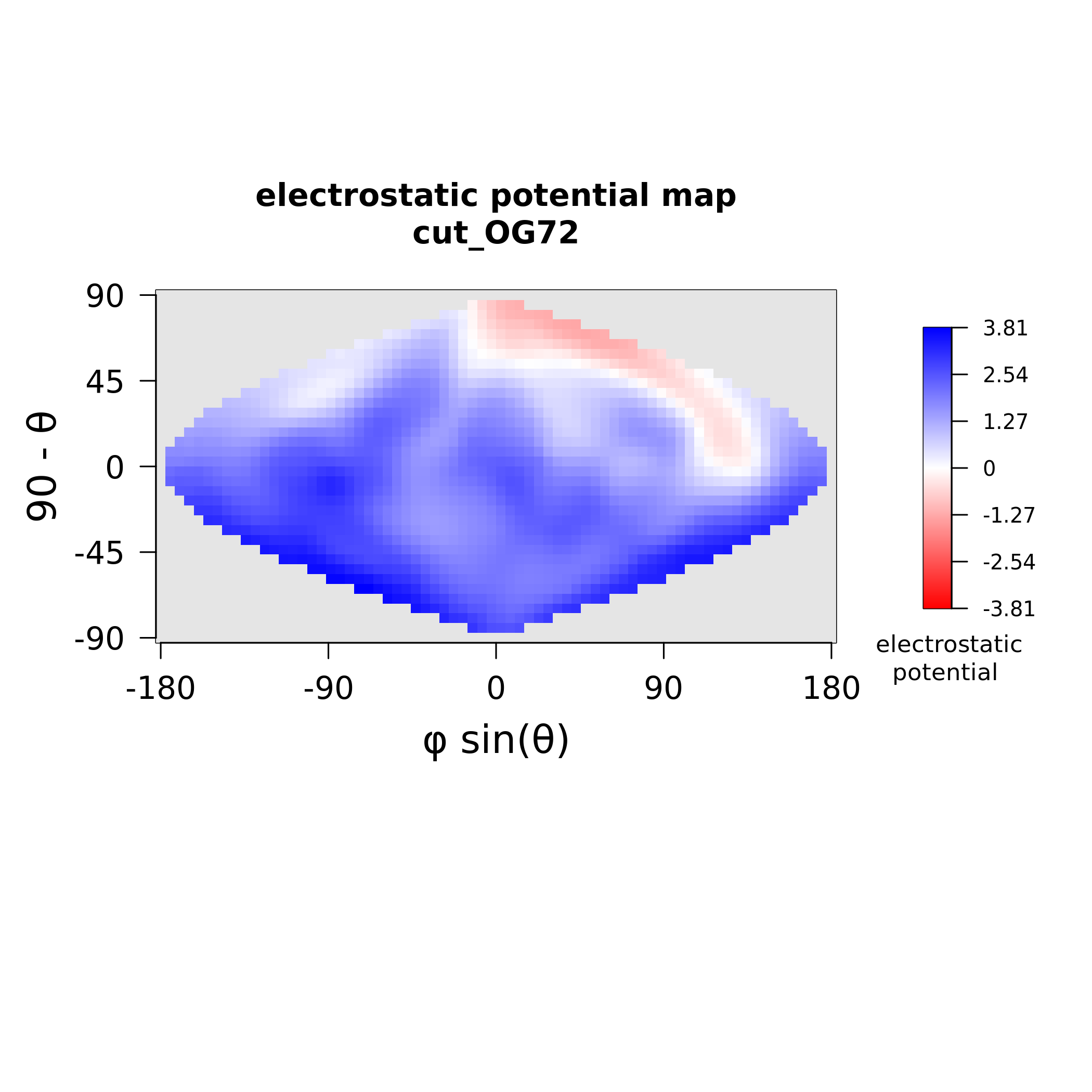

### MAX72_stickiness.png

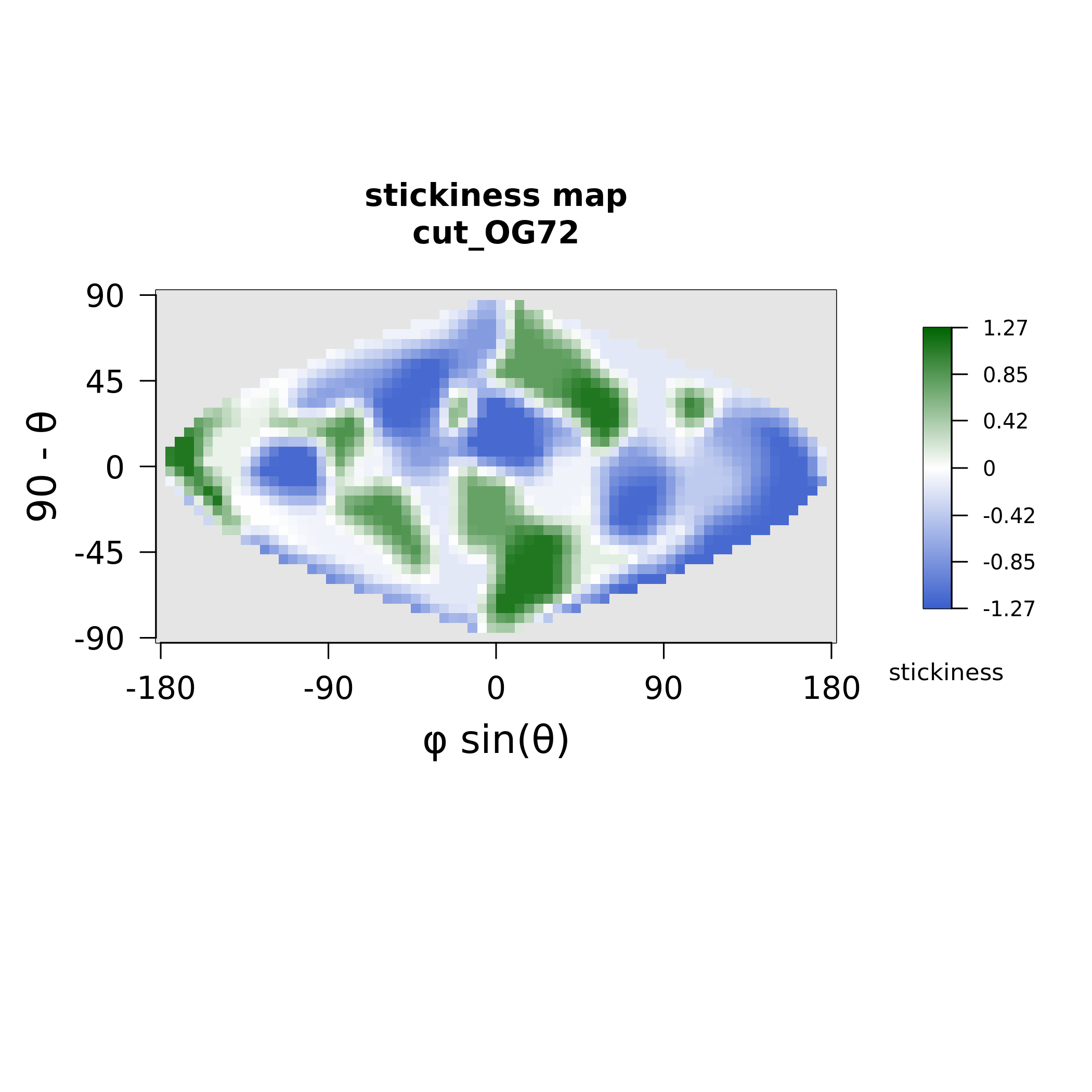

### MAX76_electrostatics.png

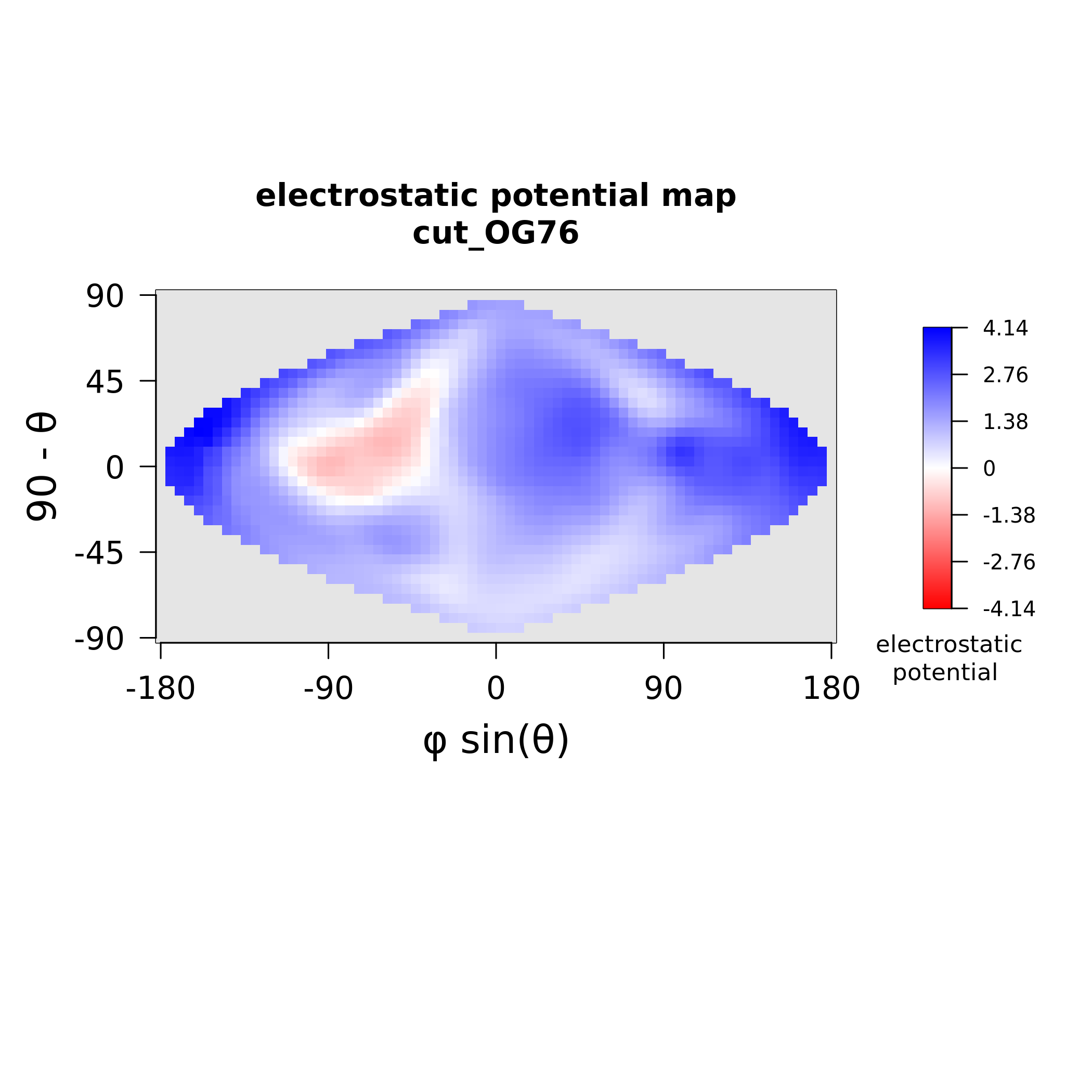

### MAX76_stickiness.png

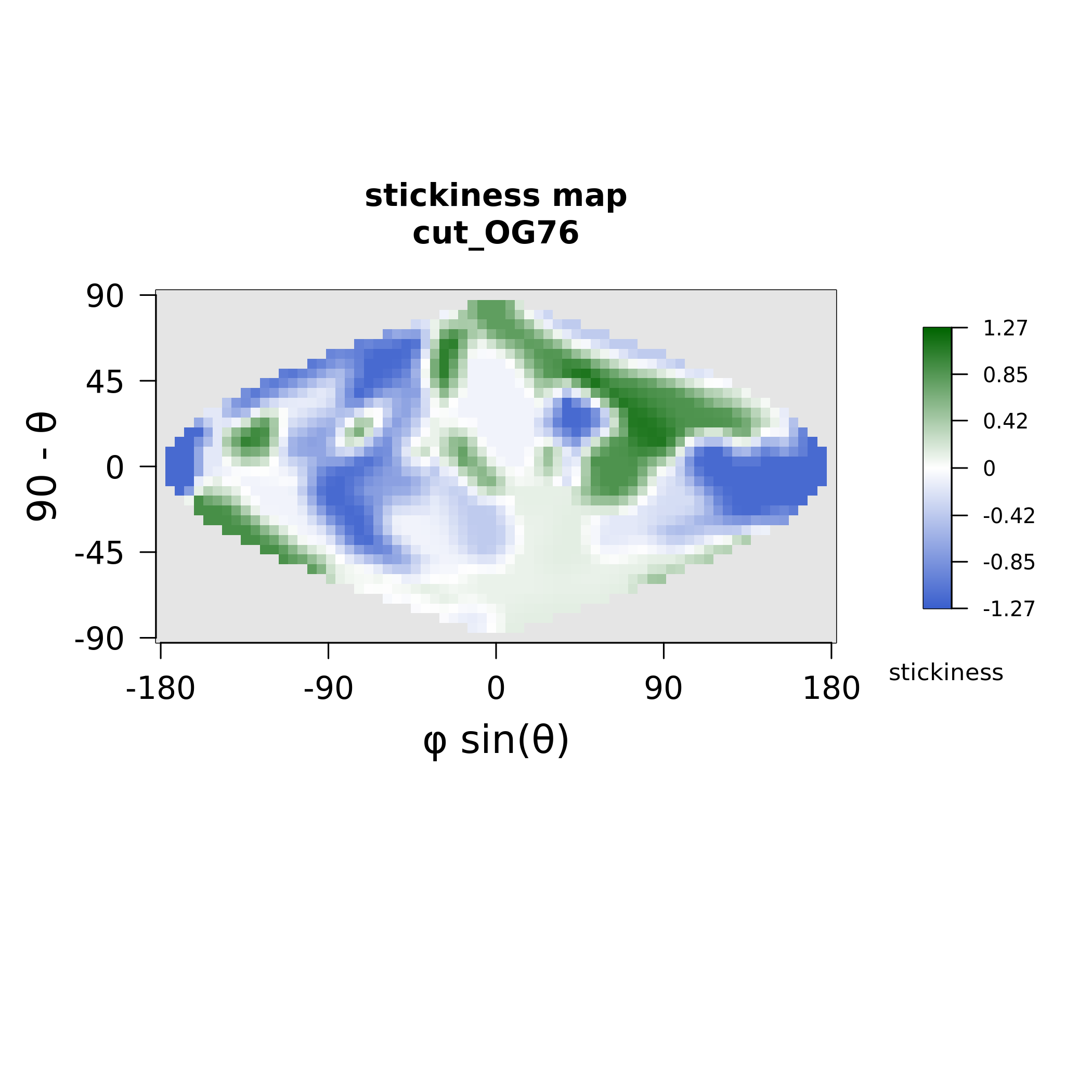

### MAX78_electrostatics.png

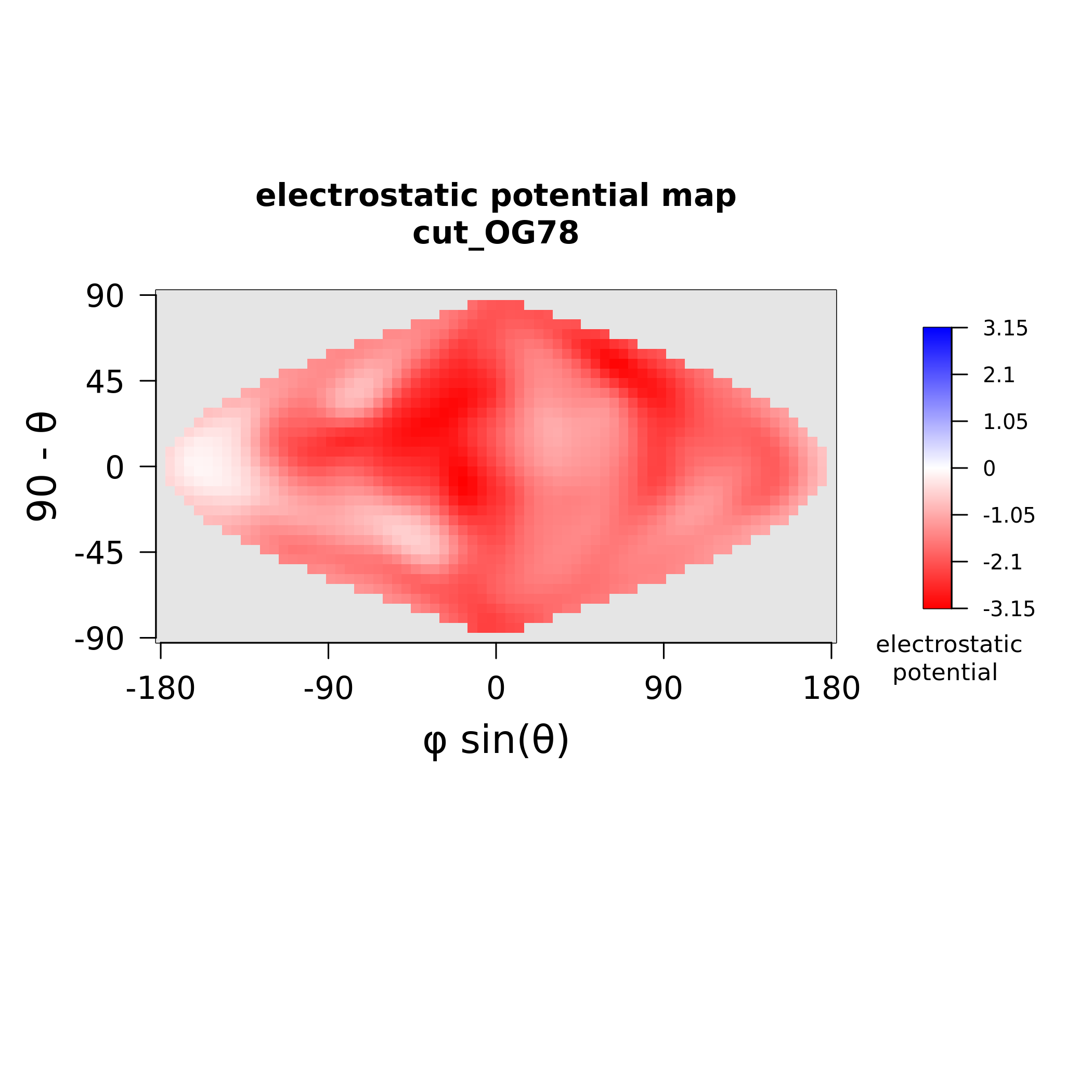

### MAX78_stickiness.png

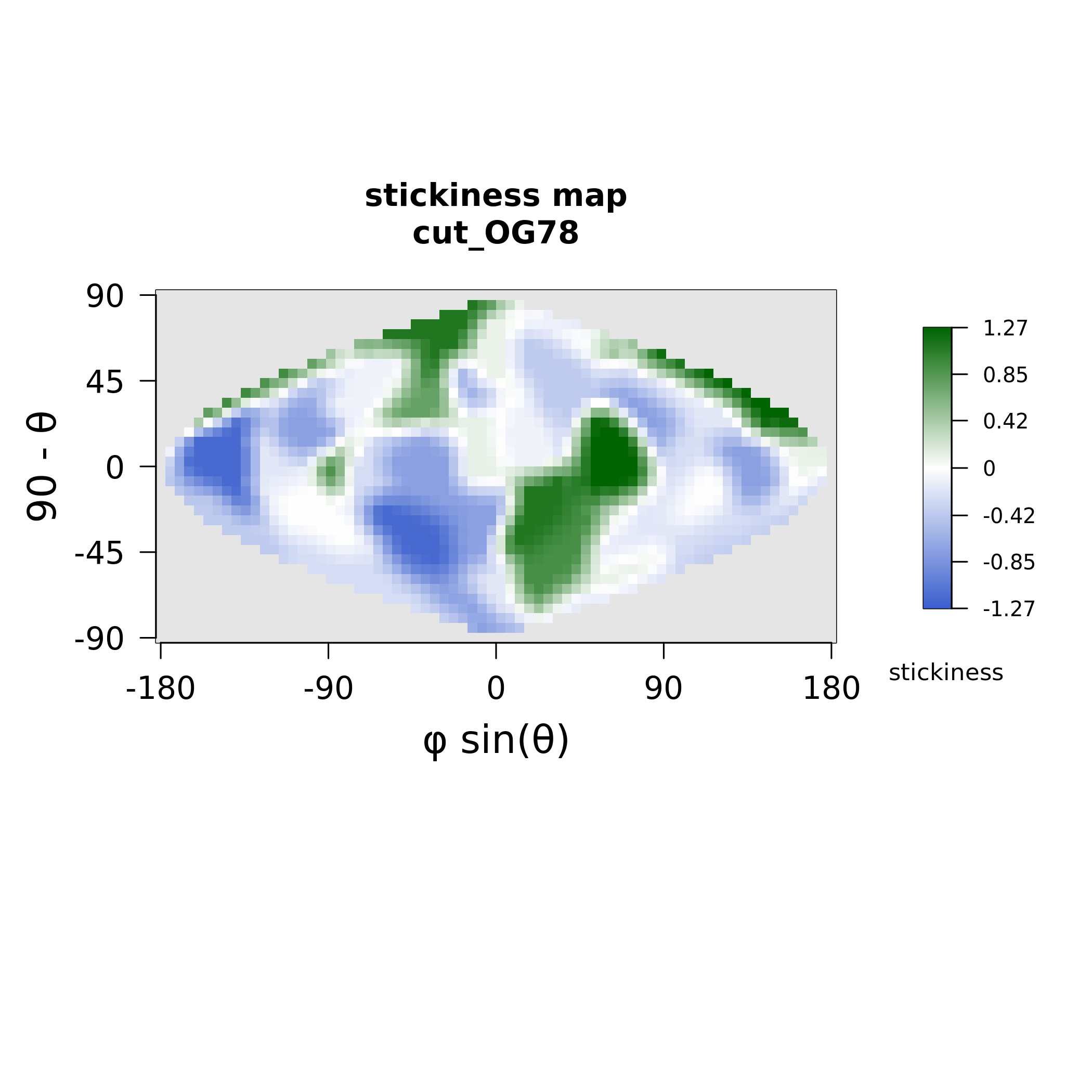

### MAX80_electrostatics.png

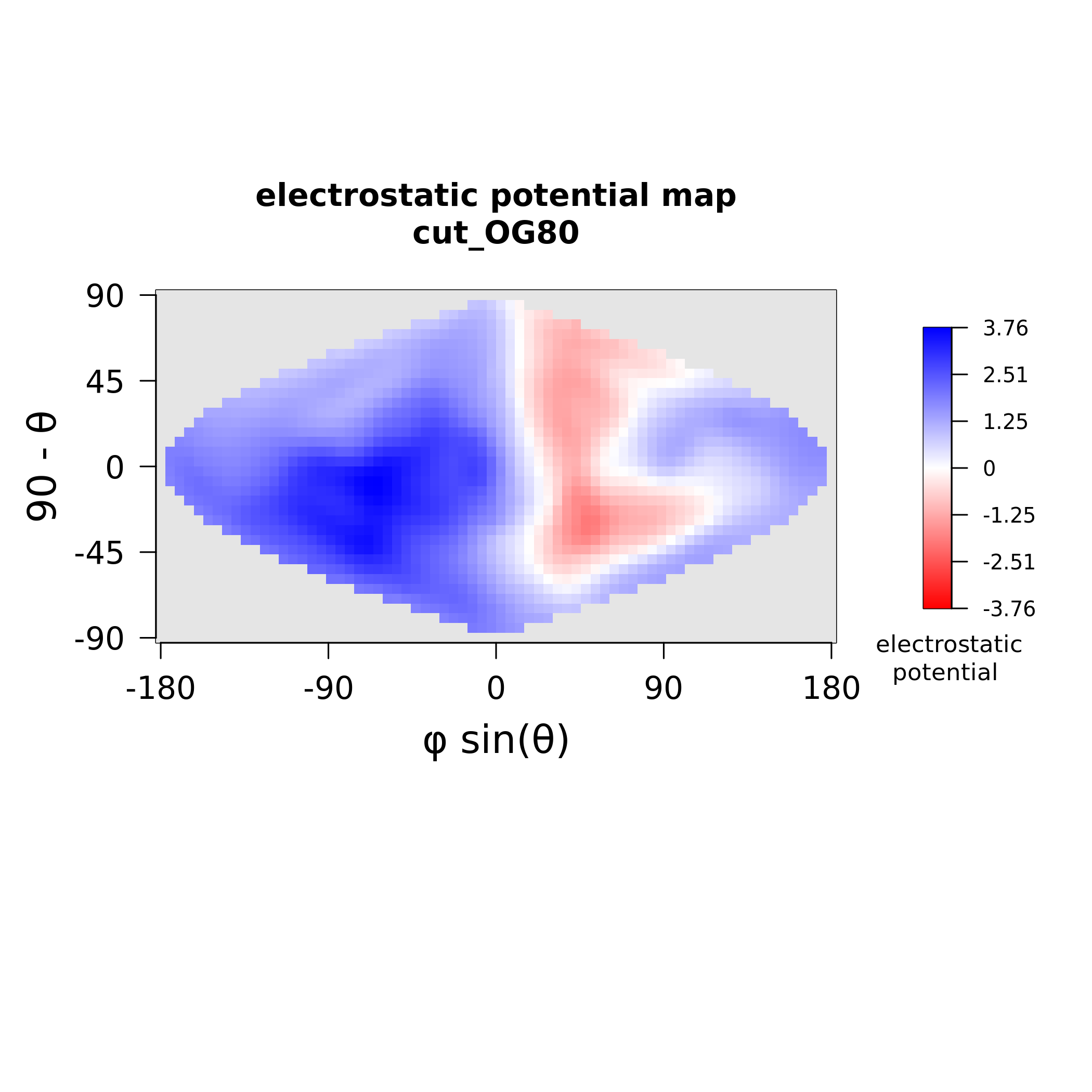

### MAX80_polymorphism.png

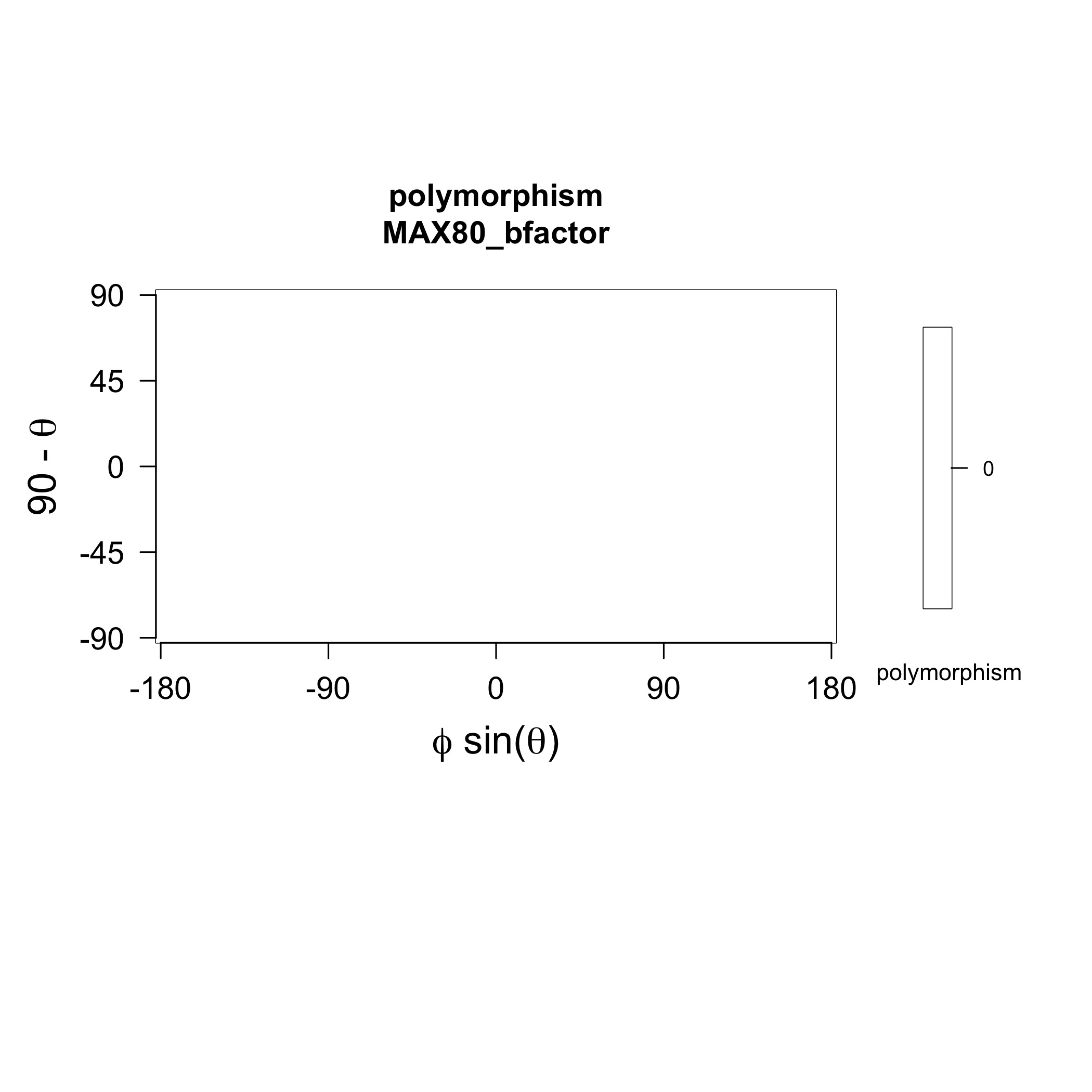

### MAX80_stickiness.png

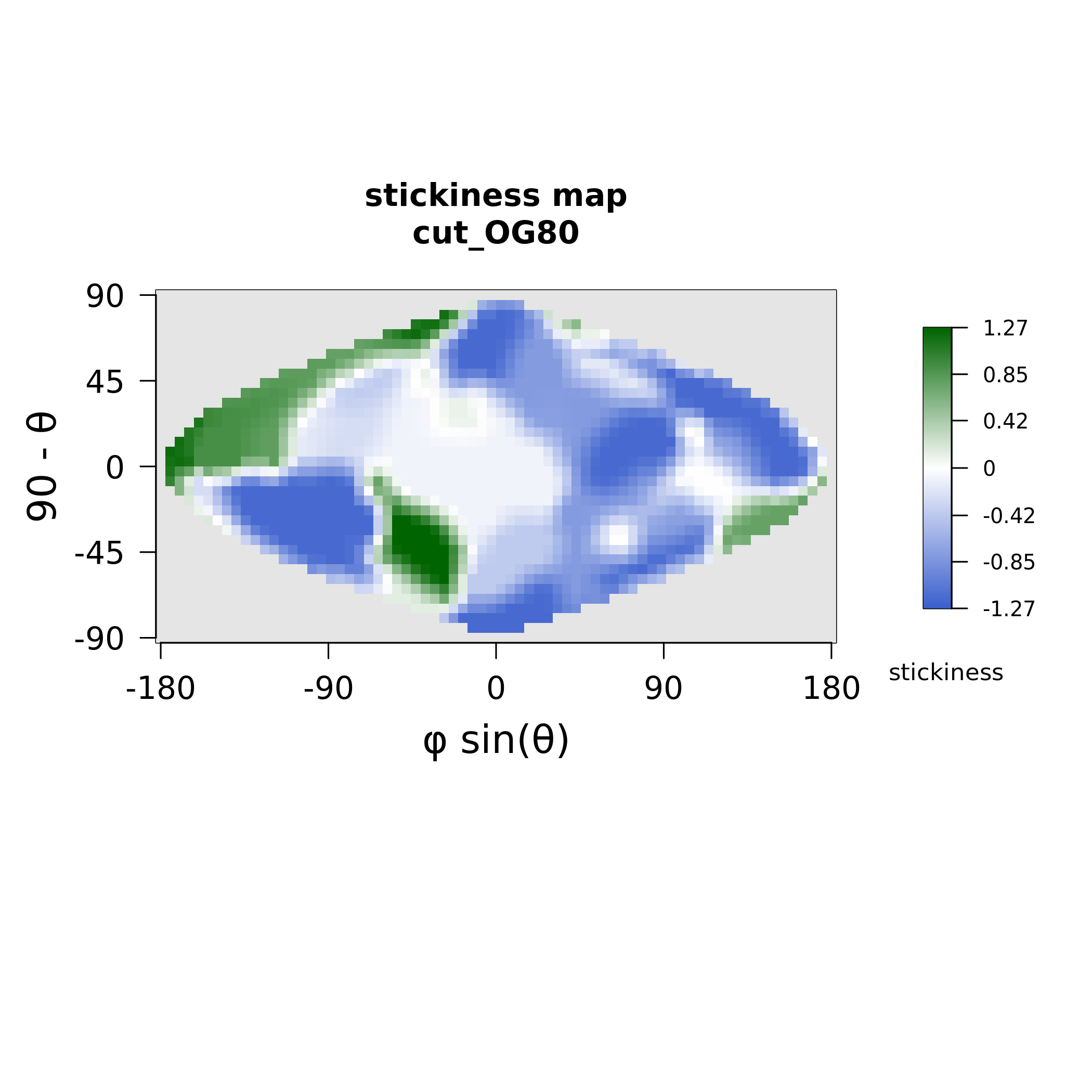

### MAX82_electrostatics.png

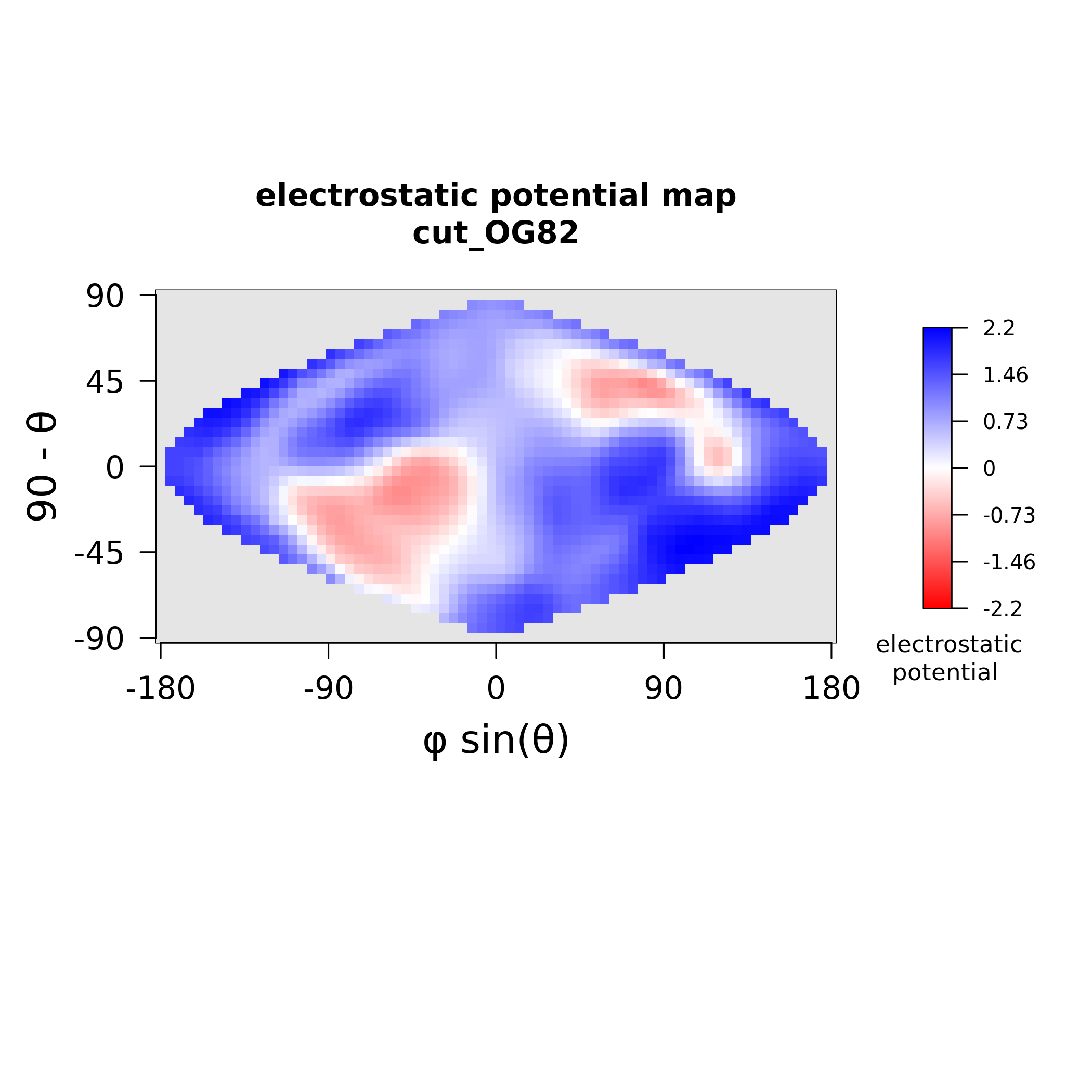

### MAX82_stickiness.png

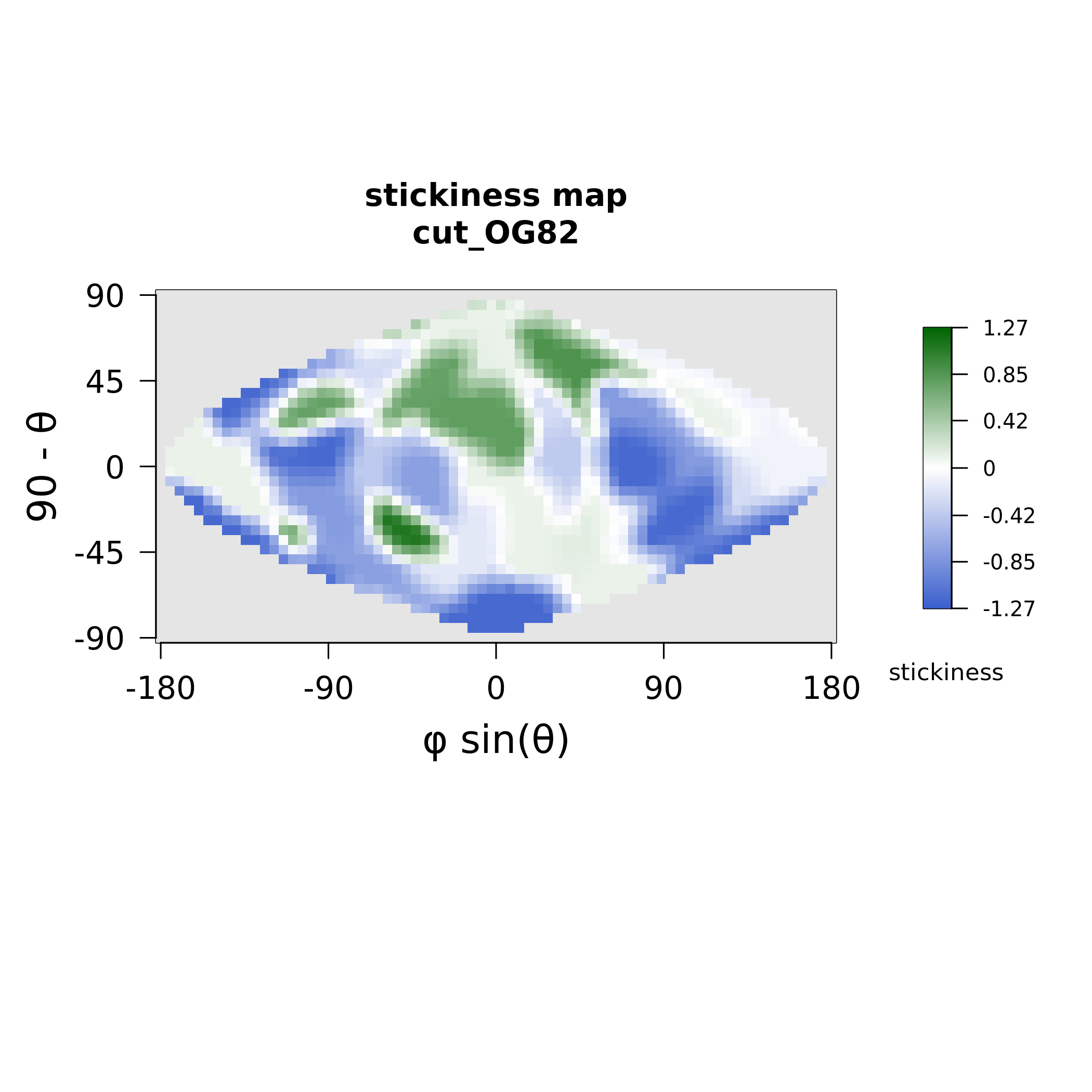

### MAX86_electrostatics.png

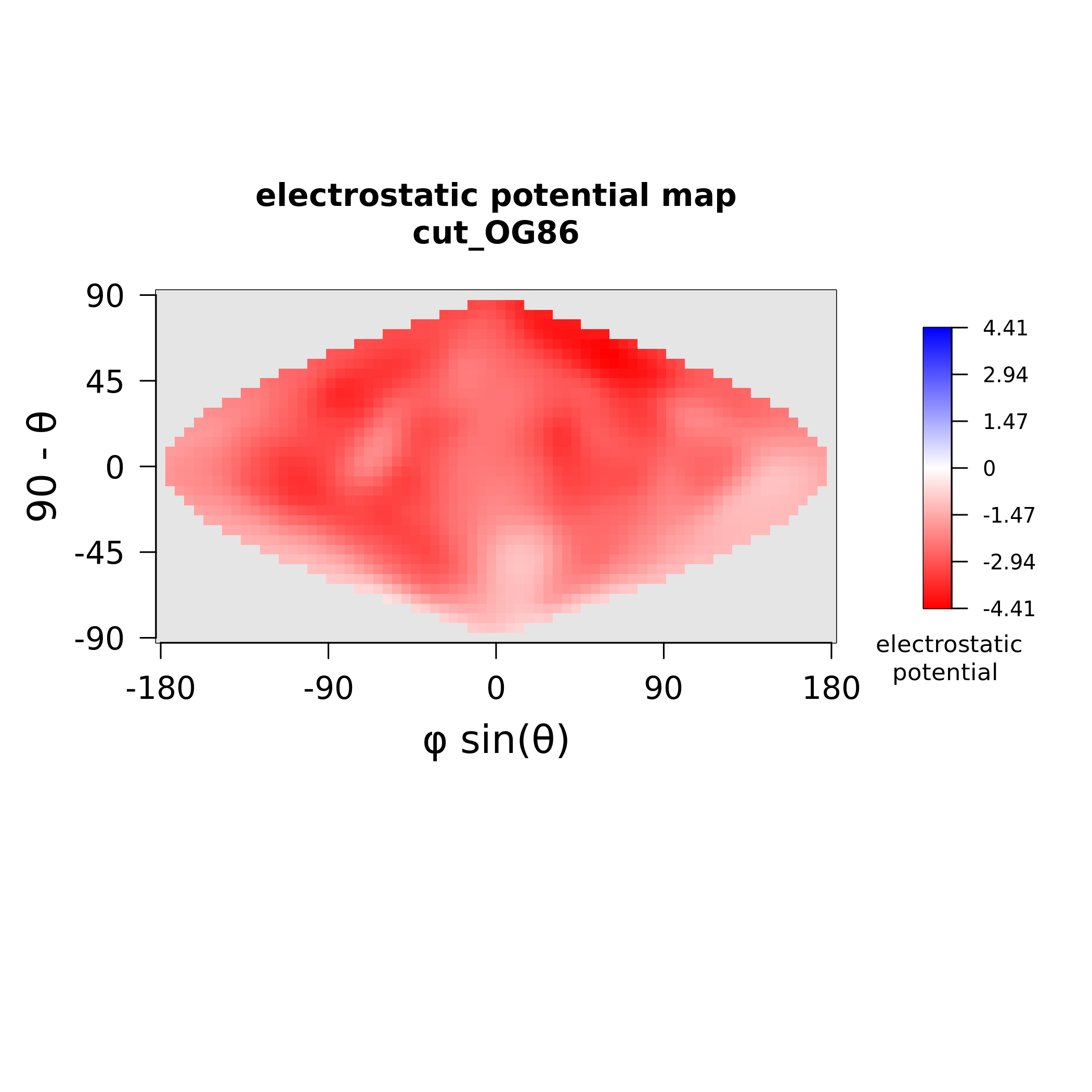

### MAX86_stickiness.png

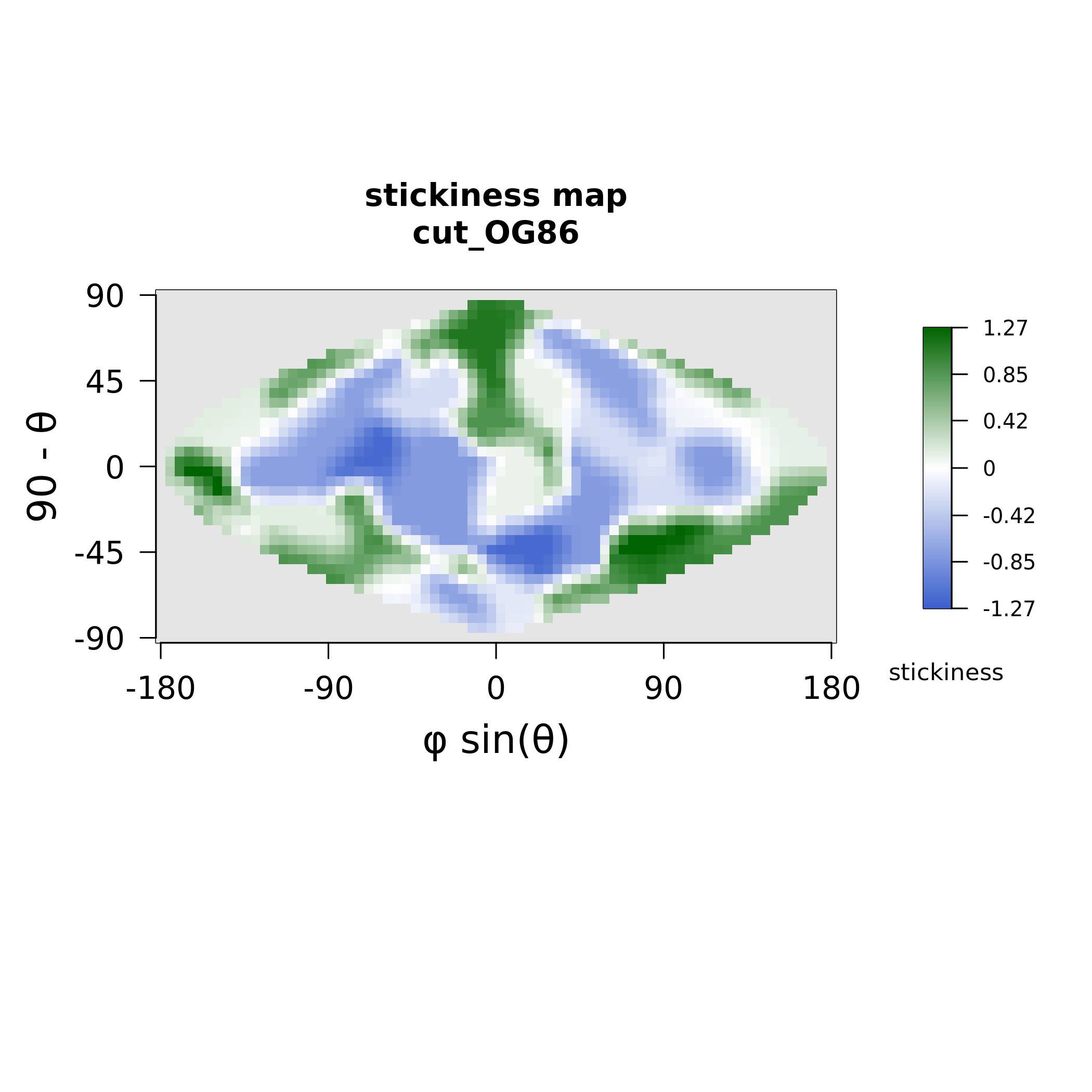

### MAX88_electrostatics.png

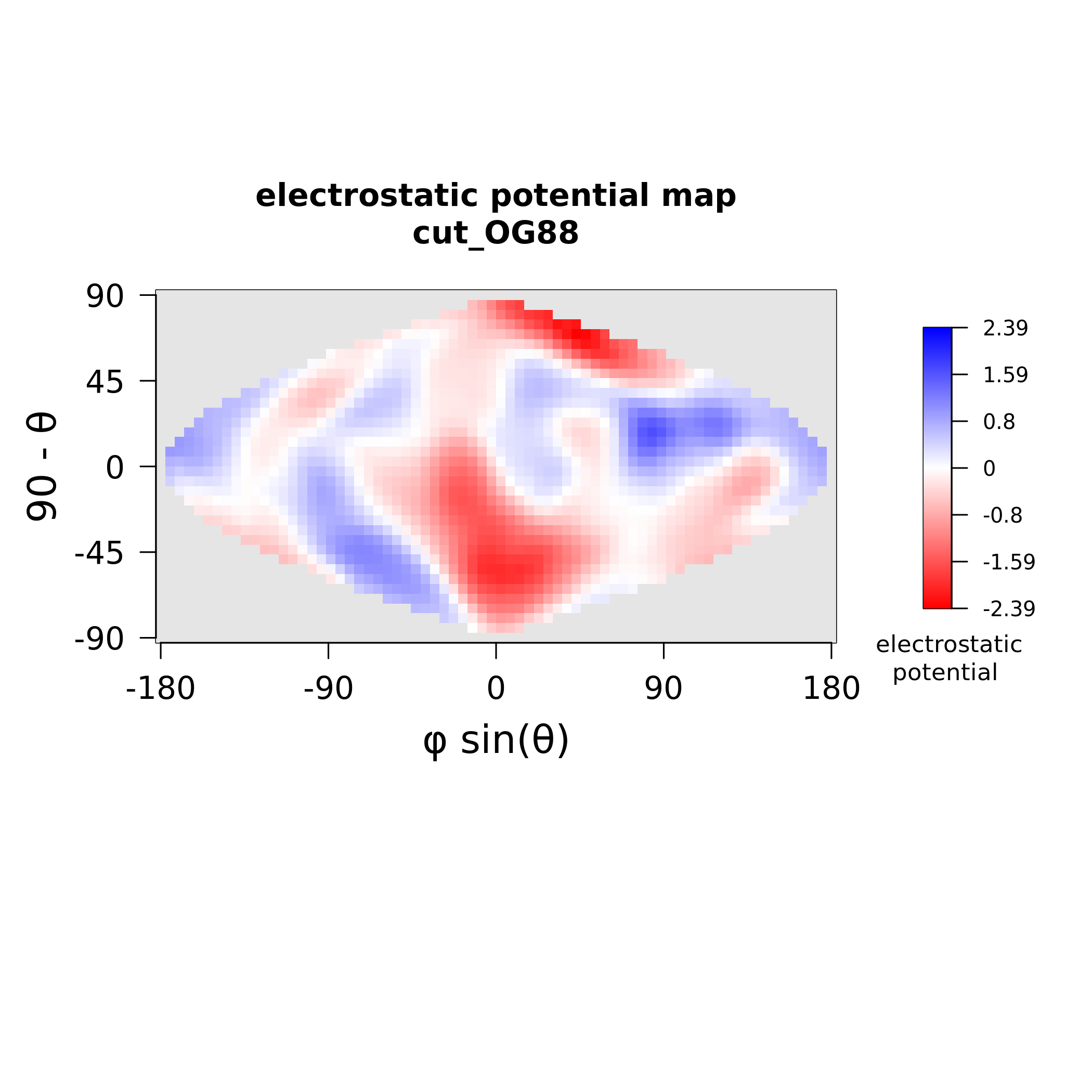

### MAX88_stickiness.png

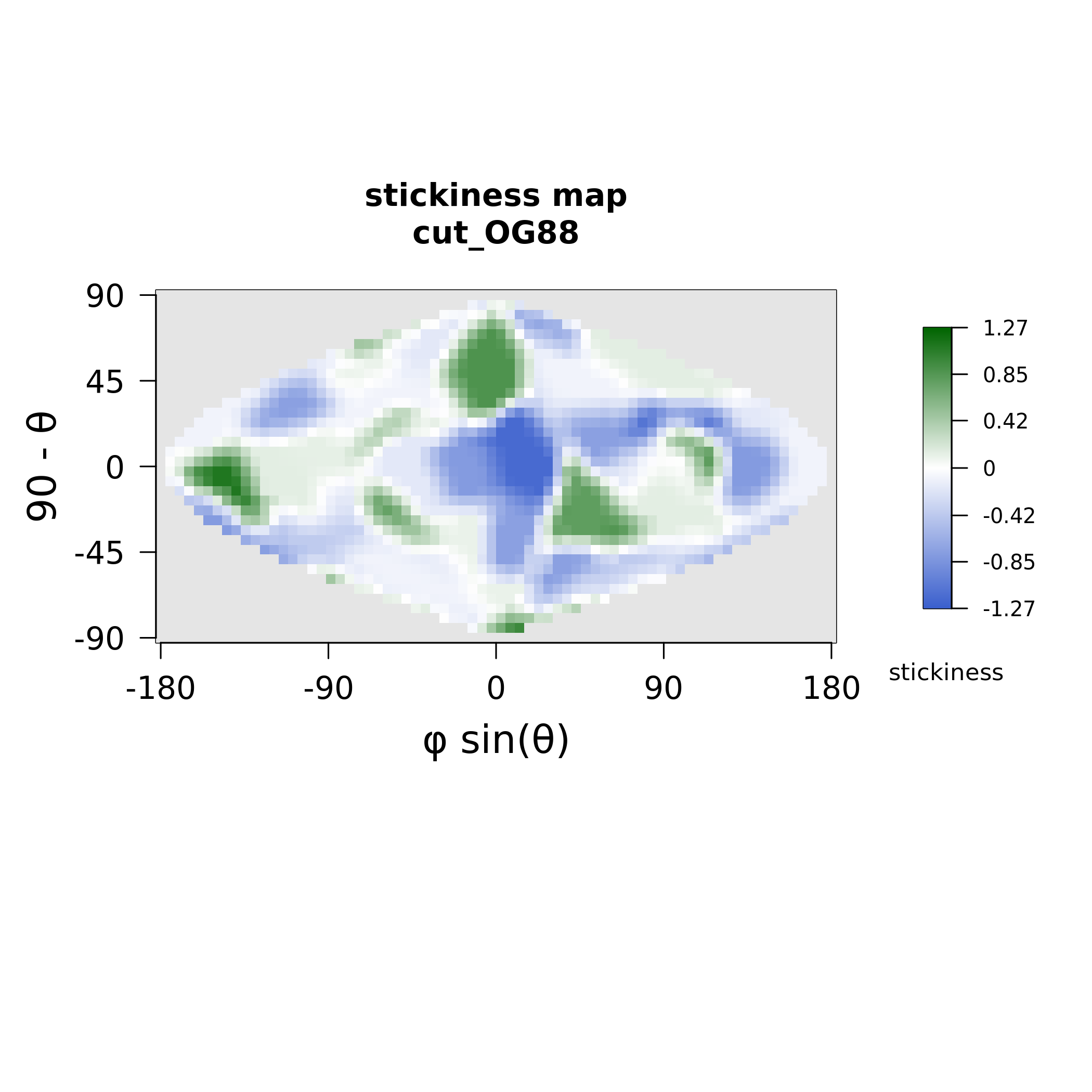

### MAX89_electrostatics.png

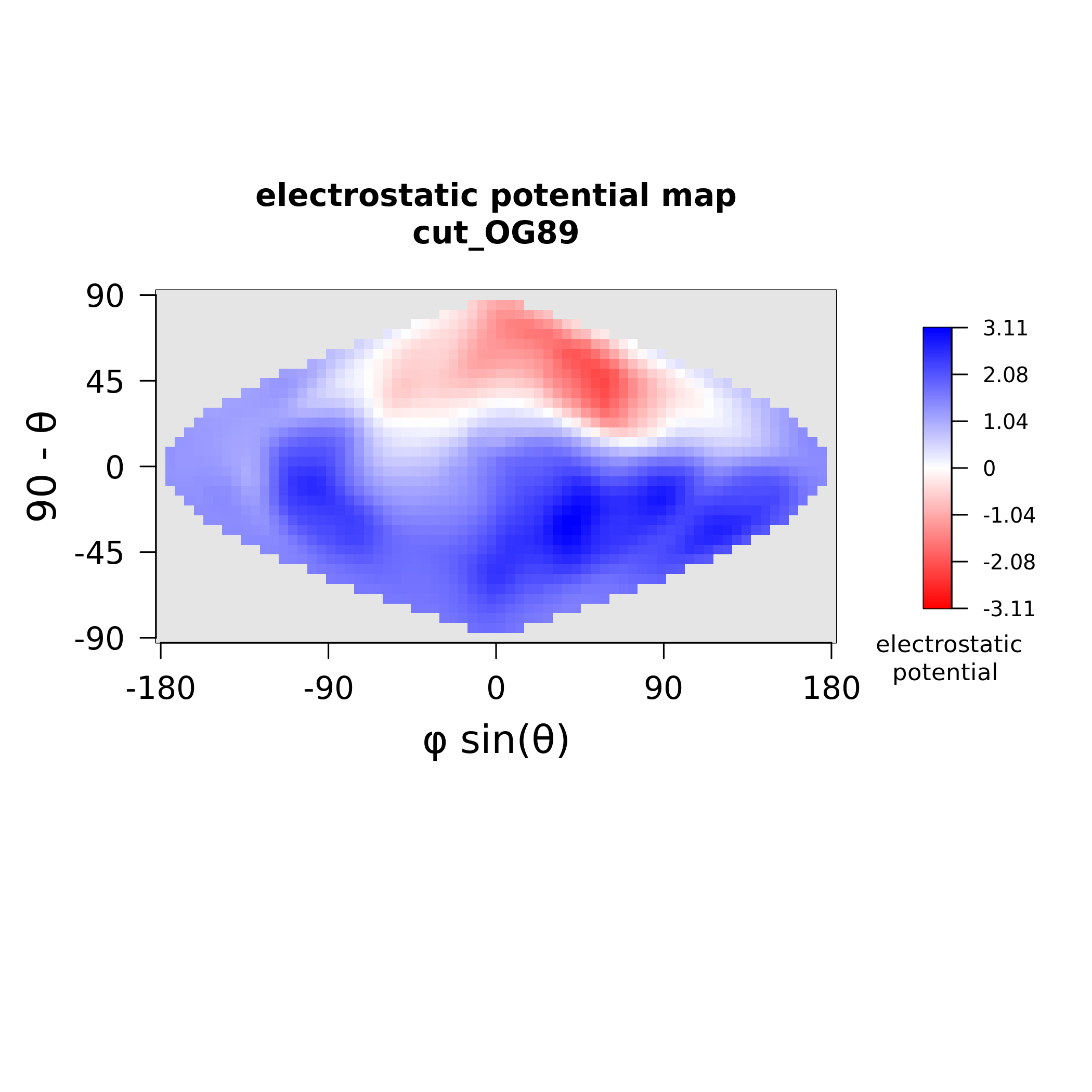

### MAX89_stickiness.png

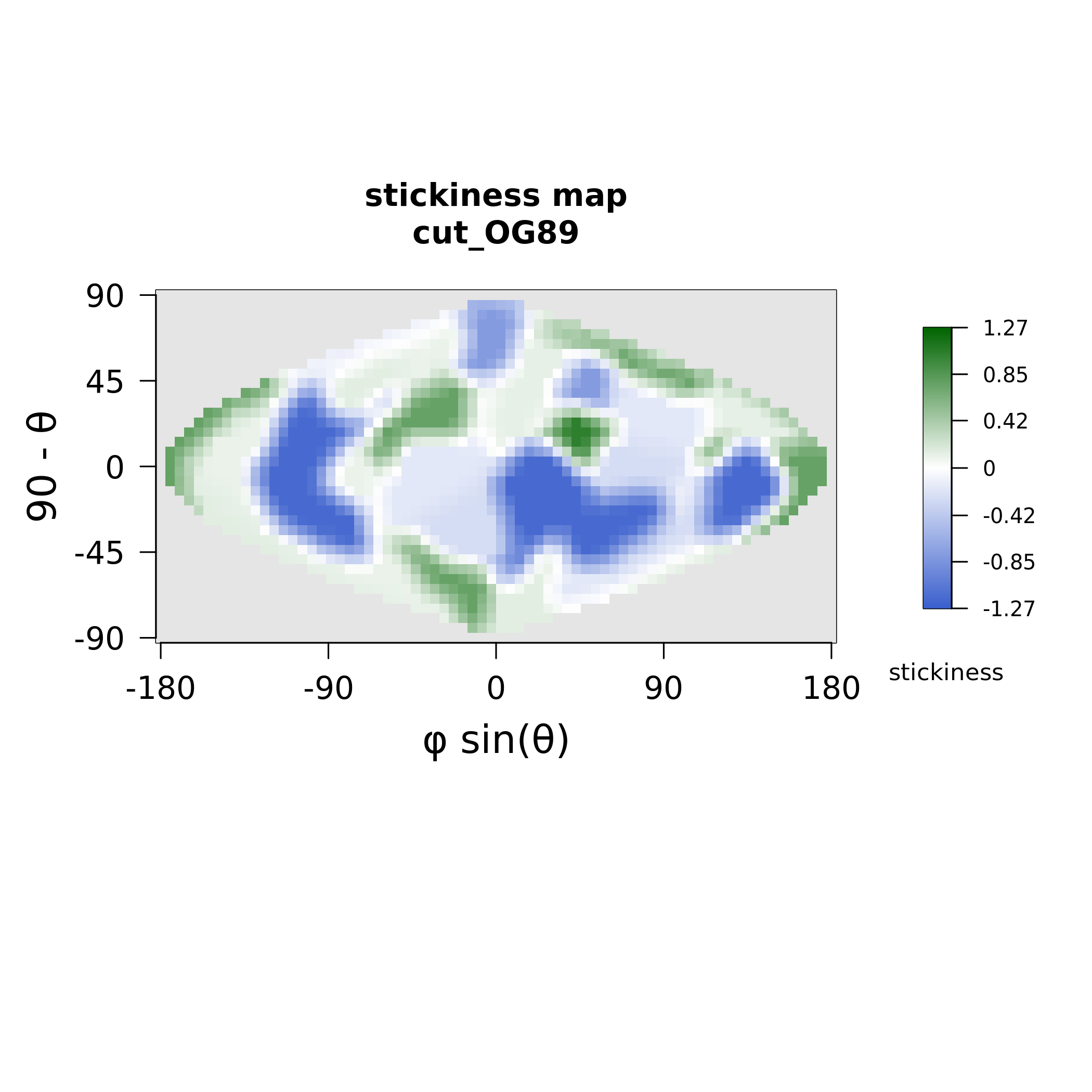

### MAX91_electrostatics.png

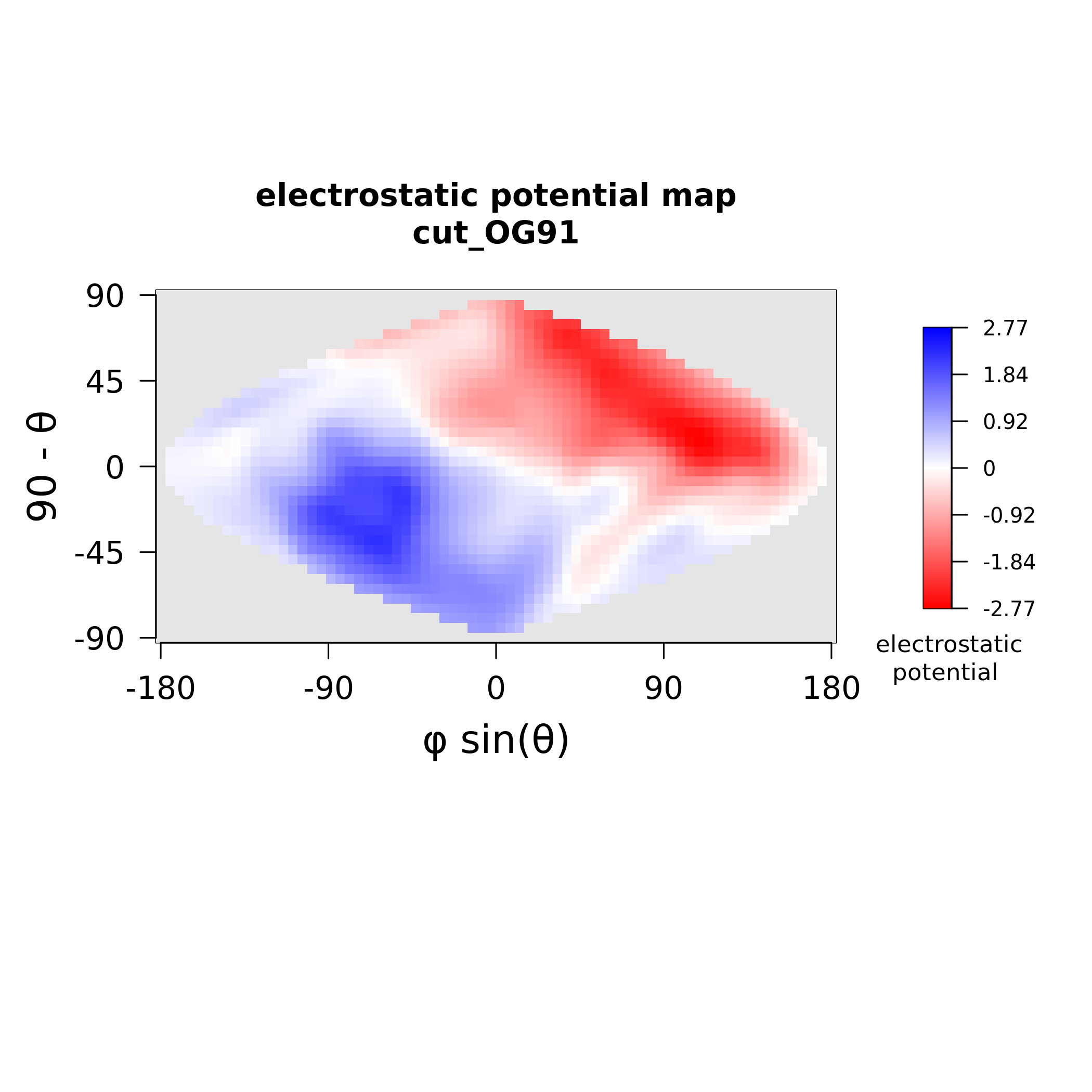

### MAX91_stickiness.png

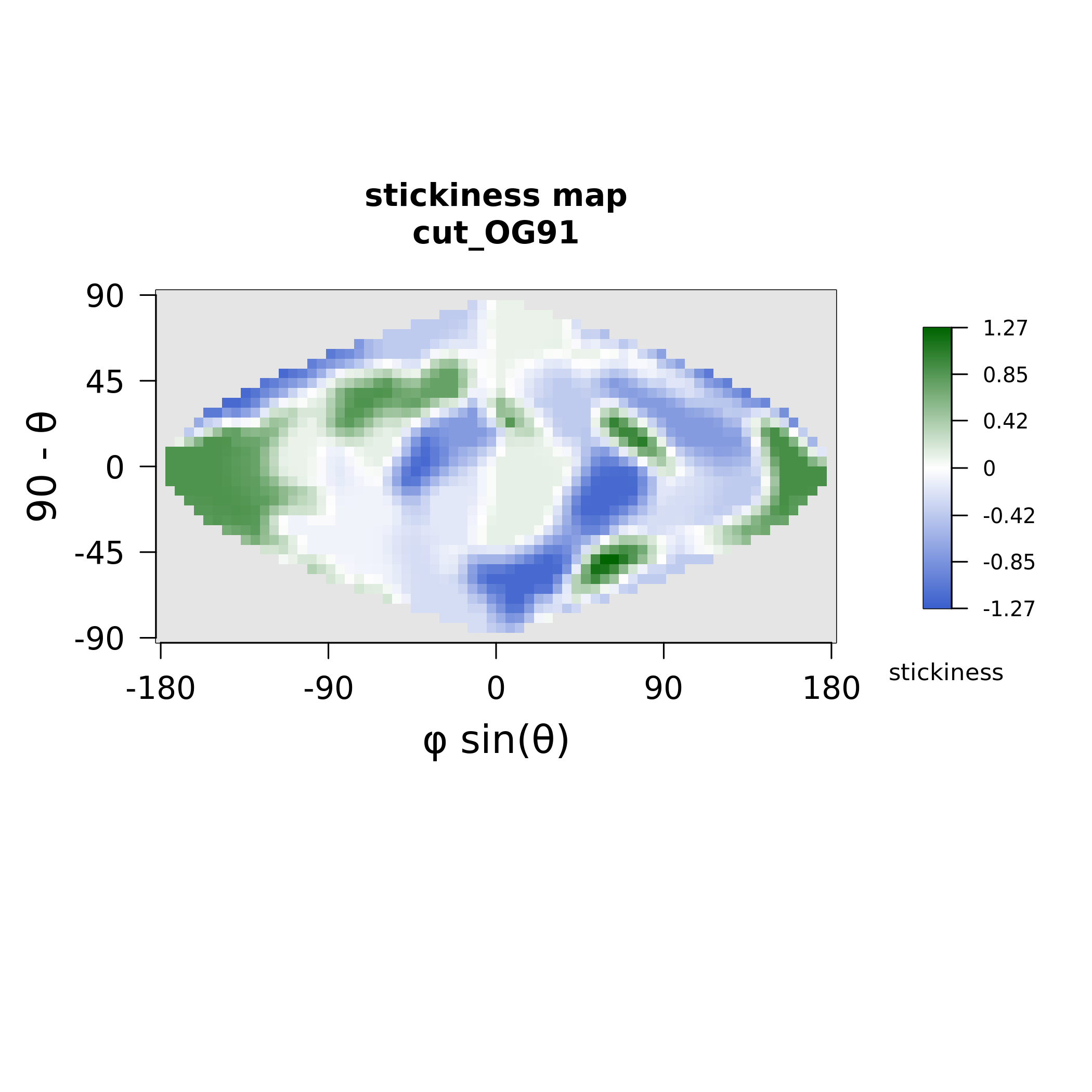

### MAX92_electrostatics.png

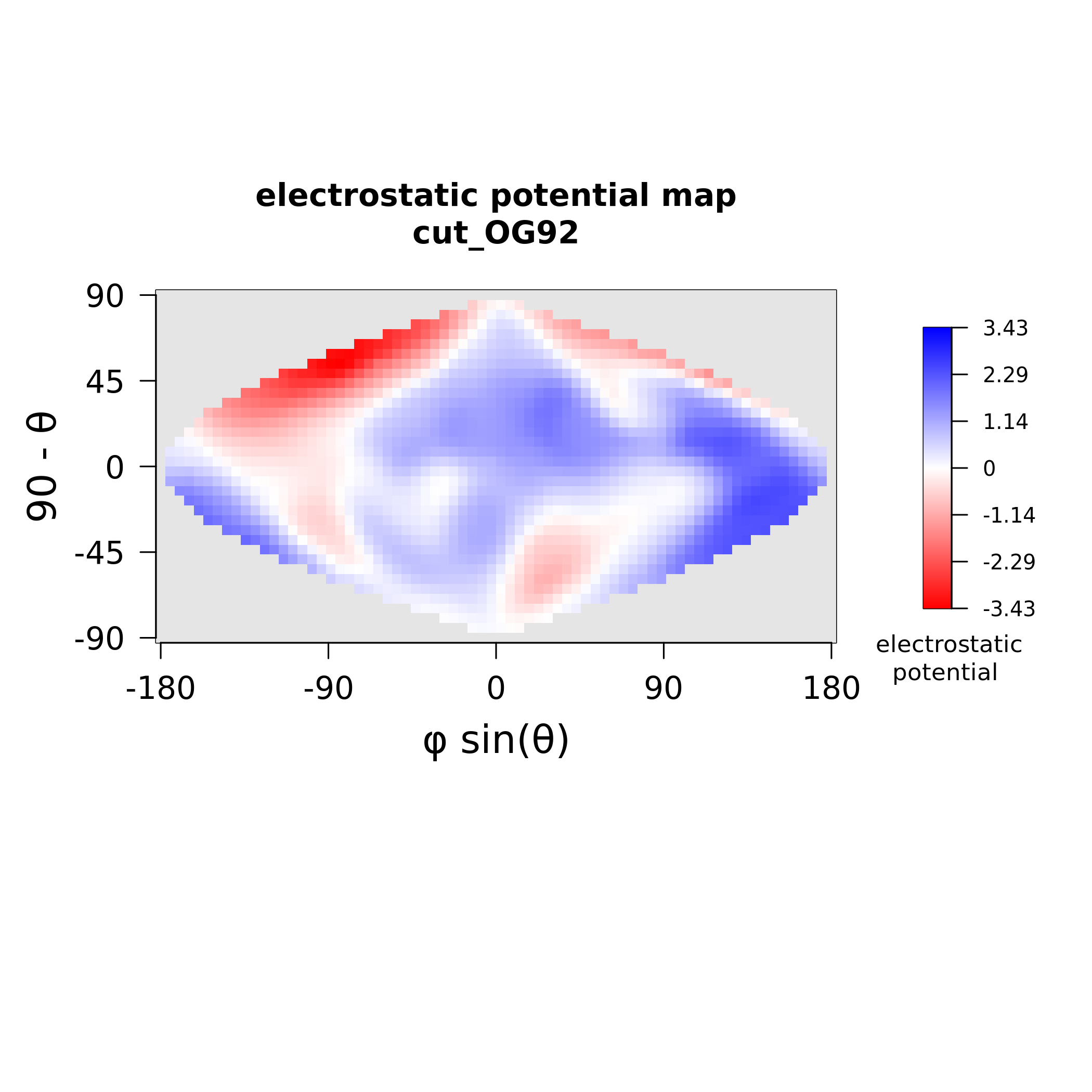

### MAX92_stickiness.png

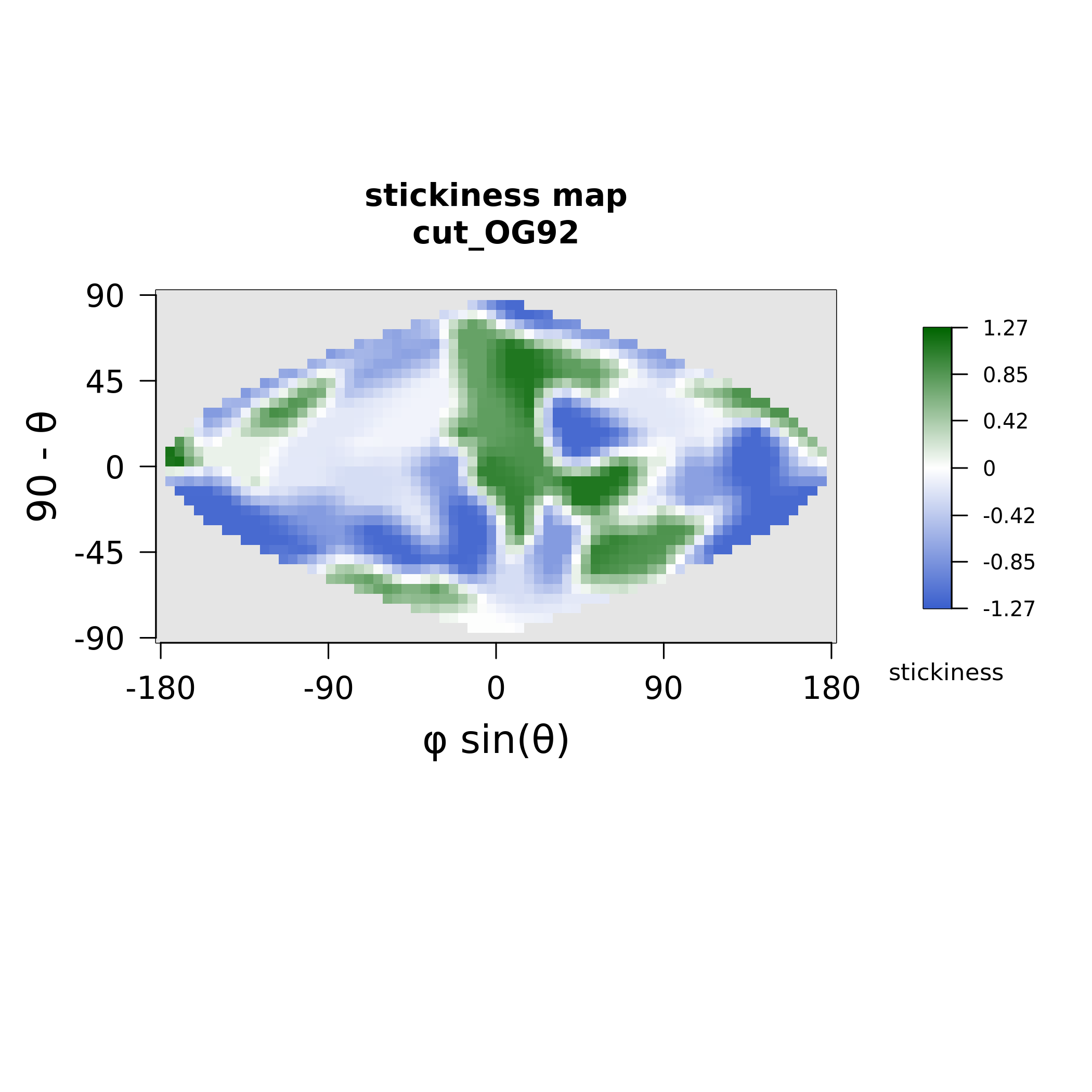

### MAX93_electrostatics.png

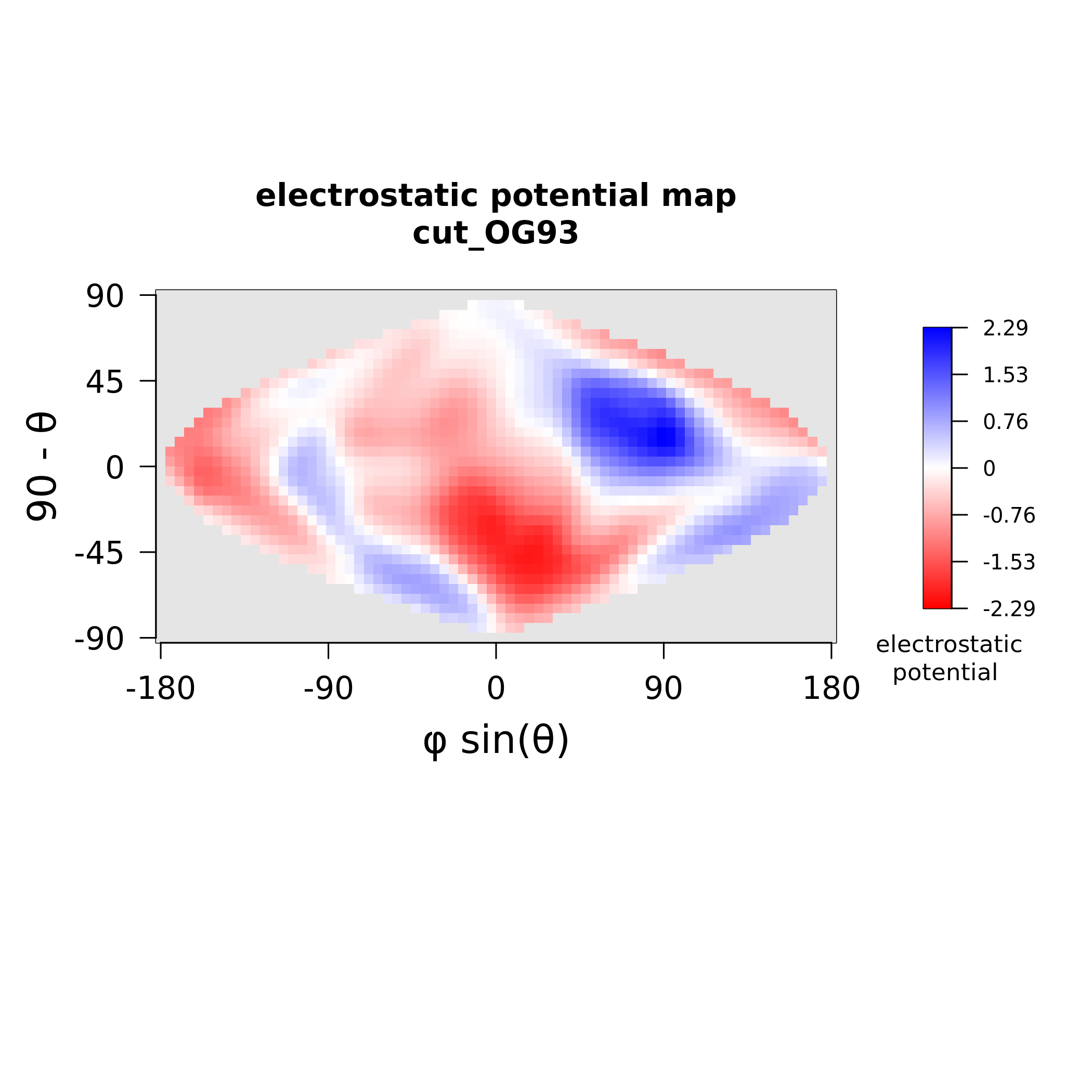

### MAX93_stickiness.png

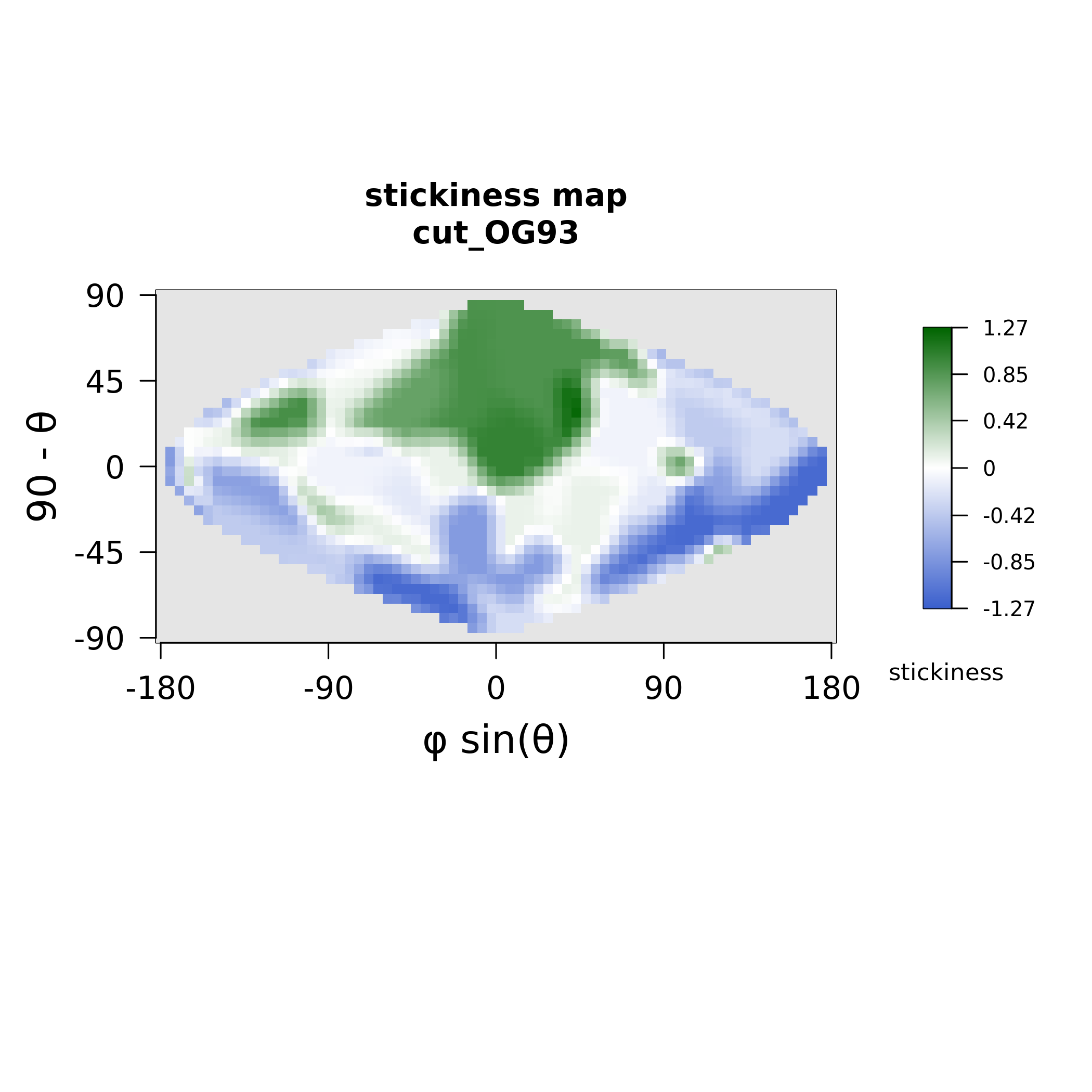
